# Chromosome-level genomes of multicellular algal sisters to land plants illuminate signaling network evolution

**DOI:** 10.1101/2023.01.31.526407

**Authors:** Xuehuan Feng, Jinfang Zheng, Iker Irisarri, Huihui Yu, Bo Zheng, Zahin Ali, Sophie de Vries, Jean Keller, Janine M.R. Fürst-Jansen, Armin Dadras, Jaccoline M.S. Zegers, Tim P. Rieseberg, Amra Dhabalia Ashok, Tatyana Darienko, Maaike J. Bierenbroodspot, Lydia Gramzow, Romy Petroll, Fabian B. Haas, Noe Fernandez-Pozo, Orestis Nousias, Tang Li, Elisabeth Fitzek, W. Scott Grayburn, Nina Rittmeier, Charlotte Permann, Florian Rümpler, John M. Archibald, Günter Theißen, Jeffrey P. Mower, Maike Lorenz, Henrik Buschmann, Klaus von Schwartzenberg, Lori Boston, Richard D. Hayes, Chris Daum, Kerrie Barry, Igor V. Grigoriev, Xiyin Wang, Fay-Wei Li, Stefan A. Rensing, Julius Ben Ari, Noa Keren, Assaf Mosquna, Andreas Holzinger, Pierre-Marc Delaux, Chi Zhang, Jinling Huang, Marek Mutwil, Jan de Vries, Yanbin Yin

## Abstract

The filamentous and unicellular algae of the class Zygnematophyceae are the closest algal relatives of land plants. Inferring the properties of the last common ancestor shared by these algae and land plants allows us to identify decisive traits that enabled the conquest of land by plants. We sequenced four genomes of filamentous Zygnematophyceae (three strains of *Zygnema circumcarinatum* and one strain of *Z. cylindricum*) and generated chromosome-scale assemblies for all strains of the emerging model system *Z. circumcarinatum*. Comparative genomic analyses reveal expanded genes for signaling cascades, environmental response, and intracellular trafficking that we associate with multicellularity. Gene family analyses suggest that Zygnematophyceae share all the major enzymes with land plants for cell wall polysaccharide synthesis, degradation, and modifications; most of the enzymes for cell wall innovations, especially for polysaccharide backbone synthesis, were gained more than 700 million years ago. In Zygnematophyceae, these enzyme families expanded, forming co-expressed modules. Transcriptomic profiling of over 19 growth conditions combined with co-expression network analyses uncover cohorts of genes that unite environmental signaling with multicellular developmental programs. Our data shed light on a molecular chassis that balances environmental response and growth modulation across more than 600 million years of streptophyte evolution.

**HIGHLIGHTS:**

1. Genomes of four filamentous algae (*Zygnema*) sisters to land plants
2. *Zygnema* are rich in genes for multicellular growth and environmental acclimation: signaling, lipid modification, and transport
3. Cell wall innovations: diversification of hexameric rosette cellulose synthase in Zygnematophyceae
4. Co-expression networks reveal conserved modules for balancing growth and acclimation

## INTRODUCTION

Plant terrestrialization changed the surface of the Earth. In a fateful event about 550 million years ago, the first representatives of the clade Embryophyta (land plants) gained a foothold on land (Delwiche and Cooper, 2015; Morris et al., 2018; Fürst-Jansen et al., 2020; Strother and Foster, 2021; Harris et al., 2022). These first land plants emerged from within the clade of Streptophyta that—until Embryophyta emerged—consisted solely of streptophyte algae. Which lineage among these freshwater and terrestrial algae is closest to land plants? Over the last decade, considerable advances have been made in establishing a robust phylogenomic framework for streptophyte evolution and the birth of embryophytes (Wodniok et al., 2011; Wickett et al., 2014; Puttick et al., 2018; Leebens-Mack et al., 2019). Moreover, a long tradition of cell biological research in streptophyte algae is available (reviewed by Domozych and Bagdan 2022). Both lines of evidence have come to the same conclusion: Zygnematophyceae are the closest algal relatives of land plants.

Zygnematophyceae is a class of freshwater and semi-terrestrial algae with more than 4,000 described species (Guiry 2021). Their unifying feature is sexual reproduction by conjugation and lack of motile stages; zygospore formation can often be observed in field samples e.g., in *Mougeotia* (Permann et al., 2021a) and *Spirogyra* (Permann et al., 2021b, 2022a). Zygnematophyceae have been recently rearranged into five orders: Spirogloeales, Serritaeniales, Spirogyrales, Desmidiales, and Zygnematales (Hess et al., 2022). Among these, filamentous growth likely evolved at least five times independently (Hess et al., 2022). So far, genome sequences are only available for unicellular Zygnematophyceae: *Penium margariatceum* and *Closterium peracerosum–strigosum–littorale* in Desmidiales (Jiao et al. 2020; Sekimoto et al., 2023), *Mesotaenium endlicherianum* in Serritaeniales, and *Spirogloea muscicola* in Spiroglocales (both Cheng et al., 2019).

Zygnematophyceae possess adaptations to withstand terrestrial stressors. Some are protected against desiccation by extracellular polymers like arabinogalactan proteins (AGPs; Palacio-Lopez et al. 2019) and homogalacturonan-rich mucilage sheaths (Herburger et al., 2019). UV-absorbing phenolic compounds were shown to occur in *Zygnema* (Pichrtová et al., 2013, Holzinger et al., 2018), *Zygogonium erictorum* (e.g.., Aigner et al., 2013, Herburger et al., 2016) and *Serritaenia* (Busch & Hess 2021). Recurrent transcriptomic and metabolomic changes have been observed following desiccation (Rippin et al. 2017) and other abiotic stresses (de Vries et al., 2018, Arc et al., 2020; Fitzek et al., 2019) in *Zygnema*, as well as under natural conditions in the uppermost layers of Arctic *Zygnema* mats (Rippin et al. 2019). The nature of these stress responses is of deep biological significance: various orthologous groups of proteins once considered specific to land plants have recently been inferred to predate the origin of Embryophyta (Nishiyama et al., 2018; Bowles et al., 2020). These include intricate transcription factor (TF) networks (Wilhelmsson et al., 2017), phytohormone signaling pathways (Bowman et al., 2019), specialized biochemical pathways (Rieseberg et al., 2022), symbiosis signaling (Delaux et al., 2015), and cell wall modifications (Harholt et al., 2016).

Accurately inferring the developmental and physiological programs of the first land plant ancestor depends on our ability to predict them in its sister group, the common ancestor of Zygnematophyceae. Robustness of this inference lays on accounting for the phylogenetic, genomic, and morphologic diversity of the Zygnematophyceae. Here, we report on the first four genomes of filamentous Zygnematophyceae (Order Zygnematales), three from strains of *Zygnema circumcarinatum* and one from *Zygnema* cf. *cylindricum*, which include the first chromosome-scale genomes for any streptophyte algae. Comparative genomics using the new *Zygnema* genomes allow inferring the genetic repertoire of land plants ancestors that first conquered the terrestrial environment. But which functional cohorts were relevant? Co-expression analyses shed light on the deep evolutionary roots of the mechanism for balancing environmental responses and multicellular growth.

## RESULTS AND DISCUSSION

### First chromosome-level genomes for streptophyte algae

The nuclear and organellar genomes of four *Zygnema* strains (*Z. circumcarinatum* SAG 698-1b, UTEX 1559, and UTEX 1560 and *Zygnema* cf. *cylindricum* SAG 698-1a, **Figure 1A-C**) were assembled using a combination of PacBio High-Fidelity (HiFi) long reads, Oxford Nanopore long reads, and Illumina short reads. In total, we sequenced 51 gigabases (Gb) (797X), 69 Gb (1042X), 6.7 Gb (103X), and 253 Gb (786X) for strains SAG 698-1b, UTEX 1559, UTEX 1560, and SAG 698-1a, respectively (**Table S1A,B**). Using chromatin conformation data (Dovetail Hi-C), we scaffolded the *Z. circumcarinatum* SAG 698-1b genome (N50=4Mb; **Table 1**) into 90 final scaffolds; 98.6% of the assembly belongs to the 20 longest scaffolds (**Table S1C**) corresponding to 20 pseudo-chromosomes (**Figure 1D**). Cytological chromosome counting at different stages of mitosis (prophase, metaphase, telophase) (**Figure 1B, Figure S1**) verified the 20 chromosomes, similar to a previous cytological study of another *Zygnema* strain that found 19 chromosomes (Prasad & Godward 1966). The total assembly size (71 megabases (Mb)) was close to genome sizes estimated by flow cytometry, fluorescence staining (Feng et al., 2021), and k-mer frequency analysis (**Figure S2, Table S1B**). The high mapping rates of UTEX 1559 and UTEX 1560 Illumina reads to the SAG 698-1b genome (97.16% and 97.12%, respectively) confirm that these genomes are from close relatives, in agreement with the strains’ history (see **Supplemental Text 1**). We thus used chromatin conformation data from SAG 698-1b to scaffold UTEX 1559 and UTEX 1560 assemblies, which resulted in 20 chromosomes containing 97.3% and 98.3% of the total assemblies, respectively. The three new *Zygnema circumcarinatum* genomes represent the first chromosome-level assemblies for any streptophyte alga (**Table 1**).

**Figure 1:**
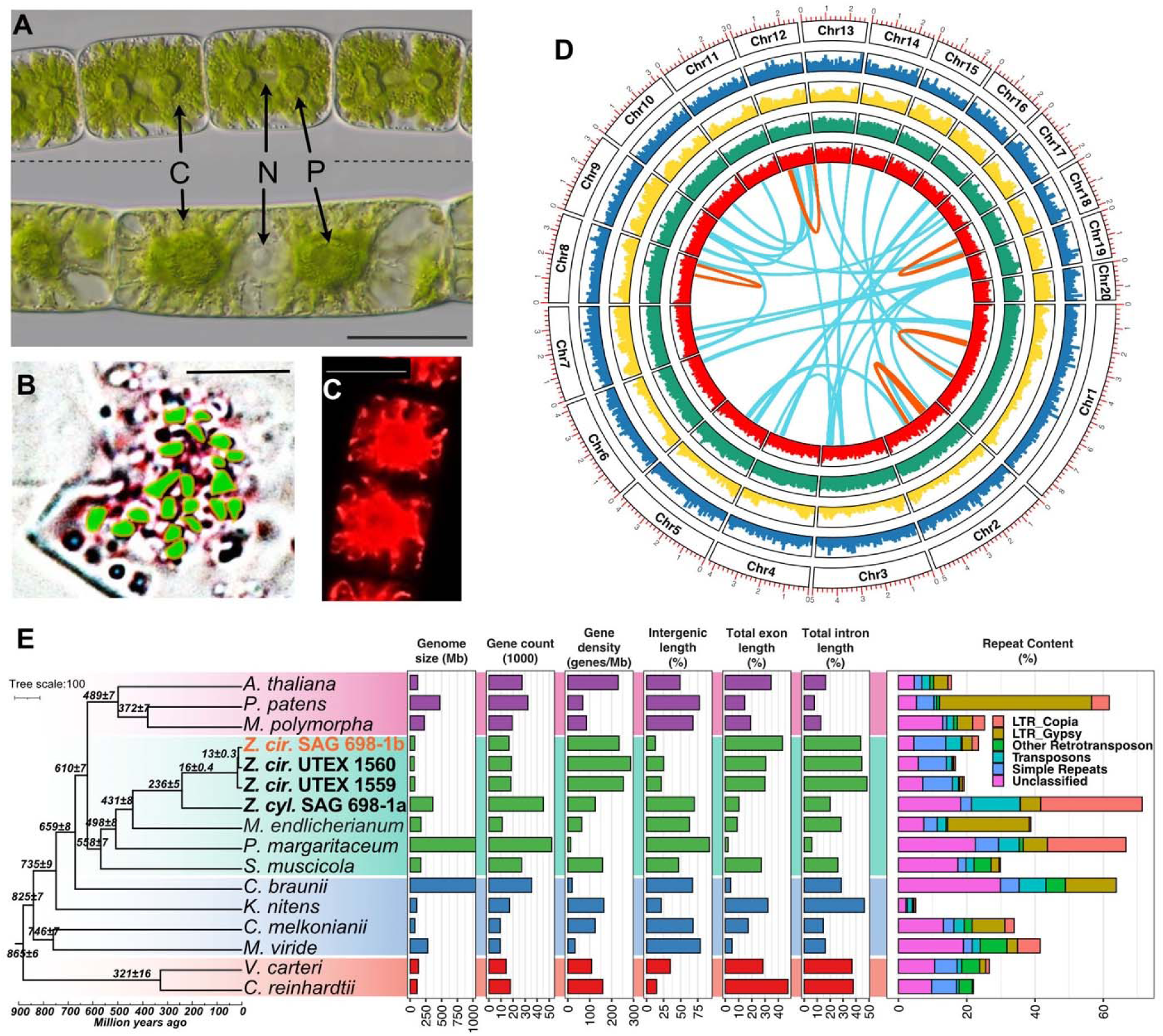
(A) Three cells of a vegetative filament of SAG 698-1b (top) compared to one cell of a vegetative filament of SAG 698-1a (bottom, both samples 1 month old). Scale bar: 20 μm; C chloroplast; N nucleus; P pyrenoid. One-cell filament contains two chloroplasts and one nucleus. (B) Chromosome counting on light micrographs of SAG 698-1b fixed and stained with acetocarmine at prophase (0.5 months old); count was also performed in metaphase and telophase. Green dots represent the 20 chromosomes which were counted after rendering a stack of ∼ 100 images (scale bar: 10 μm); see Figure S1 for the original images. (C) Confocal laser scanning image of one SAG 698-1b cell (0.5 month). Scale bar: 20 μm. (D) Chromosome-level assembly of SAG 698-1b genome. Concentric rings show chromosome (Chr) numbers, gene density (blue), repeat density (yellow), RNA-seq mapping density (log10(FPKM) (dark green), and GC% density (red). Red and green links show respectively intra- and inter-chromosomal syntenic blocks. (E) Comparison of genome properties for 13 algal and 3 land plant species. The time-calibrated species tree was built from 493 low-copy genes (all nodes supported by >97% non-parametric bootstrap; numbers at nodes are estimated divergence times (mean ± standard deviation) (see **Table S1F** for details). Data for bar plot can be found in **Table S1I,J.**

**Table 1:**
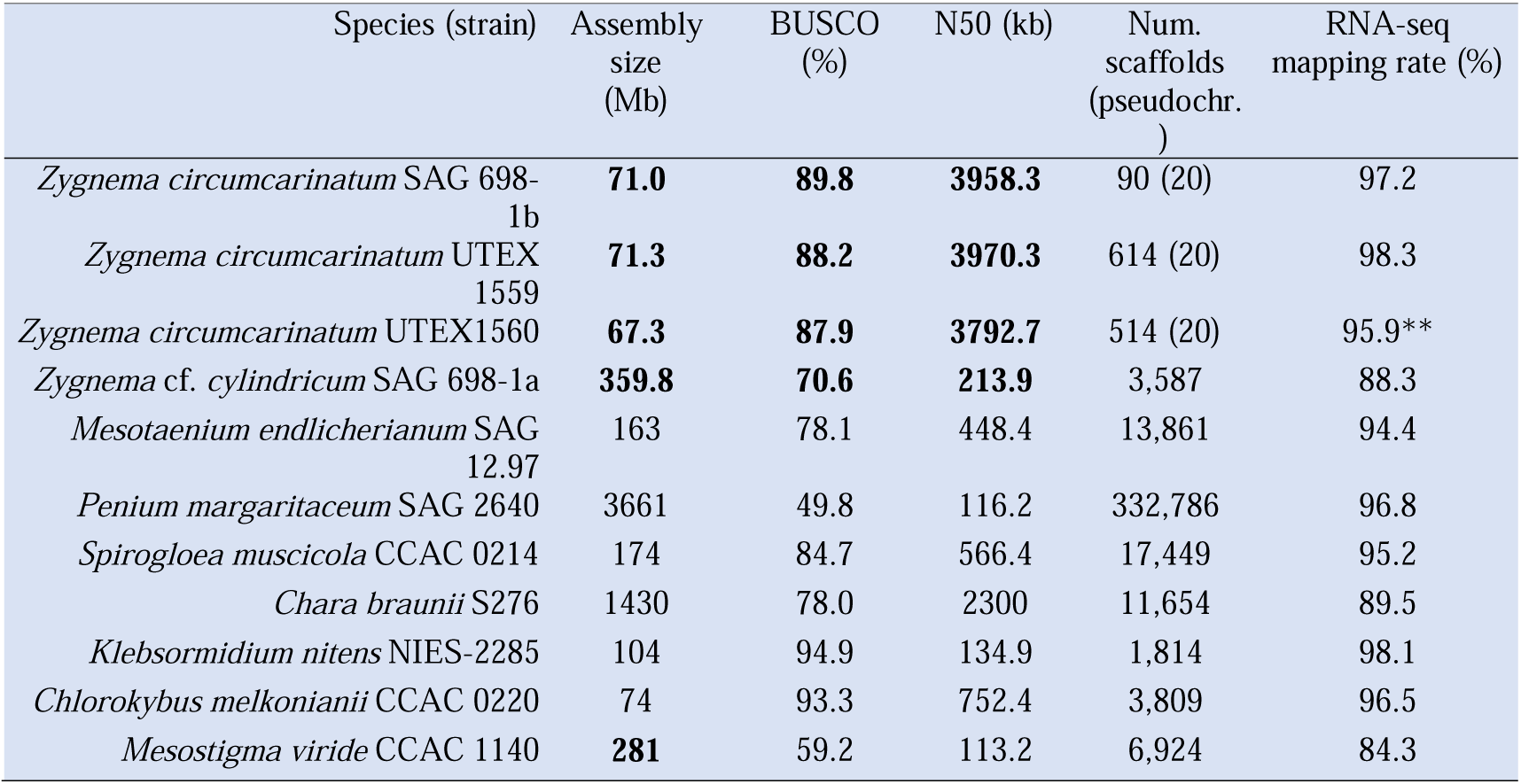
Genome assembly statistics for the new Zygnema genomes and available streptophyte algae (Hori et al., 2014; Nishiyama et al., 2018; Cheng et al., 2019; Jiao et al., 2020; Wang et al., 2020) (see Table S1B-E for further details). Mapping rate of Z. circumcarinatum UTEX 1560 was calculated by using SAG 698-1b RNA-seq reads mapped to UTEX 1560 genome.

The plastome (157,548 base pairs (bp)) and mitogenome (216,190 bp, **Figure S3**) of SAG 698-1b were assembled into complete circular genomes (**Figure S3**). The plastome of SAG 698-1b is identical to those of UTEX 1559 (GenBank ID MT040697; Orton et al., 2020) and UTEX 1560. The mitogenomes of SAG 698-1b (OQ319605) and UTEX 1560 are identical but slightly longer than that of UTEX 1559 (MT040698, 215,954 bp; Orton et al., 2020) due to extra repeats (**Figure S4**). These observations agree with the history of the strains (see Supplementary Information).

The nuclear genome assembly of SAG 698-1a is four times larger (360 Mb) than those of *Z. circumcarinatum* (**Table 1, Figure 1E**). K-mer frequencies (**Figure S2**) strongly suggest that SAG 698-1a is a diploid organism with an estimated heterozygosity rate of 2.22%. This supports previous reports of frequent polyploidy in Zygnematophyceae (Allen, 1958). The marked genome size differences further support the notion that they are two different species (Table 1, pictures in **Figure 1A**). Following a recent study (Feng et al. 2021), we refer to SAG 698-1a as *Zygnema* cf. *cylindricum*. Our molecular clock analyses suggest that *Z.* cf. *cylindricum* (SAG 698-1a) and *Z. circumcarinatum* (SAG 698-1b, UTEX 1559, UTEX 1560) diverged from one another around 236 Ma (**Figure 1E, Table S1F**).

The plastome of SAG 698-1a was available (Turmel et al., 2005) and we here determined its mitogenome (OQ316644) (**Figure S5**), which, at 323,370 bp in size, is the largest known among streptophyte algae. Compared to SAG 698-1b, the mitogenome of SAG 698-1a contains more and much longer introns (**Table S1G,H**).

### *Z. circumcarinatum* has the smallest sequenced streptophyte algal genome and no recent whole genome duplications

The three *Z. circumcarinatum* genomes reported here have the smallest nuclear genomes of all streptophyte algae sequenced thus far (**Table 1**). They have the highest protein coding gene density, smallest percentage of intergenic regions, highest exon percentage, and lowest repeat content in Zygnematophyceae (**Figure 1E, Table S1I**). The genome of SAG 698-1b contains 23.4% of repeats (**Table S1J**), mostly consisting of simple repeats (6.4%) and transposable elements of the MITE (4.3%), Gypsy (2.9%), and Copia (1.9%) families. For comparison, the *Zygnema* cf. *cylindricum* SAG 698-1a genome contains 73.3% of repeats, consisting of Copia (29.8%), MITE (11.6%), Gypsy (5.9%), and simple repeats (2.1%). The phylogenetic position and genome sizes of *Z. circumcarinatum* suggest genomic streamlining in this species, as shown also for *K. nitens* (Hori et al. 2014) and perhaps *C. melkonianii* (Wang et al., 2020). While the mechanisms of genomic streamlining are obscure, it clearly occurred independently in different streptophytes; genome shrinkage may be a signature of adaptations to new ecological niches (Bhattacharya et al., 2018).

No evidence for whole genome duplication (WGD) was found in SAG 698-1b. MCscan (Wang et al., 2012) was used to identify regions of conserved synteny, finding 28 syntenic blocks (≥ 4 genes per block, distance between two colinear regions < 20 genes) totaling 236 genes (1.44% of the 16,617 annotated genes) (**Figure S6A**). Increasing the distance among blocks to < 30 genes identified 190 syntenic blocks containing 1,298 (7.9%) genes (**Figure S6B**). The distribution of synonymous distances among paralogous gene pairs (Ks) in syntenic blocks found a single peak at Ks ∼ 0.2 (**Figure S6C**). This agrees with the apparent lack of WGD in *Z. circumcarinatum*. Because chromosome-level assemblies are so far lacking for other streptophyte algae, similar analyses were conducted for the moss *P. patens*, reported to have had at least two WGDs (Lang et al., 2018; Rensing et al., 2008). *P. patens* had more syntenic blocks (**Figure S6D: 20%** genes in syntenic blocks **and S6E:** 24% genes in syntenic blocks), and a single peak Ks ∼ 0.8 (**Figure S6F**) that is likely a fusion of the two known WGDs (Ks ∼ 0.5–0.65 and Ks ∼ 0.75–0.9 (Lang et al., 2018)). In the absence of evidence for WGD in SAG 698-1b, synteny blocks are likely the result from segmental duplications. Congruently, the top enriched Pfam domains in these regions are related to retrotransposons.

### Comparisons of the three *Z. circumcarinatum* genomes

Our phylogenetic analyses show that SAG 698-1b and UTEX 1560 are closer to each other than to UTEX 1559 (**Figure 1E**), a result confirmed by seven- and four-genome phylogenies using thousands of single copy genes (**Figure S7**). Gauch (1966) reported that UTEX 1559 was a non-functional mating type (+) whereas UTEX 1560 and SAG 698-1b were functional mating type (-). This agrees with our conjugation experiments that failed to conjugate UTEX 1559 with UTEX 1560 or SAG 698-1b. Whole genome alignments of UTEX 1559 and UTEX 1560 against SAG 698-1b respectively, identified high alignment coverages and regions of conserved synteny across most chromosomes (**Figure S8A,B**). SAG 698-1b showed higher alignment coverage (less genomic rearrangements) with UTEX 1560 than to UTEX 1559, although aligned regions are slightly more similar to UTEX 1559 (**Figure S8C,D**). Among the three *Z. circumcarinatum* genomes, chromosomes 20, 13, and 16 differ the most (see **Figure S8E** for a three-way alignment of Chr20), suggesting that they might contain sex/mating determination loci. The mating loci in *Zygnema* are so far unknown and we did not identify homologs of the sex hormone proteins (PR-IP and its inducer) described in *Closterium* (but homologs were found in *Penium margaritaceum*) (**Table S1K**). Gene content comparison found 17,644 core genes, i.e., shared by all three *Z. circumcarinatum* genomes (**Figure S8F**). Most of the unique and shell genes have no known Pfam domains and no hits in NR (NCBI non-redundant protein sequence database) and thus remain functionally unknown.

### Comparative genomics identifies significantly enriched orthogroups and domains in Zygnematophyceae and land plants

Comparative genomics were performed with annotated proteins from 16 representative algal and plant genomes, clustered into orthogroups (groups of orthologs or OGs) by OrthoFinder. 4,752 orthogroups contained at least one representative of Chlorophyta, Embryophyta, Zygnematophyceae, and other streptophyte algae (**Figure 2A,B,C,D**). We identified clade-specific orthogroups according to the species phylogeny (**Figure 2A**); the enrichments in gene ontology (GO) terms (**Figure 2C**) and Pfam domains (**Figure 2B**) were inferred by binomial test q-values against the background of GO terms and Pfam domains present in the general set of 4,752 orthogroups (**Figure 2D**). In the ancestor of Zygnematophyceae and Embryophyta, we inferred an overrepresentation of Pfam domains (**Figure 2B)** including (i) Chal_sti_synt_C (found in the key enzyme of the flavonoid pathway chalcone synthase (CHS)), (ii) Methyltransf_29 (found in an *Arabidopsis thaliana* gene (AT1G19430) required for cell adhesion (Krupková et al. 2007)), (iii) PPR domains involved in organellar RNA binding and editing, and (iv) domains related to plant immunity such as LRR and Peptidase_S15 (Muszewska et al., 2017), PK_Tyr_Ser-Thr, and Thioredoxins (Kumari et al., 2021). Some of these orthogroups and domains overrepresented in Zygnematophyceae + Embryophyta could be the results of horizontal gene transfer (HGT), such as Chal_sti_synt_C (Ma et al., 2022) and O-FucT (**Figure S9**). O-FucT is the GDP-fucose protein O-fucosyltransferase domain (PF10250), and its homologs in streptophytes are likely transferred from arbuscular mycorrhizal (AM) fungi Mucoromycotina and Glomeromycota (**Figure S9**). In SAG 698-1b, the O-FucT domain-containing gene (Zci_09922) is expressed under drought and cold stresses.

**Figure 2:**
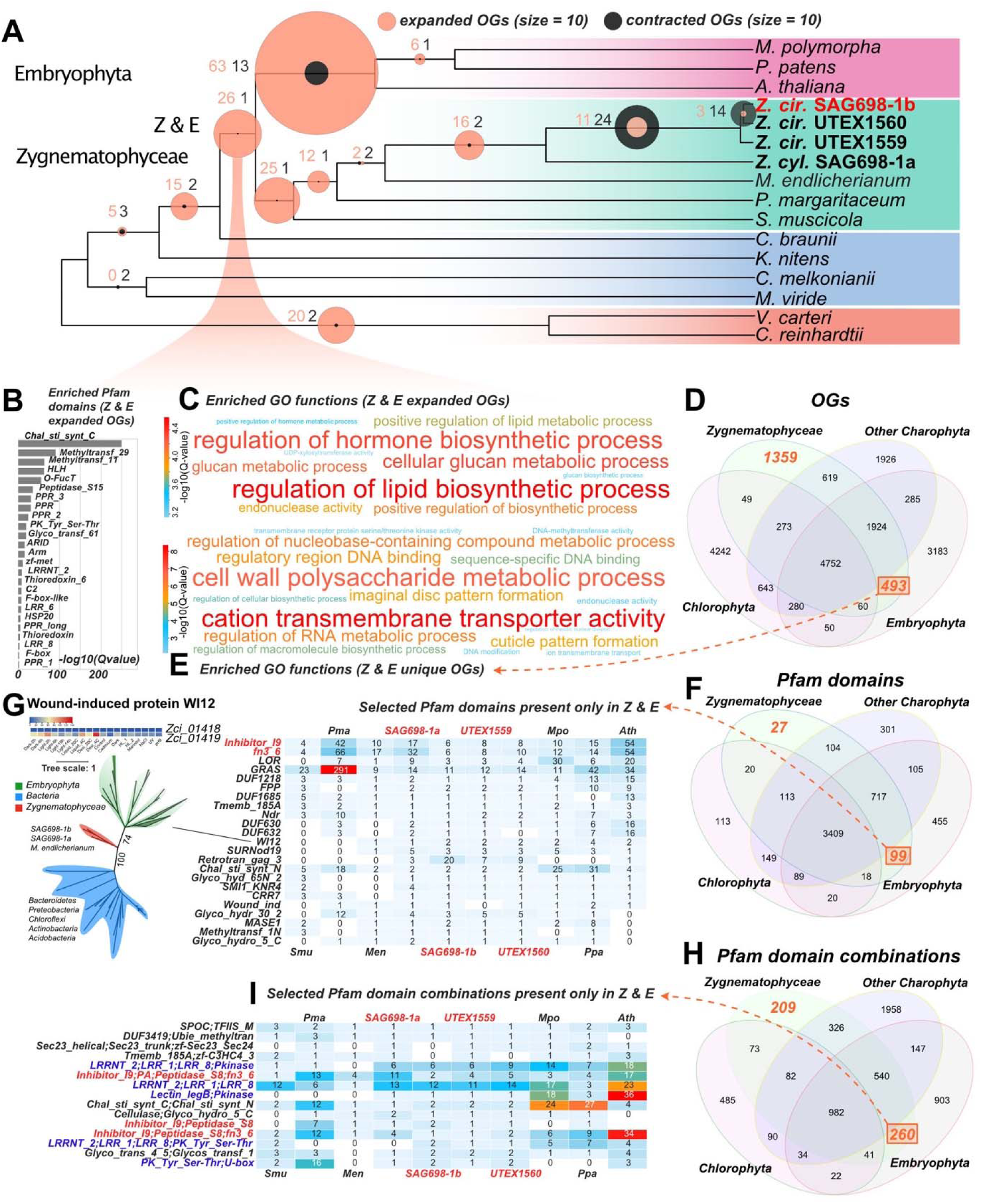
Comparative genomics of 13 algal and 3 land plant genomes. (A) Gene family expansion and contraction patterns estimated by CAFE using Orthofinder-identified orthogroups and the time-calibrated phylogeny of Figure 1E. Key nodes are indicated on the tree and circles denote significant expansions and contractions (circle size reflects the number of expanded/contracted orthogroups/OGs). (B) Pfam domain enrichment for genes on the node leading to Zygnematophyceae and Embryophyta (Z&E). (C) Functional (GO) enrichment for the Z&E node. (D) Orthogroups overlap among Chlorophyta, Embryophyta, Zygnematophyceae, and other streptophyte algae. (E) Enriched GO terms in the 493 orthogroups exclusive to Zygnematophyceae and Embryophyta. (F) Pfam domain overlap among Chlorophyta, Embryophyta, Zygnematophyceae, and other streptophyte algae. (G) Exclusive Pfam domains found only in Zygnematophyceae and Embryophyta. One Pfam family WI12 was studied with phylogenetic analysis, suggesting a possible horizontal gene transfer from bacteria and expression response to stresses. (H) Pfam domain combination overlap among Chlorophyta, Embryophyta, Zygnematophyceae, and other streptophyte algae. (I) Exclusive Pfam domain combinations in Zygnematophyceae and Embryophyta.

The Zygnematophyceae + Embryophyta (Z+E) ancestor showed enriched GO terms related to biosynthesis of pytohormones, lipids, and glucan (**Figure 2C).** 493 orthogroups were exclusively found in Z+E and not in any other studied species (**Figure 2D**), and these were enriched in GO terms “cation transmembrane transporter activity” and “cell wall polysaccharide metabolic process”. 1,359 orthogroups were Zygnematophyceae-specific, with enriched GO terms “phosphorylation”, “pyrophosphatase activity”, “transmembrane receptor protein serine/threonine kinase activity”, “cellular response to abscisic acid stimulus”, and “polysaccharide biosynthetic process” (**Figure 2E**).

Regarding Pfam domains, we found 3,409 to be present in at least one representative of Cholorophyta, Embryophyta, Zygnematophyceae, and other streptophyte algae, whereas 99 were exclusive to Z+E, and 27 were Zygnematophyceae-specific (**Figure 2F**). Unlike the Pfam domains enriched in the Z+E ancestor (**Figure 2B)**, the 99 or 27 domains were not found outside of Zygnematophyceae and Embryophyta. The most prevalent domains that are exclusively of Z+E (**Figure 2G**) include several transcription factors (more details below). In some cases, the domains exclusive to Z+E could be the result of HGT. For example, the Inhibitor_I9 and fn3_6 domains are among the most abundant in Z+E (**Figure 2G**) and often co-exist with Peptidase_S8 domain in plant subtilases (SBTs) (**Figure 2I**), which has been reported to originate by HGT from bacteria (Xu et al., 2019). The WI12 domain, named after the plant wound-induced cell wall protein WI12, was also possibly gained by HGT from bacteria (**Figure 2G**). WI12 expression is induced by wounding, salt, methyl jasmonate, and pathogen infection in the facultative halophyte ice plant (Yen et al., 2001), and plays significant roles in reinforcing cell wall and is involved in the defense to cyst nematodes in soybean (Dong and Hudson, 2022). In SAG 698-1b, one of the two genes containing the WI12 domain (Zci_01419) is significantly upregulated under desiccation and cold stresses.

Because new functions can arise through domain combinations, we searched for lineage-specific Pfam domain combinations in our dataset (**Figure 2H)**. 982 Pfam domain combinations are shared by all studied genomes, 260 being unique to Z+E and 209 to Zygnematophyceae. Among those exclusive to Z+E **(Figure 2I),** we found Lectin_legB and Pkinase domains, which despite having older evolutionary origins, were only combined into a protein in the Z+E ancestor (e.g., Zci_10218). A search for the 99 and 260 domain combinations by BLASTP against the NR database (**Table S2A,B**) failed to identify such combinations in sequenced chlorophyte or streptophyte algal genomes but were occasionally found in fungi, prokaryotes, and viruses. This pattern could be the product of HGT, functional convergence, rampant gene loss, or contamination. Combining existing protein domains is a powerful mechanism for functional innovation, as shown for, cell adhesion, cell communication and differentiation (Itoh et al., 2007; Vogel et al., 2004).

### Orthogroup expansions reveal increased sophistication and resilience

We inferred 26 significantly expanded orthogroups in the Z+E ancestor (**Figure 2A; Table S3A,B**). Three are related to phytohormone signaling: ethylene-responsive element (ERE)-binding factors (OG 22) and PP2Cs (OG 548 and OG 830), regulatory hubs for diverse responses, including to ABA, whose evolutionary origin—albeit with an ABA-independent role in algae—was already pinpointed (de Vries et al., 2018; Sun et al., 2019; Cheng et al., 2019). Several expansions suggest more sophisticated gene networks featuring calcium signaling (putative calcium-dependent protein kinases; OG 19) and ubiquitin-mediated proteolysis (RING/U-box proteins; OG 78), both cornerstones in plant stress response and environmental signaling (Yee and Goring, 2009; Reddy et al., 2012). We inferred expansion of transmembrane transporters such as major facilitator superfamily proteins (OG 169), amino acid transporters (OG 333), acetate channels (OG 353), RAB GTPases (OG 174), and Golgi SNAREs (OG 963). Two orthogroups might be involved in interactions with microbes and fungi, including subtilases (OG 23; Xu et al. 2019) and glycosyl hydrolases with a chitin domain (OG 857; (Parrent et al. 2009)). Growth and development are underpinned by expanded beta-glucosidases (OG 85) involved in xyloglucan biosynthesis and/or plant chemical defense (Morant et al. 2008), and developmental regulators root hair defective 3 GTP-binding proteins (OG 996) (Yuen et al., 2005) and ARID/BRIGHT DNA binding TFs (transcription factors) (OG 2169). A comprehensive analysis of transcription associated proteins (TAPs) with TAPscan v.3 (Petroll et al. 2021) showed higher numbers of TFs in land plants as compared to algae, as expected due to their more complex bodies (Fig. 5B; Table S3B). *Zygnema* species had comparatively more TAPs than other algae (525-706 vs. 269 in *Ulva* and 371 in *Chlorokybus*; Table 5B; Table S3B). All four studied *Zygnema* strains show similar TAP profiles, with the exception of MYB-related TF family and PHD (plant homeodomain), which were, respectively, 2- and ≥ in SAG 698-1a.

The common ancestor of Zygnematophyceae displayed 25 significantly expanded orthogroups (**Figure 2A; Table S3B**). Most expanded are alpha-fucosyltransferases (OG 89) involved in xyloglucan fucosylation (Faik et al., 2000). We found ethylene sensors and histidine kinase-containing proteins (OG 94), bolstering the idea that two-component signaling is important and active in filamentous Zygnematophyceae (Ju et al., 2015; Bowman et al., 2019). Several orthogroups were associated with typical terrestrial stressors: aldo-keto reductases closely related to *M. polymorpha* Mp2g01000 (OG 942) that could reflect ROS scavenging machinery (Stiti et al. 2021), DNA helicases for DNA repair and recombination (OG 1471), and methyltransferases (OG 269, OG 369) that could underpin specialized metabolism of phenylpropanoids for stress response (Lam et al., 2007). The phenylpropanoid pathway is in fact a well-known response to terrestrial stressors (Dixon and Paiva, 1995) and its enzymes have deep roots in streptophyte evolution (de Vries et al., 2021; Rieseberg et al., 2023); *Zygnema* has a set of phenylpropanoid enzyme homologs comparable to those reported before (**Figure S14**). We found expansions in putative light-oxygen-voltage sensitive (LOV)-domain containing proteins (OG 1897), photoreceptors mediating responses to environmental cues (Glantz et al., 2016) that imply a more elaborate response to rapidly changing light regimes typical for terrestrial habitats (see also below). Several expanded orthogroups relate to developmental processes. Expanded signaling and transport, possibly related to filamentous growth, include calcium signaling (OG 56), zinc-induced facilitators (OG 258), cysteine-rich fibroblast growth factor receptors found in the Golgi apparatus (OG 518), and cation/H+ antiporters (OG 809). Cation/H+ antiporters are closely related to *A. thaliana* nhx5/nhx6 that act on pH and ion homeostasis in the endosome and are key for membrane trafficking in the trans-Golgi network (Bassil et al., 2011; McKay et al., 2022), and diverse developmental processes (Dragwidge et al., 2018). A dynein homolog (OG 72) was significantly contracted in Zygnematophyceae (**Table S3B**), which might be associated with cytokinesis of cilia and flagella and thus in line with the loss of motile gametes in Zygnematophyceae and Embryophyta (OG 72 was also identified as contracted in that ancestor).

*Zygnema* is noteworthy among algae for its stress resilience and can grow in extreme habitats such as the Arctic, where it is abundant (Pichrtová et al., 2018; Rippin et al., 2019). Its broad ecological amplitude is confirmed by numerous studies on temperature and light stress (summarized in Permann et al., 2022b) and desiccation stress (Becker et al., 2020). The common ancestor of all four *Zygnema* strains is inferred to have had 16 significantly expanded orthogroups (**Figure 2A; Table S3B**), including PP2Cs (OG 548), early light-inducible proteins (ELIPs; OG 97), and low-CO2 inducible proteins (LCICs, OG 459). Previous studies found ELIPs to be among the top upregulated genes when *Zygnema* is exposed to environmental challenges, including growth in the Arctic (Rippin et al., 2017; 2019; de Vries et al., 2018). We further found expansion of HSP70 chaperones (OG 277) involved in protein folding and stress response as previously observed in *Mougeotia* and *Spirogyra* (de Vries et al., 2020), as well as Leucine-rich repeat (LRR) proteins (OG 35 and OG 995) related to plant-microbe interactions. Two orthogroups were significantly contracted: GTP binding elongation factor *Tu* family (OG 251) and seven transmembrane MLO family protein (OG 320). On balance, the evolution of gene families reflects *Zygnema*’s resilience in the face of challenging habitats.

### Gene gains facilitated major cell wall innovations

The earliest land plants had to overcome a wide range of stressors (Fürst-Jansen et al., 2020) and cell walls are the first layer of protection from the environment. We reconstructed the evolutionary history of 38 cell wall-related enzyme families (**Table S1L**). Large gene families were split into 76 well-supported subfamilies (>70% non-parametric bootstrap supports) according to phylogenetic trees (Data S1, homolog counts are in **Table S1M** and **Figure 3A)**. Most subfamilies belong to carbohydrate active enzyme (CAZyme) families known for the synthesis and modifications of celluloses, xyloglucans, mixed-linkage glucans, mannans, xylans, arabinogalactan proteins (AGPs), and pectins (Table **S1L,M**, **Figure 3A**). CAZymes include glycosyl transferases (GTs), glycosyl hydrolases (GHs), carbohydrate esterases (CEs), and polysaccharide lyases (PLs). Analyzing the 76 enzyme subfamilies revealed that (i) Z+E share all the major enzymes for the synthesis and modifications of the diverse polysaccharide components, including those for sidechains and modifications (**Figure 3B**, 42-56 subfamilies in Zygnematophyceae vs 63-69 in Embryophyta); (ii) many of the enzymes for cell wall innovations, especially for polysaccharide backbone synthesis, have older evolutionary origins in the common ancestor of Klebsormidiophyceae, Charophyceae, Zygnematophyceae, and Embryophyta (**Figure 3B**, 34-69 subfamilies vs 5-7 in Chlorokybophyceae and Mesostigmatophyceae). Many of such subfamilies are expanded in Zygnematophyceae (**Figure 3B % of genes,** e.g., GH16_20, GT77, CE8, CE13 in **Figure 3A**); (iii) genes involved in the syntheses of different cell wall polymers (backbones and sidechains) are co-expressed in SAG 698-1b (**Figure 3C**); (iv) phylogenetic patterns (**Data S1**) suggest that HGT could have played an important role in the origin of the enzymatic toolbox for cell wall polysaccharide metabolism (**Figure 3A, Supplemental Text 2**). HGT is more common for degradation enzymes (e.g., GH5_7, GH16_20, GH43_24, GH95, GH27, GH30_5, GH79, GH28, PL1, PL4) but it is also observed for GT enzymes; (v) frequent gene loss creates scattered distribution of homologs in Streptophyta (**Figure 3A**; e.g., *Zygnema* lacks GH5_7, GH35, GT29, GT8, CE8, GH28, PL1).

**Figure 3:**
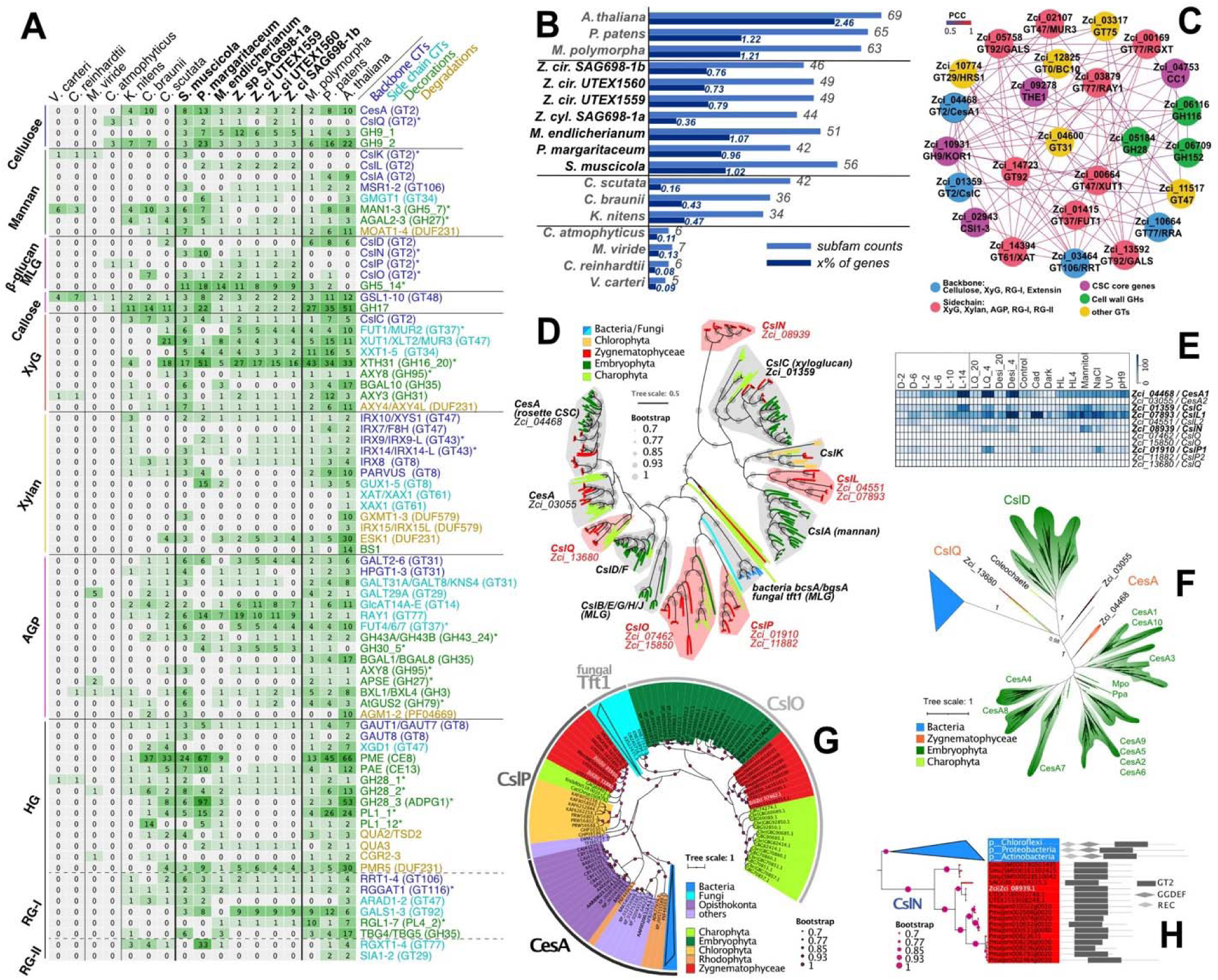
Cell wall innovations revealed by protein family analyses. (**A**) Heatmap of homolog presence in 76 enzyme subfamilies (rows) across 17 plant and algal genomes (except for *Coleochaete scutata* that we used its transcriptome). Enzyme subfamilies are grouped by polysaccharide and colors indicate their biochemical roles; phylogenetic patterns compatible with gene gain that might have involved horizontal gene transfer (HGT) are with asterisks. (**B**) Counts of subfamilies and gene percentages (with respect to the total annotated genes) across the 17 species. Shown in the plot is the gene percentage x 100. (**C**) Co-expression network of SAG 698-1b containing 25 genes (most belonging to the 76 analyzed subfamilies) involved in cell wall polysaccharide syntheses. (**D**) Phylogeny of GT2 across the 17 species. Major plant CesA/Csl subfamilies are labeled by the SAG 698-1b homolog and newly defined subfamilies are in red. Ten bacterial beta-glucan synthase (BgsA), and fungal mixed-linkage glucan (MLG) synthase (Tft1) homologs are included to show their relationships with plant CesA/Csl subfamilies. (**E**) Gene expression of 11 SAG 698-1b GT2 genes across 19 experimental conditions (3 replicates each); highly expressed genes are in red. (**F**) Phylogeny of GT2 with ZcCesA1 (Zci_04468) homologs retrieved by BLASTP against NCBI’ NR (E-value < 1e-10); colors follow D and >5,000 bacterial homologs from >8 phyla are collapsed (blue triangle; the three major phyla are indicated). (**G**) Phylogeny of GT2 with ZcCslP1 (Zci_0910) homologs retrieved by BLASTP against NCBI’s NR (E-value < 1e-40); colors follow D and 279 bacterial CesA homologs (blue triangle) and 363 fungal MLG synthase homologs (turquoise triangle) are collapsed (see **Data S1-12** for details). (**H**) Phylogeny of GT2 with ZcCslN (Zci_08939) homologs retrieved by BLASTP against NCBI’s NR (E-value < 1e-10); colors follow D and bacterial homologs are collapsed (blue triangle); Pfam domain organization are shown on the right (see **Data S1-13** for details).

We performed a careful analysis of the GT2 family, which contains major cell wall synthesis enzymes such as cellulose synthase (CesA) for the synthesis of cellulose and Csl (CesA-like) for hemicellulose backbones (**Figure 3D**). Among the 11 SAG 698-1b CesA/Csl homologs, ZcCesA1 (Zci_04468), ZcCslL1 (Zci_07893), ZcCslC (Zci_01359), ZcCslN (Zci_08939), ZcCslP1 (Zci_0910) are highly expressed in response to various stresses (**Figure 3E**), in agreement with Fitzek et al (2019) who showed response to osmotic stress. The two CesAs homologs in SAG 698-1b (**Figure 4D, 4F, Data S1-1**) and all other Zygnematophyceae homologs are orthologs of land plant CesA; Zygnematophyceae also have a second CesA homolog (ZcCesA2; Zci_03055) not found in land plants but shared with other streptophytes. The presence of land plant-like CesAs only in Zygnematophyceae suggests that the CSC (cellulose synthase complex) structure in a six-subunit rosette typical of land plants evolved in their common ancestor; this agrees with observations made by electron microscopy and isotope labeling that found an hexameric rosette CSC in Zygnematophyceae but no in other algae (Tsekos, 1999). ZcCesA1 (Zci_04468) is co-expressed with four known plant primary cell wall CSC component core genes: KOR (Zci_10931), CC1 (Zci_04753), CSI1 (Zci_02943), THE (Zci_09278) (**Figure 3C**). This extends previous observations (Lampugnani et al., 2021) suggesting that co-expression of CSC component genes is evolutionarily conserved since the common ancestor of Zygnematophyceae and land plants.

**Figure 4.**
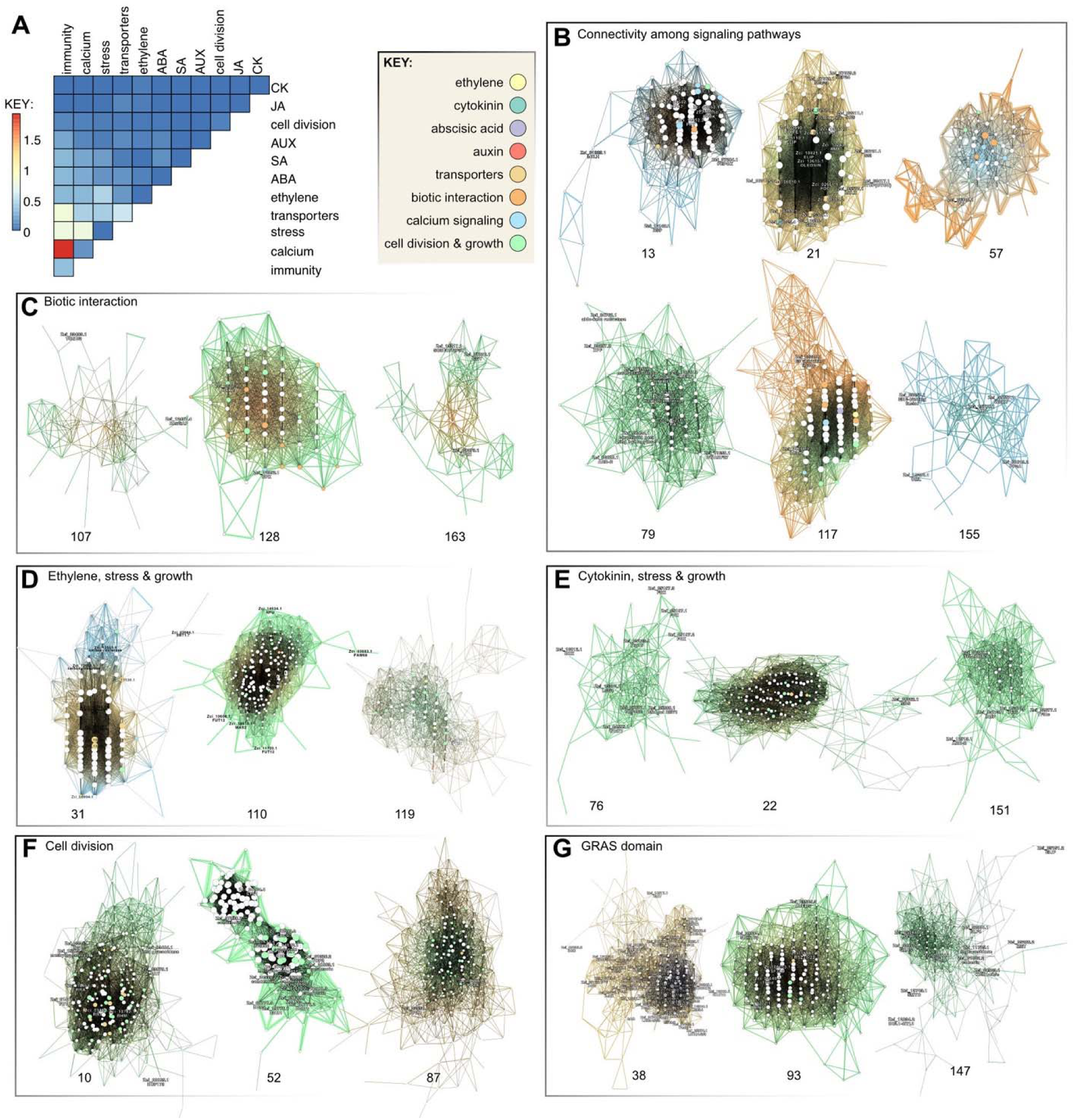
Gene co-expression modules in *Zygnema circumcarinatum* SAG 698-1b. (A) Heatmap of per-module co-occurrence frequencies among genes associated with plant-microbe (p-m) interaction, calcium signaling, phytohormone, stress, transporters, cell division, and diverse phytohormones (ethylene, cytokinin, abscisic acid/ABA, auxin/AUX, jasmonic acid/JA, salicylic acid/SA); based on 150 out of 406 total modules showing co-ocurrence of at least two functional categories. (B) Modules reflecting connectivity among signaling pathways, (C) biotic interaction, (D) ethylene, stress, and growth, (E) cytokinin, strees & growth, (F) cell division, (G) GRAS-domain containing genes. In gene networks, nodes are genes (size proportional to number of neighbors) and edges reflect co-expression (width proportional to Pearson’s correlation coefficient and colors are those of interconnected genes); numbers below indicate network number; gradient colors of the edges highlight the two dominant gene categories indicated in the KEY. The full gene co-expression results can be accessed in our online portal (https://zygnema.sbs.ntu.edu.sg/).

Our GT2 family analyses identified new Csl families (**Figure 3D; Data S1**) that are likely involved in the synthesis of cellulose, mannan, mixed-linkage glucan (MLG), or other beta-glucans. CslQ clade represents the ancestral form of all streptophyte CesAs (possibly also CslD) (**Figure 3D**), and was likely gained by an ancient HGT from bacteria into the common ancestor of Streptophyta algae (**Figure 3F**). ClsK was thought to be restricted to Chlorophyta (Yin et al., 2009; Yin et al., 2014) but has recently been found in Zygnematophyceae (Mikkelsen et al., 2021). CslL is a new Zygnematophyceae-specific family. CslK and CslL might be responsible for the mannan backbone synthesis in green algae (Chlorophyta and Zygnematophyceae) given their closer relationship to CslA than CslC (**Data S1-1, Figure 3D**). CslO, also found in non-seed land plants (**Figure 3G**), contains a *Physcomitrium pattens* homolog (Pp3c12_24670) involved in the synthesis of the *P. patens*-specific polysaccharide arabinoglucan (AGlc) (Roberts et al., 2018). CslO is closely related to a large fungal clade containing Tft1 (XP_748682.1) characterized for the synthesis of MLG, a fungal cell wall component (Samar et al., 2015). This opens the possibility that CslO was horizontally transferred from fungi (**Figure 3G**). CslP is absent in land plants, and together with CslO further clustered with known CesAs from oomycetes, tunicates, amoeba, and red algae (Blanton et al., 2000; Blum et al., 2010; Matthews et al., 2010; Matthysse et al., 2004) (**Figure 3G**). The close affinity with a large bacterial clade of BcsA/CesAs (>10,000 proteins from >10 phyla; BLASTP of ZcCslP1 against NR, E-value < 1e-30), including experimentally characterized cyanobacterial CesAs, suggests that eukaryotic CslP-like proteins could have originated via HGT (**Figure 3G, Data S1-12**). CslO evolved from CslP-like proteins and was subjected to duplication and functional diversification in eukaryotes (e.g., MLG synthesis in fungi and likely also in various microalgae, AGlc synthesis in mosses), while the CesA function of CslP-like proteins is conserved in oomycetes, tunicates, amoeba, red algae, and possibly also in green algae with CslP. CslN homologs are restricted to Zygnematophyceae and bacteria (no significant NCBI NR hits outside them at E-value < 1e-40), which suggests a possible HGT from bacterial CesA or other beta-glucan synthases (**Figure 3H, Data S1-13**) that often contain N-terminal GGDEF and REC domains. The bacterial BcsA-like CslN and CslP, if biochemically characterized as CesAs in Zygnematophyceae in the future, are clearly of a distinct origin than land plant CesAs, thus representing an example of convergent evolution.

Fucosyltransferase (FUT, member of the GT37 family), is significantly expanded in Zygnematophyceae (**Data S1-49,** OG 89 is the most expanded orthogroup; see below). Xyloglucan fucosylation is implicated in stress response and was long thought to be land-plant-specific but was recently shown to be present in the Zygnematophyceae *Mesotaenium caldariorum* (Tryfona et al., 2014; Mikkelsen et al., 2021). Our analyses show a sparse phylogenetic distribution of FUT homologs in *Zygnema* and embryophytes and distant homologs in *Chara* and *Klebsormidium*, as well as in many soil saprotroph Mortierellaceae fungi (Telagathoti et al., 2021) (**Data S1-19**). Phylogenetic patterns could be compatible with an ancient HGT between plants and fungi (**Data S1-19**). The enzyme XTH (GH16_20), which degrades the xyloglucan backbone, shows a similar pattern compatible with HGT between fungi and streptophytes (**Data S1-20, S1-21**), in agreement with a recent report (Shinohara and Nishitani, 2021). Type II fucosidase (AXY8, GH95), involved in degrading fucosylated xyloglucans and arabinogalactan proteins (Léonard et al., 2008; Wu et al., 2010), might have been acquired by HGT from bacteria (**Data S1-22**). Overall, these findings agree with an earlier origin of xyloglucan backbone enzymes and a later origin of side chain modification enzymes in the common ancestor of Zygnematophyceae and Embryophyta (Del-Bem, 2018; Mikkelsen et al., 2021). Recent reports have investigated the occurrence of homogalacturonans (e.g., Herbruger et al. 2019) and AGPs in *Zygnema* (Palacio-Lopez et al. 2019) and the filamentous Zygnematophyceae *Spirogyra*, where recently a rhamnogalactan protein has been described (Pfeifer et al., 2022); further, the specific cell wall composition of zygospores of the filamentous Zygnematophyceae *Mougeotia* and *Spirogyra* have been described (Permann et al., 2021; 2022a). Cell wall modifications by endotransgylcoylases and novel transglycosylation activities between xyloglucan and xylan, xyloglucan and galactomannan were described previously (Herburger et al., 2018). Overall, the phylogenetic analyses of key cell wall enzymes (**Data S1, Supplemental Text 2, Table S1L**) highlighted the importance of ancient HGTs contributing to evolutionary innovations of cell walls, similarly to what has been proposed for other traits (Yue et al., 2012; Cheng et al., 2019; Ma et al., 2022).

### Gene co-expression networks in *Zygnema circumcarinatum* SAG 698-1b

We explored functional gene modules in *Zygnema circumcarinatum* (SAG 698-1b) by inferring gene co-expression networks from RNA-seq data from a diverse set of 19 growth conditions (various day-light cycles and dark, liquid and agar cultures at 20°C, pH=9 and terrestrial stressors including cold treatment at 4°C, desiccation at 4°C, high light, high light at 4°C, or UV; see Methods). We obtained 406 clusters containing 17,881 out of the 20,030 gene isoforms annotated in the genome. Gene co-expression networks are available through the CoNekT web portal (Proost and Mutwil, 2018; https://zygnema.sbs.ntu.edu.sg/). We searched for homologs of genes related to (i) cell division and development, (ii) multicellularity, (iii) stress response, (iv) transporters, (v) phytohormones, (vi) calcium signaling, and (vii) plant-microbe interaction. Candidate genes were drawn from the literature and the expanded orthogroups. 150 out of 406 clusters showed co-occurrence of at least two such functional categories, the most frequent co-occurrence being plant-microbe interaction and calcium signaling, followed by plant-microbe interaction and stress (Figure 4A).

### On the deep evolutionary roots of the plant perceptron

Land plants are sessile. Sensing environmental conditions and modulating growth alongside mounting appropriate physiological responses is vital for plants, especially under adverse conditions. Several of our co-expression clusters reveal the concerted action of diverse signaling pathways to sense, process, and respond to environmental cues. This agrees with the concept of the plant perceptron (Scheres & van der Putten 2017) that views plant biology as a molecular information-processing network—formed by genes and their biochemical interactions—that enable adequate response given a combination of input cues. This implies a high degree of connectivity among signaling pathways to modulate a response to highly complex biotic and abiotic signals. Such connectivity establishes the foundation for an adaptive advantage of multicellular morphogenesis, where cell differentiation can be fine-tuned for acclimation to environmental cues.

In SAG 698-1b, many gene modules reflect a complex connectivity among signaling pathways including phytohormones, calcium signaling, transporters, and cell division and developmental genes. This shows that the plant perceptron has deeper evolutionary roots dating back at least to the common ancestor of land plants and Zygnematophyceae. Interestingly, even though some phytohormones are likely not present in *Zygnema* (see below), we observe gene homologs of their biosynthesis and signaling co-expressing with well-known effectors, thus suggesting these genes were already involved in response to abiotic and biotic stresses before the appearance of some phytohormones (see discussion in Fürst-Jansen et al., 2020). Several clusters (**Figure 4B**) contain early light-induced proteins (ELIPs), light-harvesting complex-related proteins that respond to light stimulus and can reduce photooxidative damage by scavenging free chlorophyll (Hutin et al., 2003) under cold stress (cluster 21) —as shown for other streptophyte algae (Han et al. 2013; de Vries et al. 2018)— but also under high light (cluster 20). In cluster 20, we identified a heat shock protein and a PsbS homolog putatively involved in non-photochemical quenching. Cluster 13 is deployed under dark conditions and shows expression of a red/far-red light receptor phytochrome in the canonical PHY2 clade (Li et al., 2015; **Figure S15**). Zygnematophyceae have a unique chimeric photoreceptor called neochrome, which is a combination of red/far-red sensing phytochrome and blue-sensing phototropin (Suetsugu et al., 2005). Neochromes have independently evolved twice, with a separate origin in hornworts (Li et al., 2014). In Zygnematophyceae, neochrome is hypothesized to be involved in chloroplast rotation to maximize light reception and to avoid photodamage (Suetsugu et al. 2005). In cluster 123, two neochrome homologs are co-expressed with zeaxanthin epoxidase and a few photosynthesis-related genes, in agreement with its role in fine-tuning photosynthesis.

Several clusters include leucine-rich repeat (LRR) proteins that might be involved in response to biotic stresses. Downstream, late embryogenesis abundant (LEA) proteins and protein phosphatases type 2C (PP2C) are often found, plant protein groups involved in diverse stress responses (Dure et al. 1981). Calcium signaling genes are also frequently expressed under stress conditions (cluster 117). Signal transducers are expected to affect many downstream genes through deployment of transcription factors (many visible in the clusters: WRKY, PSRP1, MYB, bHLH, mTERF, SCR), thereby allowing a tightly orchestrated response in time and space. As expected from the involvement of several transcription factors in cell division (e.g., MYB, SCR), we identify master cell cycle regulators (e.g., CYCP in cluster 21) and genes involved in cell division and development. Among phytohormone pathways, we highlight the histidine phosphotransfer protein (HPT) putatively involved in cytokinin signaling (AHP homolog; cluster 155), various ethylene responsive element-binding factors (clusters 57, 155), ABI1/2 homologs of the ABA biosynthesis (cluster 21), and the PYL homolog (cluster 13) that is an ABA receptor in land plants but probably not in algae (Sun et al., 2019). The expression of different ethylene response factors under desiccation (cluster 57) or other stress conditions (cluster 155) reveals functional specialization. In cluster 79, we identify ROS scavenging (cluster 79; Stiti et al. 2021; OG 942 expanded in Zygnematophyceae), various genes of the carotenoid pathway, and DNA repair machinery. Under cold conditions (cluster 21) and osmotic stress (cluster 103), oleosins are expressed for the formation of lipid droplets. In line with the need to tightly control metabolism, various genes of the carbon metabolism are also co-expressed, particularly under dark and cold conditions (clusters 13 and 21). Overall, the co-expression data suggest a joint action of genes to sense the environment and modulate growth.

### Gene modules associated with cell division and development

We compiled an extended list of 270 genes with experimental evidence for their involvement in mitosis and cytokinesis in *Arabidopsis thaliana* (including cytoskeleton and endomembrane transport and upstream regulators of cell division; Table S3D). Homolog distribution across streptophyte and chlorophyte genomes revealed secondary loss in *Zygnema* of microtubule plus tip CLASP and SPIRAL1 genes key to microtubule dynamics. There are important changes in the AUGMIN complex involved in microtubule nucleation, and GIP1 and GIP2 (γ-tubulin complex interactors) are missing from all analyzed Zygnematophyceae genomes. Homologs of AUGMIN2 and EDE1-like show weak conservation in Zygnematophyceae. Together, these data suggest that the Zygnematophyceae, and especially *Zygnema*, exhibit major differences in microtubule dynamics.

Some cell division genes are land plant-specific, including TANGLED (see Nishiyama et al., 2018), TRM (TON1-recruiting motif), SOSEKI, and EPF1 (epidermal patterning factor), which have pivotal roles in cell polarity and division and may have underpinned the multicellularity and increased morphological complexity of land plant. We also identified genes that likely originated in the common ancestor of Z+E: UGT1, SUN1, SUN2, and LONESOME HIGHWAY. These genes did not have reciprocal best BLAST hits in other streptophyte algae, although likely paralogs were found in *Klebsormidium nitens* (Table S3D). The clearest cases of genes originating in the Z+E ancestor are GRAS (Cheng et al. 2019), containing pro-orthologs of SCARECROW (SCR), SCARECROW-like, and SHORTROOT transcription factors, which in land plants are essential to control cell division orientation and tissue formation and they have also been associated with abiotic stress response. These genes allow branching of cell filaments (i.e., formative cell divisions), which are occasionally found in multicellular Zygnematophyceae.

GRAS homologs co-express with genes involved in cell division, cell cycle regulation, and cell wall functions (cluster 147, 38 and 93). All three clusters contain genes associated with abiotic stress responses, such as an ELIP homolog (OG 97 is expanded in *Zygnema*), beta-glucosidase (OG 85 expanded in the Z+E ancestor), calcium cation channel (DMI1/Pollux/Castor) and other calcium signaling components, and LRR receptor-like protein kinases. Links to various phytohormone pathways exist via the co-expression of a regulatory protein of ethylene receptor activity (TPR1) with an ABA signal transducer (AIP2) (cluster 93), and ABA4, gibberellin with auxin-related gene ARF10 (cluster 38). ARF10 often co-expresses with other TFs, some of which (e.g., CAMTA) have also been linked to biotic and abiotic stress responses (Xiao et al., 2021). Cluster 38 showcases links between photosynthesis, ABA signaling (ABA4), and the xanthophyll cycle (LUT2) that can mitigate photosynthetic stress. Photooxidative protection is also suggested by the co-expression of MPH2 photosynthetic acclimation factor and various redox proteins. The involvement of GRAS transcription factors in developmental and environmental signaling networks speaks of the evolution of a complex network to coordinate growth and stress since the common ancestor of Z+E.

In gene co-expression analysis, various gene modules reflect cell division and development. They included homologs of phragmoplastin (cluster 87), kinesins motor proteins (e.g., clusters 52, 87), spindle assembly proteins (cluster 52), RAB GTPases (cluster 10, 87), SNARE components (clusters 52, 87), components of the cargo complex (clusters 10, 87), and cell division-related protein kinases. Cluster 52, deployed during long day (14h) cycles that likely reflects steady-state growth, includes cyclin and cyclin-dependent protein kinases involved in cell cycle regulation. Cell wall formation genes often co-express, including cellulose or 1,4-beta-glucan synthases or transferases (cluster 10), beta-glucosidase (cluster 10; OG 85 expanded in the Z+E ancestor), or fucosyltransferases (clusters 10, 52; OG 89 expanded in Zygnematophyceae; see also **Figure 3C**). The co-expression of plastid division genes (e.g., components of MinD, MinE, FtsZ1 and FtsZ2, ARC6 regulatory protein) speak of the coordinated division of plastids. Further indicators of active cell division are genes related to DNA packaging and segregation such as DNA topoisomerase, DNA helicase complex, chromatin remodeling factors, and components of the condensin I and II complexes. Co-expression of protein phosphatases (PP2A; cluster 87) and methyltransferases (clusters 10 and 52; OG 369 expanded in Zygnematophyceae) point to a functional link between stress-responsive and cell division genes.

### Symbiotic genes are not co-expressed in Zygnematophyceae

The symbiotic association with fungi was one of the key innovations that allowed plants to colonize land (Rich et al. 2021). Phylogenetic analyses of land plant genes known for their symbiotic functions revealed four of such genes in Zygnematophyceae (Delaux et al. 2015). It was thus proposed that these genes could either form a pathway directly exapted for symbiosis in Embryophyta or that their genetic interactions evolved along with the symbiotic habit. All four genes were found in *Zygnema*: DMI2/SYMRK pro-ortholog (Zci_05951, DMI1/POLUX (Zci_12099), DMI3/CCaMK (Zci_01672) and IPD3/CYCLOPS (Zci_13230). Co-expression analyses show that these genes likely belong to different modules (clusters 134, 78, 172, and 159, respectively). Given the diversity in putative functions of these additional genes, it is not possible to propose a function for these clusters. The lack of co-expression between these genes in Zygnematophyceae supports the hypothesis that the evolution of symbiosis in Embryophytes recruited genes from diverse pathways rather than directly co-opting an existing pathway into a new function (Delaux et al. 2012).

### Plant-microbe interactions and calcium signaling

The first layer of plant responses to microbes hinges on their perception by receptors of extracellular or intracellular signals (Wang et al. 2019); some of the receptors also act in the perception of mutualistic partners (Plett and Martin, 2018). When encountering pathogens, depending on the signal and recognition either pattern-triggered (PTI) or effector-triggered (ETI) immune responses are initiated. PTI and ETI have been traditionally separated; yet they share many signaling pathways and downstream responses (Cunha et al. 2006, Wang et al. 2019, Yuan et al. 2021). In PTI, molecules such as chitin or flagellin are first recognized by pattern-recognition receptors (PRRs), whereas ETI is triggered by pathogen isolate-specific effectors recognized by nucleotide-binding LRR receptors (NLRs). Typical immune responses are callose deposition and ROS production. In *Zygnema*, we observed LRR domain-containing genes often co-expressed with two callose synthase homologs (CALS; Zci_11575, Zci_11576; cluster 117), SOBER1/TIPSY1, which suppress ETI (Zci_13211; cluster 163), and a TOM2B homolog (Zci_03483; cluster 107), which in *Arabidopsis thaliana* is associated with multiplication of tobamoviruses (Tsujimoto et al. 2003). LRR proteins also co-express with immunity-associated receptor-like protein kinases (clusters 107, 128, 163) and a ROP-activating protein RenGAP (Zci_12487; cluster 107). Cluster 163 is mostly deployed under UV and other stresses and shows coexpression of LRR and SOBER1/TIPSY1 with a protein of the photosystem II assembly (LPA1; Zci_08073) and a plastidal protease (EGY; Zci_01876). Overlap of some degree of defense and UV-B stress response is shown in land plants (VandenBussche et al. 2018). In cluster 128, which is deployed most strongly under desiccation conditions at 4°C, some LRR proteins have been involved in cell division (BAK1; Zci_06073 and SRF3; Zci_03265). An interesting pattern is the frequent co-expression of LRR protein-encoding and calcium signaling genes (Fig. 4A). For example, the calcium sensor and kinase (CPK; Zci_12352) in cluster 128 or CDPKs in cluster 117; cluster 117, showcasing the “plant perceptron” concept, also features PP2Cs and LRR proteins (in fact, the most connected node is a LRR protein). Indeed, calcium signaling has recently been proposed to link plant PTI and ETI (Jacob et al. 2021; Bjornson & Zipfel, 2022) but is also important in mutualistic interactions (Plett and Martin, 2018). Overall, this highlights the interconnectivity of the signaling cascades in *Zygnema*.

### Ancestry and diversity of *Zygnema* MADS-box genes

In flowering plants, MADS-box genes control many developmental processes, from root to flower and fruit development. Of special relevance is a lineage of Type II MADS-box genes termed MIKC-type genes, which encode transcription factors with a characteristic domain-structure that includes a keratin-like (K) domain that facilitates dimerization and enables tetramerization of these transcription factors, yielding Floral Quartet-like Complexes (FQCs) (Puranik et al., 2014; Theißen et al., 2016). The increase and diversification of these factors during land plants evolution is tightly associated with the establishment of evolutionary novelties, especially in seed plants (Theißen et al., 2016).

We found one MADS-box gene each in the *Zygnema* genomes, which are closely related and do not encode K domains (**Figure S16**). In transcriptomic data (One Thousand Plant Transcriptomes Initiative, 2019, Sayers et al., 2021), however, we found MADS-box genes encoding a K domain in other Zygnematophyceae including a *Zygnema* species, which form a separate clade. This suggests the presence of two MADS-box gene in the Zygnematophyceae ancestor: (i) an ancestral Type II that did not acquire the K-domain yet and is thus very likely unable to form FQCs, and (ii) the MIKC-type, for which in one case FQC formation has already experimentally been demonstrated in vitro (Rümpler et al., 2022). The clade of the K-domain encoding genes was apparently lost in the *Zygnema* species sequenced here (**Figure S16**). The ancestor of Zygnematophyceae thus had a higher basic diversity of Type II MADS-box genes than embryophytes, but the number of MADS-box genes in *Zygnema* is similarly low than in other streptophyte algae (Tanabe et al., 1995; Nishiyama et al., 2018), indicating that the boost of MADS-box genes is a “synapomorphy” of land plants.

### Deep evolutionary roots of phytohormone biosynthesis and signaling pathways

Phytohormone biosynthesis and signaling networks have deep evolutionary roots. Except for gibberellins and jasmonic acid that likely originated in land plants (Bowman et al. 2019), all other phytohormone pathways were present in green lineage ancestors. Looking at the number of phytohormone-associated genes, land plants have more homologs than algae, as expected for their more complex signaling pathways (Wang et al., 2015), and Zygnematophyceae are overall similar to other streptophyte algae (**Figure 5**). We therefore explored how phytohormone-related genes are woven into *Zygnema*’s co-expression networks. It must be noted that the identification of homologs known to be involved in phytohormone biosynthesis and signaling in land plants cannot account for pathway variations that might happen in algae.

**Figure 5.**
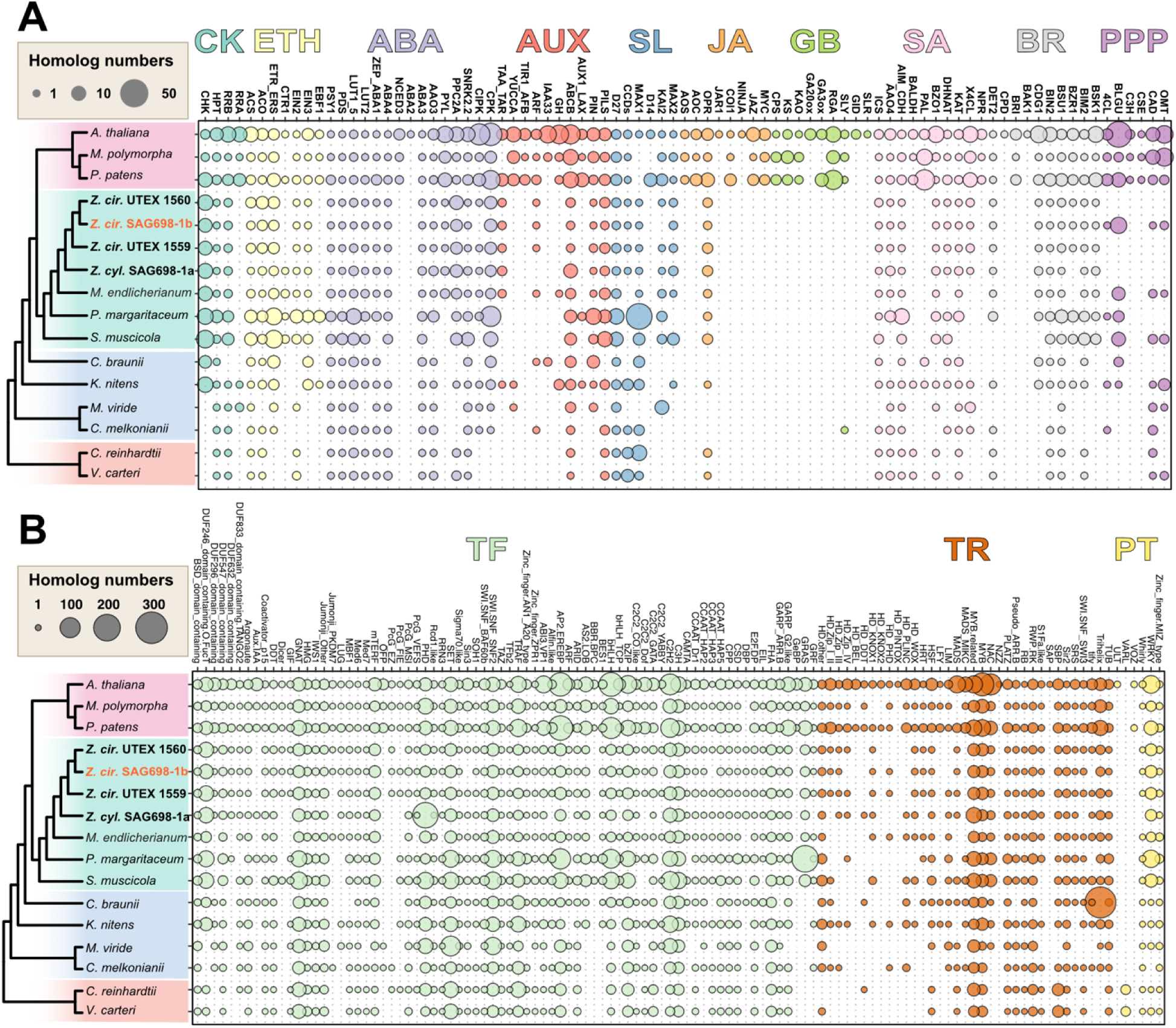
Phylogenetic distribution of (A) proteins involved in phytohormone biosynthesis, signaling, and phenylpropanoid biosynthesis. and (B) transcription factors. CK, cytokinin; ETH, ethylene; ABA, abscisic acid; AUX, auxin; SL, strigolactone; JA, jasmonic acid; GB, gibberellic acid; SA, salicylic acid; BR, brassinosteroids; PPP, phenylpropanoid; TF transcription factors; TR transcriptional regulators; PT putative transcription-associated proteins. For phytohormone-related proteins, homolog numbers were inferred from maximum likelihood gene family trees estimated from significant BLASTP hits (E-value<1e-6) using *Arabidopsis* queries. Note that the high number of homologs found in *Penium margaritaceum* are likely due to the large genome of 3.6 Gb and >50K annotated proteins. Transcription factors were identified by TAPscan.

#### Cytokinin

Cytokinin is involved in plant growth and stress. In *Zygnema*, one out of four co-orthologs of the *A. thaliana* cytokinin receptor AHK2-4 contains the CHASE domain required for binding cytokinin (Zci_13126.1). Other CHASE-domain-containing histidine kinase (CHK) homologs result from more ancient duplications in streptophytes, where homologs previously identified *Spirogyra* and *Mougeotia* likely belong (de Vries et al. 2020). Two *Zygnema* CHKs were found to co-express with cell division and development genes such as cell cycle regulators, plate maturation factors, cell wall-relevant cellulose synthase, DNA replication factors and the only identified MADS-box gene (cluster 22, 76). Among response regulators, we found that the *Zygnema* strains lack Type-A *Arabidopsis* response regulators (RRA) and have only two Type-B *Arabidopsis* response regulators (RRB), and their involvement in cytokinin signaling of Zygnematophyceae is unclear; in fact RRA showed low responsiveness to exogenous cytokinin in *Spirogyra pratensis* (de Vries et al. 2020). Moreover, the paucity of RRA/RRB genes in non-vascular plants supports the notion that this cytokinin-mediated induction of response regulators might be a feature of seed plants (Brenner and Schmülling 2015). Yet, RRB homologs are co-expressed with genes involved in cell division (cluster 151) and stress response (cluster 79).

#### Ethylene

In accordance with its deep evolutionary roots (Ju et al., 2015; Bowman et al., 2019), the chassis for ethylene biosynthesis and signaling is present in Zygnematophyceae. EBF1, which binds EIN3 and is responsible for the last step in the signaling cascade, shows an almost land-plant specific distribution, with the exceptions of EBF1, which is not found in *Zygnema* but homologs are found in *Penium* and *Klebsormidium*. It is not yet known whether these algal homologs bind EIN3 in physiological conditions. In land plants, ethylene triggers cell wall matrix modification, reduces chlorophyll biosynthesis and photosynthesis, and activates abiotic stress responses. Exogenous ethylene treatment in the zygnematophyte *Spirogyra* triggered similar transcriptomic responses and cell elongation (Van de Poel et al., 2016) and *Spirogyra* genes can complement *Arabidopsis* ethylene signaling KO lines (Ju et al., 2015). Several abiotic stress conditions have been shown to stimulate cell elongation in an ethylene-dependent manner. Congruently, cluster 110, deployed under stress (high light, osmotic stress, UV, high pH), shows ACS—a key regulator of ethylene biosynthesis—co-expressed with cell division and circadian clock genes, genes related to photosystem II assembly, starch metabolism, and immunity. Two homologs of ethylene receptor ETR1 co-express with genes involved in cell and plastid division, the PIN auxin effector transporter, calcium signaling SIEL, a regulator of plasmodesmata intercellular trafficking (cluster 119). Cluster 31, deployed largely under darker growing conditions (2h and 6h) and to a lesser extent under high salt or high pH, shows co-expression of an ethylene-responsive factor with genes involved in cell wall remodeling (transferases and beta-glucosidases), photosynthesis, and stress responses (calcium signaling genes, subtilisin-like protease). These clusters illustrate well the effects of ethylene in modifying the cell wall matrix, downregulating photosynthesis, and activating abiotic stress responses as known from land plants.

#### Abscisic acid

Major aspects of the ABA signaling network are conserved across land plants (Cuming et al., 2007; Umezawa et al., 2010; Eklund et al., 2018). The four new *Zygnema* genomes contain a complete set of homologous genes to the ABA signaling cascade, including *Pyrabactin Resistance 1 /PYR1-like /Regulatory Component of ABA Receptor* (PYR/PYL/RCAR) receptors, as previously identified (de Vries et al. 2018; Cheng et al., 2019). Functional data showed that Zygnematophyceae PYL regulates downstream phosphatases in an ABA-independent manner (Sun et al., 2019). Among ABA biosynthetic genes, two key *A. thaliana* enzymes lacked homologs outside of land plants (NCED3, Nine-cis-Epoxycarotenoid dioxygenase 3 and ABA2, SHORT-CHAIN DEHYDROGENASE/ REDUCTASE 1). Yet, we detected about 1.01±0.13 ng/g ABA in SAG 698-1b cultures by LC-MS (**Figure S13**), suggesting that these reactions occur by alternative routes, perhaps *via* an ABA1-independent biosynthetic pathway starting upstream of zeaxanthin as suggested by Jia et al. (2022). An interesting observation is the expansion of clade A PP2Cs in *Zygnema* (5 homologs), akin to the 9 homologs found in *Arabidopsis*. The expansion of PP2Cs was also detected by orthogroup expansion analyses (OG 548 and OG 830), but it must be noted that the high PP2C numbers in *Arabidopsis* and *Zygnema* derive from independent duplications.

#### Auxin

Auxin is the major morphogenic phytohormone. Its polar distribution—largely based on the PIN proteins (Adamowski and Friml, 2015)—leads to gradients along plant bodies, shaping various developmental processes in all land plants (e.g., Friml et al., 2003; Weijers and Wagner, 2016). While all components of the canonical auxin signaling pathway likely first came together in land plants (Flores-Sandoval et al., 2018; Mutte et al., 2018; Martin-Arevalillo et al., 2019), PIN-mediated polar auxin transport might have emerged earlier (Żabka et al., 2016); that said, not all streptophyte algal PINs localize polarly (Skokan et al., 2019; Vosolsobě et al. 2020). While TAA homologs involved in auxin biosynthesis are found in several streptophyte algae including *Zygnema*, no YUCCA homologs were found in any Zygnematophyceae. In fact, embryophyte YUCCAs might have been acquired by HGT from bacteria (Yue et al. 2014). Similarly, several signaling proteins (e.g., TIR, AUX, GH) are also absent from *Zygnema* and distant homologs (never co-orthologs) are sometimes present in other streptophyte algae. These patterns suggests that auxin signaling might be land plant specific.

#### Strigolactones

Strigolactone biosynthesis has been well established in land plants (Proust et al. 2011, Delaux et al. 2012, Waters et al; 2017, Kodama et al. 2022). The streptophyte alga *Nitella* has been shown to produce and respond to strigolactone although the actual biosynthetic pathway might differ from the canonical one of embryophytes (Bowman et al. 2019, Nishiyama et al. 2018). Indeed, no clear orthologs of carotenoid cleavage dioxygenases (CCD7 and CCD8; **Figure S10**), key biosynthetic enzymes, were found in its transcriptome nor in the genome of the close relative *Chara braunii* (Nishiyama et al. 2018). Strigolactones were also detected in some chlorophyte algal species for which no genomes data are available (Smýkalová, I. et al. 2017). We find close homologs of strigolactone biosynthesis genes in all *Zygnema* genomes (including CCD7 orthologs) as well as across the whole Chloroplastida. Yet, all CCD7-8 homologs outside of embryophytes do not show conserved amino acid motifs proposed to be important for substrate specificity (Messing, et al. 2010; **Figures S11, S12**). The angiosperm strigolactone sensor D14 has likely evolved in the seed or vascular plant ancestors through neofunctionalization of karrikin-sensing F-box proteins (Kodama et al. 2022) and thus no homologs are found in any algae. Our results consolidate the hypothesis that strigolactone biosynthesis and signaling differ between embryophytes and streptophyte algae.

### Microexons have evolved during plant terrestrialization

Microexons are very short (1∼15 basepairs (bp)) exons that can be evolutionarily conserved and crucial for gene functions in plants (Yu et al., 2022). To study how microexons have evolved in streptophytes, we predicted 45 microexon-tags in 16 plant genomes using *MEPmodeler* (https://github.com/yuhuihui2011). Land plants typically have more than 20 of 45 microexon-tag clusters. In Zygnematophyceae genomes, we found 10-20 microexon-tag clusters (only 6 clusters in *P. margaritaceum* probably due to the fragmented genome assembly, **Table 1**), <5 in other streptophytes, and none in Chlorophyta (**Figure 6**). Zygnematophyceae and land plants have the highest number of microexons. For example, a 1 bp microexon of cluster 2 was found in Vps55 (Vacuolar protein sorting-associated protein 55, Zci_4861) (**Figure 6B**), and two adjacent microexons (5 and 12 bp) of cluster 7 were found in a Peptidase M1 family gene (Zci_04270) (**Figure 6C**), which are all supported by RNA-seq read mapping. Aligning the orthologous genes of the Peptidase M1 family gene (Zci_04270) across multiple genomes, we found that the two adjacent microexons are in the context of a 108 bp coding region spanning five exons in the *Arabidopsis* gene (AT1G63770.5). The five-exon structure of this coding region is only conserved in land plants and *Zygnema* (**Figure 6D**). In *M. endlicherianum* the last two exons (including the 112 bp) are fused, while in earlier branching algae the five-exon structure exists as two or three exons with the two adjacent microexons (5 and 12 bp) of cluster 7 are always fused. It appears that during terrestrialization, at least for this Peptidase M1 family gene, there is a gradual intronization process that creates the higher abundance of microexons in land plants.

**Figure 6:**
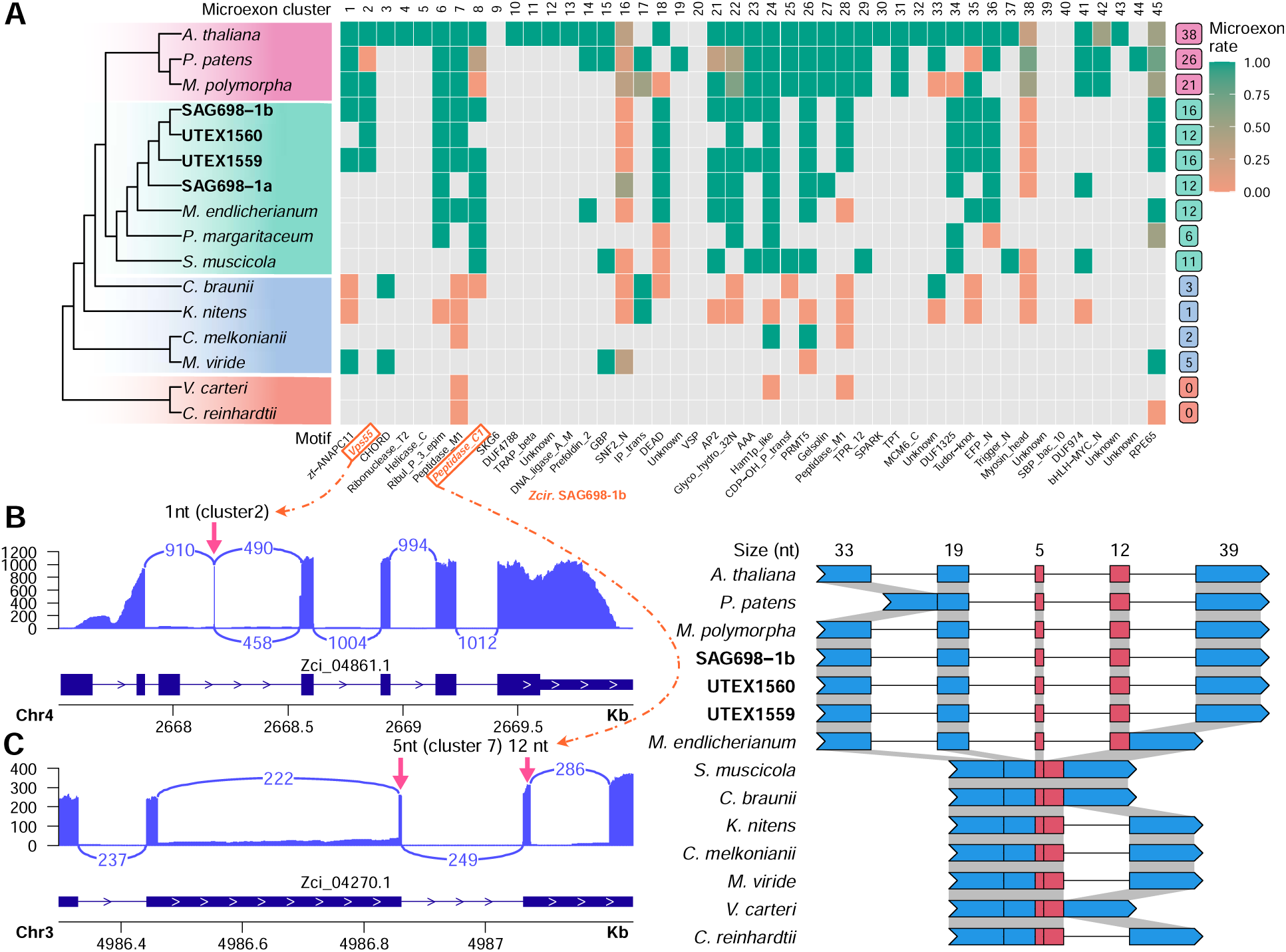
Microexon prediction in 16 plant and algae genomes. (**A**) Heatmap of 45 conserved microexon-tags predicted by MEPmodeler with default parameters. Microexon rate is the rate of true microexons among all predicted results in the cluster, e.g., green cell indicates that 100% microexons with both two flanking introns are present, red indicates all microexon sequences are parts of large exons and none of them could be considered as microexons, and the others are between 0 and 1. A gray cell indicates missing data (a microexon-tag could not be found). Numbers on the right column indicate the predicted clusters containing at least one true microexon (see Yu et al., 2022 for more detail). (**B**) RNA-seq evidence of the 1 bp microexon in Cluster 2. (**C**) RNA-seq evidence of 1 bp microexon in Cluster 2 two adjacent microexons (5 and 12 bp) in Cluster 7. In **B** and **C**, the RNA-seq of condition p881sControl2 was used; RNA-seq read depth and gene annotation are shown; the number in each intron indicates the junction reads and the arrows point to microexons. (**D**) Exon-intron structures of microexon-tag Cluster 7 in 14 plant genomes. The structure was predicted by relaxing the stringency in *M. viride* genome and by doing TBLASTX search in *S. muscicola* genome (all the three copies are intronless in this microexon-tag), respectively. The others are predicted with default parameters.

## Conclusion

One of the defining features of land plants is the plastic development of their multicellular bodies, ever adjusting to altered environmental conditions. We here generated chromosome-level genome assemblies for a representative of the filamentous algal sister lineage to land plants and performed exhaustive co-expression network analyses. Our data underscore earlier notions of a deep evolutionary origin of important plant signaling cascades for acclimation to environmental cues and suggest a deep conservation of interconnections with regulation of growth. The plant perceptron connected environmental input with development before embryophytes began their conquest of land.

## Acknowledgements

This work was funded by the U.S. National Science Foundation (NSF) CAREER award (DBI-1652164), the Nebraska Tobacco Settlement Biomedical Research Enhancement Funds as part of a start-up grant of the University of Nebraska Lincoln, the Research & Artistry Award of Northern Illinois University, the Joint Genome Institute Community Science Program (CSP), the United States Department of Agriculture (USDA) award (58-8042-9–089), the National Institutes of Health (NIH) awards (R21AI171952) and (R01GM140370) all to Y.Y., and by the German Research Foundation grant 440231723 (VR 132/4-1) to J.d.V., TH417/12-1 to GT and FR, and 440540015 (BU 2301/6-1) to H.B. within the framework of the Priority Programme “MAdLand – Molecular Adaptation to Land: Plant Evolution to Change” (SPP 2237), and grant 410739858 in the frame of the project CharMod to K.v.S., as well as RE 1697/16-1 (CharMod) and 18-1 (CharKeyS) to S.A.R. J.d.V. further thanks the European Research Council for funding under the European Union’s Horizon 2020 research and innovation programme (Grant Agreement No. 852725; ERC-StG “TerreStriAL”). The work was further supported by Austrian Science Fund project P34181-B to A.H.; A.D., A.D.A., M.J.B., and J.M.S.Z. are grateful for being supported through the International Max Planck Research School (IMPRS) for Genome Science; J.M.R.F.-J. and T.P.R. gratefully acknowledge support by the Ph.D. program “Microbiology and Biochemistry” within the framework of the “Göttingen Graduate Center for Neurosciences, Biophysics, and Molecular Biosciences” (GGNB) at the University of Goettingen. The work (proposal: 10.46936/10.25585/60001088) conducted by the U.S. Department of Energy Joint Genome Institute (https://ror.org/04xm1d337), a DOE Office of Science User Facility, is supported by the Office of Science of the U.S. Department of Energy operated under Contract No. DE-AC02-05CH11231. P-M.D is supported by the project Engineering Nitrogen Symbiosis for Africa (ENSA) currently funded through a grant to the University of Cambridge by the Bill & Melinda Gates Foundation (OPP1172165) and the UK Foreign, Commonwealth and Development Office as Engineering Nitrogen Symbiosis for Africa (OPP1172165), by the “Laboratoires d’Excellence (LABEX)” TULIP (ANR-10-LABX-41)” and by the European Research Council (ERC) under the European Union’s Horizon 2020 research and innovation program (Grant agreement No. 101001675).

## Authors’ contributions

A.H., J.d.V., Y.Y. secured funding.

M.L., J.M.A., J.d.V., Y.Y. provided resources and materials.

A.H., J.d.V., Y.Y. provided supervision.

J.d.V., Y.Y. conceptualized the study.

J.d.V., K.B., Y.Y. performed project administration.

X.F., E.F., W.S.G., T.D., J.M.R.F.-J performed experimental work, including algal culturing, DNA/RNA extraction, microscopy.

J.Z., X.F., and Y.Y. generated and annotated the draft genomes and transcriptomes. I.I., S.d.V., J.d.V. analyzed phenylpropanoid metabolism-related genes.

R.D.H., I.V.G. coordinated genomes deposition/annotation in PhycoCosm.

L.B. performed and evaluated Illumina assemblies.

K.B., C.D. coordinated Illumina sequencing for UTEX 1559 and UTEX 1560.

L.G., F.R., and G.T. annotated and analyzed MADS-box genes

I.I., J.M.S.Z., T.P.R., A.D.A., A.A., A.M., P.M.-D. analyzed phytohormone-related genes.

J.B.A., N.K, A.M. performed ABA measurements.

H.B. analyzed data on cell division.

F.W.-L. analyzed photoreceptor genes.

K.v.S. helped in the initial phase of the project in strain purifications and mating experiments.

J.K. and P.-M.D. performed phylogenetic analyses of symbiotic genes

M.J.B., A.A. analyzed RNA-seq data.

X.F., J.Z., B.Z., T.L., O.N., I.I., J.d.V., Y.Y. analyzed data and generated data figures and tables.

J.Z., X.W., N.F.-P., S.A.R., Y.Y. conducted the WGD analysis.

S.A.R., R.P. performed TAPscan analysis.

S.A.R., F.H. performed contamination analyses.

N.R. and C.Pe. established the protocols and performed the chromosome stainings, staining

X.F., J.P.M. performed the organellar genome assembly and analysis.

X.F., J.Z., B.Z., J.H., Y.Y. performed the cell wall and HGT analysis.

H.Y., C.Z. performed the microexon analysis.

Z.A., M.M. built the *Zygnema* gene co-expression database.

X.F., I.I., J.Z., J.d.V., Y.Y. wrote the original draft.

All authors helped discussing the results and writing the paper.

## METHODS

### Resource table

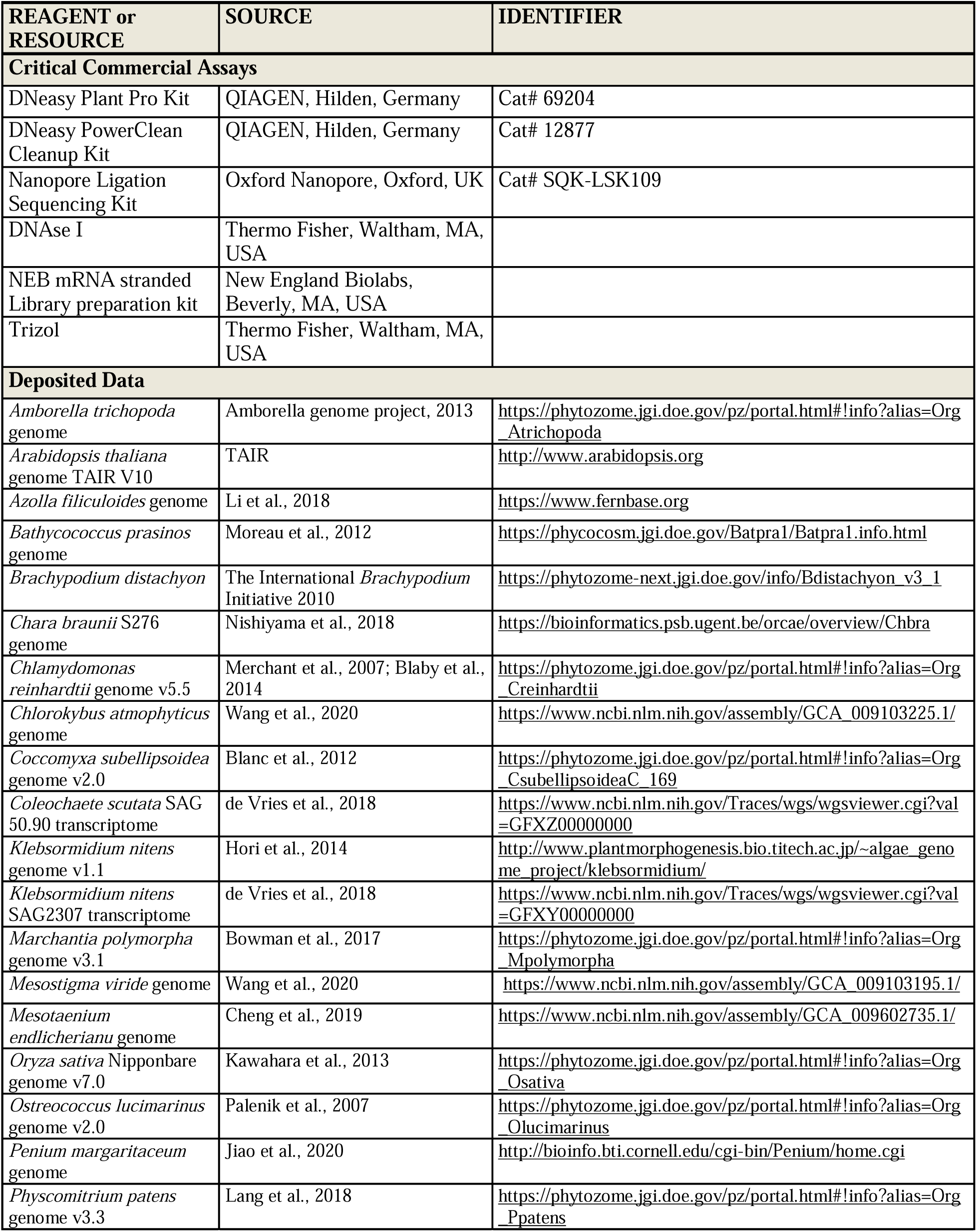
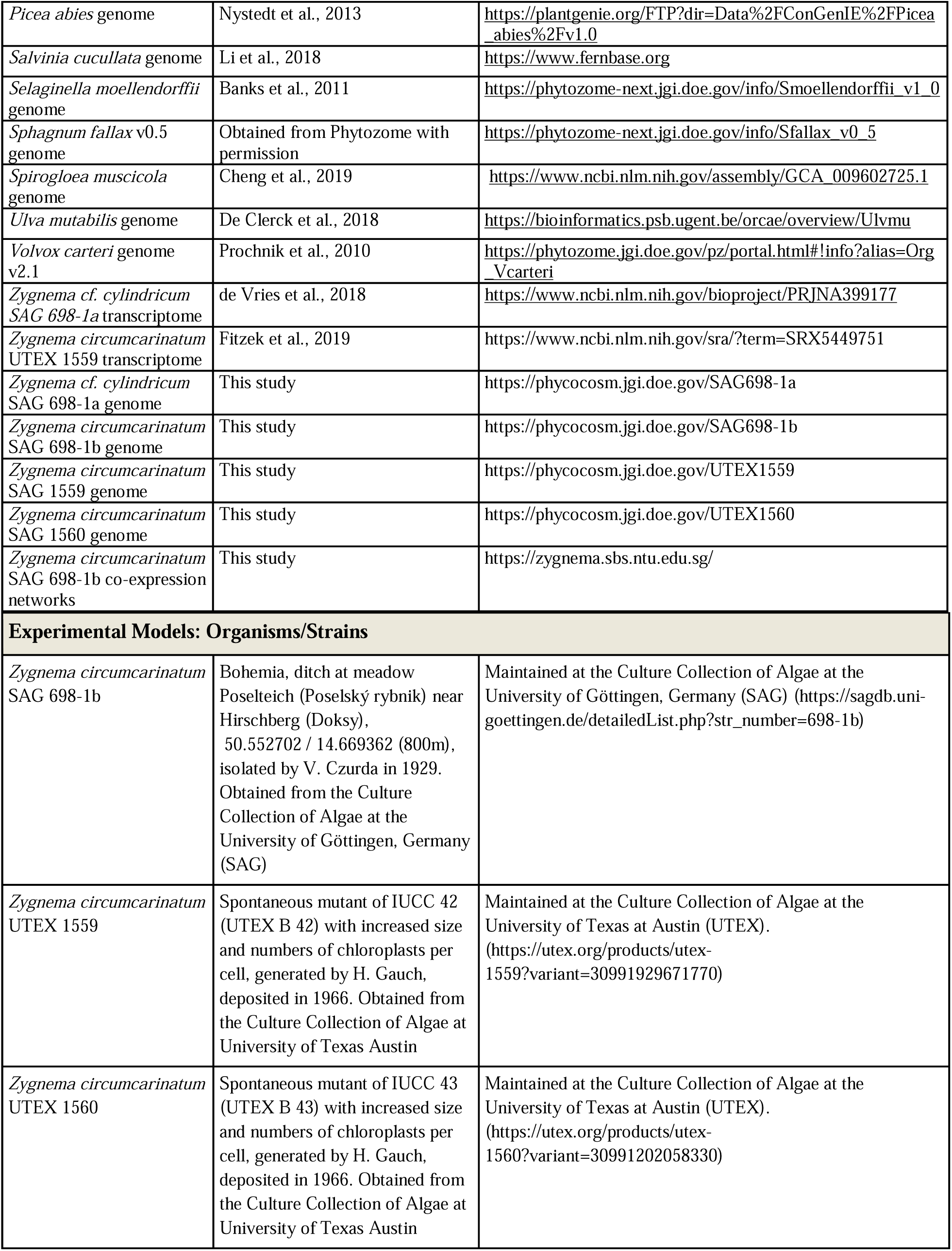
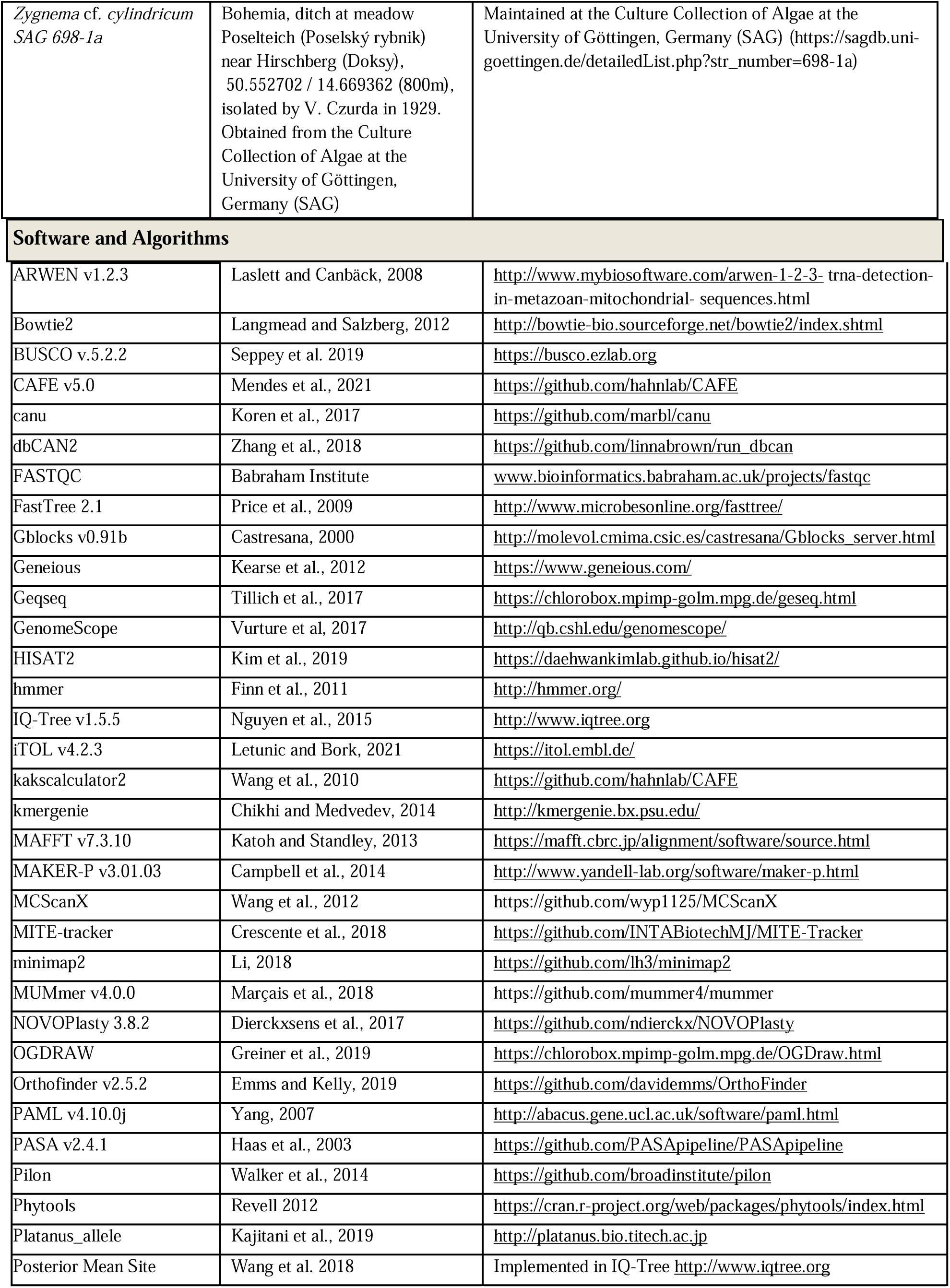
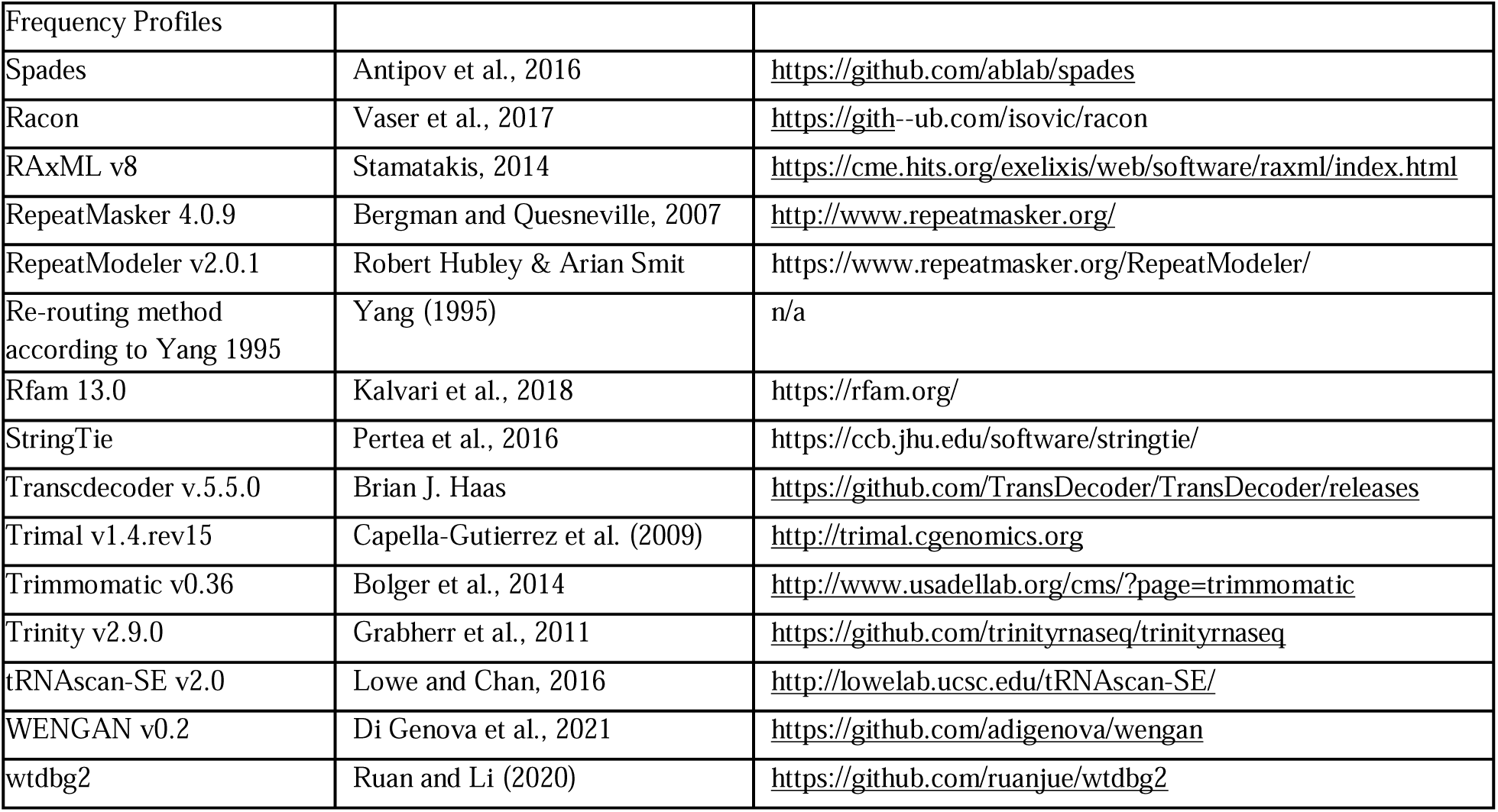

### Resource availability

#### Lead contact

Further information and requests for resources and reagents should be directed to and will be fulfilled by the lead contacts, Jan de Vries and Yanbin Yin.

### Materials availability

This study did not generate new unique reagents.

### Data and code availability

The four *Zygnema* genomes, raw DNA reads of all four strains, and rRNA-depleted RNA-seq of SAG 689-1b can be accessed through NCBI BioProject PRJNA917633. The raw DNA read data of UTEX1559 and UTEX1560 sequenced by the Joint Genome Institute can be accessed through BioProjects PRJNA566554 and PRJNA519006, respectively. RNA-seq data of UTEX1559 can be accessed through BioProject PRJNA524229. Poly-A enriched RNA-seq data of SAG 689-1b can be accessed through BioProject PRJNA890248 and the Sequence Read Archive (SRA) under the accession SRR21891679 to SRR21891705. *Zygnema* genomes are also available through the PhycoCosm portal (Grigoriev et al., 2021; https://phycocosm.jgi.doe.gov/SAG698-1a; https://phycocosm.jgi.doe.gov/SAG698-1b; https://phycocosm.jgi.doe.gov/UTEX1559; https://phycocosm.jgi.doe.gov/UTEX1560). No original code was used; all computational analyses were performed with published tools and are cited in the Methods section.

### Organisms

#### Algal strains

*Z. circumcarinatum* SAG 698-1b and *Z. cf. cylindricum* SAG 698-1a were obtained from the Culture Collection of Algae at Göttingen University (SAG) (https://sagdb.uni-goettingen.de). *Z. circumcarinatum* UTEX 1559 and UTEX 1560 were obtained from the UTEX Culture Collection of Algae at UT-Austin (https://utex.org/). The algae were cultured with Bold’s basal media (BBM) or modified Bold’s basal media (MBBM), supplemented with 0.02% L-arginine, 0.1% peptone and 0.5% sucrose (Feng et al., 2021). The filaments were grown for two or three weeks on a rotary shaker platform (Fermentation Design, 125 rpm) in Plant Growth Chambers (Conviron PGR15) with conditions: 16:8 of light: dark cycle, 20°C, ∼50 µmol of quanta m-2 ·s-1 (Feng et al., 2021). Some cultures were also maintained on 1% agar or liquid MBBM in the same conditions.

### Chromosome staining

*Zygnema circumcarinatum* SAG 698-1b were collected at the end of the light phase and pre-fixed in 2 mM 8-hyroxyquinoline (Roth, Karlsruhe, Germany) for 1h at room temperature (RT) and 1h at 4°C and then fixed in Carnoy’s fluid (3 parts 100% ethanol, 1 part 100% acetic acid) for 12h at 4°C. Then, the totally bleached cells were stained in 1% acetocarmine (in 99% acetic acid, Morphisto, Offenbach, Germany) by boiling for 5 min over open flame. Samples were then transferred to microscope slides in 1% acetocarmine and examined at a Zeiss Axiovert 200 M microscope (Carl Zeiss, Jena, Germany) and images captured with an Axiovision HRc camera (Carl Zeiss, Jena, Germany). Stacked models of 100-150 images were rendered by Helicon Focus software (HeliconSoft Ltd., Kharkiv, Ukraine) and further processed with ImageJ software (Fiji, open source).

### Chromosome counting of Z. cricumcarinatum SAG 698-1b

Z. strain SAG 698-1b was grown in liquid Bold’s basal medium (BBM) in axenic culture under controlled conditions (20°C, ∼50 µmol photons m-2 s-1). Exposition to a 10:14 h light-dark cycle led to synchronization of the cell cycle. The period where most cell divisions took place was from the last hour of the light cycle to the first 5 h of the dark cycle, as previously reported cell division occurs almost solely in the dark phase in *Zygnema* (Staker, 1971). After a two-week incubation, algal filaments were harvested and pretreated for 1 h at RT, followed by 1 h at 4°C in 2 mM 8-hydroxychinolin (Roth, Karlsruhe, Germany) in darkness, resulting in depolymerization of microtubules and an increased condensation of chromosomes. Cells were fixed in 1:3 glacial acetic acid - ethanol solution (Carnoy’s fluid) for 12 h until all chlorophyll was extracted. Fixed cells were stored in 70% ethanol at −20°C until staining, which was performed by boiling in acetocarmine (99% acetic acid, 1% carmine; Morphisto, Offenbach, Germany) for 5 min. To maximize the visualization of the chromosomes the filaments on the prepared slides were slightly crushed, and stacks of 50 to 100 images per filaments with well stained chromosomes were taken at a Zeiss Axiovert 200 M microscope (100x, 1.3 NA, objective lens) with a Zeiss Axiocam HRm Rev.3 camera (Carl Zeiss, Jena, Germany). Stacked models were rendered by the software Helicon Focus (HeliconSoft Ltd.). Final count of chromosomes was done with ImageJ.

### DNA extraction

Detailed protocols have been published elsewhere (Fitzek et al., 2019) (Orton et al., 2020) (Feng et al., 2021). Briefly, algae were grown for two weeks and harvested using a vacuum filtration with Whatman #2 paper (GE Healthcare 47 mm), washed with distilled water (three times), frozen in liquid nitrogen and stored in −80°C. Frozen algae were lyophilized overnight and total genomic DNA was extracted with DNeasy PowerPlant Pro Kit (Qiagen, Germany) using the following workflow: lyophilized algae were first chopped with a spatula into fine powder and mixed well with beads solution and RNase A. The mixture was homogenized on a vortex adapter at maximum speed for 5 min. Then the DNeasy PowerPlant Pro Kit protocol was followed and the extracted DNA was further purified with DNeasy PowerClean CleanUp Kit (Qiagen, Germany).

Before DNA extraction, to reduce chloroplast and mitochondria derived DNA (up to > 60% of total DNA), a modified nucleus isolation method was used (Zhang et al., 2012). Briefly, fresh algae tissues were grinded into fine powder in liquid nitrogen with precooled mortar and pestle. After that, the powder was transferred into a beaker containing nucleus isolation buffer. This mixture was homogenized well on ice, and then were vacuum filtered with two layers of miracloth (Thermo Fisher Scientific, USA). The remained nuclei were pelleted by centrifugation with speed of x 800 g at 4 °C for 10 min, and extracted with DNeasy PowerPlant Pro Kit (Qiagen, Germany).

After DNA extraction, quality and quantity of purified DNA was evaluated by using 1% agarose gel electrophoresis, NanoDrop 2000/2000c Spectrophotometers, and Qubit 3.0 Fluorometer (Thermo Fisher Scientific).

### Stress treatments and RNA-seq

We subjected *Z. circumcarinatum* SAG 698-1b to 19 growing and stress conditions, after which RNA-seq was obtained for the construction of a gene co-expression network. Stress and RNA-seq experiments were done in three baches. The first batch followed Pichrtová et al. (2014) and de Vries et al. (2018) with modifications. Three-week algae were sub-cultured in 12 flasks of liquid BBM with 0.02% L-arginine and grown for two weeks under standard conditions: 16/8 of light/dark cycle at 20°C and ∼50 µmol of quanta m-2 ·s-1. Then the algae were treated for 24 h under four conditions: (i) 20°C in liquid medium (standard control); (ii) 4°C in liquid medium, (iii) desiccation at 20°C, and (iv) desiccation at 4°C. Four treatments each with three replicates were performed. For desiccation treatments, algae were harvested using a vacuum filtration with Glass Microfiber Filter paper (GE Healthcare, 47 mm) and 20 µl of MBBM was added on the filter paper. Papers with algae were then transferred onto a glass desiccator containing saturated KCl solution (Pichrtová et al., 2014) and the desiccator was sealed with petroleum jelly and placed in the growth chamber under standard culture conditions. Cultures grown in liquid conditions were harvested using a vacuum filtration with Whatman #2 paper (GE Healthcare, 47 mm). After 24 h of treatment, the twelve samples were transferred into 1.5 ml Eppendorf tubes and immediately frozen in liquid nitrogen and stored in −80°C. For the second batch (6 diurnal experiments), the algae were grown with the same control conditions as the above mentioned (16/8 of light/dark cycle, 20°C, ∼50 µmol of quanta m-2 ·s-1), and samples were collected every 4 hours: (v) diurnal dark 2h, (vi) diurnal dark 6h, (vii) durnal light 2h, (viii) diurnal light 6h, (ix) diurnal light 10h, (x) diurnal light 14h. For the third batch (9 stress experiments): SAG 698-1b was pre-cultivated at 20°C, 16:8 hrs light:dark cycle at 90 µmol photons/m2s on a cellophane discs (folia Bringmann, Germany) for 8 days. For certain treatments (NaCl, Mannitol, CadmiumCl) the culture was transferred to a new petri dish where the medium was supplemented with the substances mentioned above. Algae where then subjected to: (xi) 150 µM NaCl (Roth, Germany) for 24h, (xii) 300 mM Mannitol (Roth, Germany) for 24h, (xiii) 250 µM CadmiumCl (Riedel-de Haën AG, Germany) for 24 h, (xiv) dark treatment for 24h, (xv) high light (HL) treatment at 900 µmol photons/m2s for 1h, (xvi) UV-A at 385 nm, 1400 µW/cm2 for 1h, (xvii) high light at 4°C (HL4) at 900 µmol photons/m2s for 1h, (xviii) pH=9 for 24h, and (xix) a corresponding control growth at 20°C on a plate.

### RNA extraction

Frozen algae were lyophilized overnight for RNA extraction with a modified CTAB method described in Chang et al. (1993) and Bekesiova et al. (1999). Specifically, the tissue was chopped with spatula into fine powder, and then 500 µl of CTAB buffer (2% CTAB, 100 mM Tris-HCl pH 8.0, 25 mM EDTA, 2 M NaCl, 2% Polyvinylpyrrolidone, 1% β-mercaptoethanol) was added and mixed well. The mixture was incubated in heating block at 65°C for 15 min. After the tubes cooled down, the solution was extracted with Chlorophorm: Isoamyl alcohol 24:1 twice. The supernatant was precipitated with 0.3 volume of 10 M LiCl that was incubated in − 20°C for 30 min. The pellet was washed with 75% ethanol twice and vacuum dried for 15 min. RNA was resuspended in 50 µl of 0.1% DEPC (diethylpyrocarbonate) water. RNA samples were treated with RNase-Free DNase I (Promega) at 37 °C for 30 min to remove any DNA residue. Quality and quantity of purified RNA was evaluated by using 1% agarose gel electrophoresis, NanoDrop 2000/2000c Spectrophotometers, Qubit 3.0 Fluorometer (Thermo Fisher Scientific) and RNA Integrity Number (RIN) (Agilent).

### Library preparation and sequencing

DNA samples were sequenced at Roy J. Carver Biotechnology Center at University of Illinois at Urbana-Champaign, using Oxford Nanopore and Illumina technologies (Table S1A). Oxford Nanopore DNA libraries were prepared with 1D library kit SQK-LSK109 and sequenced with SpotON R9.4.1 FLO-MIN106 flowcells for 48h on a GridIONx5 sequencer. Base calling was performed with Guppy v1.5 (https://community.nanoporetech.com). Illumina shotgun genomic libraries were prepared with the Hyper Library construction kit (Kapa Biosystems, Roche). Libraries had an average fragment size of 450 bp, from 250 to 500 bp, and sequenced with 2×250 bp pair-end on HiSeq 2500. Additional DNA samples were sequenced at the Genome Research Core in University of Illinois at Chicago and JGI. The Illumina shotgun genomic libraries were prepared with Nextera DNA Flex Library Prep Kit. The libraries had an average fragment size of 403bp and sequenced with 2×150 bp pair-end on HiSeq 4000 (Table S1A). RNA samples were sequenced at the Genome Research Core in University of Illinois at Chicago. The libraries were prepared by rRNA depletion with Illumina Stranded Total RNA kit plus Ribo-Zero Plant (https://www.illumina.com/products/by-type/sequencing-kits/library-prep-kits/truseq-stranded-total-rna-plant.html), and 2×150 bp pair-end sequencing was performed on HiSeq 4000. RNA from the third batch of stress experiments were sequenced at the NGS-Integrative Genomics Core Unit of the University Medical Center Göttingen, Germany. Stranded mRNA libraries were prepared with the Illumina stranded mRNA kit and paired-end sequencing of 2×150 bp reads was carried out on an Illumina HiSeq4000 platform.

RNA-seq data for SAG 698-1a and UTEX 1559 have been previously published (Table S1A).

### Transcriptome assembly

Raw RNA-seq reads (Table S1A) were quality checked with FastQC v.0.11.9 (http://www.bioinformatics.babraham.ac.uk/projects/fastqc/) (Andrews, 2010), trimmed with TrimGalore (https://github.com/FelixKrueger/TrimGalore), and were inspected again with FastQC. All reads were combined, and *de novo* assembled with Trinity version 2.9.0 (Grabherr et al., 2011; Haas et al., 2013).

### K-mer frequency analysis

The trimmed DNA Illumina reads were filtered out with BLASTP using plastomes and mitogenomes from *Zygnema* as references. Remaining (putatively nuclear) were used to predict the best k-mer size by kmergenie (http://kmergenie.bx.psu.edu/) (Chikhi and Medvedev, 2014). The histogram of the best k-mer was then uploaded to GenomeScope for viewing the genome plot (http://qb.cshl.edu/genomescope/) (Vurture et al., 2017) (Table S1B and Figure S2).

### Genome assembly and scaffolding

To assemble the genome of SAG 698-1b, a total of 5.4 Gb (82x) of Oxford Nanopore nuclei DNA reads were assembled with wtdbg (Ruan and Li, 2020) (https://github.com/ruanjue/wtdbg). Assembled contigs were polished by Racon (Vaser et al., 2017) and three iterations of pilon (Walker et al., 2014) with Illumina paired-end reads. The polished genome was scaffolded by Dovetail Genomics HiRise software with Hi-C sequencing data (https://dovetailgenomics.com/). Genome contamination was examined by BLASTX against NCBI’s NR database and contaminated scaffolds were removed.

To assemble the UTEX 1559 genome, an initial assembly was done with SPAdes (Antipov et al., 2016) using Illumina paired-end reads (2×150), three mate-pair libraries (insert size: 3-5 kb; 5-7 kb and 8-10 kb) and Oxford Nanopore reads (Table S1A). Assembled contigs were further scaffolded by two rounds of Platanus (Kajitani et al., 2019) with Illumina paired-end reads (2x 250), three mate-pair libraries (insert size 3-5 kb; 5-7 kb and 8-10 kb) and Oxford Nanopore reads. For the UTEX 1560 genome, Illumina paired-end (2×150 bp) and PacBio HiFi reads were used for assembly with SPAdes and further scaffolded with Platanus. Scaffolds with contaminations were identified by BLASTX against NR and removed. The genomes of UTEX1559 and UTEX1560 were scaffolded by Dovetail Genomics HiRise software with Hi-C sequencing data from SAG 698-1b.

The genome of SAG 698-1a was sequenced with PacBio HiFi long reads (40Gb), Nanopore long reads (4Gb), and Illumina short reads (>100Gb). The k-mer analysis using Illumina reads revealed two peaks in the k-mer distribution, suggesting that SAG 698-1a exists as a diploid organism with an estimated heterozygosity rate of 2.22% (Figure S2). All Illumina short reads and the Nanopore reads were first assembled into contigs using SPAdes. Then, WENGAN was used to assemble HiFi long reads and Illumina paired-end reads (2×150bp) using the SPAdes contigs as the reference. Lastly, the resulting WENGAN contigs were scaffolded and gaps were closed with Platanus-allee using all the Nanopore, HiFi, and Illumina reads to derive a consensus pseudo-haploid genome.

To evaluate the quality of assembled genomes (Table S1D), raw RNA-seq reads, Oxford Nanopore and Illumina DNA reads were mapped to the assembly with hisat2 (Kim et al., 2019), minimap2 (Li, 2018), and bowtie2 (Langmead and Salzberg, 2012), respectively. To assess genome completeness, a BUSCO (Seppey et al., 2019) analysis was performed with the ‘Eukaryota odb10’ and ‘Viridiplantae odb10’ reference sets.

### Repeat annotation

Repetitive DNA was annotated using the homology strategy with repeat libraries generated with RepeatModeler. RepeatModeler integrates RepeatScout, RECON, LTRharvest and LTR_retriever tools (version 2.0.1; http://www.repeatmasker.org/RepeatModeler/) (Flynn et al., 2020). The MITE (Miniature inverted-repeat transposable elements) library was identified with MITE-tracker (Crescente et al., 2018) software. These two identified libraries were combined and incorporated into RepeatMasker (version 4.0.9; http://www.repeatmasker.org/) for repeat annotation.

### Genome annotation

In all four genomes, protein coding genes were predicted by the MAKER-P pipeline (Campbell et al., 2014) which integrates multiple gene prediction resources, including *ab initio* prediction, protein homology-based gene prediction and transcripts-based evidence. First, repetitive elements were masked by RepeatMasker with a custom repeat library generated by RepeatModeler. Rfam with infernal and tRNA-Scan2 were used to analyze non-coding RNA and tRNA. For the transcripts evidence, total of 103,967 transcripts were assembled by Trinity (reference-free) and StringTie (reference-based) with RNA-seq reads. Transcriptome assembly were used for generating a complete protein-coding gene models using PASA. Proteins from *M. endlicherianum, S. muscicola* and *A. thaliana* (TAIR10) were used as homology-based evidence. Then, the resulting protein-coding gene models from the first iteration of MAKER-P pipeline were used as training date set for SNAP and Augustus models, which were fed into MAKER for the second iteration of annotation. After three rounds of gene prediction, MAKER-P combined all the protein-coding genes as the final annotated gene sets.

### Plastome and mitogenome assembly and annotation

NOVOPlasty 3.8.2 (https://github.com/ndierckx/NOVOPlasty) (Dierckxsens et al., 2017) was used to assemble plastomes. The contiguity of assembled plastomes was examined in Geneious software (https://www.geneious.com/) (Kearse et al., 2012) with read mapping. For SAG 698-1b mitogenome assembly, Oxford Nanopore reads were assembled with Canu (Koren et al., 2017), where one long mitogenome contig of 238,378 bp was assembled. This contig was circularized and polished with three rounds of pilon (Walker et al., 2014), that was further corrected with Illumina raw reads and compared with mitogenome of UTEX 1559 (MT040698; Orton et al., 2020) in Geneious. For SAG 698-1a, PacBio HiFi reads were used for the assembly of its mitogenome.

Plastome and mitogenome annotation was performed with GeSeq (Tillich et al., 2017) (https://chlorobox.mpimp-golm.mpg.de/geseq.html). For plastome annotation, BLAT search and HMMER profile search (Embryophyta chloroplast) were used for coding sequence, rRNA and tRNA prediction; ARAGORN v1.2.38, ARWEN v1.2.3 and tRNAscan-SE v2.0.5 were used for tRNA annotion. For mitogenome annotation, Viridiplantae was use for BLAT Reference Sequences. The annotated gff files were uploaded for drawing circular organelle genome maps on OGDRAW (https://chlorobox.mpimp-golm.mpg.de/OGDraw.html) (Greiner et al., 2019).

#### Comparison of *Z. circumcarinatum* genomes (SAG 698-1b, UTEX 1559, UTEX 1560)

Two approaches were used to compare the three genomes (Figure S8). The first approach was based on the whole genome alignment (WGA) by using MUMMER. The parameters “-- maxmatch -c 90 -l 40” were set to align the three genomes and then “-i 90 -l 1000” were set to filter out the smaller fragments. The second approach focused on the gene content comparisons. Orthofinder was used to obtain ortholog groups (orthogroups) from genomes’ annotated proteins. Orthofinder results led to a Venn diagram with unique genes (orthogroups with genes from only 1 genome), cloud genes (orthogroups with genes from only 2 genomes), and core genes (orthogroups with genes from all 3 genomes), which collectively form the pan-genome. Orthofinder could have failed to detect homology between very rapidly evolved orthologous genes, which leads to an under-estimation of core genes. Also, gene prediction may have missed genes in one genome but found them in other genomes. To address these issues, the raw DNA reads of each genome were mapped to the unique genes and cloud genes using BWA. This step was able to push more unique genes and cloud genes to core genes or push some unique genes to cloud genes. The following criteria were used to determine if an orthogroup needed to be re-assigned: (i) the number of reads and coverage calculated by bedtools are >10 and >0.8 for a gene, respectively, and (ii) >60% coverage of genes in the orthogroup find sequencing reads from the other genomes. After this step the final Venn diagram was made (Figure S8F), showing the counts of the final core genes, cloud genes, and unique genes.

### Whole genome duplication (WGD) analysis

To identify possible WGDs, Ks and 4dtv values were calculated for each genome. First, all paralog pairs were identified using RBBH (Reciprocal Best BLAST Hit) method using protein sequences (E-value < 1e-6), following the method described by Bowman et al. (2017). RBBH paralog pairs were aligned with MAFFT (Katoh and Standley, 2013) and the corresponding nucleotide alignments were generated. Using RBBH alignments of paralog pairs, KaKs_Calculator2.0 (Wang et al., 2010b) with the YN model and the calculate_4DTV_correction.pl script were run to calculate Ks and 4dtv values for each alignment, respectively. Ks = 0 and 4dtv = 0 values were filtered. The Ks and 4dtv distributions were fitted with a gaussian kernel density model using the seaborn package. For the SAG 698-1b chromosome-level genome, MCscan (Wang et al., 2012) was run to identify syntenic block regions with default parameters.

### Species phylogeny and divergence time analysis

Sixteen representative genomes were selected, including two chlorophytes (*Volvox carteri, Chlamydomonas reinhardtii*), seven Zygnematophyceae (*Zygnema circumcarinatum* SAG 698-1b, UTEX 1559, UTEX 1560, *Z.* cf. *cylindricum* SAG 698-1a, *Mesotaenium endlicherianum*, *Penium margaritaceum*, *Spirogloea muscicola*), four additional streptophyte algae (*Chara braunii, Klebsormidium nitens, Chlorokybus melkonianii, Mesostigma viride*), two bryophytes (*Marchantia polymorpha, Physcomitrium patens*) and a vascular plant (*Arabidopsis thaliana*). Orthogroups were generated by OrthoFinder version 2.5.2 (Emms and Kelly, 2019) and 493 low-copy orthogroups containing ≤ 3 gene copies per genome were aligned with MAFFT v7.310 (Katoh and Standley, 2013). Gene alignments were concatenated and gaps were removed by Gblocks version 0.91b (Castresana, 2000). Phylogenetic tree was built using RAxML v.8 (Stamatakis, 2014) with the “-f a” method and the PROTGAMMAJTT model, and support with 100 pseudoreplicates of non-parametric bootstrap. The tree was rooted on Chlorophyta.

Using the above methodology, additional phylogenetic analyses were performed with (i) the four *Zygnema* strains and (ii) the seven Zygnematophyceae genomes, in order to obtain a higher number of single-copy loci, 5,042 and 204, respectively (**Figure S7**).

Divergence time estimation was carried out with MCMCTree implemented in the PAML package version 4.10.0j (Yang, 2007). The 493 low-copy orthogroup protein sequence alignment was converted to the corresponding nucleotide alignment for MCMCTree, in which 10 MCMC (Markov Chain Monte Carlo) chains were run, each for 1,000,000 generations (Table S1F). Three calibration were set in the reference tree according to (Morris et al., 2018) on the nodes Viridiplantae (972.4∼669.9 Ma), Streptophyta (890.9∼629.1 Ma) and Embrophyta (514.8∼473.5 Ma).

### Gene family phylogenetic analysis

CAZyme families were identified with dbCAN2 (Zhang et al., 2018) with default parameters (E-value < 1e-10 and coverage > 0.35). Whenever needed, dbCAN2 was rerun by using more relaxed parameters. The experimentally characterized cell wall enzymes were manually curated from the literature (Data S1 and Table S1L). Reference genes were included into the phylogenies to infer the presence of orthologs across the 16 genomes and guide the split of large families into subfamilies. Phylogenetic trees were built by using FastTree (Price et al., 2009) initially, and for some selected families, RAxML (Price et al., 2009) and IQ-Tree (Nguyen et al., 2015) were used to rebuild phylogenies to verify topologies.

### Orthogrup expansion and contraction analysis

We inferred expanded and contracted gene families with CAFE v.5 using orthogroups inferred with Orthofinder v.2.4.0 and the previously inferred time-calibrated species phylogeny. CAFE v.5 was run with default settings (base) using the inferred orthogroups and a calibrated species phylogeny. Two independent runs arrived to the same final likelihood and lambda values. The first eight orthogroups (OG0-7) were excluded from the analysis due to too drastic size changes between branches that hampered likelihood calculation; excluded orthogroups were mostly exclusive to a single *Zygnema* or *Chara* genome and likely represented transposable elements, as judged by results of BLASTP against NR.

### Phytohormones

Proteins involved in phytohormone biosynthesis and signaling were identified by BLASTP against annotated proteomes (e-value<1e-6) using *Arabidopsis* genes as queries. For genes with ubiquitous domains (e.g. CIPK, CPK3, SNRK2, CDG1, BAK1), hits were filtered by requiring BLASTP coverage ≥50% of the query. Significant hits were then aligned (MAFFT auto) and maximum likelihood gene trees were inferred in IQ-Tree using best-fit models and 1000 replicates of SH-like aLRT branch support (‘-m TEST -msub nuclear -alrt 1000’). The final sets of homologs were identified by visually inspecting gene trees and identifying the most taxonomically diverse clade (with high support of SH-aLRT>0.85) that included the characterized *Arabidopsis proteins*. Bubble plot was generated with ggplot2 in R.

### Screening for symbiotic genes and phylogenetic analysis

Symbiotic genes were screened against a database of 211 plant species across Viridiplantae lineage (see Table) using proteins of the model plant *Medicago truncatula* as queries in BLASTP v2.11.0+ (Camacho et al., 2009) searches with default parameters and an e-value < 1e-10. Initial alignments of all identified homologs was performed using the DECIPHER package (Wright, 2015) in R v4.1.2 (R Core Team). Positions with >60% gaps were removed with trimAl v1.4 (Capella-Gutiérrez et al., 2009) and a phylogenetic analysis was performed with FastTree v2.1.11 (Price et al., 2009). Clades corresponding to *M. truncatula* orthologs queries were extracted and a second phylogeny was performed. Proteins were aligned using MUSCLE v3.8.1551 (Edgar, 2004) with default parameters and alignments cleaned as described above. Tree reconstruction was performed using IQ-TRee v2.1.2 (Minh et al., 2020) based on BIC-selected model determined by ModelFinder (Kalyaanamoorthy et al., 2017). Branch supports was estimated with 10,000 replicates each of both SH-aLRT (Guindon et al., 2010) and UltraFast Bootstraps (Hoang et al., 2017). Trees were visualized and annotated with iTOL v6 (Letunic and Bork, 2021). For the GRAS family, a subset of 42 species representing the main lineages of Viridiplantae was selected (see Table) and GRAS putative proteins screened using the HMMSEARCH program with default parameters and the PFAM domain PF03514 from HMMER3.3 (Johnson et al., 2010) package. Phylogenetic analysis was then conducted as described above.

### Screening for CCD homologs and phylogenetic analysis

Annotated proteins from 21 land plant genomes (Amborella trichopoda, Anthoceros agrestis, Anthoceros punctatus, Arabidopsis lyrata, Arabidopsis thaliana, Azolla filiculoides, Brachypodium distachyon, Brassica rapa, Lotus japonicus, Marchantia polymorpha, Medicago truncatula, Oryza sativa, Physcomitrium patens, Picea abies, Pisum sativum, Salvinia cucullata, Selaginella moellendorffii, Sphagnum fallax, Spinacia oleracea, Gnetum montanum, Crocus sativus), 7 streptophyte algal genomes (Spirogloea muscicola, Penium_margaritaceum, Mesotaenium ‘endlicherianum’, Mesostigma viride, Klebsormidium nitens, Chlorokybus melkonianii, Chara braunii, Zygnema circumcarinatum), 6 chlorophyte genomes (Ulva mutabilis, Ostreococcus lucimarinus, Micromonas pusilla, Micromonas sp., Chlamydomonas reinhardtii, Coccomyxa subellipsoidea, Chlorella_variabilis), 5 cyanobacterial genomes (Trichormus azollae, Oscillatoria acuminata, Nostoc punctiforme, Gloeomargarita lithophora, Fischerella thermalis), as well as the transcriptome of Coleochaete scutata (de Vries et al., 2018). The representative A. thaliana protein was used as query for BLASTP searches against the above annotated proteins (E-value < 1e-5). Homologs were aligned with MAFFTv7.453 L-INS-I approach (Katoh and Standley, 2013) and maximum likelihood phylogenies computed with IQ-Tree v.1.5.5 (Nguyen et al., 2015), with 100 bootstrap replicates and BIC-selected model (WAG+R9) with ModelFinder (Kalyaanamoorthy et al., 2017). Functional residue analyses were done based on published structural analysis (Messing et al., 2010), and the alignments were plotted with ETE3 (Huerta-Cepas et al., 2016).

### Phylogeny of MADS-box genes

MADS-domain proteins were identified by Hidden Markov Model (HMM) searches (Eddy, 1998) on annotated protein collections. MADS-domain sequences of land plants and opisthokonts were taken from previous publications (Marchant et al., 2022; Gramzow et al., 2010). MADS domain proteins of other streptophyte algae were obtained from the corresponding genome annotations and transcriptomic data (OneThousandPlantTranscriptomesInitiative, 2019). Additional MADS-domain proteins of Zygnematophyceae were identified by BLAST against transcriptome data available at NCBI’s sequence read archive (SRA) (Sayers et al., 2021). MADS-domain-protein sequences were aligned using MAFFTv7.310 (Katoh and Standley, 2013) with default options. Sequences with bad fit to the MADS domain were excluded and the remaining sequences realigned, and trimmed using trimAl (Capella-Gutierrez et al., 2009) with options “-gt .9 -st .0001”. A maximum likelihood phylogeny was reconstructed using RAxMLv8.2.12 (Stamatakis 2014) on the CIPRES Science Gateway (Miller et al., 2011).

### LC-MS/MS analysis of abscisic acid

Abscisic acid was determined in *Physcomitrium patens* samples using the LC-MS/MS system which consisted of Nexera X2 UPLC (Shimadzu) coupled QTRAP 6500+ mass spectrometer (Sciex). Chromatographic separations were carried out using the Acclaim RSLC C18 column (150×2.1 mm, 2.2µm, Thermo Scientific) employing acetonitrile/water+0.1% acetic acid linear gradient. The mass spectrometer was operated in negative ESI mode. Data was acquired in MRM mode using following transitions: 1) ABA 263.2->153.1 (-14 eV), 263.2->219.1 (-18 eV); 2) ABA -D6 (IS) 269.2->159.1 (-14 eV), 269.2->225.1 (-18 eV); declustering potential was -45 V. Freeze-dried moss samples were ground using the metal beads in homogenizer (Bioprep-24) to a fine powder. Accurately weighted (about 20mg) samples were spiked with isotopically labeled ABA -D6 (total added amount was 2 ng) and extracted with 1.5 ml acetonitrile/water (1:1) solution acidified with 0.1% formic acid. Extraction was assisted by sonication (Elma S 40 H, 15 min, two cycles) and solution was left overnight for completion of extraction. Liquid was filtered through 0.2 µm regenerated cellulose membrane filters, evaporated to dryness upon a stream of dry nitrogen and redissolved in 100 µl extraction solution.

