## Supplementary material for "Chromosome-level genomes of multicellular algal sisters to land plants illuminate signaling network evolution": Data S1

### Data S-1

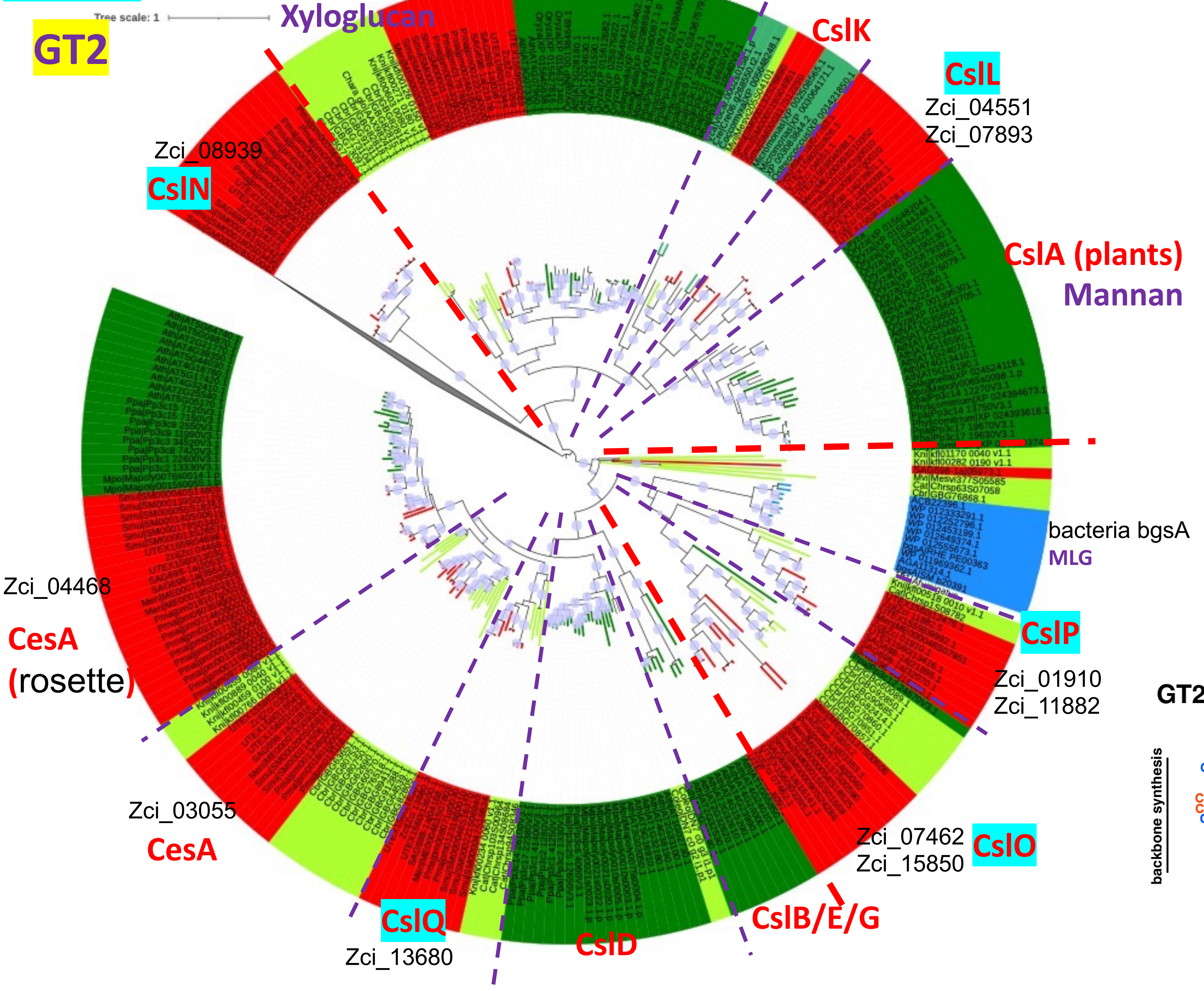


https://itol.embl.de/tree/979822487247861669516257

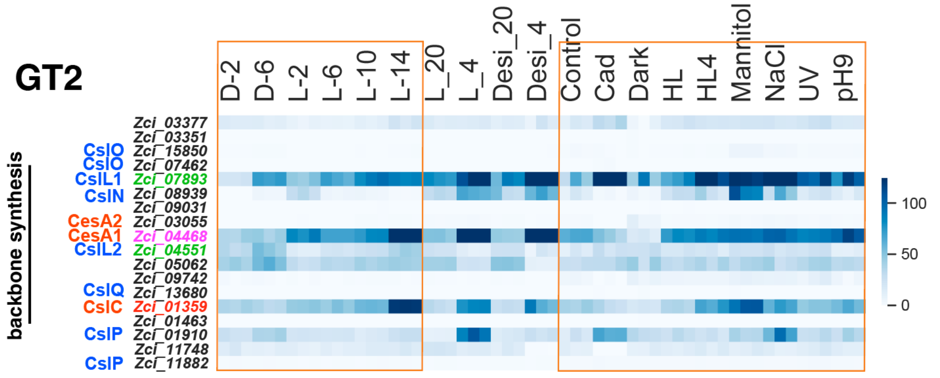

Data S-2

GH9: cellulase

Zci_02466 (subclass C)

Zci_12102 (subclass A)

GH9B7 (At1g75680)
GH9B5 (At1g19940)
subclass B

Zci_03863
Zci_03857
Zci_03866
Zci_01575
(subclass C)

(Urbanowicz et al., 2007)

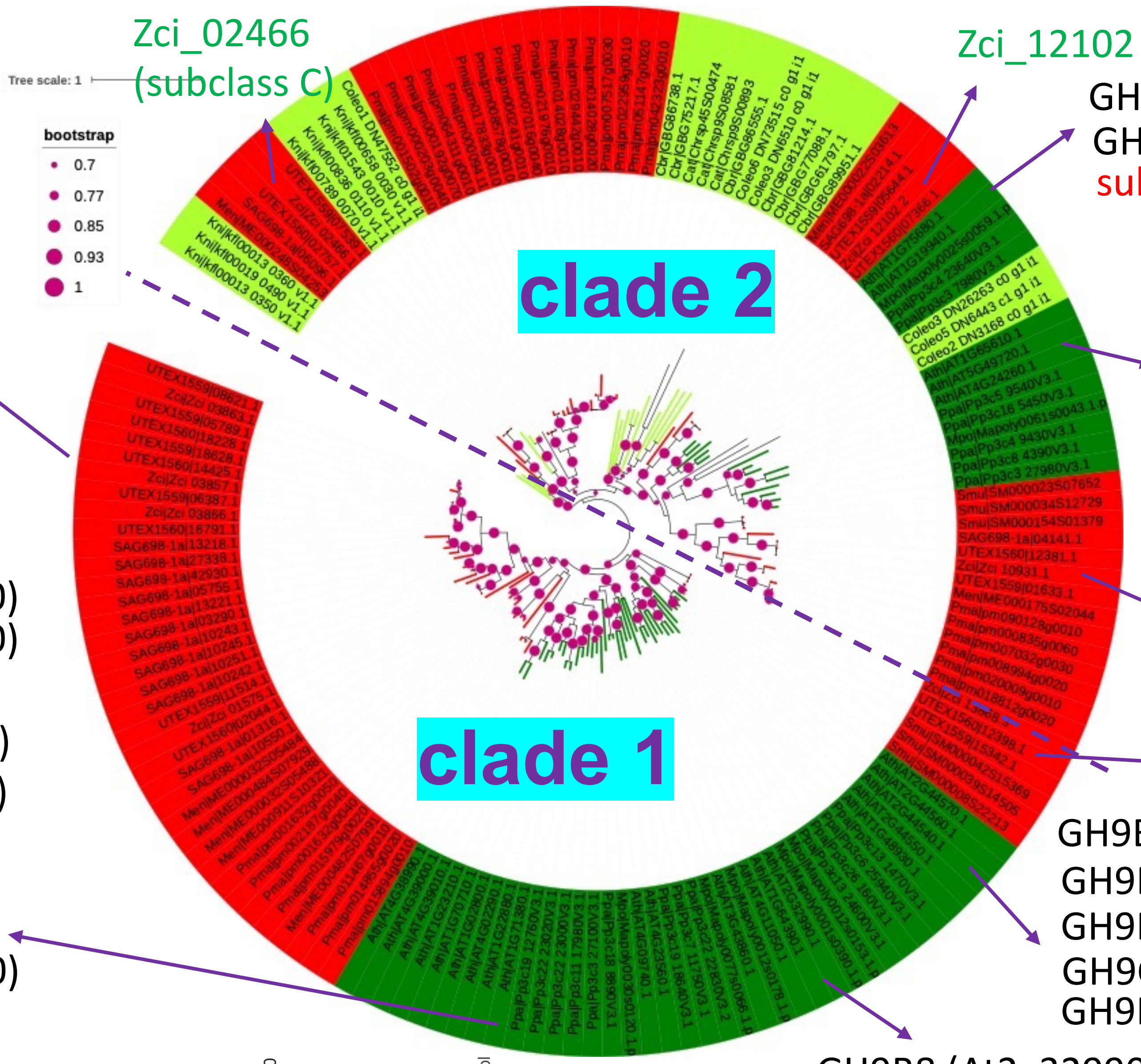


KOR3/GH9A2 (At1g65610)
**KOR1/GH9A1 (AT5G49720)**
KOR2/GH9A3 (AT4G24260)
subclass A

Zci_10931 (subclass A)

Zci_13868 (subclass C)

GH9B14 (At4g09740)
GH9B15 (At4g23560)
GH916 (At4g38990)
GH9B17 (At4g39000)
GH9B18 (At4g39010)
GH9B6 (At1g23210)
GH9B1 (At1g70710)
GH9B2 (At1g02800)
GH9B13 (At1g02290)
GH9B4 (At1g22880)
GH9B3 (At1g71380)
GH9 (At3g43860)
subclass B

GH9B12 (At2g44570)
GH9B11 (At2g44560 )
GH9B9 (At2g44540 )
GH9C1 (At1g48930)
GH9B10 (At2g44550 )

GH9B8 (At2g32990)
GH9C2 (At1g64390)
GH9C3 (At4g11050)
subclass B&C

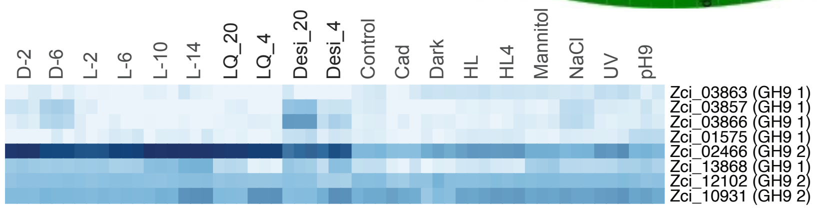


https://itol.embl.de/tree/2081272402221277416673566291

Data S-3a

### Mannan

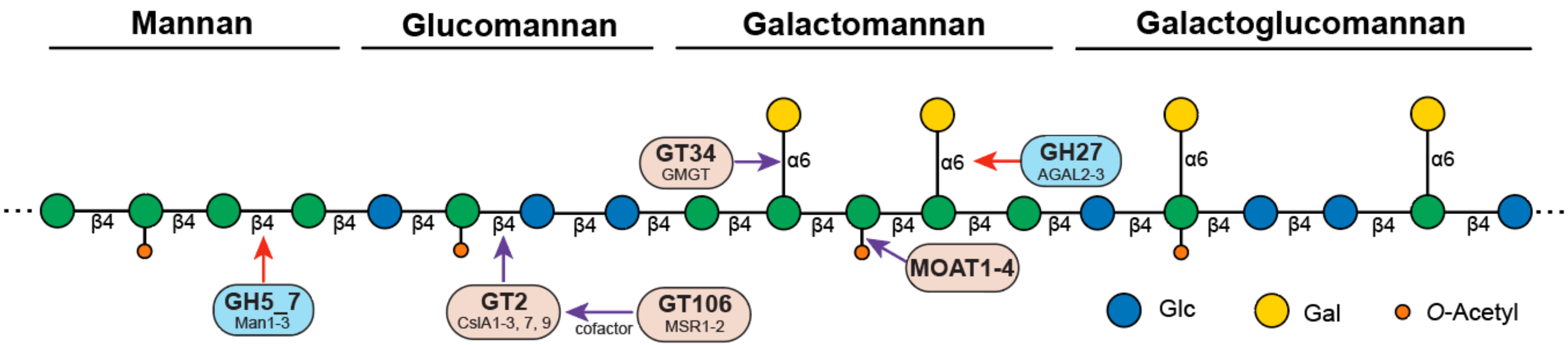

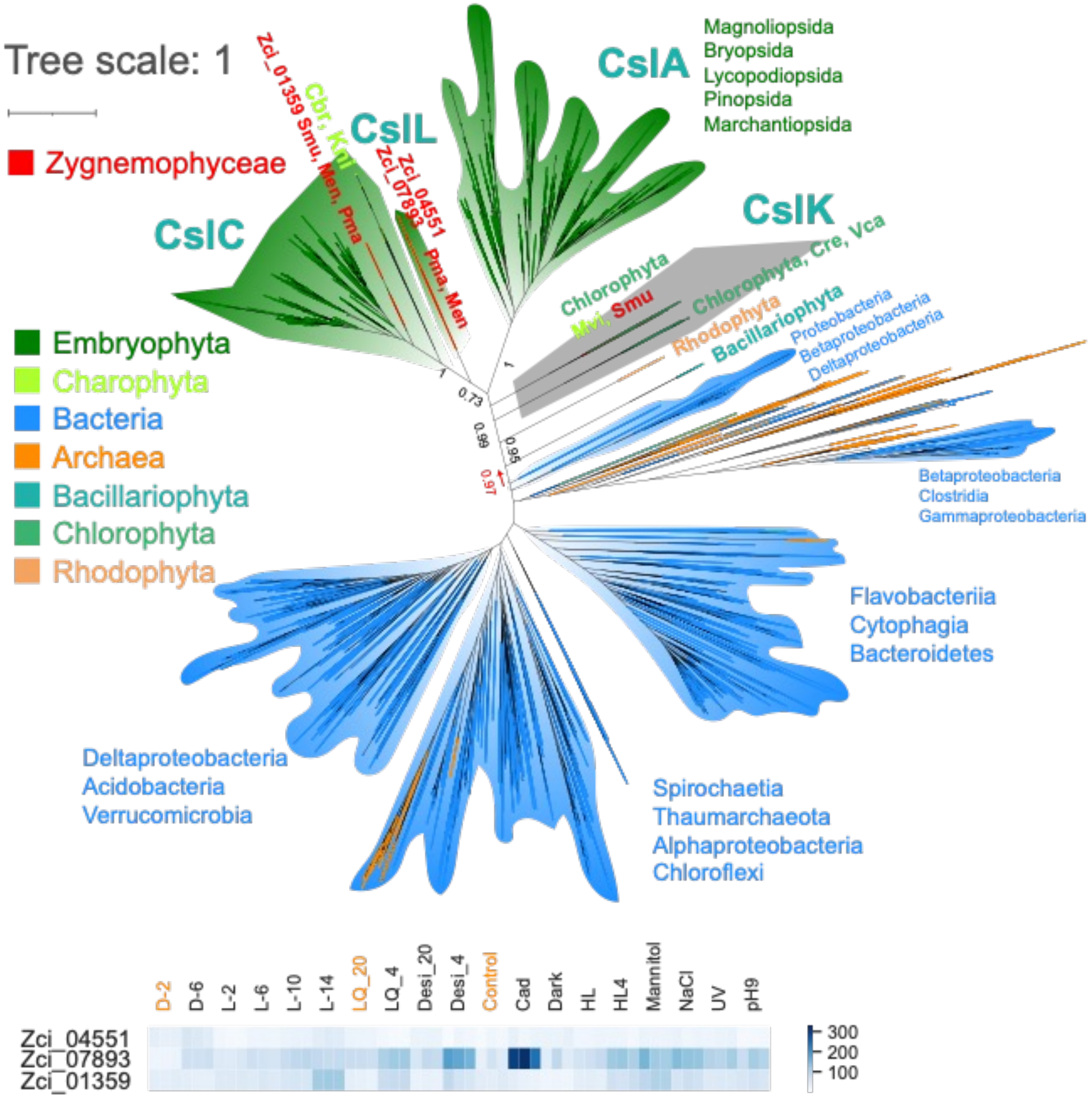

Tree scale: 1
Zygnemophyceae
Embryophyta
Charophyta
Bacteria
Archaea
Bacillariophyta
Chlorophyta
Rhodophyta
CslA
Magnoliopsida
Bryopsida
Lycopodiopsida
Pinopsida
Marchantiopsida
CslC
CslL
CslK
Zci_01359 Smu, Men, Pma
Cbr, Kni
Zci_04551
Zci_07893
Pma, Men
Chlorophyta
Mvi, Smu
Chlorophyta, Cre, Vca
Rhodophyta
Bacillariophyta
Proteobacteria
Betaproteobacteria
Deltaproteobacteria
Betaproteobacteria
Clostridia
Gammaproteobacteria
Flavobacteriia
Cytophagia
Bacteroidetes
Deltaproteobacteria
Acidobacteria
Verrucomicrobia
Spirochaetia
Thaumarchaeota
Alphaproteobacteria
Chloroflexi
0.73
0.99
0.95
0.97
1
D-2
D-6
L-2
L-6
L-10
L-14
LQ_20
LQ_4
Desi_20
Desi_4
Control
Cad
Dark
HL
HL4
Mannitol
NaCl
UV
pH9
Zci_04551
Zci_07893
Zci_01359
300
200
100

Data S-4

Superimposed 3D structures with ligands

**blue**: 4HG6 (bcsA, solved PDB structure, PubMed: 23222542)

**red**: Zci_07893 (predicted ZcCslL1 3D structure from AlphaFold)

**yellow** mesh: GDP-mannose docked into ZcCslL1 structure

**cyan** sticks: UDP-glucose in bcsA structure

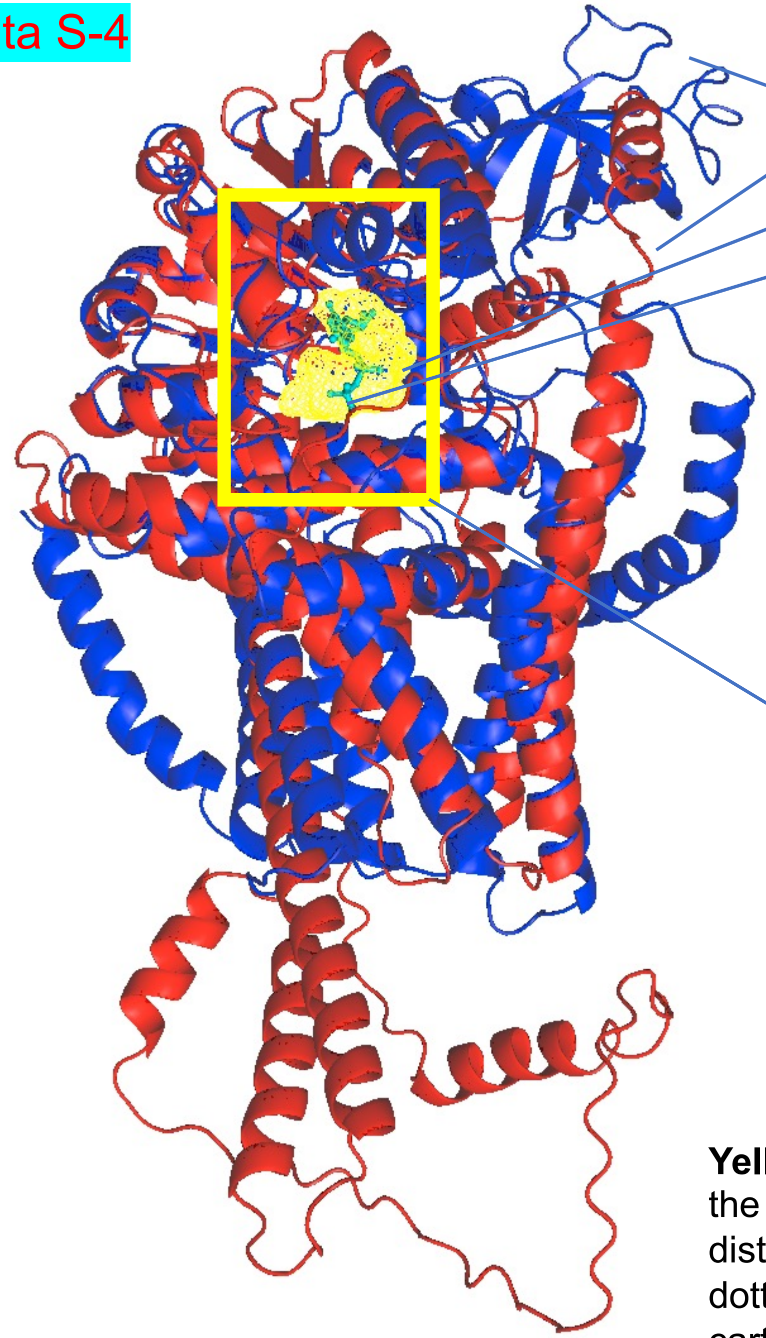

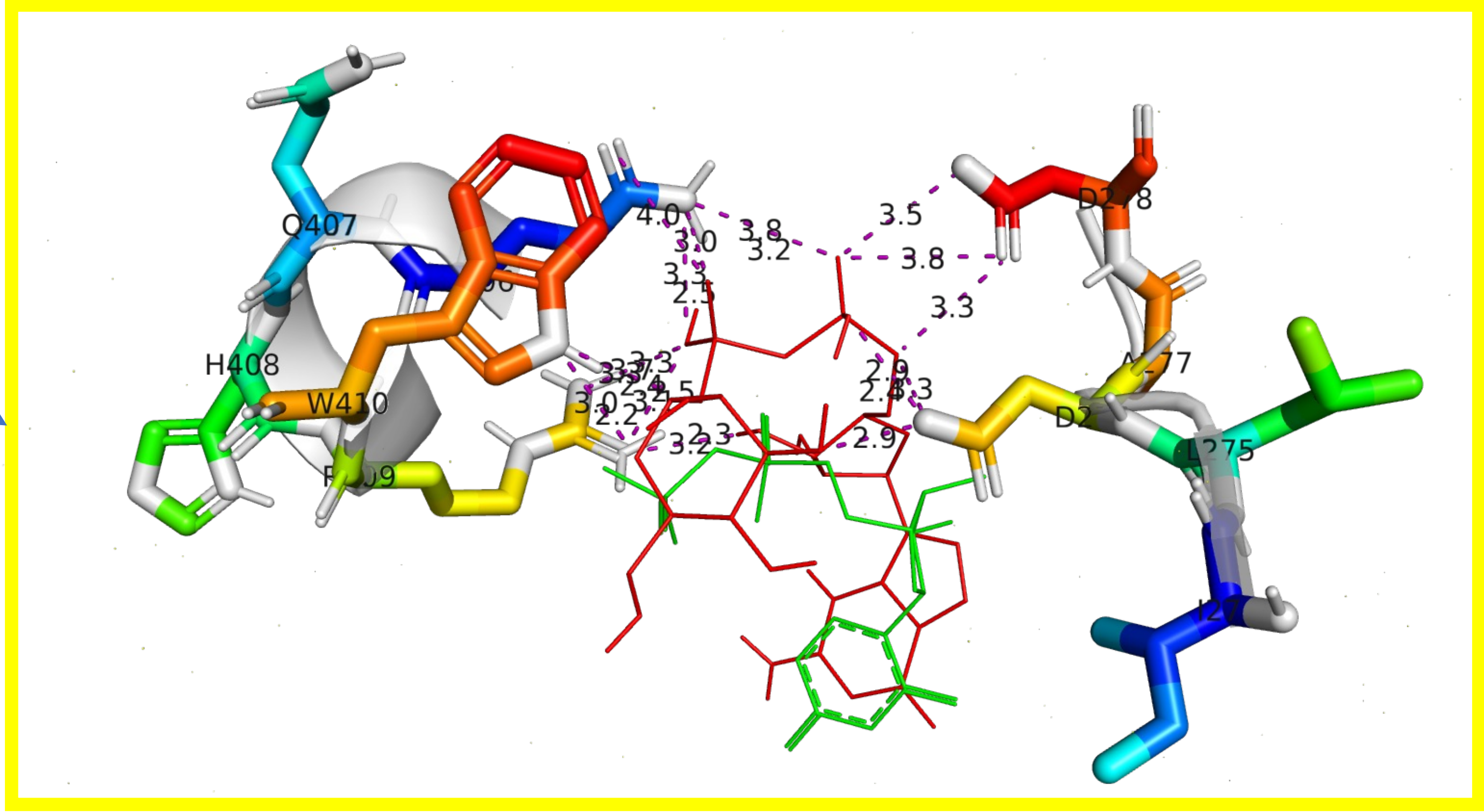


**Yellow** rectangle zoomed-in view of the binding sites of GDP-mannose (red lines) in ZcCslL1: sticks on the left are the QQHRW motif (406-410), on the right are the ILDADD motif (274-278). The RMSD distances between the closest atoms in GDP-mannose and in the two motifs are show indicated with dotted lines. The superimposed binding motifs of UDP-glucose (green lines) in 4HG6 are also shown as cartoon view: on the left are the QRGRW motif (379-383), on the right are the VFDAD motif (244-248).

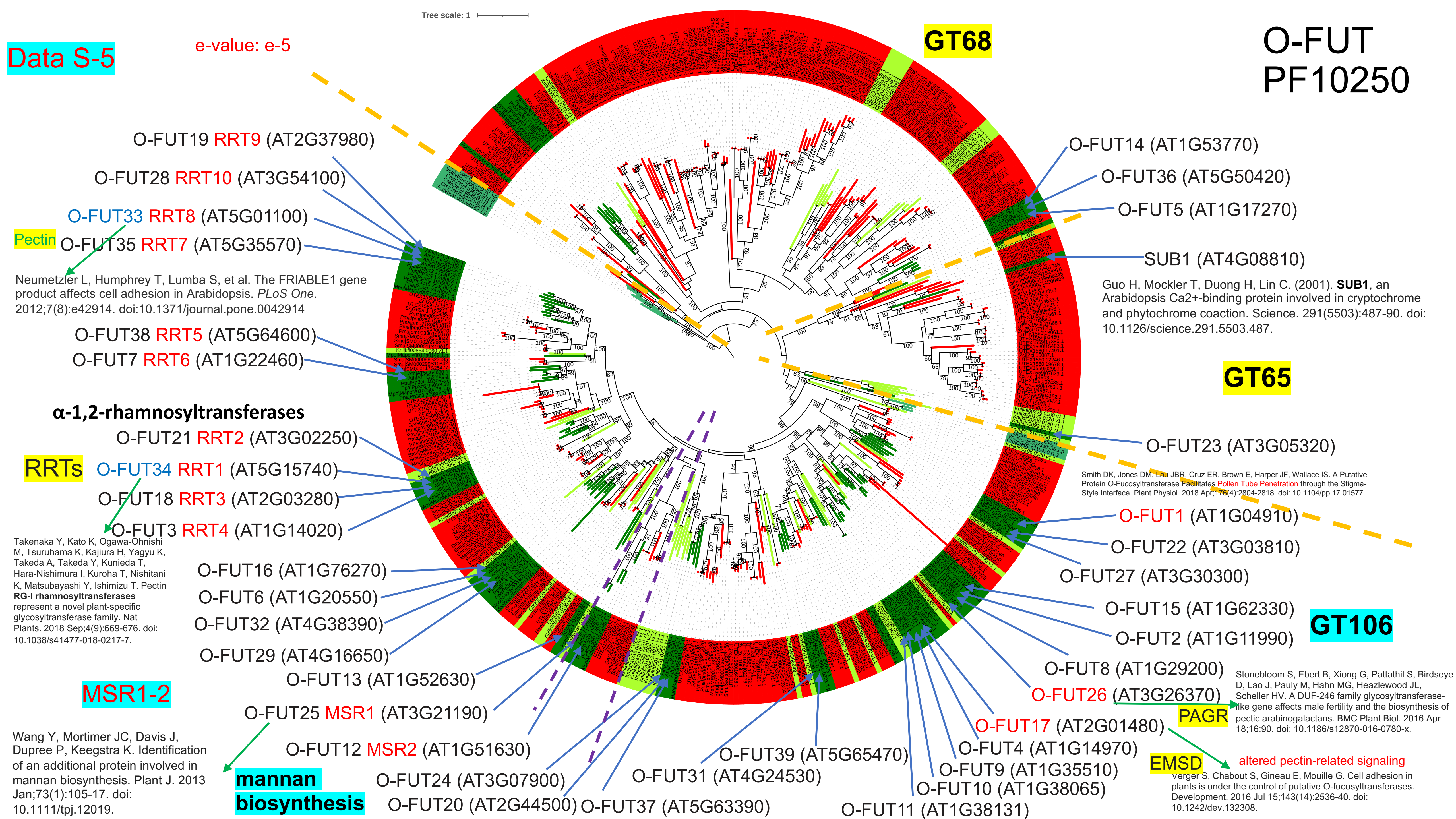

Data S-5
e-value: e-5
Tree scale: 1
GT68
O-FUT
PF10250
O-FUT19 RRT9 (AT2G37980)
O-FUT28 RRT10 (AT3G54100)
O-FUT33 RRT8 (AT5G01100)
Pectin
O-FUT35 RRT7 (AT5G35570)
Neumetzler L, Humphrey T, Lumba S, et al. The FRIABLE1 gene product affects cell adhesion in Arabidopsis. PLoS One. 2012;7(8):e42914. doi:10.1371/journal.pone.0042914
O-FUT38 RRT5 (AT5G64600)
O-FUT7 RRT6 (AT1G22460)
α-1,2-rhamnosyltransferases
O-FUT21 RRT2 (AT3G02250)
RRTs
O-FUT34 RRT1 (AT5G15740)
O-FUT18 RRT3 (AT2G03280)
O-FUT3 RRT4 (AT1G14020)
Takenaka Y, Kato K, Ogawa-Ohnishi M, Tsuruhama K, Kajiura H, Yagyu K, Takeda A, Takeda Y, Kunieda T, Hara-Nishimura I, Kuroha T, Nishitani K, Matsubayashi Y, Ishimizu T. Pectin RG-I rhamnosyltransferases represent a novel plant-specific glycosyltransferase family. Nat Plants. 2018 Sep;4(9):669-676. doi: 10.1038/s41477-018-0217-7.
O-FUT16 (AT1G76270)
O-FUT6 (AT1G20550)
O-FUT32 (AT4G38390)
O-FUT29 (AT4G16650)
O-FUT13 (AT1G52630)
MSR1-2
O-FUT25 MSR1 (AT3G21190)
O-FUT12 MSR2 (AT1G51630)
Wang Y, Mortimer JC, Davis J, Dupree P, Keegstra K. Identification of an additional protein involved in mannan biosynthesis. Plant J. 2013 Jan;73(1):105-17. doi: 10.1111/tpj.12019.
mannan
biosynthesis
O-FUT24 (AT3G07900)
O-FUT20 (AT2G44500)
O-FUT37 (AT5G63390)
O-FUT31 (AT4G24530)
O-FUT39 (AT5G65470)
O-FUT11 (AT1G38131)
O-FUT10 (AT1G38065)
O-FUT9 (AT1G35510)
O-FUT4 (AT1G14970)
O-FUT17 (AT2G01480)
EMSD
altered pectin-related signaling
Verger S, Chabout S, Gineau E, Mouille G. Cell adhesion in plants is under the control of putative O-fucosyltransferases. Development. 2016 Jul 15;143(14):2536-40. doi: 10.1242/dev.132308.
O-FUT26 (AT3G26370)
PAGR
Stonebloom S, Ebert B, Xiong G, Pattathil S, Birdseye D, Lao J, Pauly M, Hahn MG, Heazlewood JL, Scheller HV. A DUF-246 family glycosyltransferase-like gene affects male fertility and the biosynthesis of pectic arabinogalactans. BMC Plant Biol. 2016 Apr 18;16:90. doi: 10.1186/s12870-016-0780-x.
O-FUT8 (AT1G29200)
GT106
O-FUT2 (AT1G11990)
O-FUT15 (AT1G62330)
O-FUT27 (AT3G30300)
O-FUT22 (AT3G03810)
O-FUT1 (AT1G04910)
Smith DK, Jones DM, Lau JBR, Cruz ER, Brown E, Harper JF, Wallace IS. A Putative Protein O-Fucosyltransferase Facilitates Pollen Tube Penetration through the Stigma-Style Interface. Plant Physiol. 2018 Apr;176(4):2804-2818. doi: 10.1104/pp.17.01577.
O-FUT23 (AT3G05320)
GT65
SUB1 (AT4G08810)
Guo H, Mockler T, Duong H, Lin C. (2001). SUB1, an Arabidopsis Ca2+-binding protein involved in cryptochrome and phytochrome coaction. Science. 291(5503):487-90. doi: 10.1126/science.291.5503.487.
O-FUT5 (AT1G17270)
O-FUT36 (AT5G50420)
O-FUT14 (AT1G53770)

Data S-6

# GT34

#### Xyloglucan

XXT5 (AT1G74380)

XXT4 (AT1G18690)

XXT3 (AT5G07720)

XXT2 (AT4G02500)

XXT1 (AT3G62720)

Faik A, Price NJ, Raikhel NV, Keegstra K. An Arabidopsis gene encoding an alpha-xylosyltransferase involved in xyloglucan biosynthesis. Proc Natl Acad Sci U S A. 2002 May 28;99(11):7797-802. doi: 10.1073/pnas.102644799.

Cavalier DM, Keegstra K. Two xyloglucan xylosyltransferases catalyze the addition of multiple xylosyl residues to cellohexaose. J Biol Chem. 2006 Nov 10;281(45):34197-207. doi: 10.1074/jbc.M606379200.

Vuttipongchaikij S, Brocklehurst D, Steele-King C, Ashford DA, Gomez LD, McQueen-Mason SJ. Arabidopsis GT34 family contains five xyloglucan α-1,6-xylosyltransferases. New Phytol. 2012 Aug;195(3):585-595. doi: 10.1111/j.1469-8137.2012.04196.x.

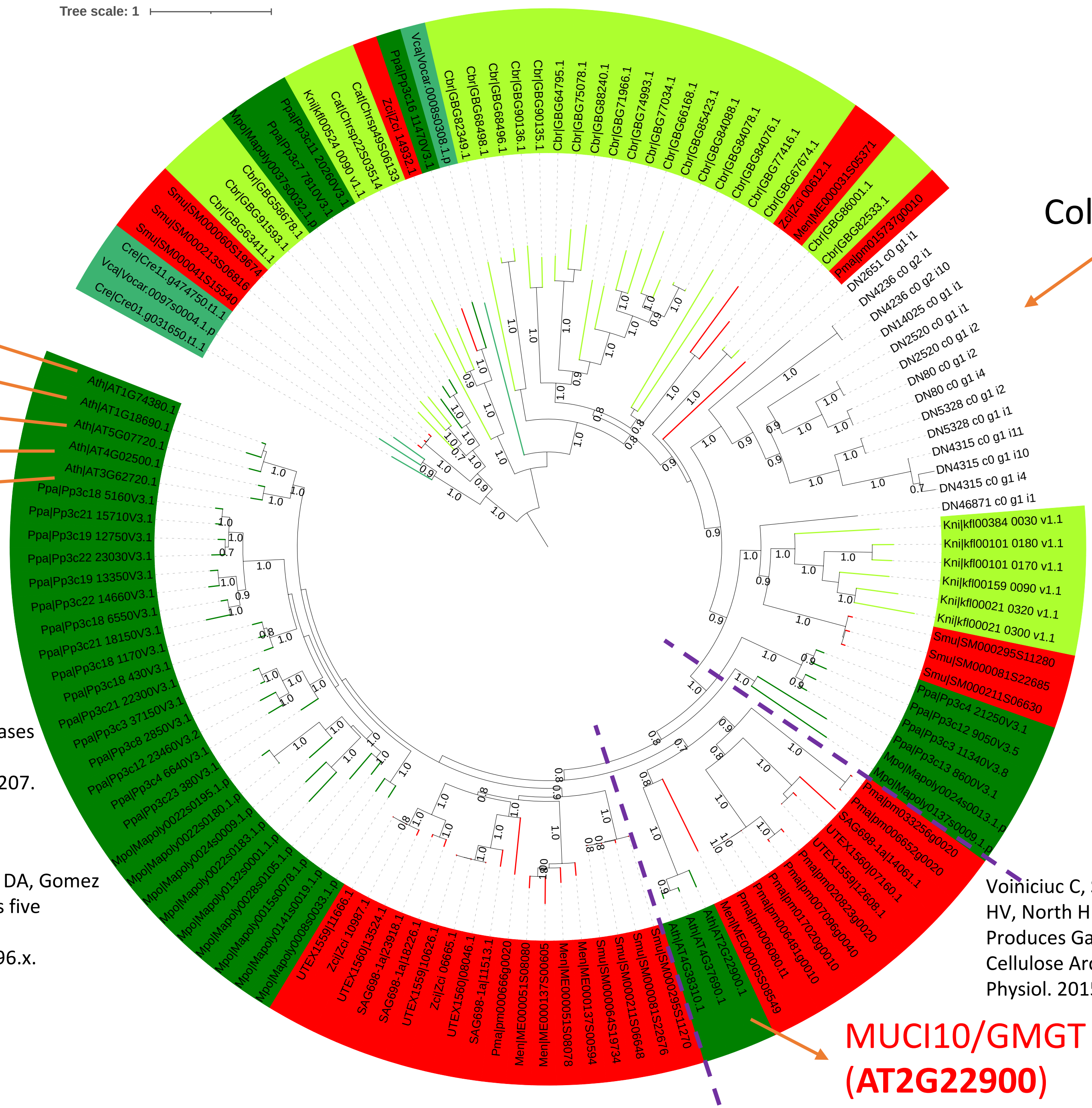


Coleochaete scutate

#### Mannan

galactomannan α-1,6-galactosyltransferase activity

Voiniciuc C, Schmidt MH, Berger A, Yang B, Ebert B, Scheller HV, North HM, Usadel B, Günl M. MUCILAGE-RELATED10 Produces Galactoglucomannan That Maintains Pectin and Cellulose Architecture in Arabidopsis Seed Mucilage. Plant Physiol. 2015 Sep;169(1):403-20. doi: 10.1104/pp.15.00851.

MUCI10/GMGT (**AT2G22900**)

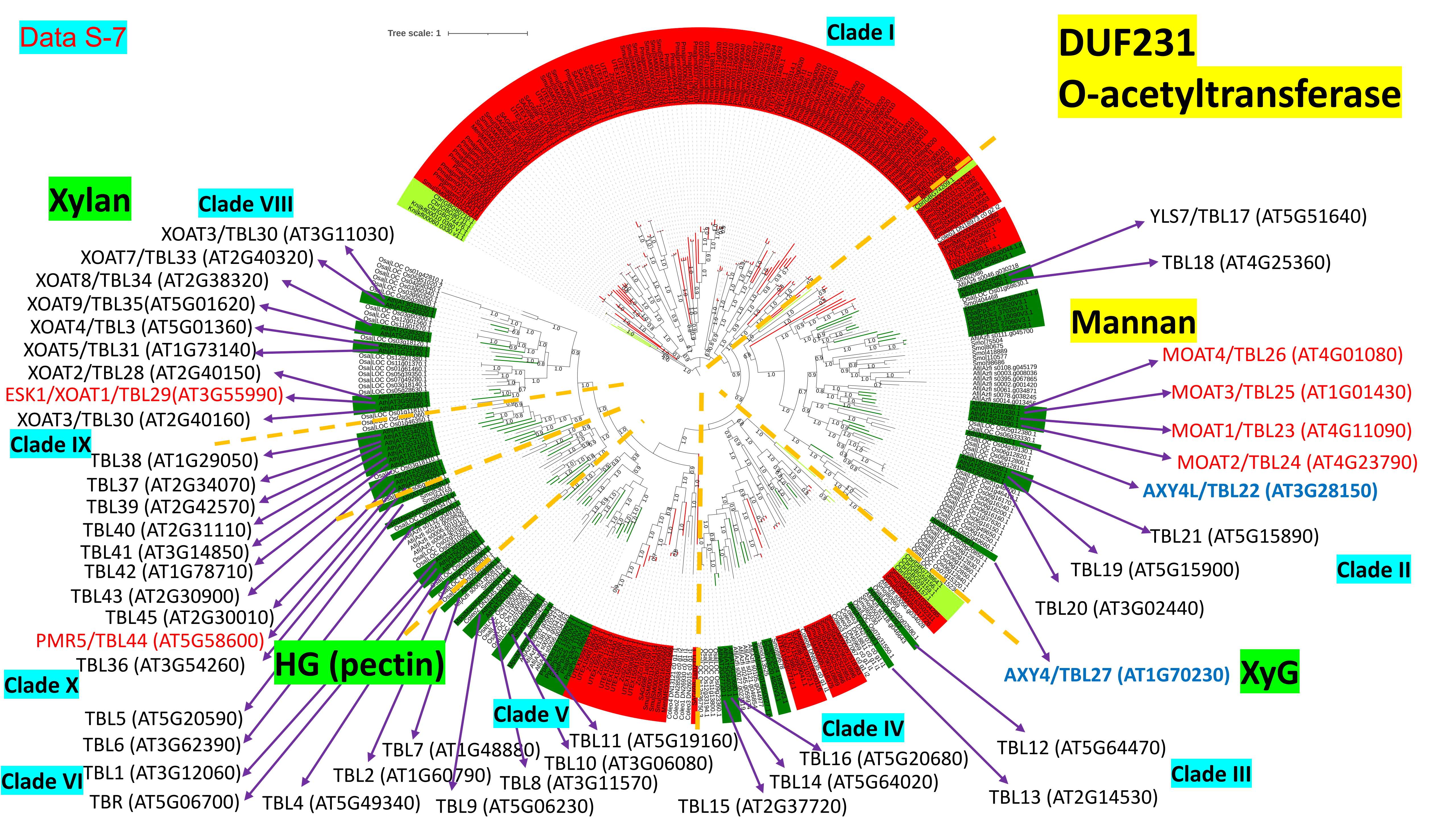
Data S-7
Tree scale: 1
Clade I
DUF231
O-acetyltransferase
Xylan
Clade VIII
XOAT3/TBL30 (AT3G11030)
XOAT7/TBL33 (AT2G40320)
XOAT8/TBL34 (AT2G38320)
XOAT9/TBL35(AT5G01620)
XOAT4/TBL3 (AT5G01360)
XOAT5/TBL31 (AT1G73140)
XOAT2/TBL28 (AT2G40150)
ESK1/XOAT1/TBL29(AT3G55990)
XOAT3/TBL30 (AT2G40160)
Clade IX
TBL38 (AT1G29050)
TBL37 (AT2G34070)
TBL39 (AT2G42570)
TBL40 (AT2G31110)
TBL41 (AT3G14850)
TBL42 (AT1G78710)
TBL43 (AT2G30900)
TBL45 (AT2G30010)
PMR5/TBL44 (AT5G58600)
TBL36 (AT3G54260)
Clade X
HG (pectin)
TBL5 (AT5G20590)
TBL6 (AT3G62390)
TBL1 (AT3G12060)
Clade VI
TBR (AT5G06700)
TBL4 (AT5G49340)
TBL2 (AT1G60790)
TBL7 (AT1G48880)
TBL9 (AT5G06230)
TBL8 (AT3G11570)
Clade V
TBL10 (AT3G06080)
TBL11 (AT5G19160)
TBL15 (AT2G37720)
TBL14 (AT5G64020)
TBL16 (AT5G20680)
Clade IV
TBL13 (AT2G14530)
TBL12 (AT5G64470)
Clade III
AXY4/TBL27 (AT1G70230)
XyG
TBL20 (AT3G02440)
TBL19 (AT5G15900)
Clade II
TBL21 (AT5G15890)
AXY4L/TBL22 (AT3G28150)
MOAT2/TBL24 (AT4G23790)
MOAT1/TBL23 (AT4G11090)
MOAT3/TBL25 (AT1G01430)
MOAT4/TBL26 (AT4G01080)
Mannan
TBL18 (AT4G25360)
YLS7/TBL17 (AT5G51640)

Data S-8

# GH5_7

**endo-β-1,4-mannanase (EC 3.2.1.78)**

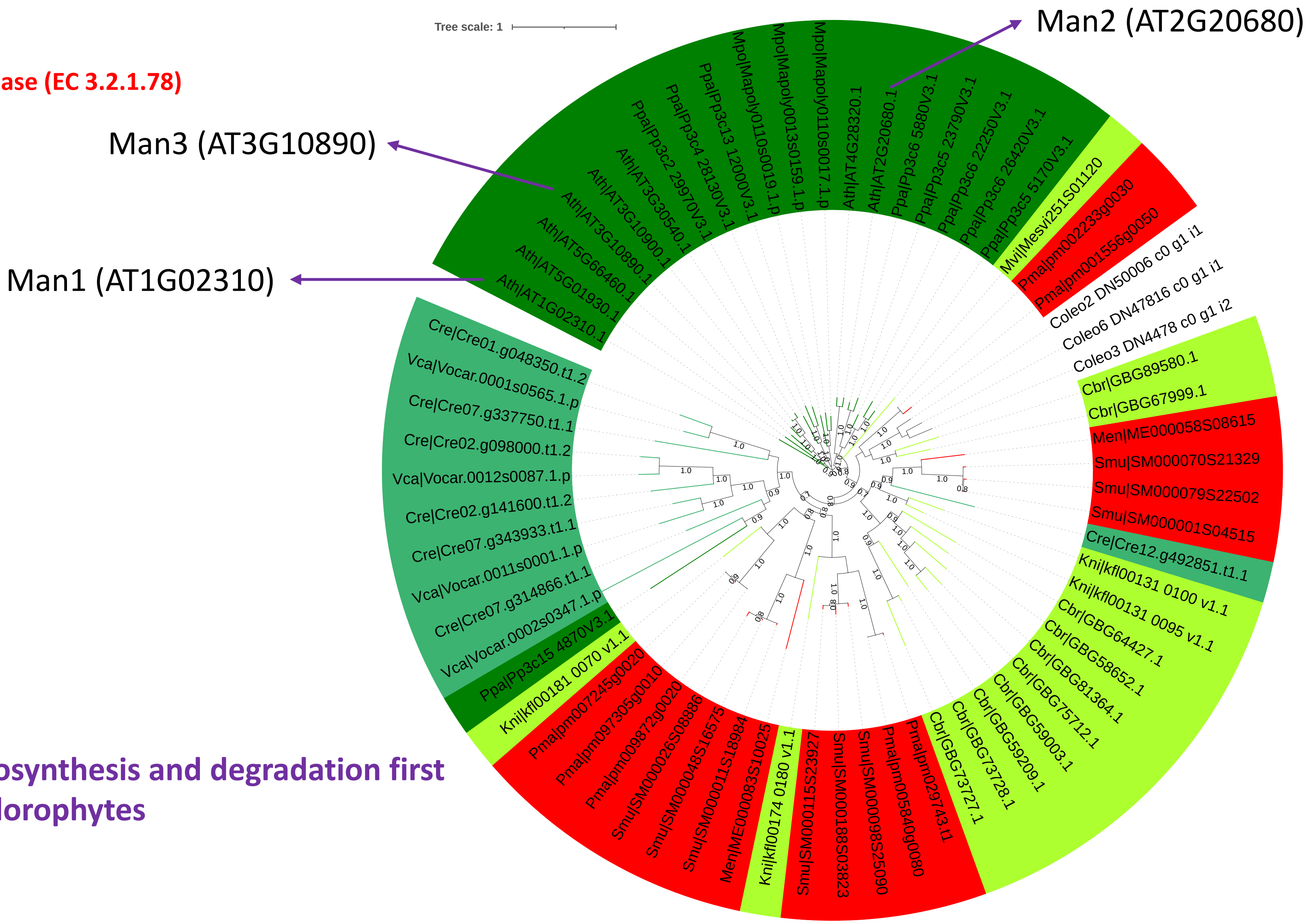


**The mannan biosynthesis and degradation first appeared in Chlorophytes**

Data S-9

# GH5_7 (nr)

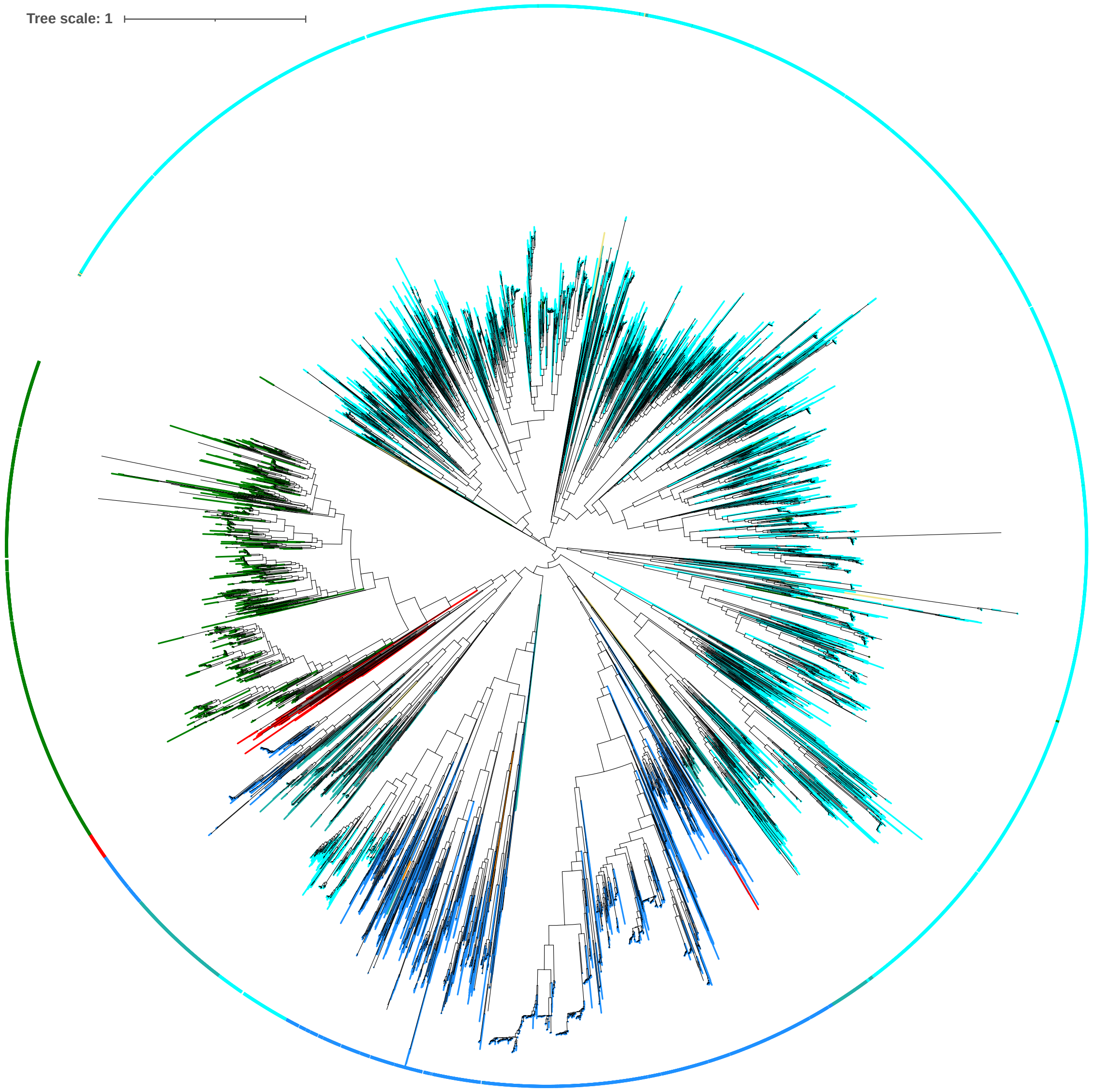


**Possible HGT from bacteria**

https://itol.embl.de/tree/134238164171357031667587420

Data S-10

# GH27 nr

Imaizumi et al. Heterologous expression and characterization of an Arabidopsis β-l-arabinopyranosidase and α-d-galactosidases acting on β-l-arabinopyranosyl residues. J Exp Bot. 2017 Jul 20;68(16):4651-4661. doi: 10.1093/jxb/erx279.

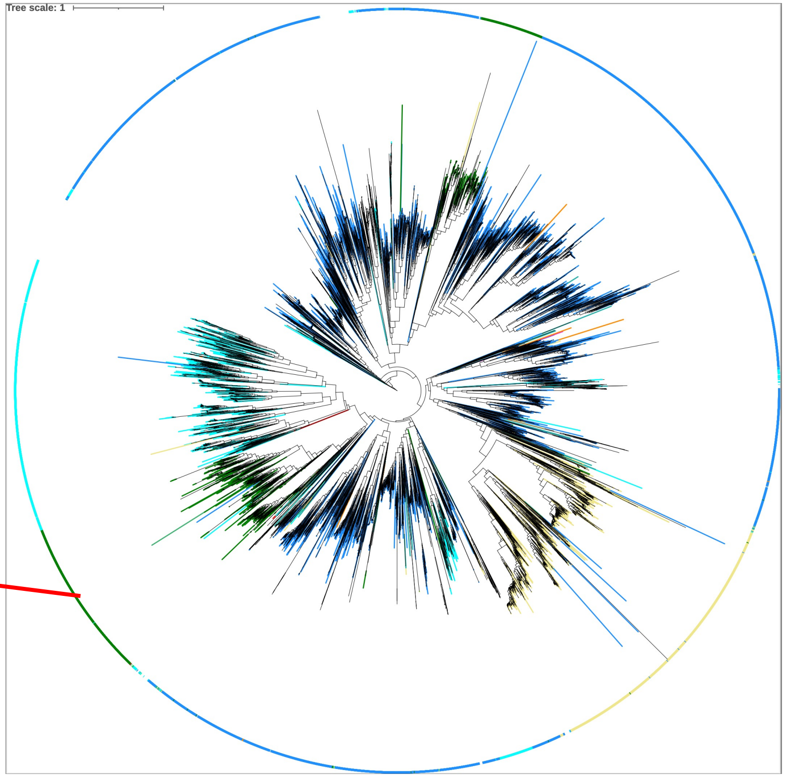


AGAL1 (AT5G08380)
AGAL2 (AT5G08370)
AGAL3 (AT3G56310)

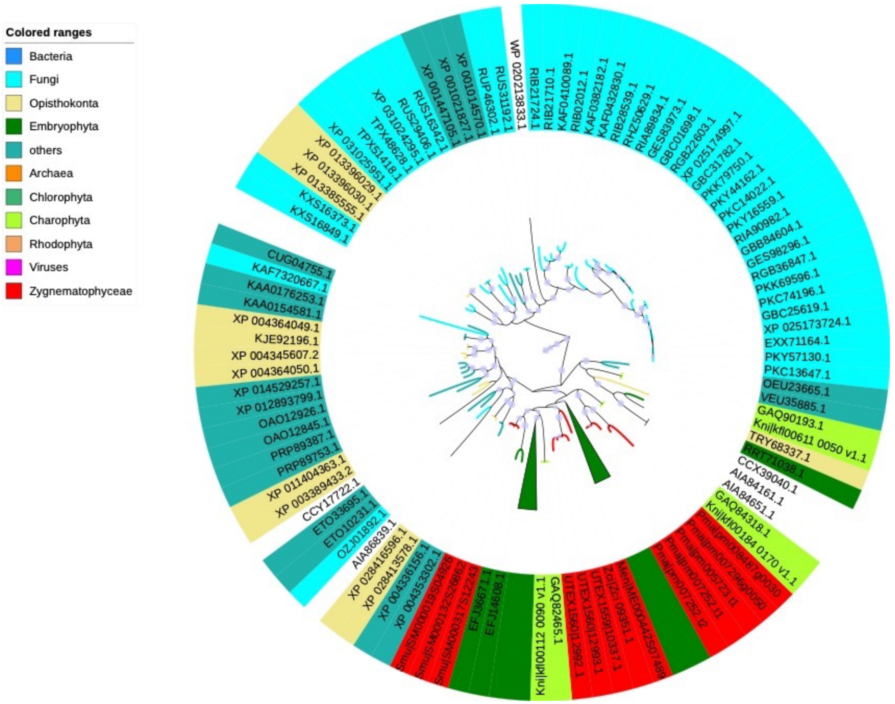


might have been gained via HGT between fungi and *Klebsormidium*

https://itol.embl.de/tree/2081272402225633166520345

Data S-11

# GH27 nr

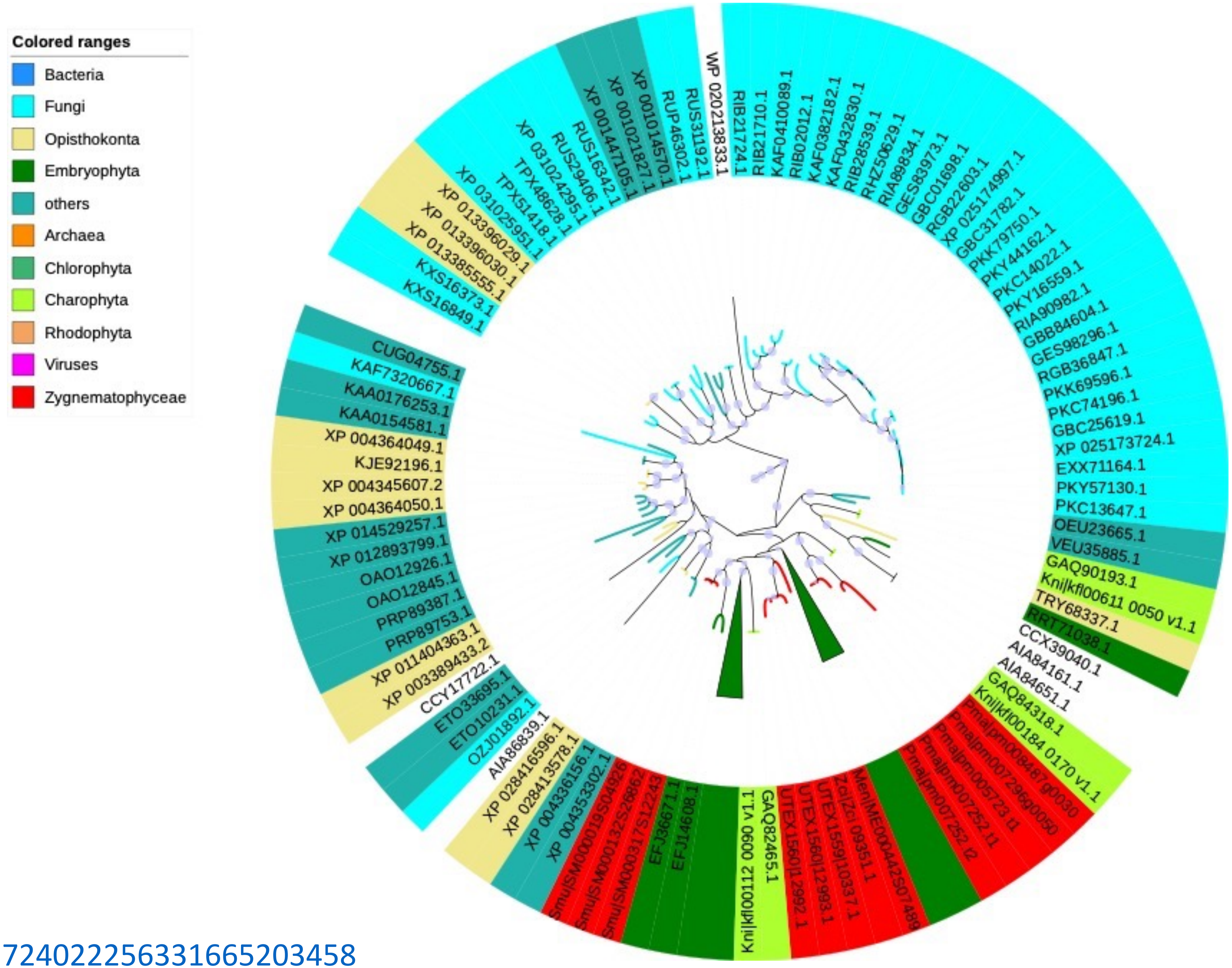


https://itol.embl.de/tree/2081272402225633166520345

Data S-12

https://itol.embl.de/tree/2081272402221437816720288870

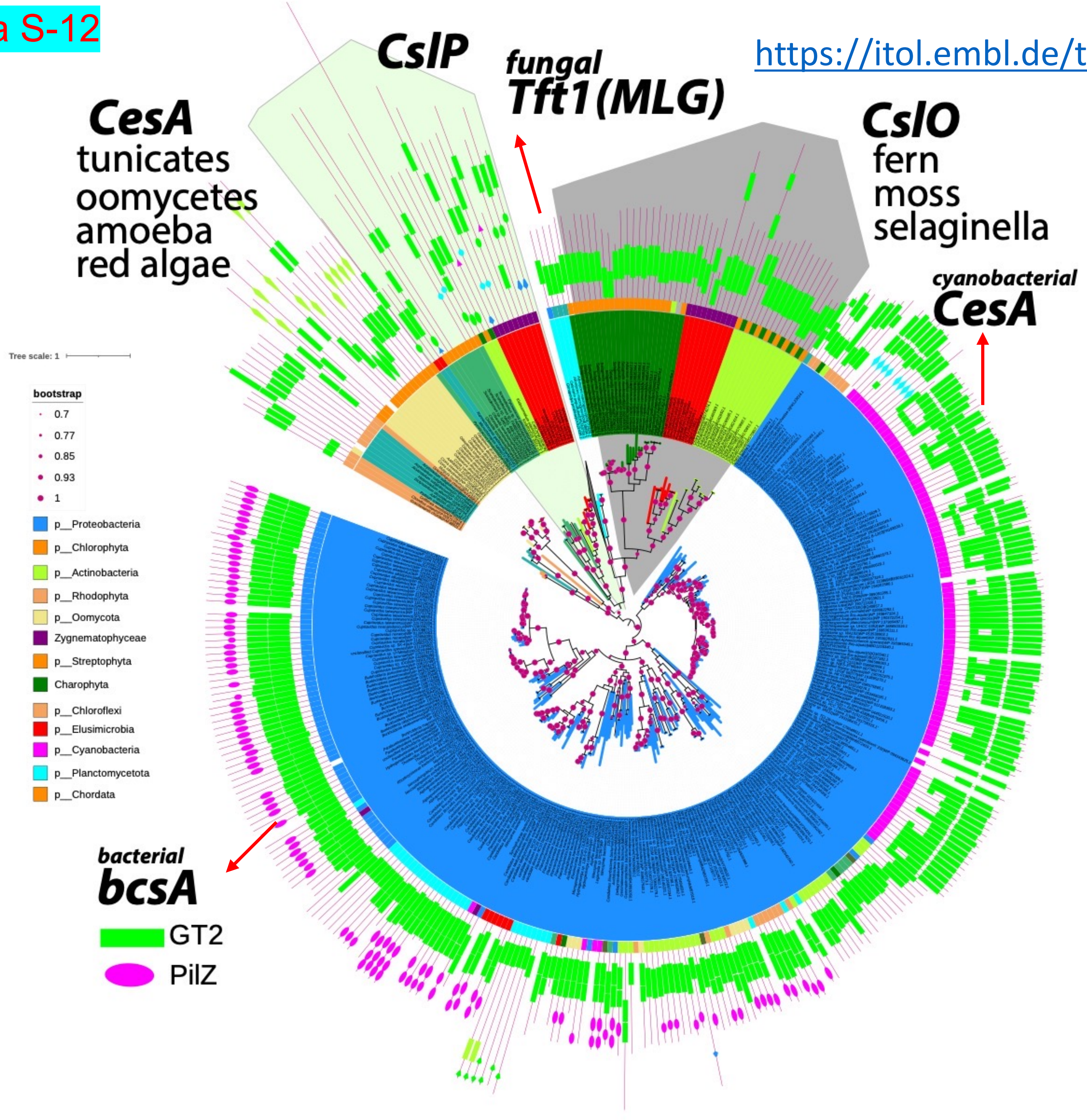

Data S-13

https://itol.embl.de/tree/2081272402224417016711125554

CslN might have been gained via HGT from bacteria (bcsA)

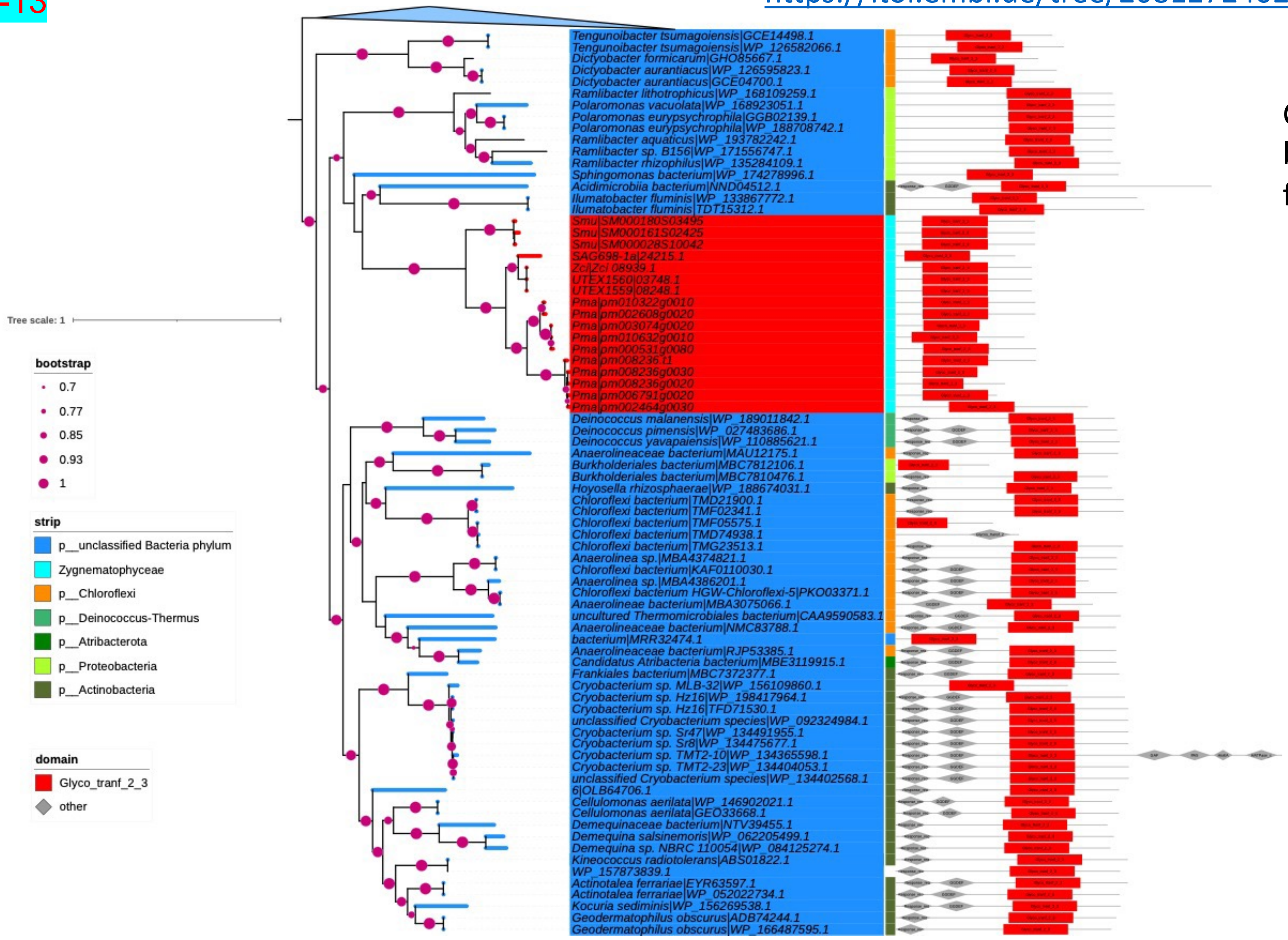

Data S-14

### Xyloglucan

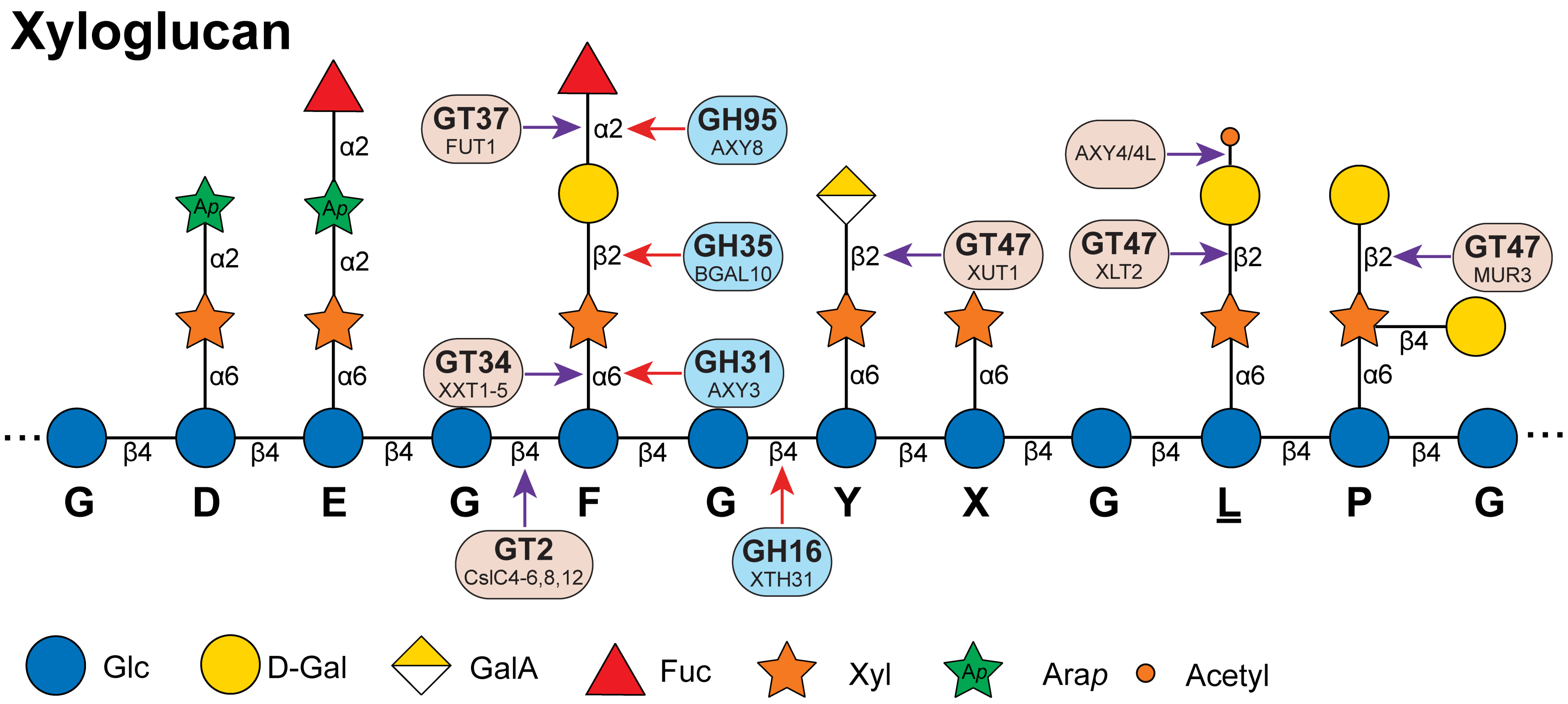

Data S-15

# GT34

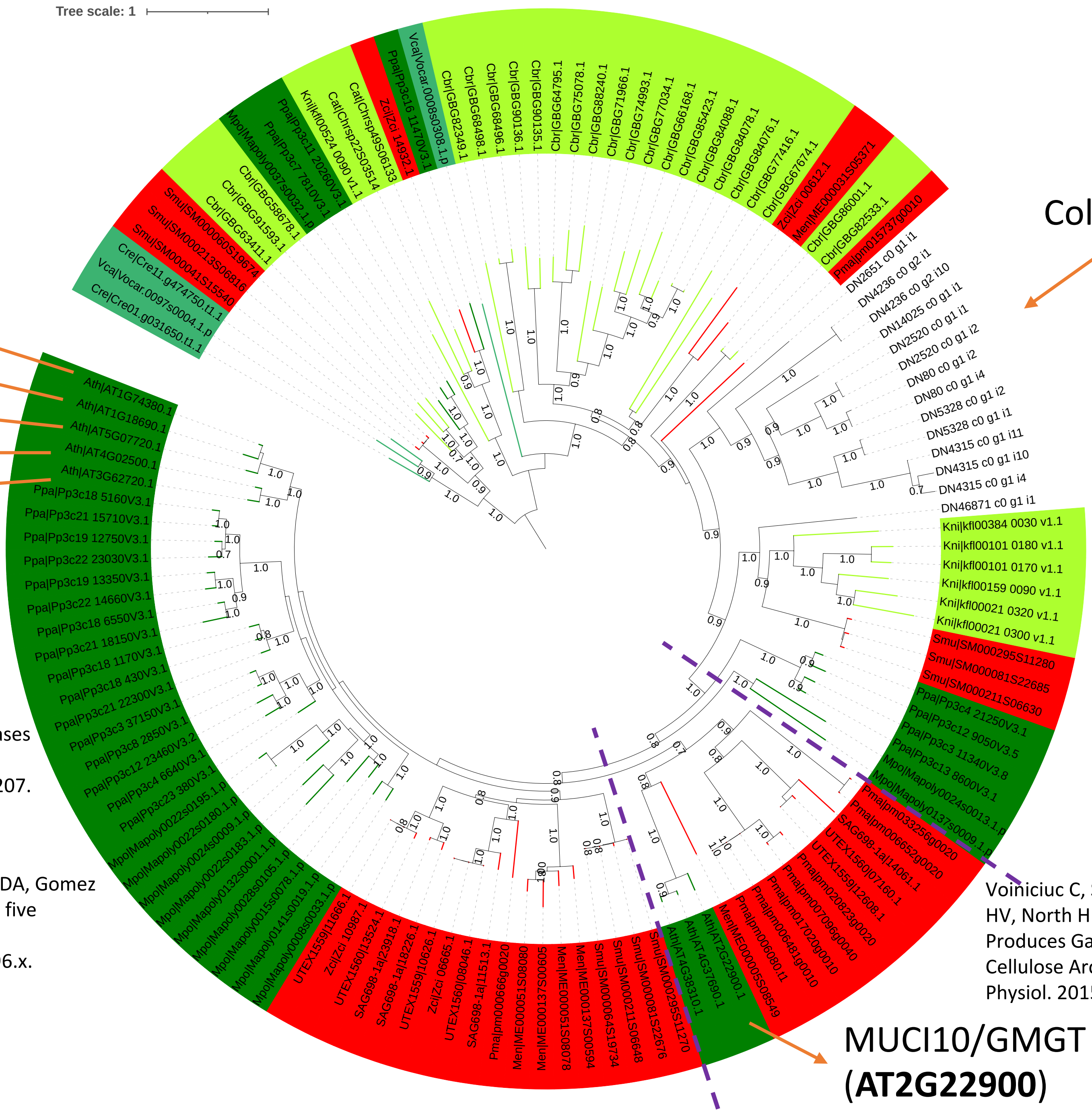


#### Xyloglucan

XXT5 (AT1G74380)

XXT4 (AT1G18690)

XXT3 (AT5G07720)

XXT2 (AT4G02500)

XXT1 (AT3G62720)

Faik A, Price NJ, Raikhel NV, Keegstra K. An Arabidopsis gene encoding an alpha-xylosyltransferase involved in xyloglucan biosynthesis. Proc Natl Acad Sci U S A. 2002 May 28;99(11):7797-802. doi: 10.1073/pnas.102644799.

Cavalier DM, Keegstra K. Two xyloglucan xylosyltransferases catalyze the addition of multiple xylosyl residues to cellohexaose. J Biol Chem. 2006 Nov 10;281(45):34197-207. doi: 10.1074/jbc.M606379200.

Vuttipongchaikij S, Brocklehurst D, Steele-King C, Ashford DA, Gomez LD, McQueen-Mason SJ. Arabidopsis GT34 family contains five xyloglucan α-1,6-xylosyltransferases. New Phytol. 2012 Aug;195(3):585-595. doi: 10.1111/j.1469-8137.2012.04196.x.

Coleochaete scutate

#### Mannan

galactomannan α-1,6-galactosyltransferase activity

Voiniciuc C, Schmidt MH, Berger A, Yang B, Ebert B, Scheller HV, North HM, Usadel B, Günl M. MUCILAGE-RELATED10 Produces Galactoglucomannan That Maintains Pectin and Cellulose Architecture in Arabidopsis Seed Mucilage. Plant Physiol. 2015 Sep;169(1):403-20. doi: 10.1104/pp.15.00851.

MUCI10/GMGT (**AT2G22900**)

Data S-16

# GT47

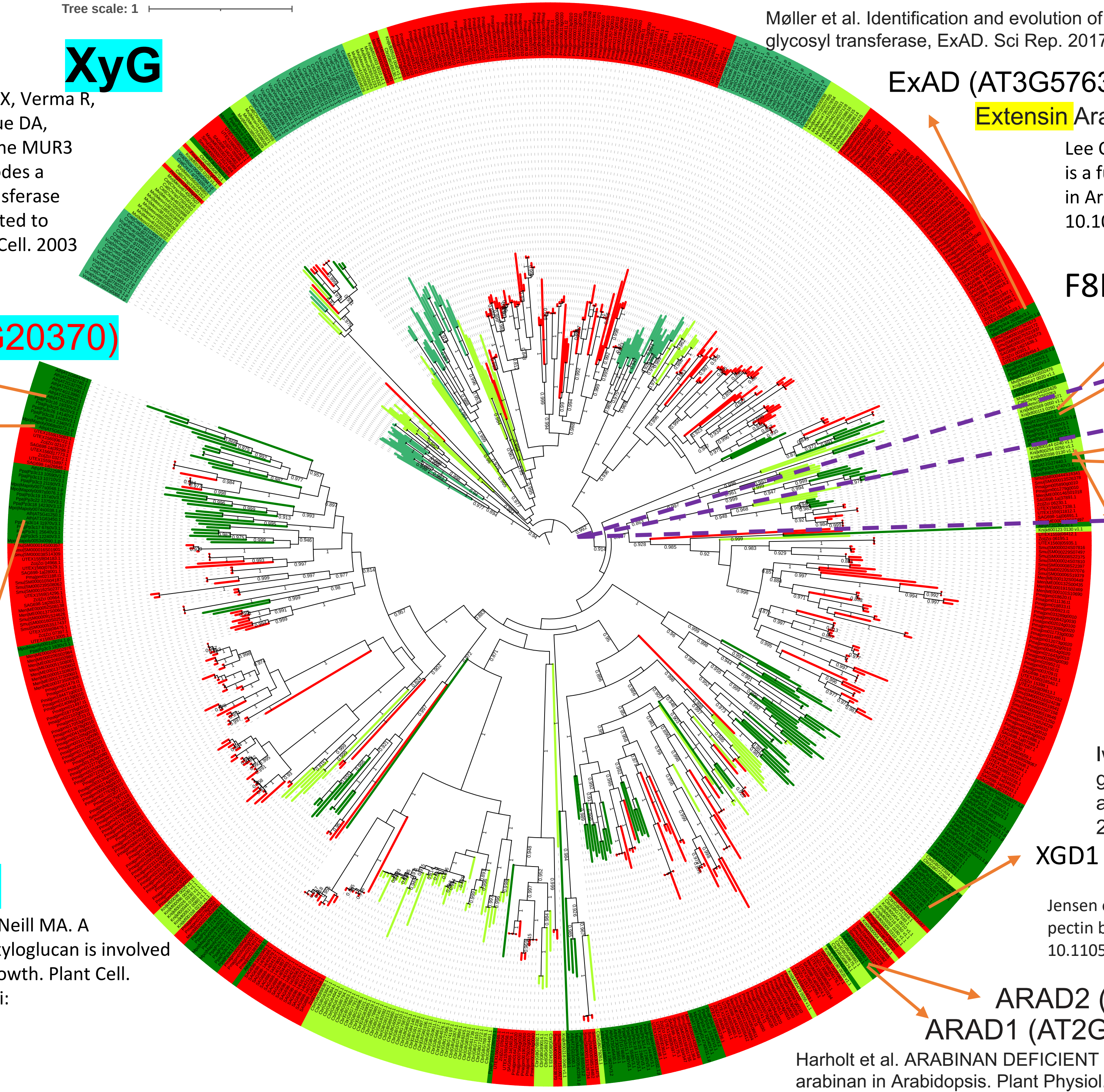


**XyG**

Madson M, Dunand C, Li X, Verma R, Vanzin GF, Caplan J, Shoue DA, Carpita NC, Reiter WD. The MUR3 gene of Arabidopsis encodes a xyloglucan galactosyltransferase that is evolutionarily related to animal exostosins. Plant Cell. 2003 Jul;15(7):1662-70. doi: 10.1105/tpc.009837.

MUR3 (AT2G20370)

XLT2 (AT5G62220)

Jensen JK, Schultink A, Keegstra K, Wilkerson CG, Pauly M. RNA-Seq analysis of developing nasturtium seeds (Tropaeolum majus): identification and characterization of an additional galactosyltransferase involved in xyloglucan biosynthesis. Mol Plant. 2012 Sep;5(5):984-92. doi: 10.1093/mp/sss032.

XUT1 (AT1G63450)

Peña MJ, Kong Y, York WS, O'Neill MA. A galacturonic acid-containing xyloglucan is involved in Arabidopsis root hair tip growth. Plant Cell. 2012 Nov;24(11):4511-24. doi: 10.1105/tpc.112.103390.

Møller et al. Identification and evolution of a plant cell wall specific glycoprotein glycosyl transferase, ExAD. Sci Rep. 2017 Mar 30;7:45341. doi: 10.1038/srep45341

ExAD (AT3G57630)

Extensin Arabinose Deficient transferase

Lee C, Teng Q, Huang W, Zhong R, Ye ZH. The F8H glycosyltransferase is a functional paralog of FRA8 involved in glucuronoxylan biosynthesis in Arabidopsis. Plant Cell Physiol. 2009 Apr;50(4):812-27. doi: 10.1093/pcp/pcp025. Epub 2009 Feb 18. PMID: 19224953.

F8H (AT5G22940)

IRX7/FRA8 (AT2G28110)

Brown, D.M., Goubet, F., Wong, V.W., Goodacre, R., Stephens, E., Dupree, P., and Turner, S.R. (2007). Comparison of five xylan synthesis mutants reveals new insight into the mechanisms of xylan synthesis. Plant J. 52: 1154–1168.

XYS1/IRX10-L (AT5G61840)

IRX10 (AT1G27440)

**Xylan**

Jensen JK, Busse-Wicher M, Poulsen CP, Fangel JU, Smith PJ, Yang JY, Peña MJ, Dinesen MH, Martens HJ, Melkonian M et al. 2018. Identification of an algal xylan synthase indicates that there is functional orthology between algal and plant cell wall biosynthesis. New Phytologist 218: 1049–1060.

Urbanowicz BR, Peña MJ, Moniz HA, Moremen KW, York WS. 2014. Two Arabidopsis proteins synthesize acetylated xylan in vitro. Plant Journal 80: 197–206.

NpGUT1 is a pectin b-glucuronyltransferase that transfers glucuronic acid to RG-II

Iwai H, Masaoka N, Ishii T, Satoh S. A pectin glucuronyltransferase gene is essential for intercellular attachment in the plant meristem. Proc Natl Acad Sci U S A. 2002 Dec 10;99(25):16319-24. doi: 10.1073/pnas.252530499.

XGD1 (AT5G33290)

**XGA**

Jensen et al. Identification of a xylogalacturonan xylosyltransferase involved in pectin biosynthesis in Arabidopsis. Plant Cell. 2008 May;20(5):1289-302. doi: 10.1105/tpc.107.050906

ARAD2 (AT5G44930)

ARAD1 (AT2G35100)

**RG-I**

Harholt et al. ARABINAN DEFICIENT 1 is a putative arabinosyltransferase involved in biosynthesis of pectic arabinan in Arabidopsis. Plant Physiol. 2006 Jan;140(1):49-58. doi: 10.1104/pp.105.072744.

Data S-17

# GT47

https://itol.embl.de/tree/20812788684772516631797540

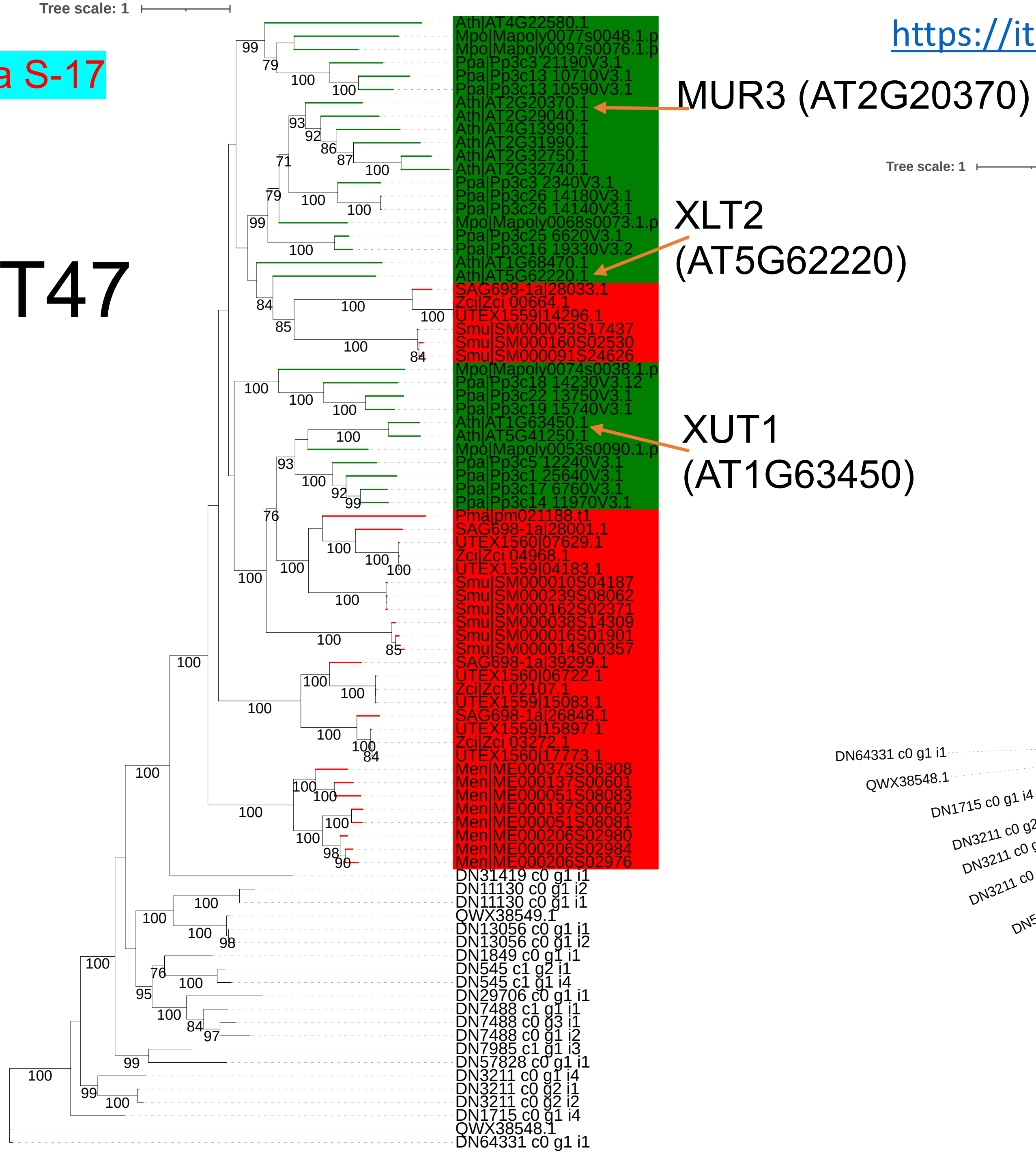


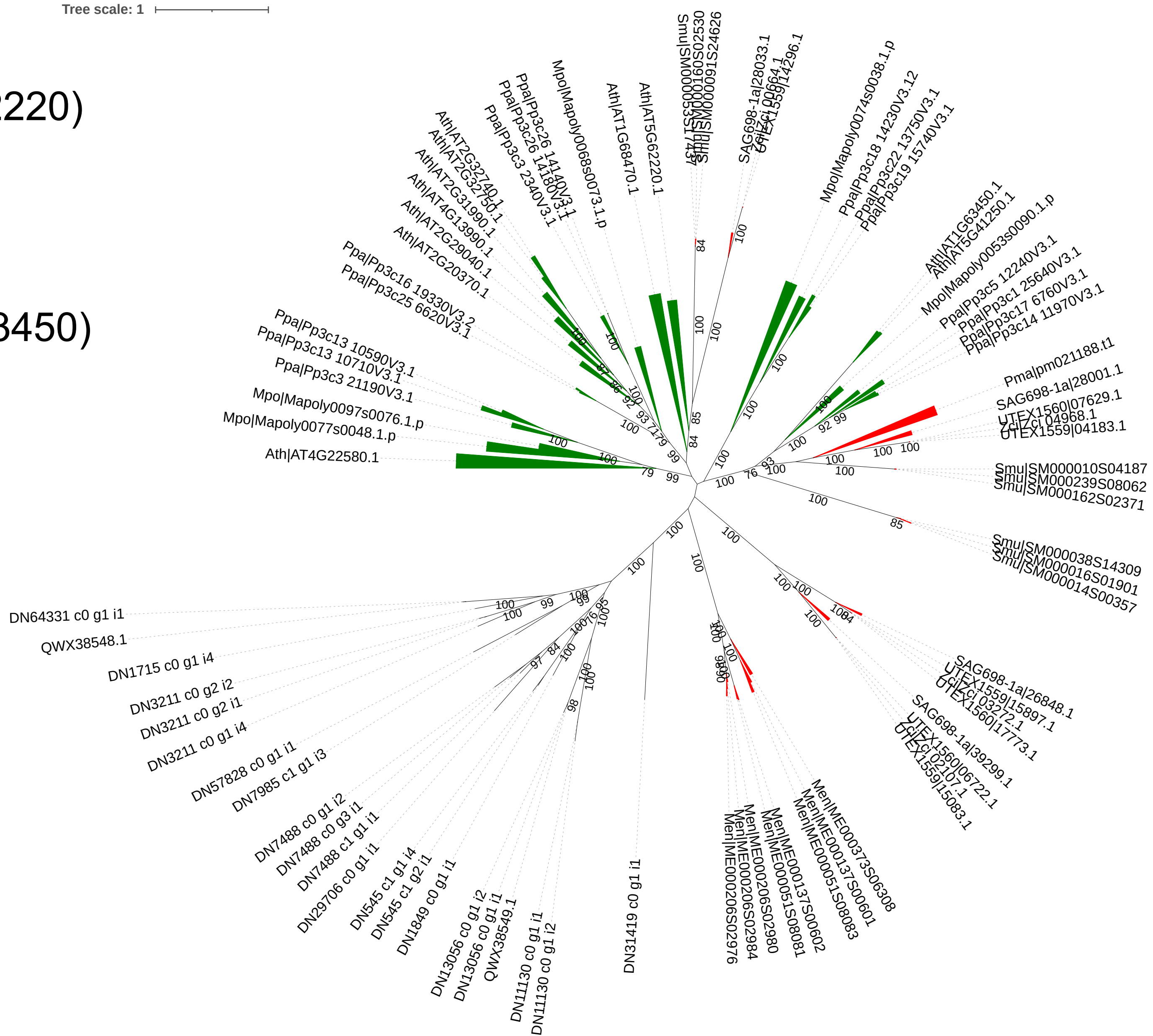

Data S-18

# GT37

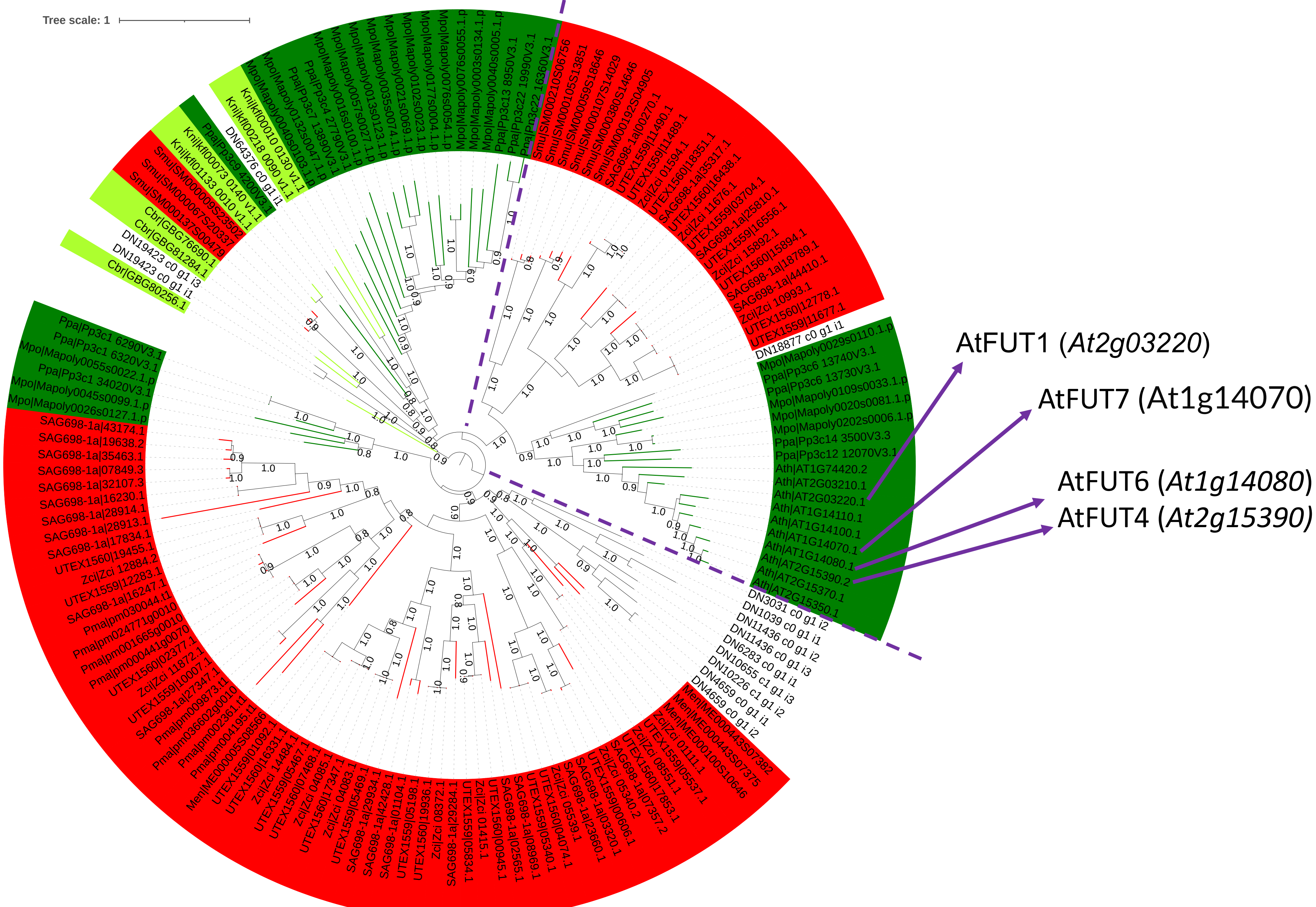

Data S-19

https://itol.embl.de/tree/134238164171257581669144581

# GT37 nr

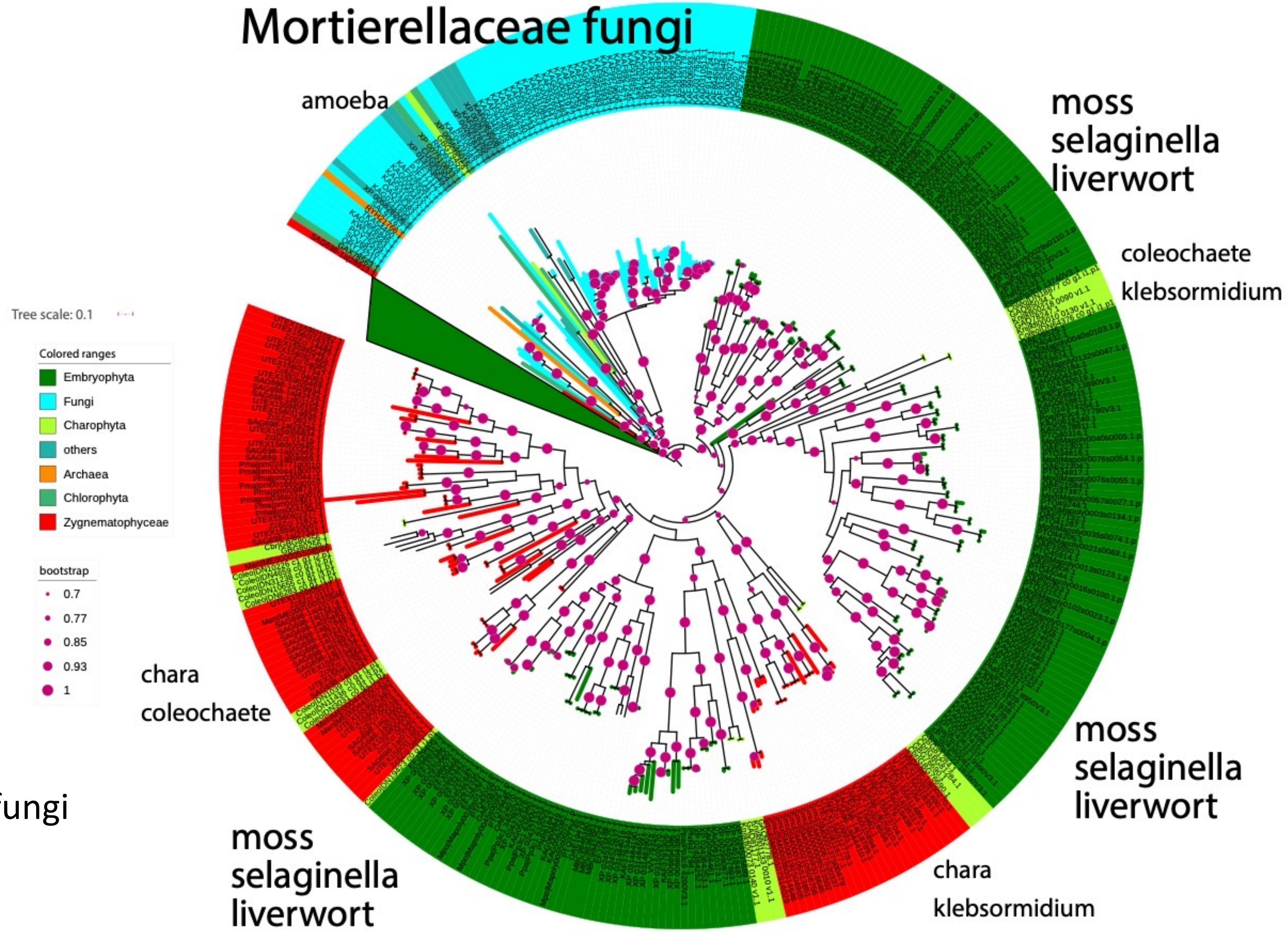


might have been
gained via HGT from fungi

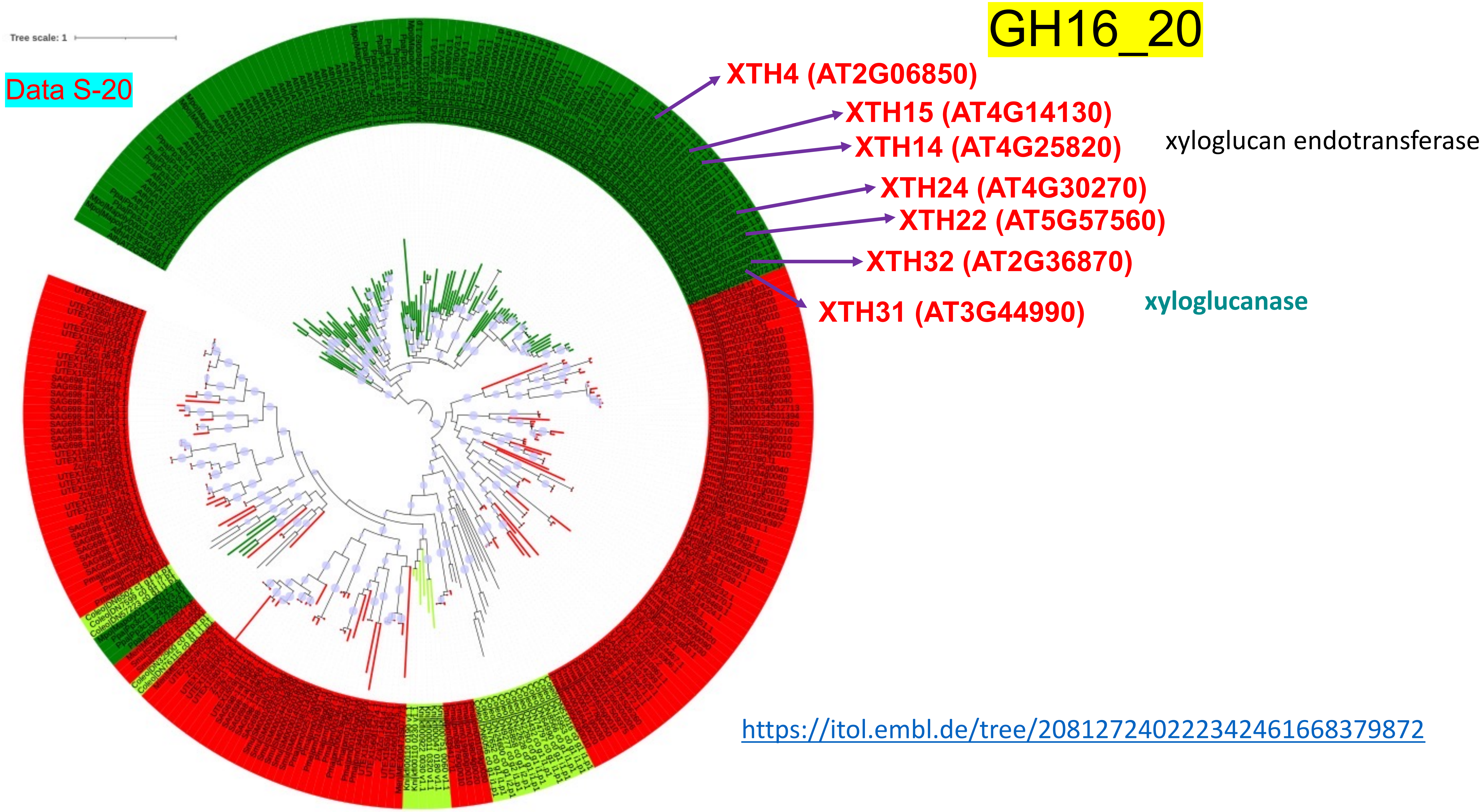


https://itol.embl.de/tree/20812724022342461668379872

https://itol.embl.de/tree/134238164171126381664207231

# GH16_20

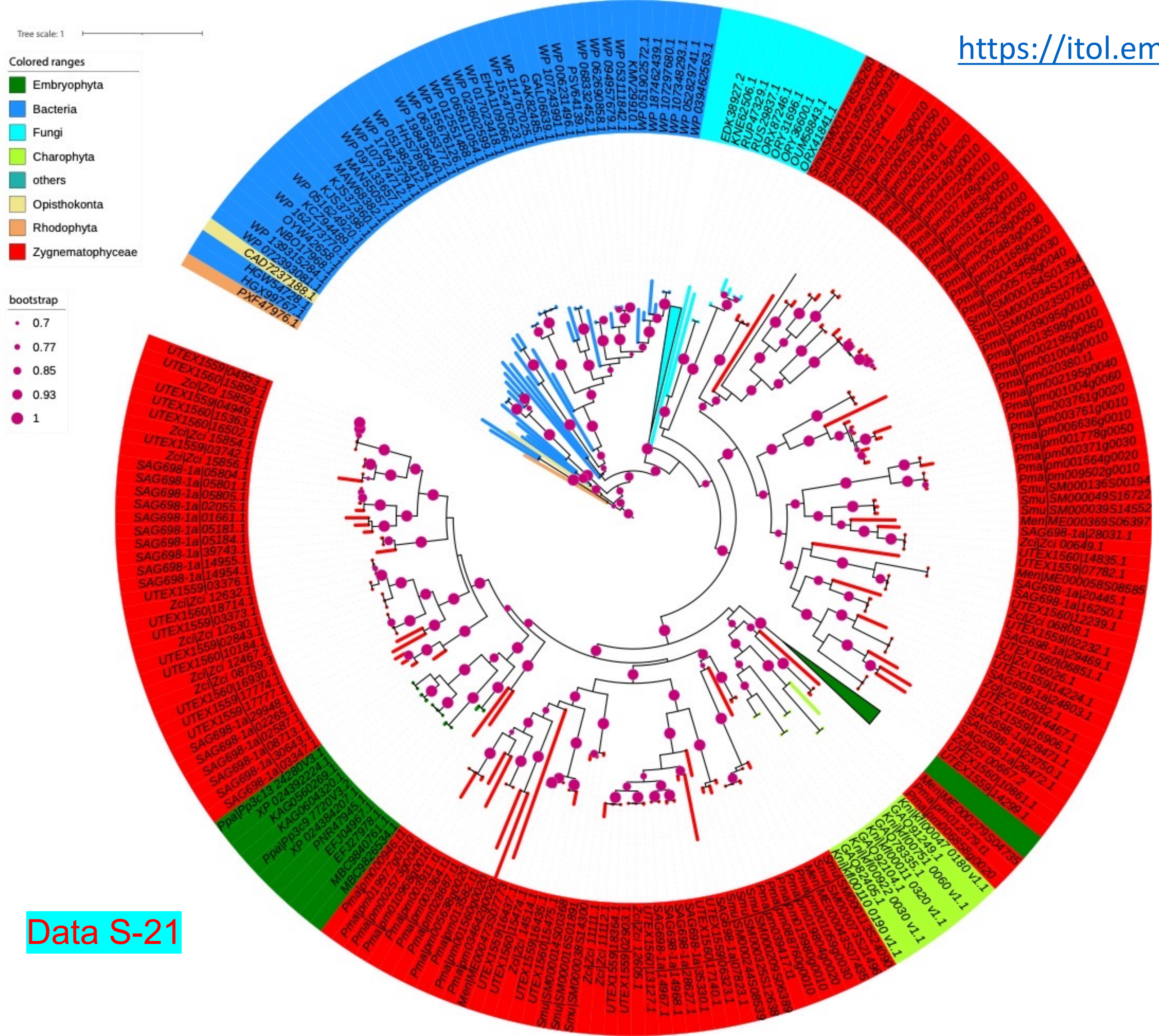


XTH might have been involved in HGT between fungi and ancient streptophyte algae (Shinohara and Nishitani, 2021).

XTH genes are expanded in Zygnemophyceae (1b has 16 genes)

Data S-21

Data S-22

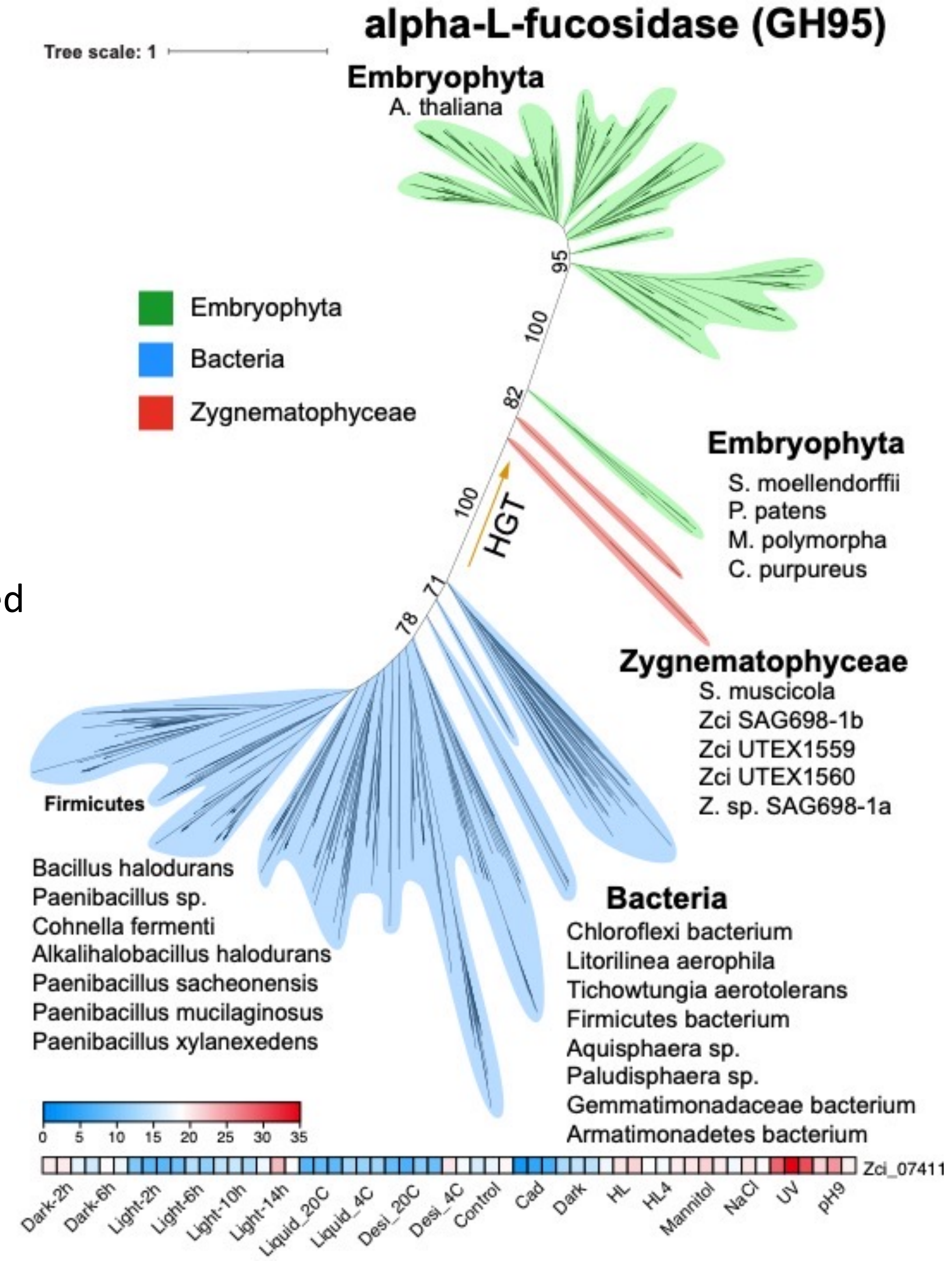


GH95: might have been gained via HGT from bacteria

### Data S-23

BGAL8 (AT3G28470)

Kotake T, Dina S, Konishi T, Kaneko S, Igarashi K, Samejima M, Watanabe Y, Kimura K, Tsumuraya Y. Molecular cloning of a {beta}-galactosidase from radish that specifically hydrolyzes {beta}-(1->3)- and {beta}-(1->6)-galactosyl residues of Arabinogalactan protein. Plant Physiol. 2005 Jul;138(3):1563-76. doi: 10.1104/pp.105.062562.

Sampedro et al. AtBGAL10 is the main xyloglucan β-galactosidase in Arabidopsis, and its absence results in unusual xyloglucan subunits and growth defects. Plant Physiol. 2012 Mar;158(3):1146-57. doi: 10.1104/pp.111.192195.

https://itol.embl.de/tree/20812724022273791665197706
https://itol.embl.de/tree/13423816417179911667574537

AtBGAL10 (AT5g63810)

Ishimaru M, Smith DL, Mort AJ, Gross KC. Enzymatic activity and substrate specificity of recombinant tomato beta-galactosidases 4 and 5. Planta. 2009 Jan;229(2):447-56. doi: 10.1007/s00425-008-0842-x.

BGAL12 (AT4G26140)

Eda M, Ishimaru M, Tada T, Sakamoto T, Kotake T, Tsumuraya Y, Mort AJ, Gross KC. Enzymatic activity and substrate specificity of the recombinant tomato β-galactosidase 1. J Plant Physiol. 2014 Oct 15;171(16):1454-60. doi: 10.1016/j.jplph.2014.06.010.

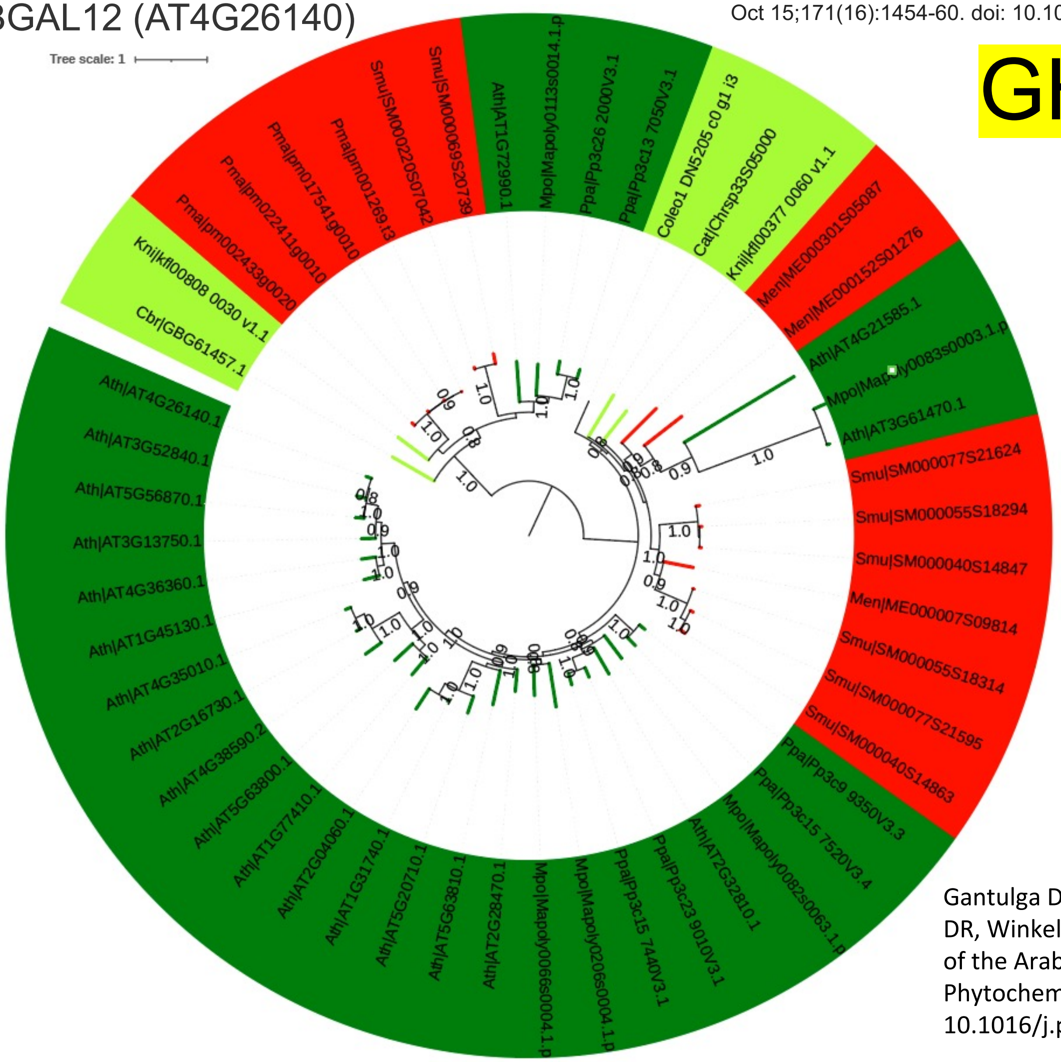


# GH35

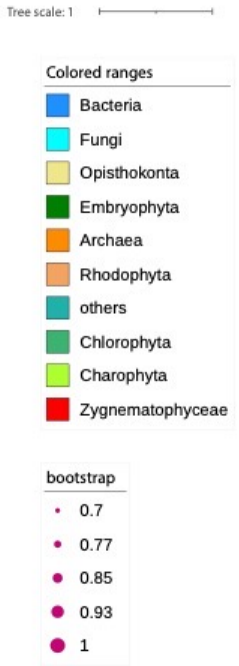


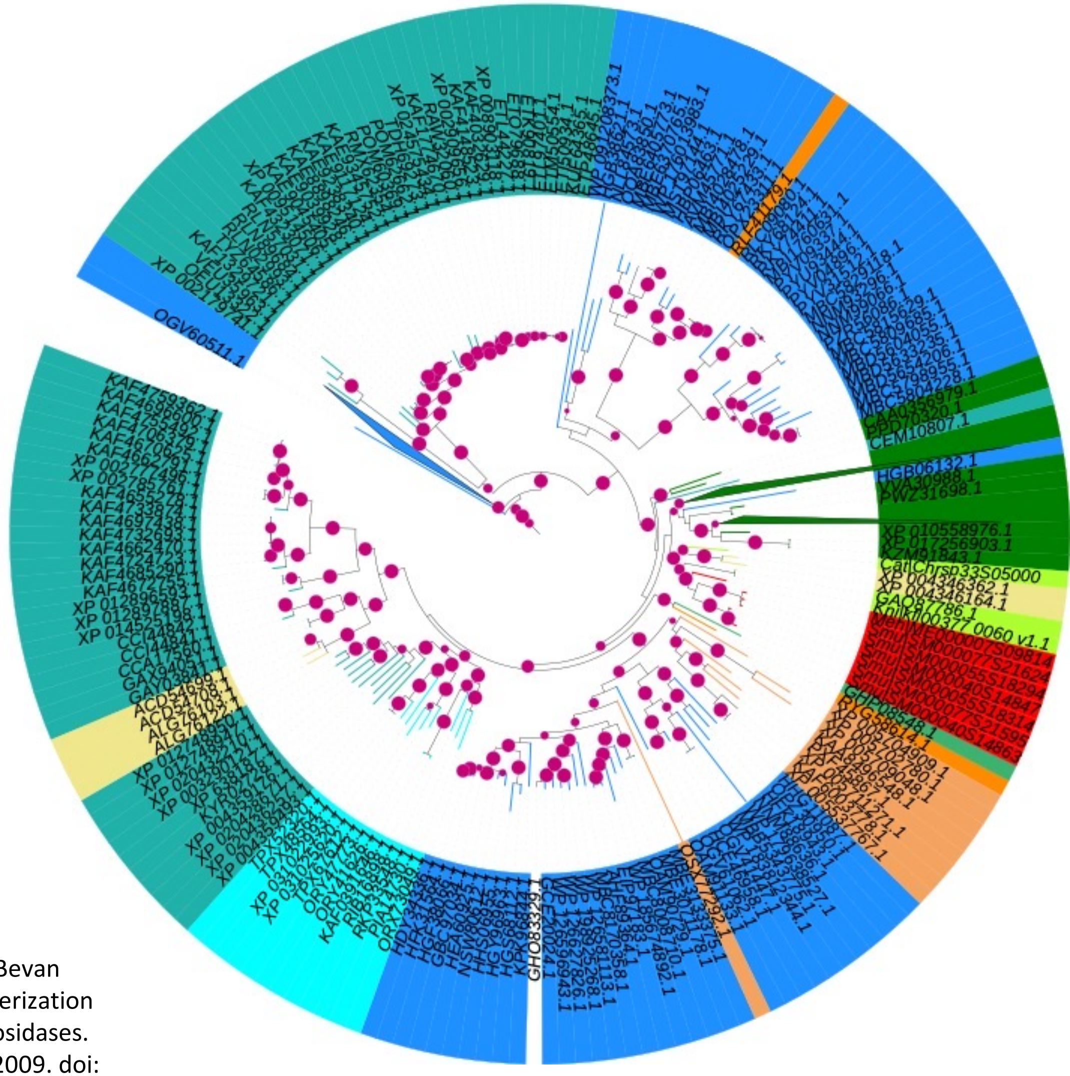

Gantulga D, Ahn YO, Zhou C, Battogtokh D, Bevan DR, Winkel BS, Esen A. Comparative characterization of the Arabidopsis subfamily a1 beta-galactosidases. Phytochemistry. 2009 Dec;70(17-18):1999-2009. doi: 10.1016/j.phytochem.2009.08.008.

Data S-24

# GH31

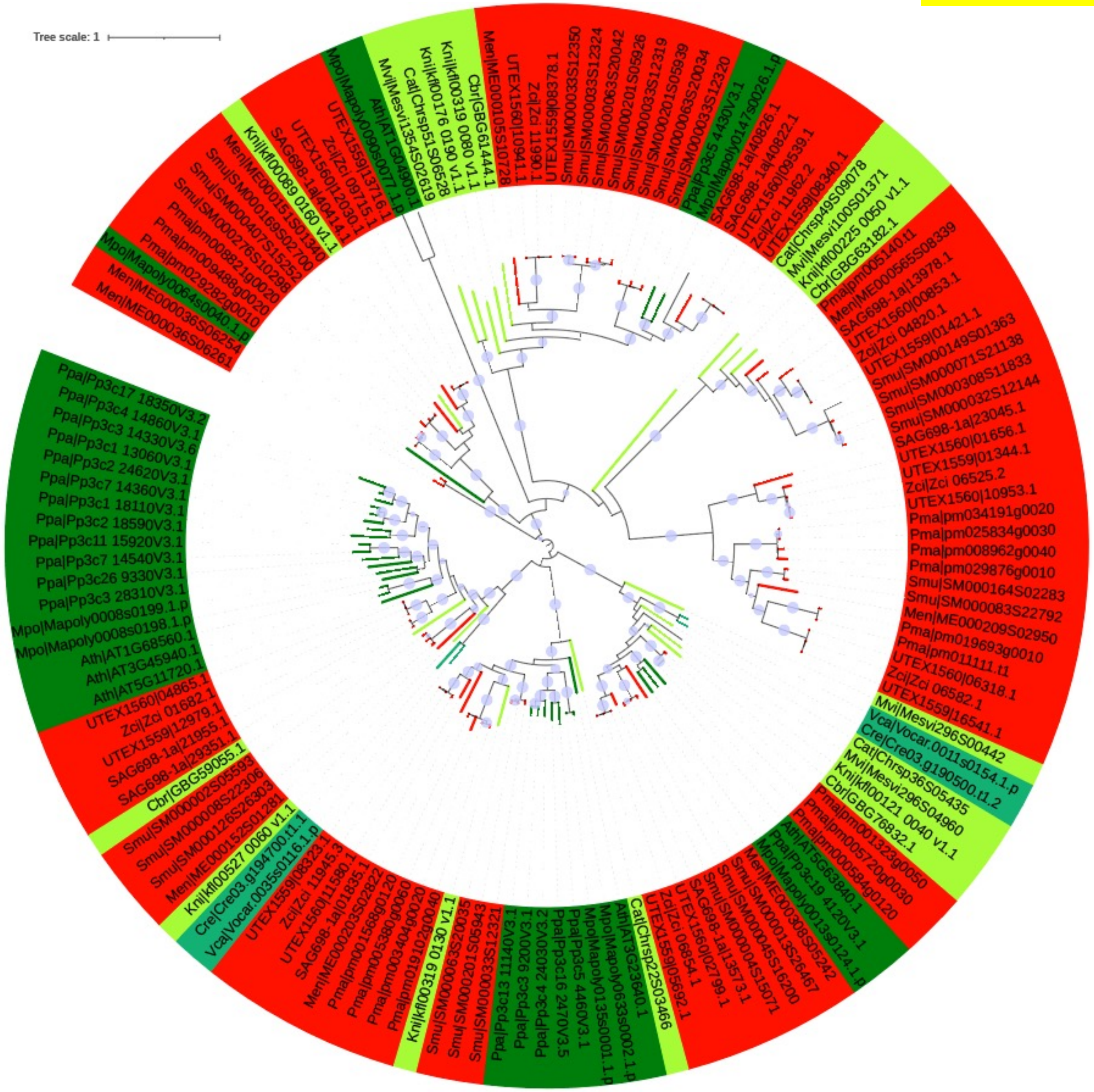


Plant Physiol . 2010 Nov;154(3):1105-15. doi: 10.1104/pp.110.163212. Epub 2010 Aug 26.

Lack of α-xylosidase activity in Arabidopsis alters xyloglucan composition and results in growth defects

**(AXY3)AT1G68560**

Javier Sampedro 1, Brenda Pardo, Cristina Gianzo, Esteban Guitián, Gloria Revilla, Ignacio Zarra
Affiliations expand
PMID: 20801759
PMCID: PMC2971592 DOI: 10.1104/pp.110.163212

Data S-25

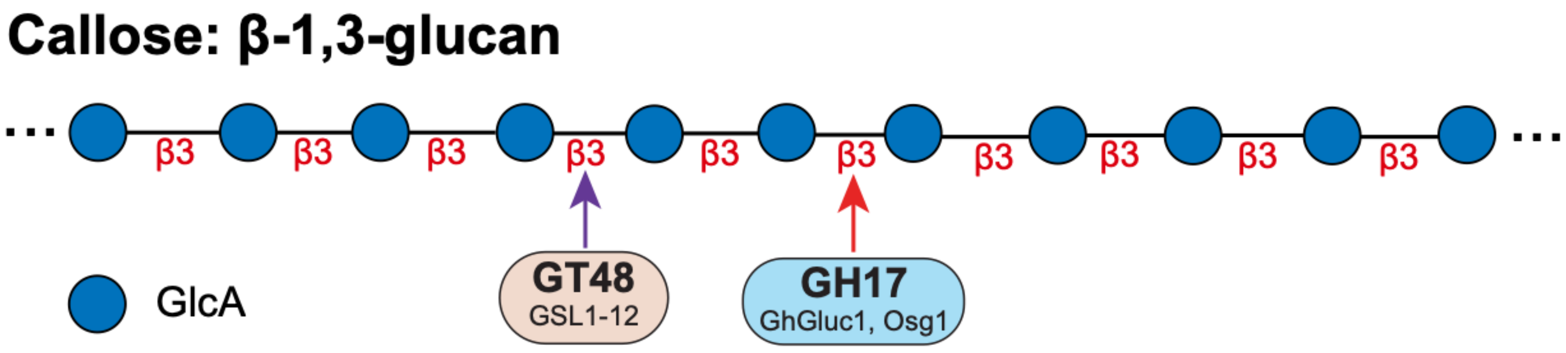


Scherp, P., Grotha, R., and Kutschera, U. (2001). Occurrence and phylogenetic significance of cytokinesis-related callose in green algae, bryophytes, ferns and seed plants. Plant Cell Reports *20*, 143-149.

Davis, D.J., Wang, M., Sorensen, I., Rose, J.K.C., Domozych, D.S., and Drakakaki, G. (2020). Callose deposition is essential for the completion of cytokinesis in the unicellular alga Penium margaritaceum. J Cell Sci *133*.

Data S-26

# GT48

#### 1,3-β-glucan synthase

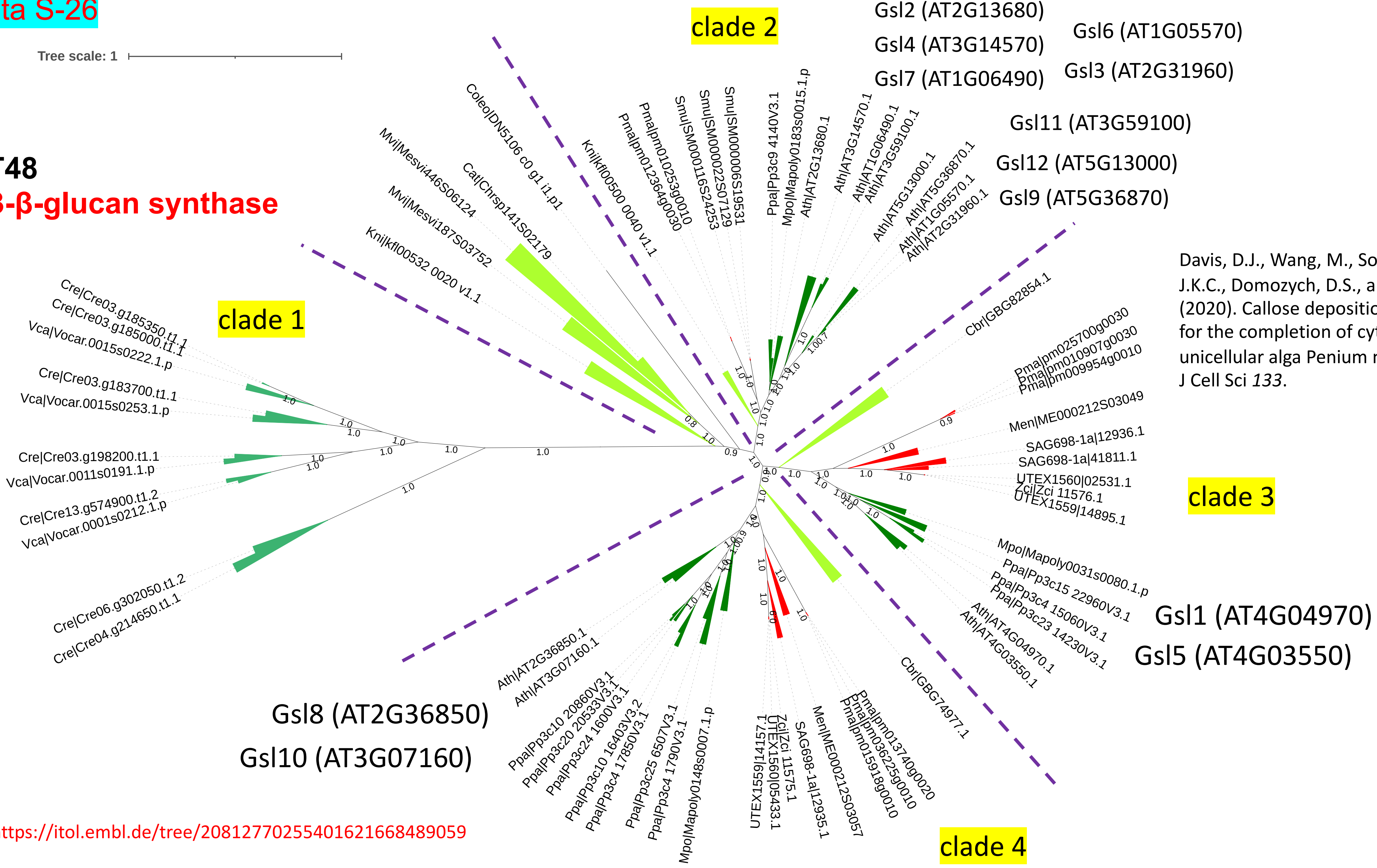


Davis, D.J., Wang, M., Sorensen, I., Rose, J.K.C., Domozych, D.S., and Drakakaki, G. (2020). Callose deposition is essential for the completion of cytokinesis in the unicellular alga Penium margaritaceum. J Cell Sci *133*.

https://itol.embl.de/tree/20812770255401621668489059

Data S-27

### GT48 nr hits

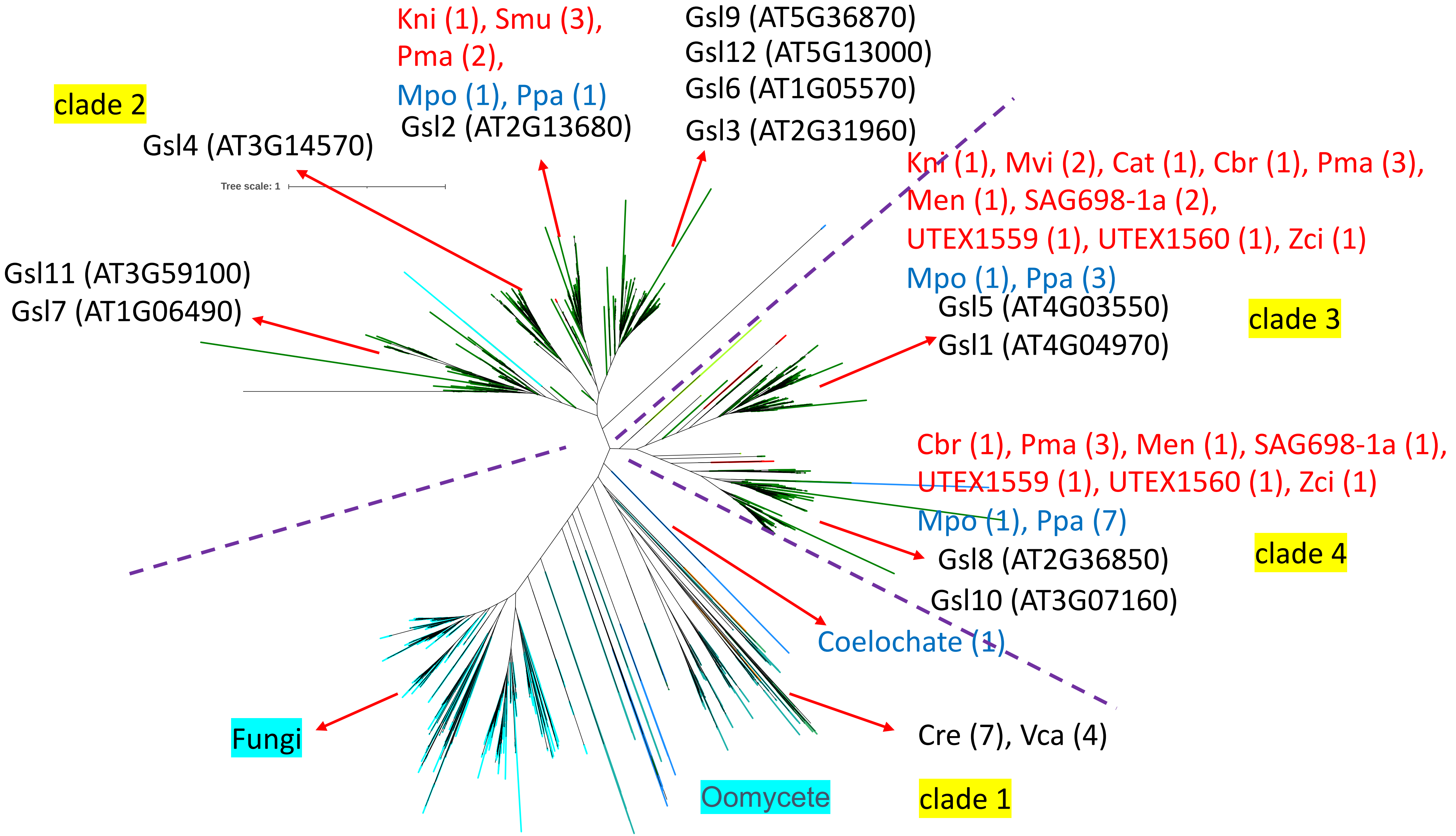


https://itol.embl.de/tree/2081279310513677166852 6914

Data S-28

# GH17

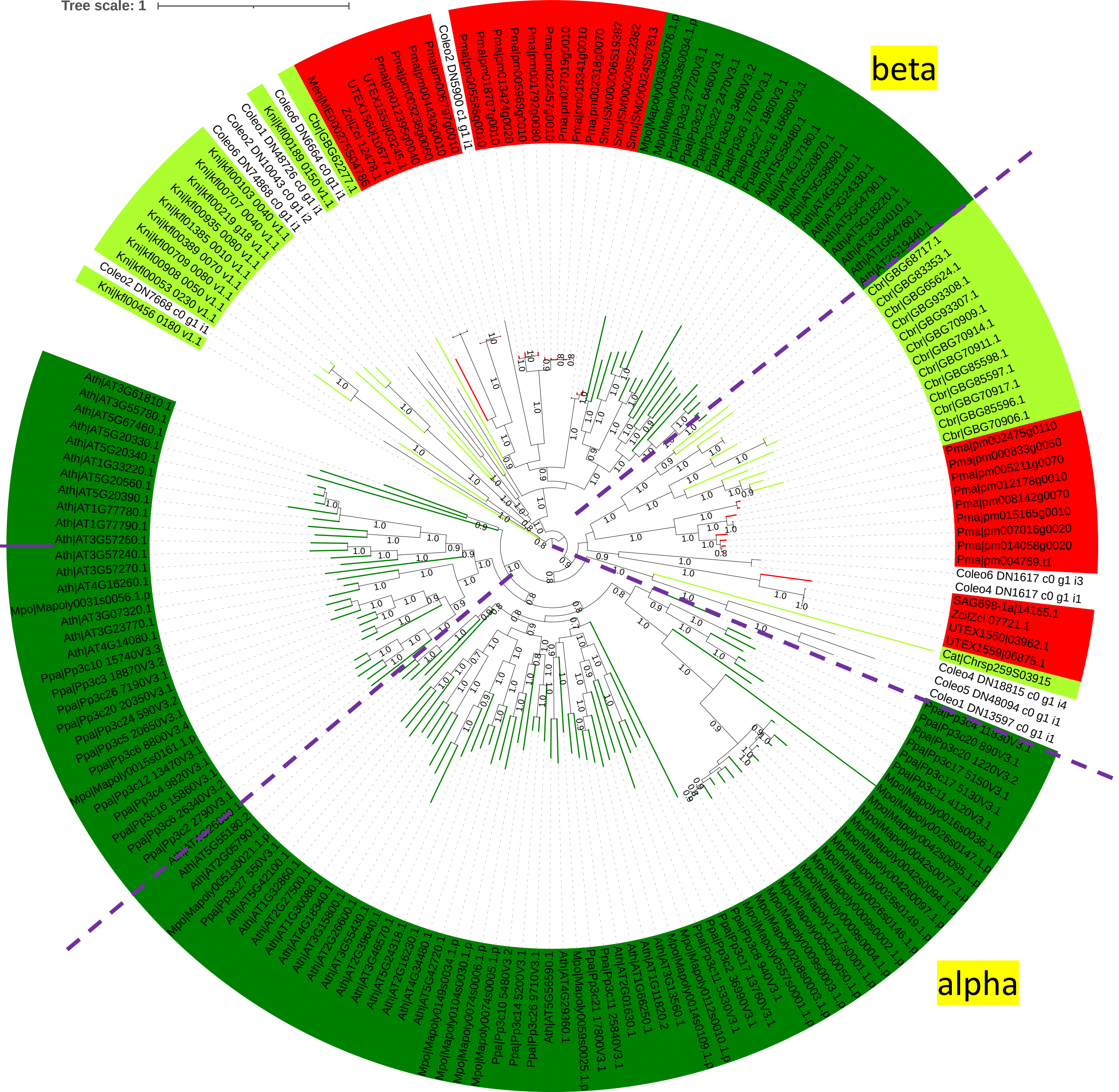


BG2
(AT3G57260)

PMID: 24904609

| class | Cre | Vca | Mvi | Cat | Kni | Cbr | Smu | Pma | Men | 1a | UTEX1559 | UTEX1560 | 1b | Mpo | Ppa | Ath |
|---|---|---|---|---|---|---|---|---|---|---|---|---|---|---|---|---|
| GH5_14 | 0 | 0 | 0 | 0 | 0 | 0 | 11 | 18 | 14 | 11 | 8 | 9 | 9 | 3 | 2 | 0 |

Data S-29

### exo-b-1,3-glucanase (EC 3.2.1.58)

https://itol.embl.de/tree/13423816417184651661223495

AT3G26130: e-value is low 9.1e-06

GH5_14 hmm

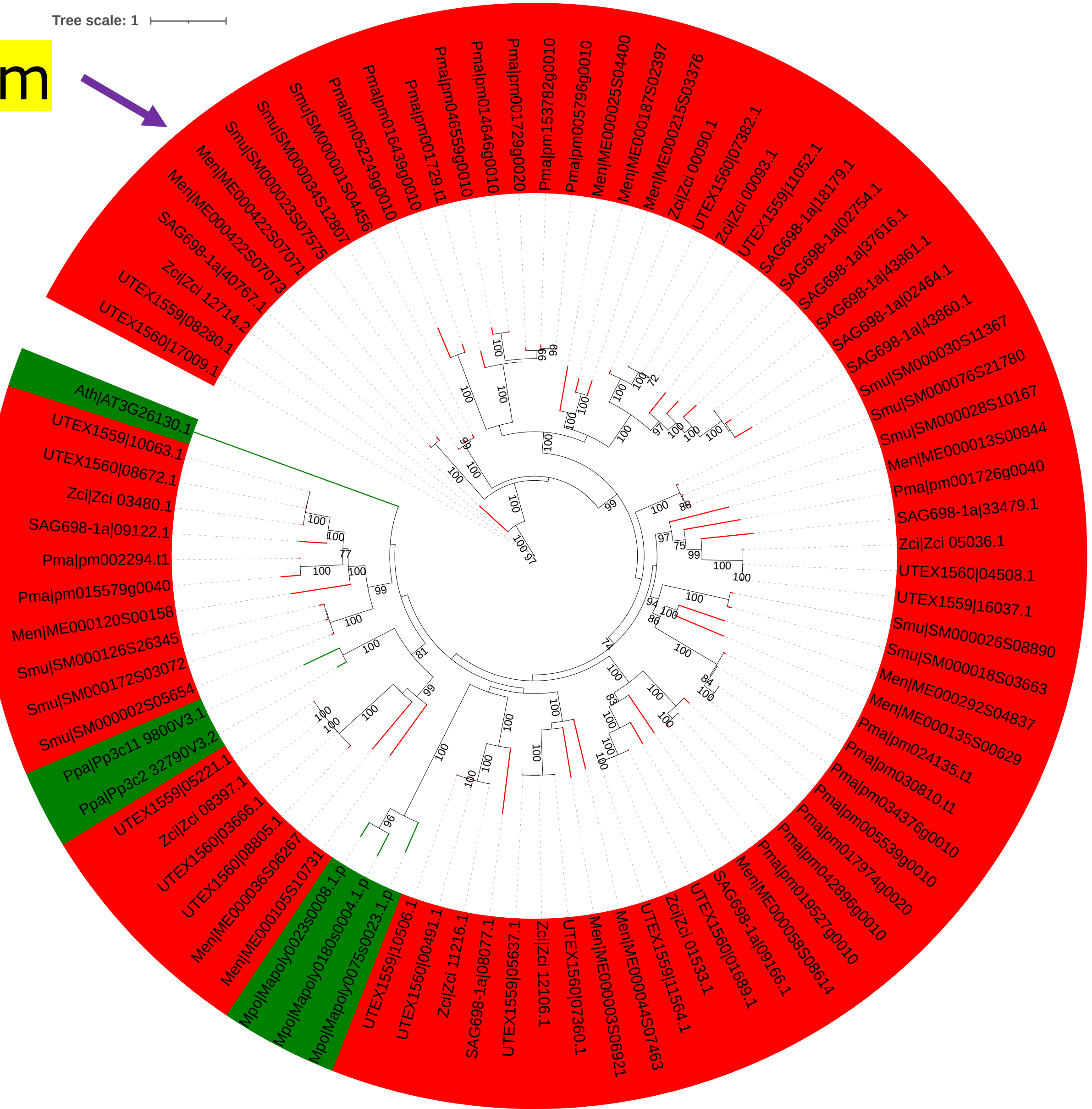


Zhou Y, Zeng L, Gui J, Liao Y, Li J, Tang J, Meng Q, Dong F, Yang Z. Functional characterizations of β-glucosidases involved in aroma compound formation in tea (Camellia sinensis). Food Res Int. 2017 Jun;96:206-214. doi: 10.1016/j.foodres.2017.03.049.

Opassiri R, Pomthong B, Akiyama T, Nakphaichit M, Onkoksoong T, Ketudat Cairns M, Ketudat Cairns JR. A stress-induced rice (Oryza sativa L.) beta-glucosidase represents a new subfamily of glycosyl hydrolase family 5 containing a fascin-like domain. Biochem J. 2007 Dec 1;408(2):241-9. doi: 10.1042/BJ20070734.

#### Glycoside Hydrolase Family 5 / Subf 14

GH5 Go GH5_14 Go ... Family Go

| | |
|---|---|
| **Activities in Sub Family** | b-glucosidase (EC 3.2.1.21);exo-b-1,3-glucanase (EC 3.2.1.58) |
| **Mechanism** | Retaining |
| **Clan** | GH-A |
| **3D Structure Status** | ( β / α ) $_8$ barrel |
| **Catalytic Nucleophile/Base** | Glu (experimental) |
| **Catalytic Proton Donor** | Glu (experimental) |
| **Note** | Once known as cellulase family A; many members have been assigned to subfamilies as described by Aspeborg et al. (2012) BMC Evol Biol. 12(1):186 (PMID: 22992189). |
| **External resources** | CAZypedia; HOMSTRAD; PROSITE; |
| **Commercial Enzyme Provider(s)** | MEGAZYME; NZYTech; PROZOMIX; |
| **Statistics** | GenBank accession (103); Uniprot accession (3); |

Summary | Download GH5_14 (83) | Taxonomic display | Characterized (2)

##### Eukaryota

| Protein Name | EC# | Reference | Organism | GenBank | Uniprot | PDB/3D | Subf |
|---|---|---|---|---|---|---|---|
| β-glucosidase 1 GH5 family (GH5BG1;CsGH5BG1) | 3.2.1.21 | pubmed | Camellia sinensis | **ARU79086.1** | A0A1Y0K2D4 | | 14 |
| exo-β-1,3-glucosidase (GH5BG;Os10g0370500) | 3.2.1.58 | pubmed | Oryza sativa Japonica Group | **AAM08614.1** AAM08821.1 AAP53379.1 BAF26372.1 BAG89316.1 BAT10560.1 | Q0IY43 Q8RU06 | | 14 |

Top

Data S-30

GH5_14 nr
e-15

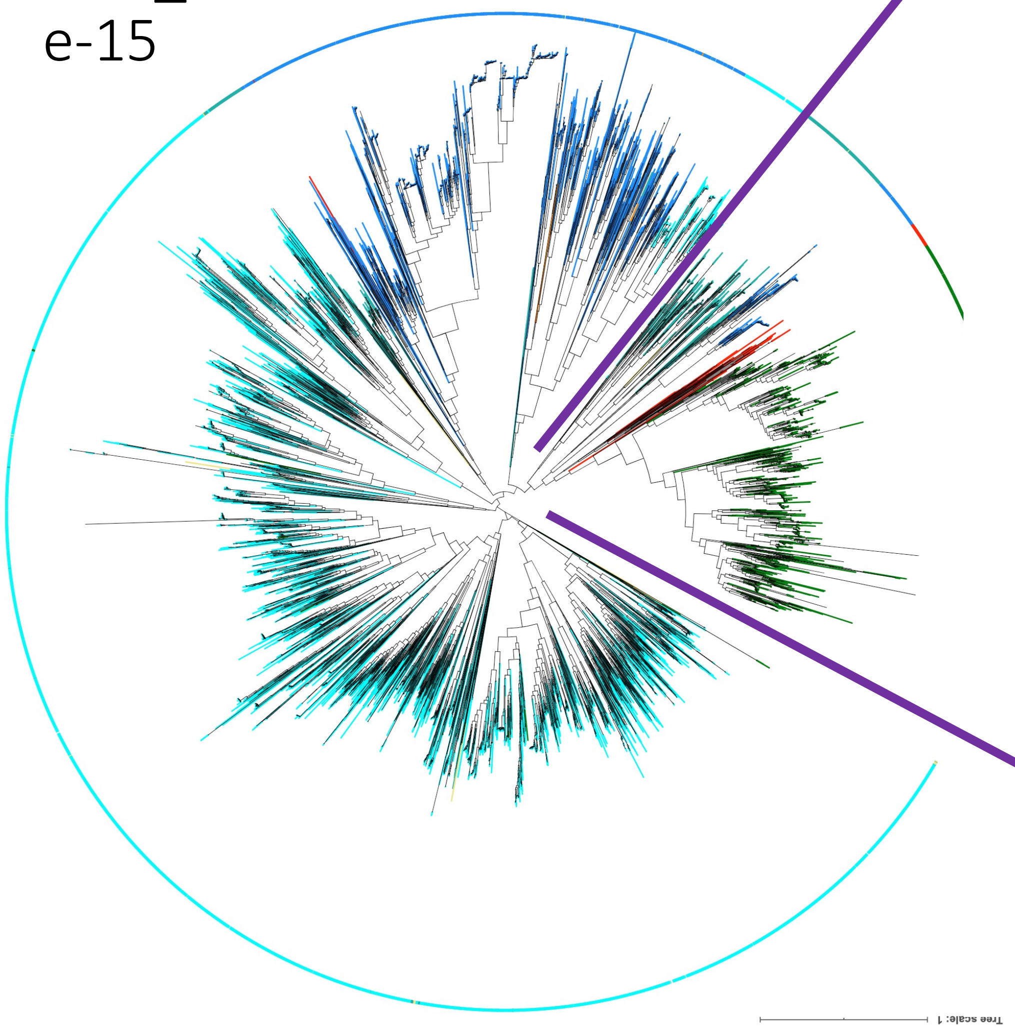


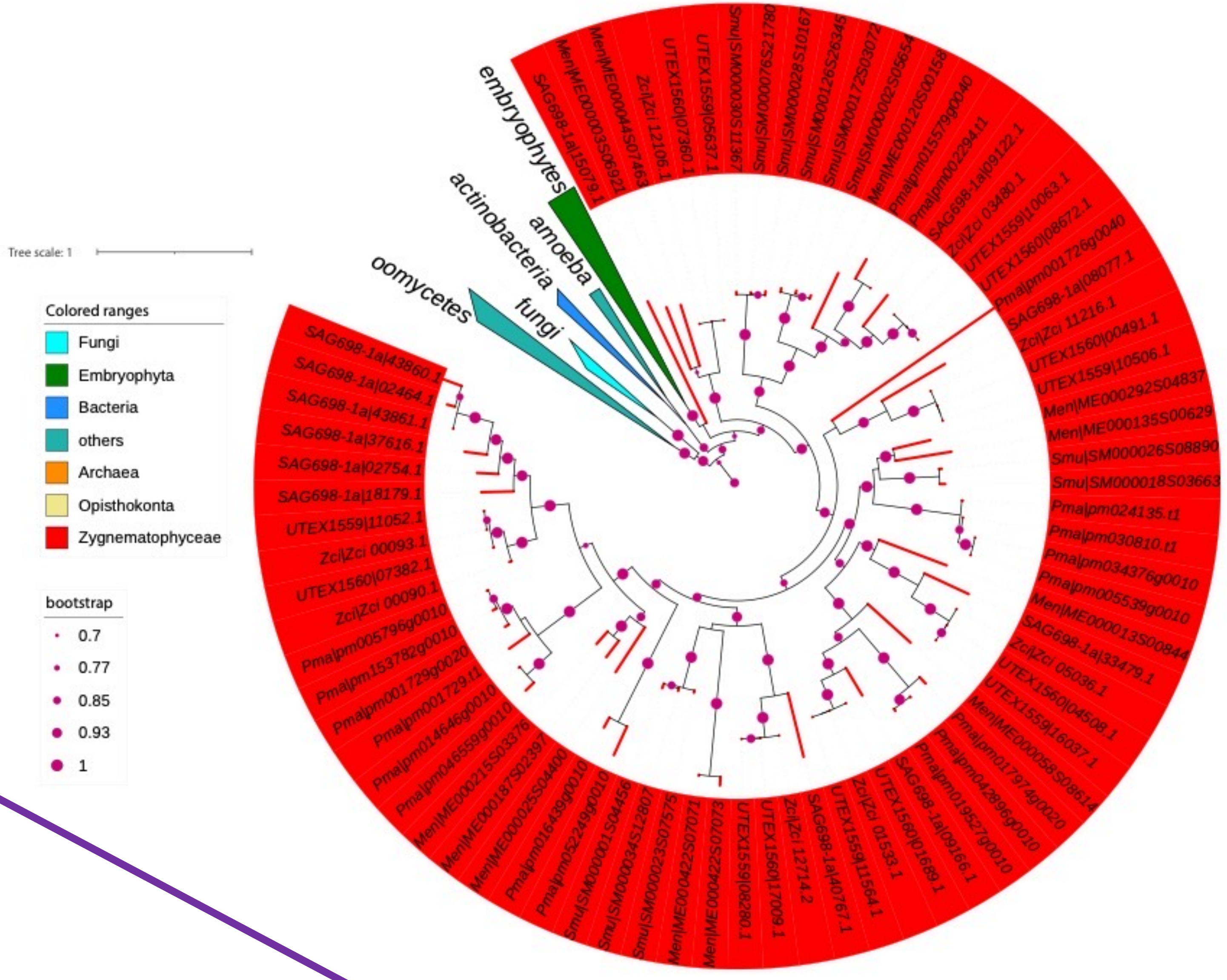


Possible gain via HGT from bacteria

Data S-31

### The combination of Fascin domain and GH5_14 is first present in LCA of Zygnematophyceae & land plants

#### Conserved domains on [gi|19920182|gb|AAM08614|]

View Standard Results

Putative Glucan 1,3-beta-glucosidase precursor [Oryza sativa Japonica Group]

**Graphical summary** Zoom to residue level show extra options »

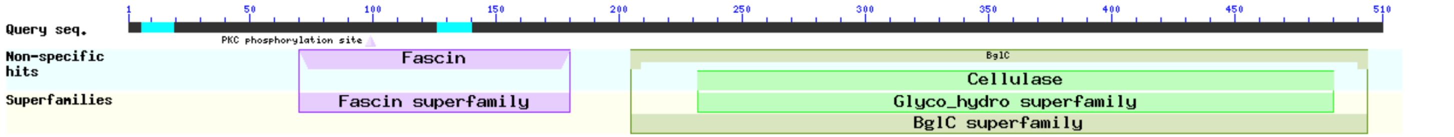


Search for similar domain architectures | Refine search

**List of domain hits**

| Name | Accession | Description | Interval | E-value |
|---|---|---|---|---|
| [+] BglC | COG2730 | Aryl-phospho-beta-D-glucosidase BglC, GH1 family [Carbohydrate transport and metabolism]; | 205-504 | 2.41e-29 |
| [+] Cellulase | pfam00150 | Cellulase (glycosyl hydrolase family 5); | 232-490 | 8.40e-14 |
| [–] Fascin | cd00257 | Fascin-like domain; members include actin-bundling/crosslinking proteins fascin, ... | 70-180 | 3.74e-03 |

Fascin-like domain; members include actin-bundling/crosslinking proteins fascin, histoactophilin and singed; identified in sea urchin, Drosophila, Xenopus, rodents, and humans; The fascin-like domain adopts a beta-trefoil topology and contains an internal threefold repeat; the fascin subgroup contains four copies of the domain; Structurally similar to fibroblast growth factor (FGF)

**Pssm-ID:** 238160 **Cd Length:** 119 **Bit Score:** 37.25 **E-value:** 3.74e-03

```
                      10        20        30        40        50        60        70        80
              ....*....|....*....|....*....|....*....|....*....|....*....|....*....|....*....|
gi 19920182  70 TQLQFKSVTQNmYLCAEQGGGTiLVANRTSASGWETFKLWRIDEDTFDLRVFDNLFVTVAGDGVTVVATvaSPGPGEAFQ 149
Cdd:cd00257   2 PQVVLRSVNGR-YLSAEAGGDK-VDANRDSLKGDETFTLEFDNTGKYALRSHDGKYLSADSDGGVQLEG--HPNADCRFT 77

                      90       100       110
              ....*....|....*....|....*....|.
gi 19920182 150 IVRNGDkTRARIRAPNGMFLQAKTSDSVTAD 180
Cdd:cd00257  78 LEFHGD-GKWALRAENGRYLGGDGSGTLKAS 107
```

rice GH5BG (Fascin domain)

#### Conserved domains on [lcl|Query_8185]

View Standard Results

Local query sequence

**Graphical summary** Zoom to residue level show extra options »

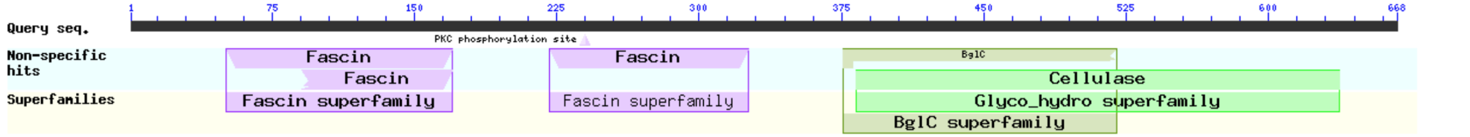


Search for similar domain architectures | Refine search

**List of domain hits**

| Name | Accession | Description | Interval | E-value |
|---|---|---|---|---|
| [+] BglC | COG2730 | Aryl-phospho-beta-D-glucosidase BglC, GH1 family [Carbohydrate transport and metabolism]; | 376-520 | 7.10e-17 |
| [+] Cellulase | pfam00150 | Cellulase (glycosyl hydrolase family 5); | 383-637 | 2.23e-09 |
| [+] Fascin | cd00257 | Fascin-like domain; members include actin-bundling/crosslinking proteins fascin, ... | 221-326 | 2.79e-09 |
| [+] Fascin | pfam06268 | Fascin domain; This family consists of several eukaryotic fascin or singed proteins. The ... | 51-170 | 1.19e-06 |
| [–] Fascin | cd00257 | Fascin-like domain; members include actin-bundling/crosslinking proteins fascin, ... | 91-170 | 4.00e-06 |

Fascin-like domain; members include actin-bundling/crosslinking proteins fascin, histoactophilin and singed; identified in sea urchin, Drosophila, Xenopus, rodents, and humans; The fascin-like domain adopts a beta-trefoil topology and contains an internal threefold repeat; the fascin subgroup contains four copies of the domain; Structurally similar to fibroblast growth factor (FGF)

**Pssm-ID:** 238160 **Cd Length:** 119 **Bit Score:** 46.11 **E-value:** 4.00e-06

```
                      10        20        30        40        50        60        70        80
              ....*....|....*....|....*....|....*....|....*....|....*....|....*....|....*....|
Query_8185   91 VAFRGPNGKLLAAHADATVTASGEASgtdipTECMFRVEKKPNKKFTIWSVsNSYYLRAWDNGTLRLHMKAPTSWELFSI 170
Cdd:cd00257  46 YALRSHDGKYLSADSDGGVQLEGHPN-----ADCRFTLEFHGDGKWALRAE-NGRYLGGDGSGTLKASSETVGPDELFEL 119
```

Zci|Zci_03480.1 (Fascin domain)

Data S-32

### Xylan / AX / GX / GAX

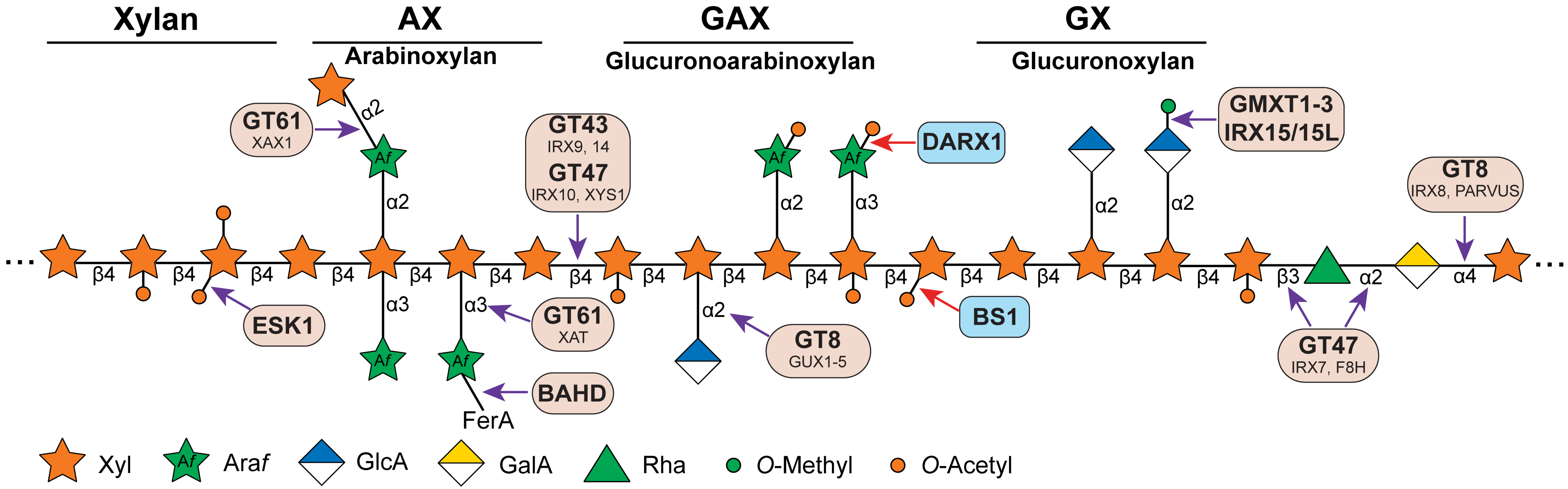

# GT47

Tree scale: 1

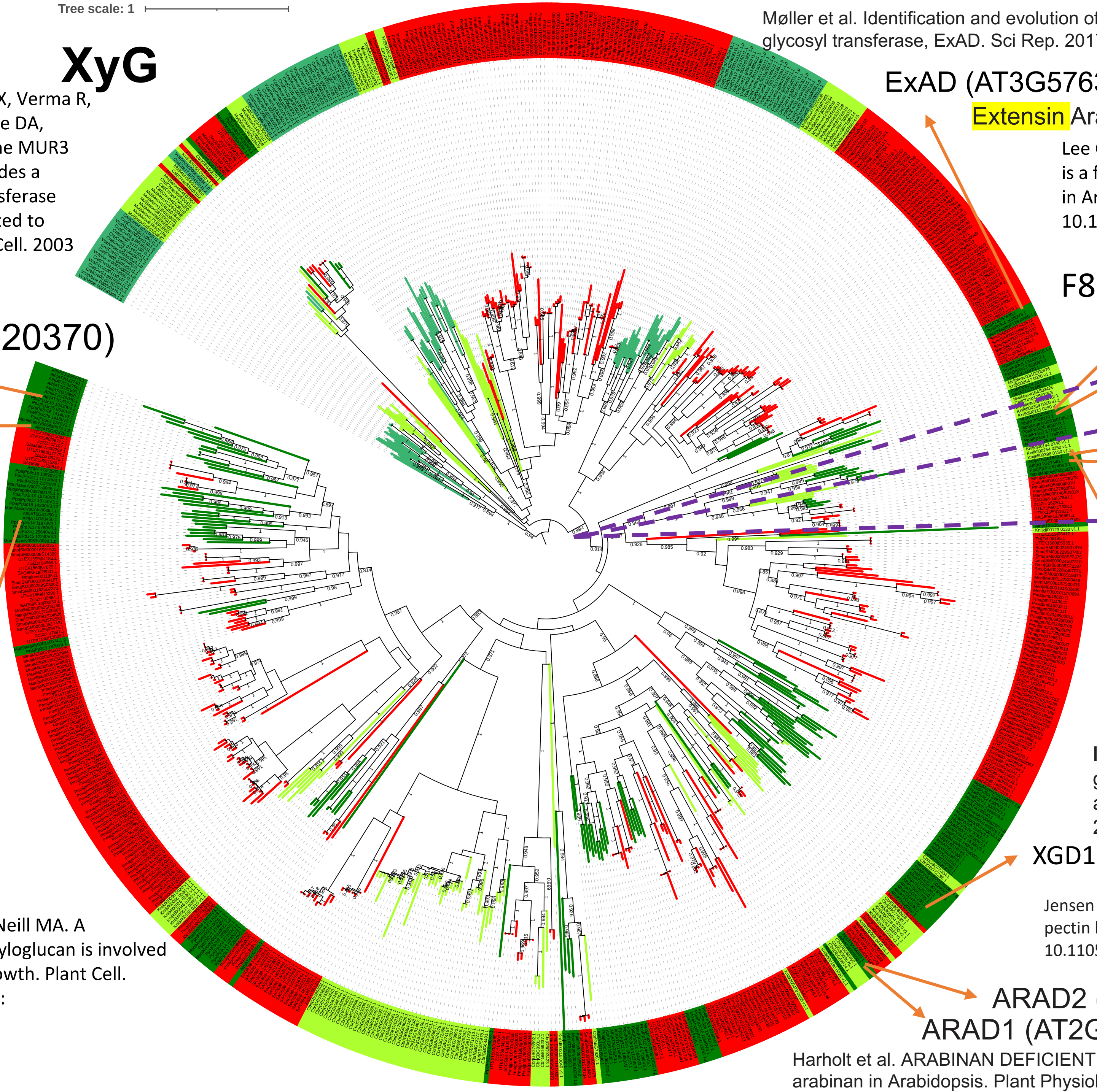

#### XyG

Madson M, Dunand C, Li X, Verma R, Vanzin GF, Caplan J, Shoue DA, Carpita NC, Reiter WD. The MUR3 gene of Arabidopsis encodes a xyloglucan galactosyltransferase that is evolutionarily related to animal exostosins. Plant Cell. 2003 Jul;15(7):1662-70. doi: 10.1105/tpc.009837.

MUR3 (AT2G20370)

XLT2 (AT5G62220)

Jensen JK, Schultink A, Keegstra K, Wilkerson CG, Pauly M. RNA-Seq analysis of developing nasturtium seeds (Tropaeolum majus): identification and characterization of an additional galactosyltransferase involved in xyloglucan biosynthesis. Mol Plant. 2012 Sep;5(5):984-92. doi: 10.1093/mp/sss032.

XUT1 (AT1G63450)

Peña MJ, Kong Y, York WS, O'Neill MA. A galacturonic acid-containing xyloglucan is involved in Arabidopsis root hair tip growth. Plant Cell. 2012 Nov;24(11):4511-24. doi: 10.1105/tpc.112.103390.

Møller et al. Identification and evolution of a plant cell wall specific glycoprotein glycosyl transferase, ExAD. Sci Rep. 2017 Mar 30;7:45341. doi: 10.1038/srep45341

ExAD (AT3G57630)

Extensin Arabinose Deficient transferase

Lee C, Teng Q, Huang W, Zhong R, Ye ZH. The F8H glycosyltransferase is a functional paralog of FRA8 involved in glucuronoxylan biosynthesis in Arabidopsis. Plant Cell Physiol. 2009 Apr;50(4):812-27. doi: 10.1093/pcp/pcp025. Epub 2009 Feb 18. PMID: 19224953.

F8H (AT5G22940)

IRX7/FRA8 (AT2G28110)

Brown, D.M., Goubet, F., Wong, V.W., Goodacre, R., Stephens, E., Dupree, P., and Turner, S.R. (2007). Comparison of five xylan synthesis mutants reveals new insight into the mechanisms of xylan synthesis. Plant J. 52: 1154–1168.

XYS1/IRX10-L (AT5G61840)

IRX10 (AT1G27440)

#### Xylan

Jensen JK, Busse-Wicher M, Poulsen CP, Fangel JU, Smith PJ, Yang JY, Peña MJ, Dinesen MH, Martens HJ, Melkonian M et al. 2018. Identification of an algal xylan synthase indicates that there is functional orthology between algal and plant cell wall biosynthesis. New Phytologist 218: 1049–1060.

Urbanowicz BR, Peña MJ, Moniz HA, Moremen KW, York WS. 2014. Two Arabidopsis proteins synthesize acetylated xylan in vitro. Plant Journal 80: 197–206.

NpGUT1 is a pectin b-glucuronyltransferase that transfers glucuronic acid to RG-II

Iwai H, Masaoka N, Ishii T, Satoh S. A pectin glucuronyltransferase gene is essential for intercellular attachment in the plant meristem. Proc Natl Acad Sci U S A. 2002 Dec 10;99(25):16319-24. doi: 10.1073/pnas.252530499.

XGD1 (AT5G33290)

#### XGA

Jensen et al. Identification of a xylogalacturonan xylosyltransferase involved in pectin biosynthesis in Arabidopsis. Plant Cell. 2008 May;20(5):1289-302. doi: 10.1105/tpc.107.050906

ARAD2 (AT5G44930)

ARAD1 (AT2G35100)

## RG-I

Harholt et al. ARABINAN DEFICIENT 1 is a putative arabinosyltransferase involved in biosynthesis of pectic arabinan in Arabidopsis. Plant Physiol. 2006 Jan;140(1):49-58. doi: 10.1104/pp.105.072744.

Data S-34

https://itol.embl.de/tree/1342381681245654166831464

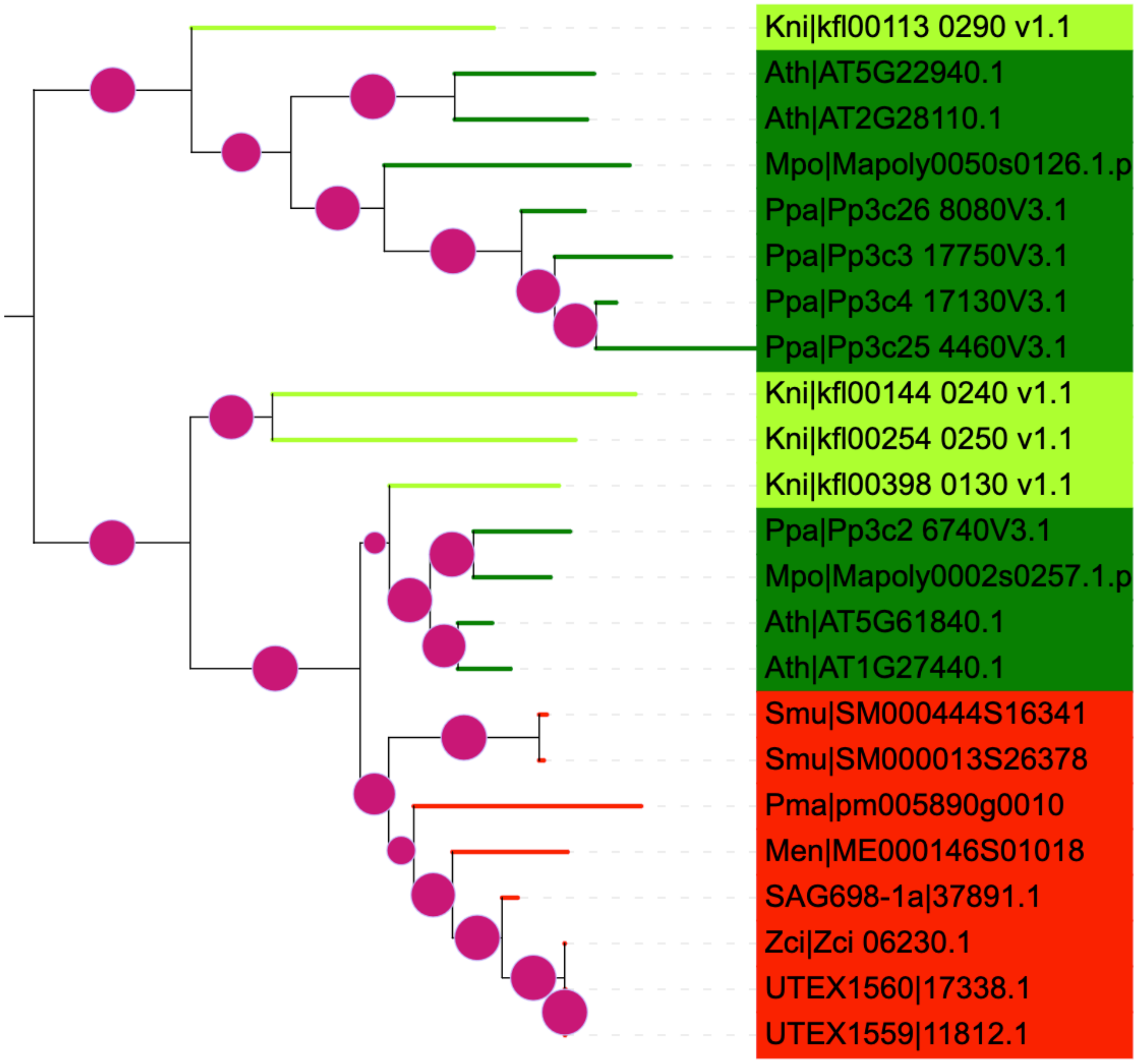


F8H (AT5G22940)
IRX7/FRA8 (AT2G28110)

XYS1/IRX10-L (AT5G61840)
IRX10 (AT1G27440)

GT47

Data S-35

https://itol.embl.de/tree/208127240222430161661878342

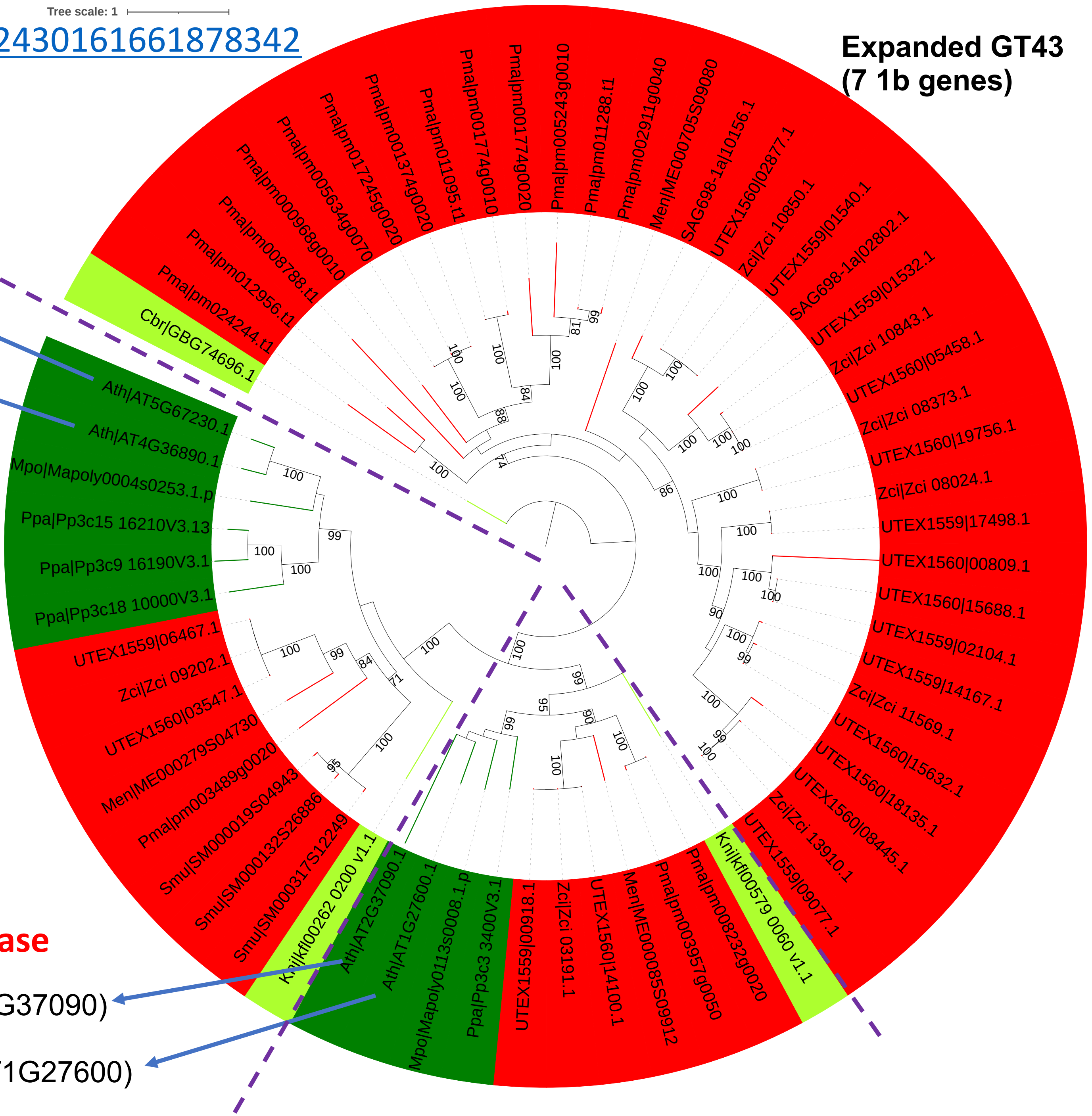


**glucuronoxylan glycosyltransferase**

IRX14-L (AT5G67230)

IRX14 (AT4G36890)

secondary wall thickening in fibers

Brown, D.M., Goubet, F., Wong, V.W., Goodacre, R., Stephens, E., Dupree, P., and Turner, S.R. (2007). Comparison of five xylan synthesis mutants reveals new insight into the mechanisms of xylan synthesis. Plant J. 52: 1154–1168.

Lee C, Teng Q, Huang W, Zhong R, Ye ZH. The Arabidopsis family GT43 glycosyltransferases form two functionally nonredundant groups essential for the elongation of glucuronoxylan backbone. Plant Physiol. 2010 Jun;153(2):526-41. doi: 10.1104/pp.110.155309.

Ren Y, Hansen SF, Ebert B, Lau J, Scheller HV. Site-directed mutagenesis of IRX9, IRX9L and IRX14 proteins involved in xylan biosynthesis: glycosyltransferase activity is not required for IRX9 function in Arabidopsis. PLoS One. 2014 Aug 13;9(8):e105014. doi: 10.1371/journal.pone.0105014.

# GT43

**xylan β-1,4-xylosyltransferase**

IRX9 (AT4G37090)

IRX9-L (AT1G27600)

Data S-36

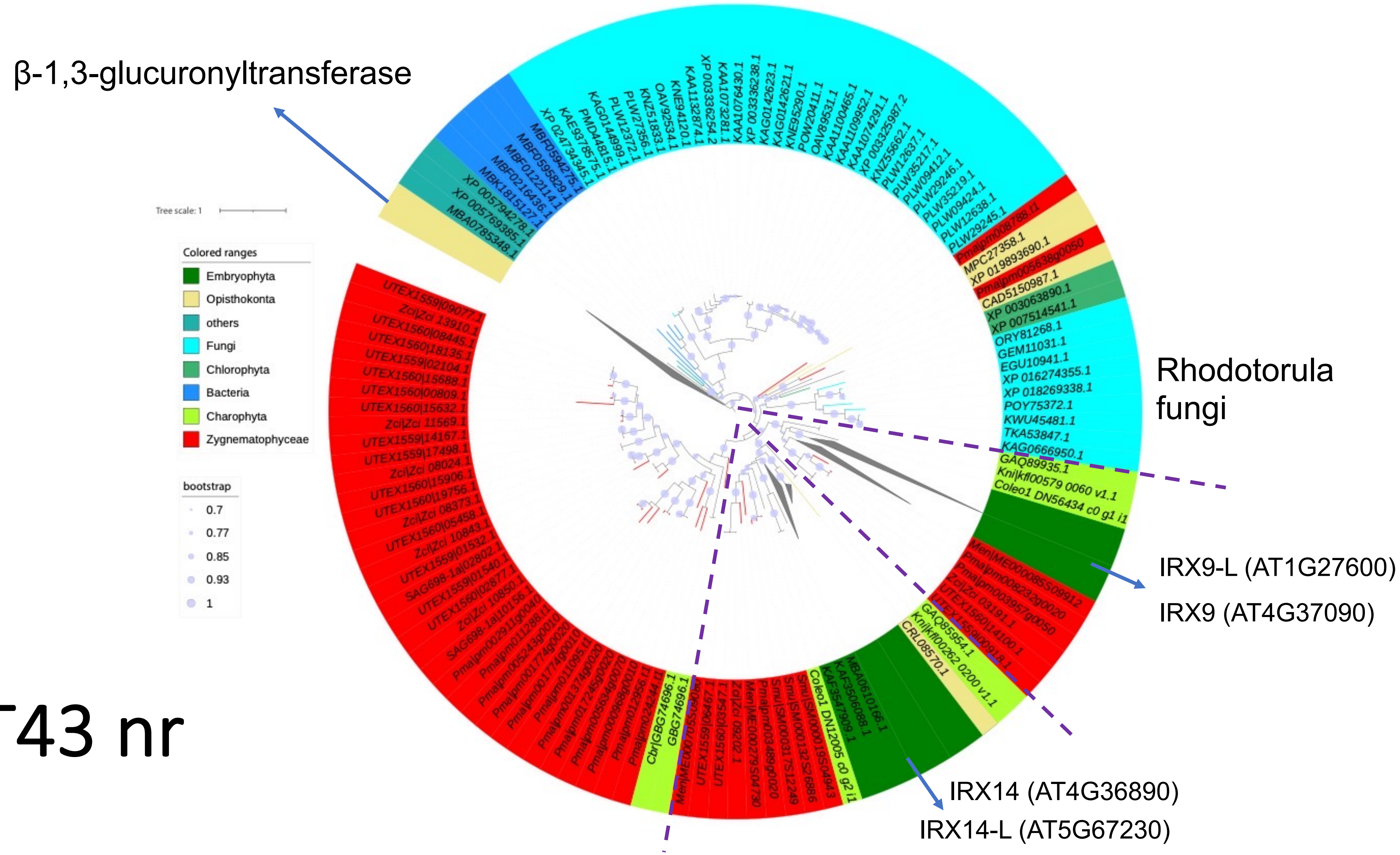


# GT43 nr

might have been gained via HGT from fungi

Data S-37

### GT8 (dbcan) + Coleochaete

#### Glucuronyltransferases of Xylan (first present in Zygnematophyceae)

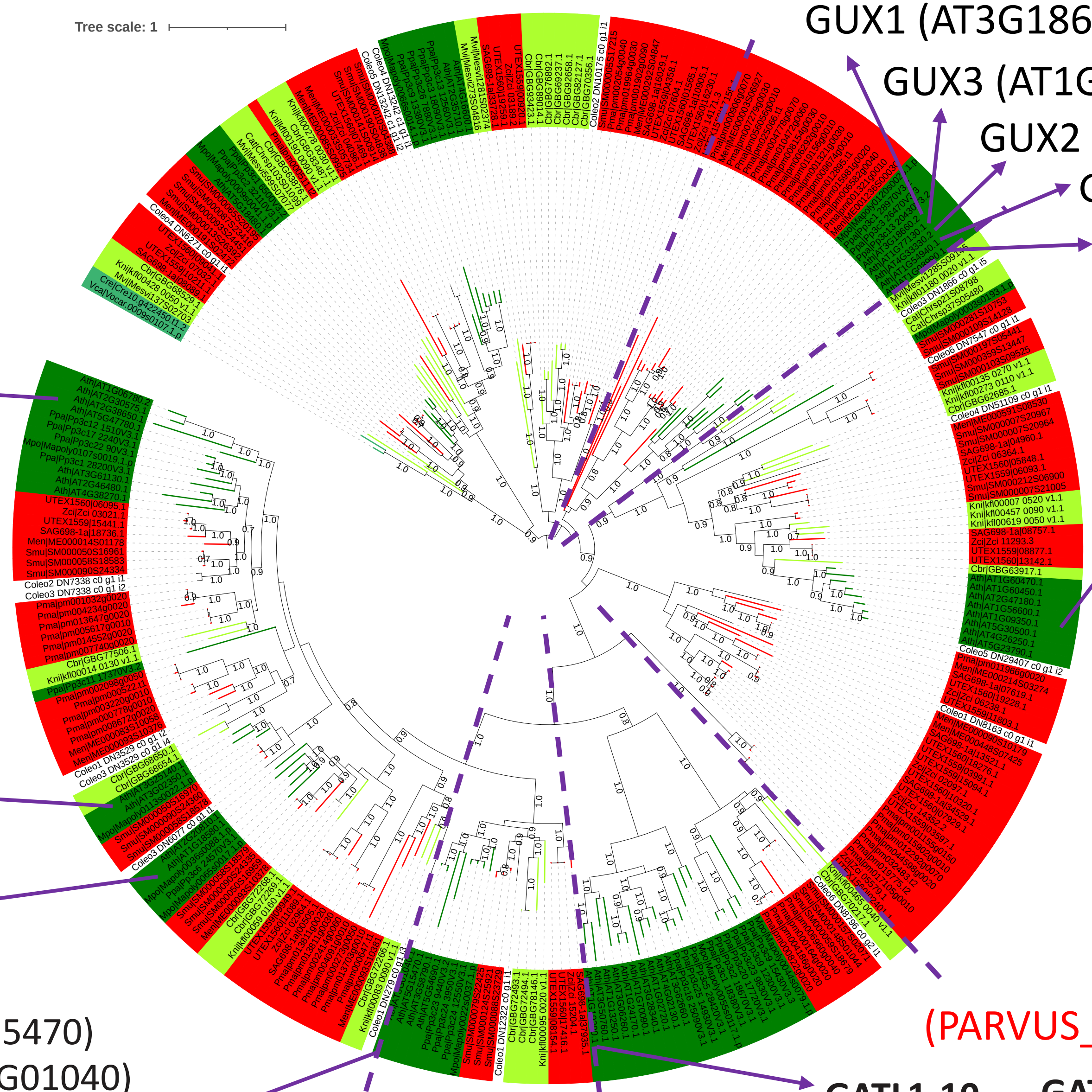


Rennie EA, Hansen SF, Baidoo EEK, Hadi MZ, Keasling JD, Scheller HV. 2012. Three members of the Arabidopsis glycosyltransferase family 8 are xylan glucuronosyltransferases. Plant Physiology 159: 1408–1417.

Data S-38

https://itol.embl.de/tree/2081272402222635216653524 15

(PARVUS_GATL: Xylan)

**GAUT12** (AT5G54690) (IRX8: Xylan)

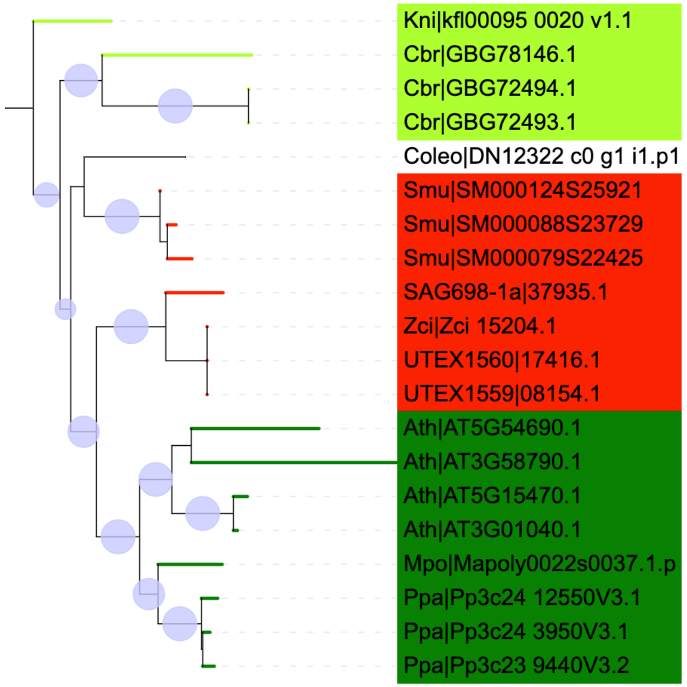


**GATL1** (AT1G19300)

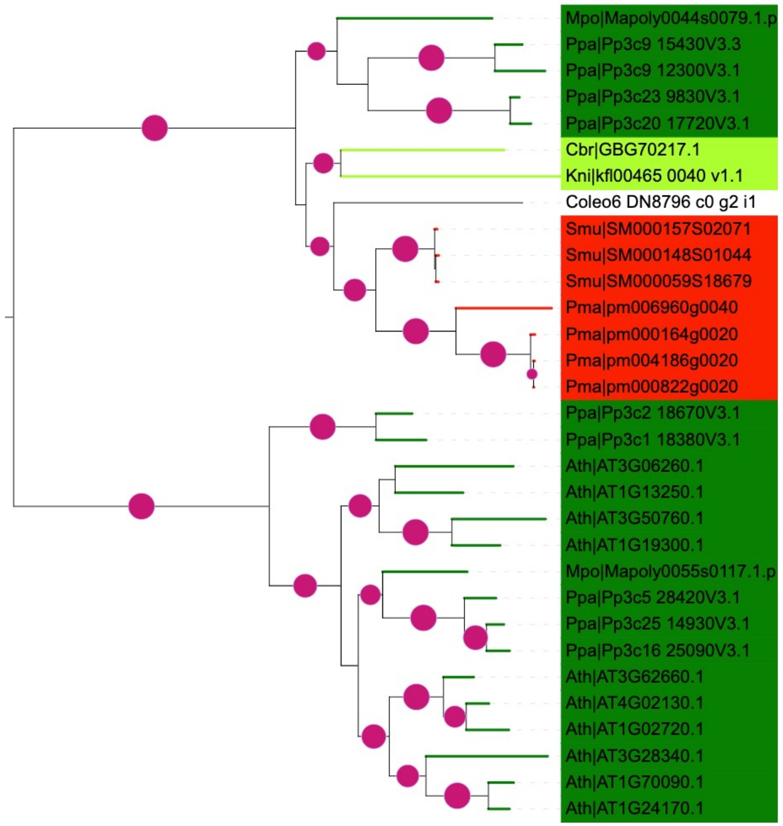


GUX1 (AT3G18660)

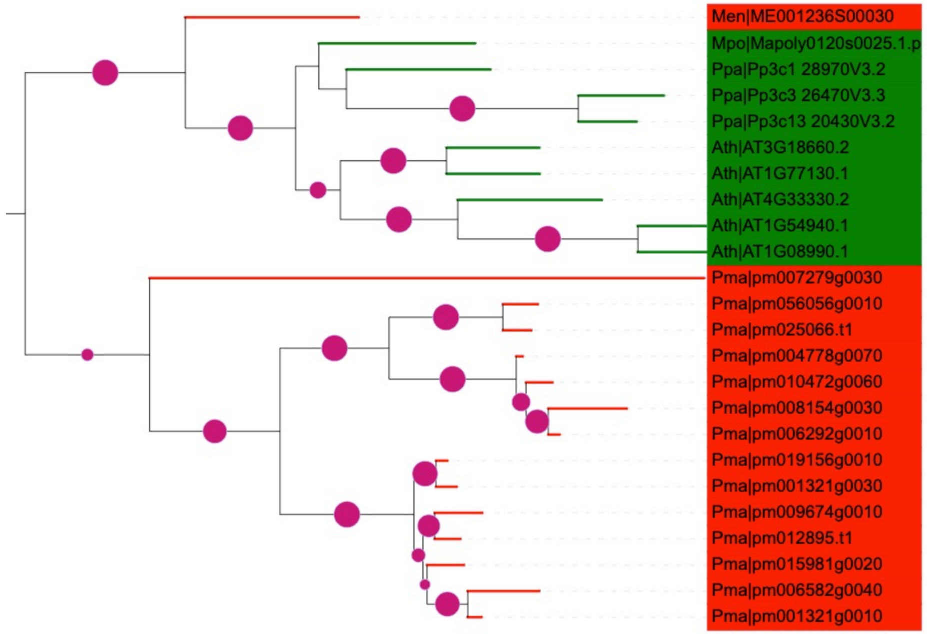

Data S-39a

### GT61 + nr AtXAT blastp

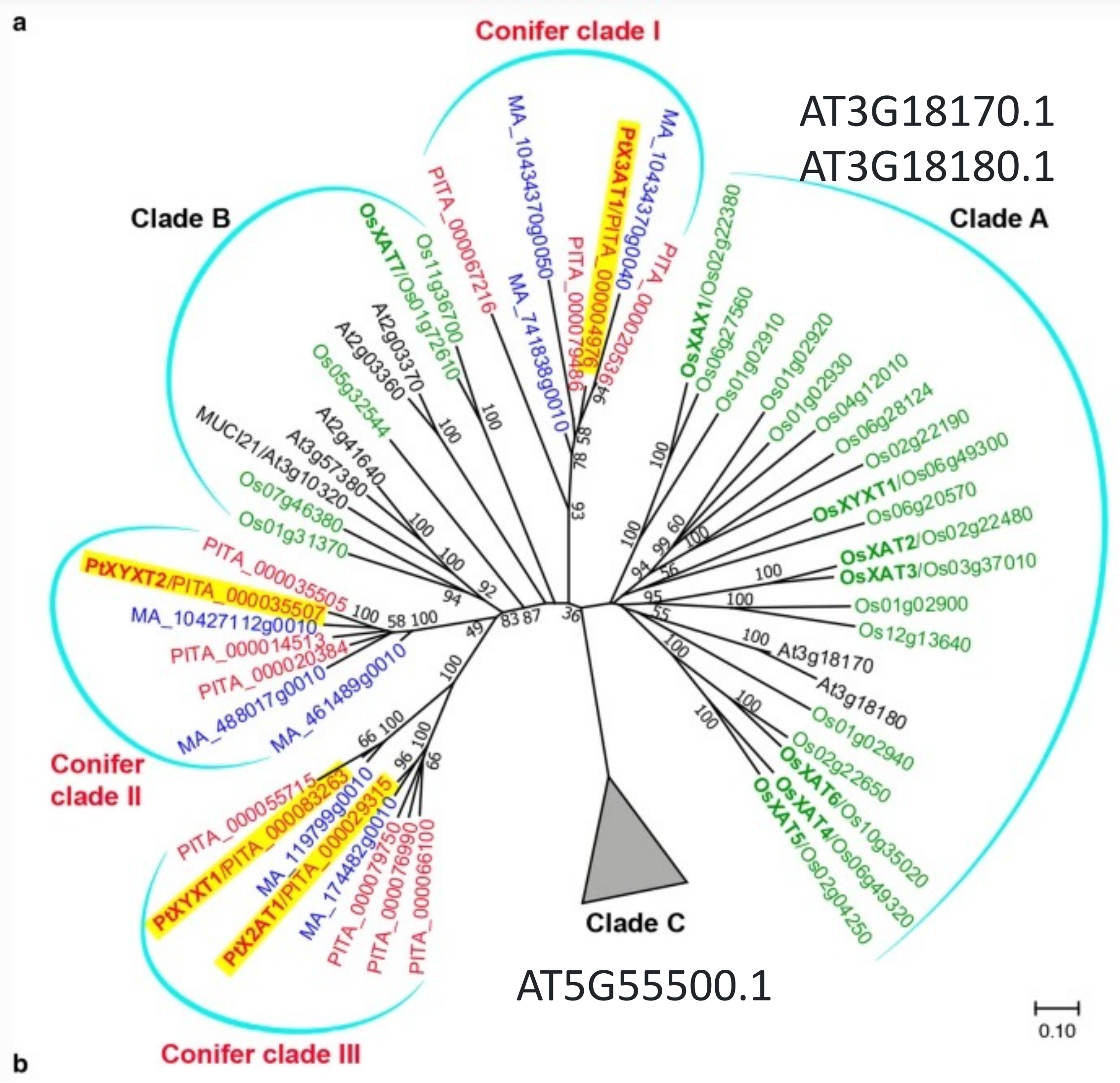


AT3G18170.1
AT3G18180.1

AT5G55500.1

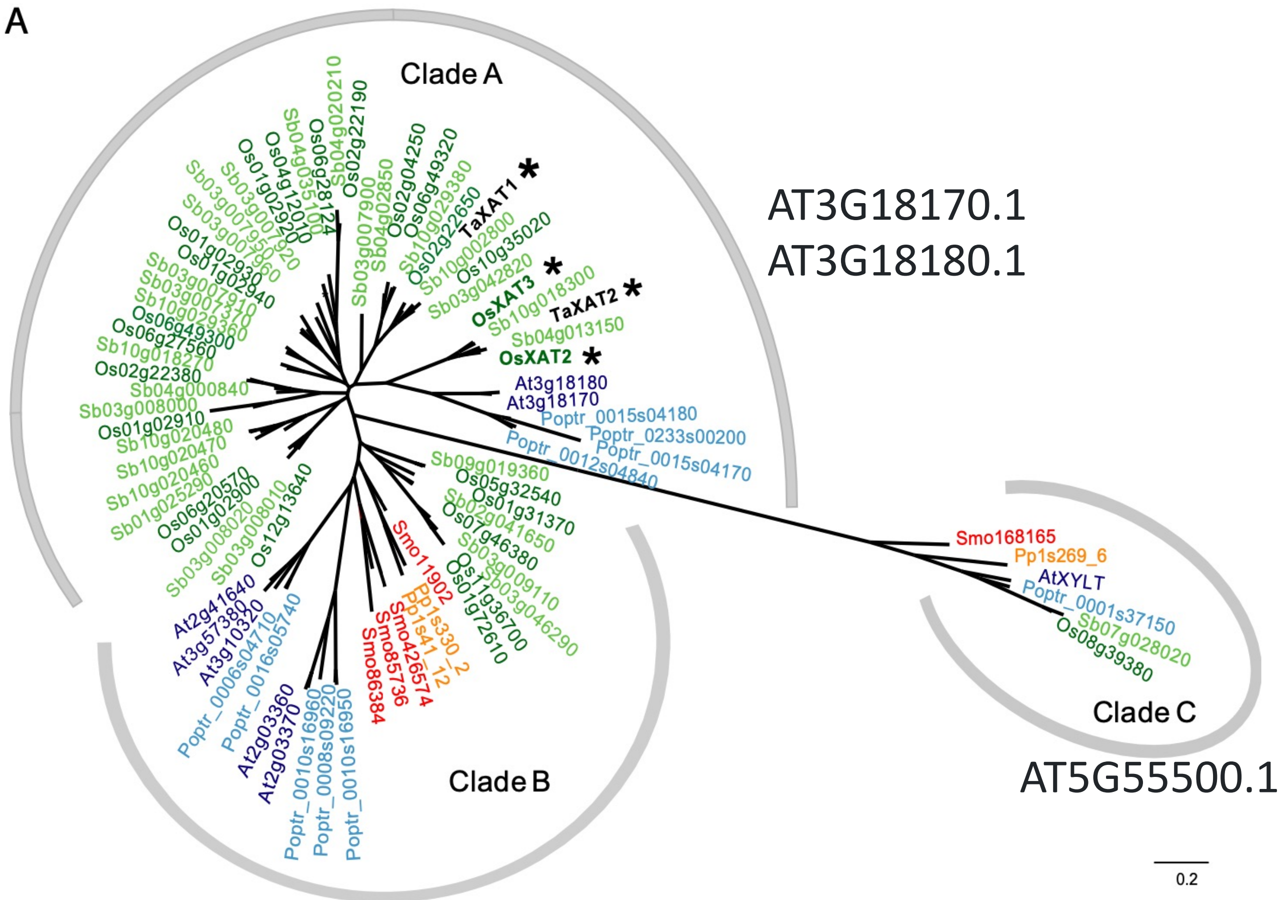


AT3G18170.1
AT3G18180.1

AT5G55500.1

**References**

https://www.ncbi.nlm.nih.gov/pmc/articles/PMC3271882/
https://link.springer.com/article/10.1007/s00425-022-03989-x
https://www.ncbi.nlm.nih.gov/pmc/articles/PMC3479505/

### Data S-39b

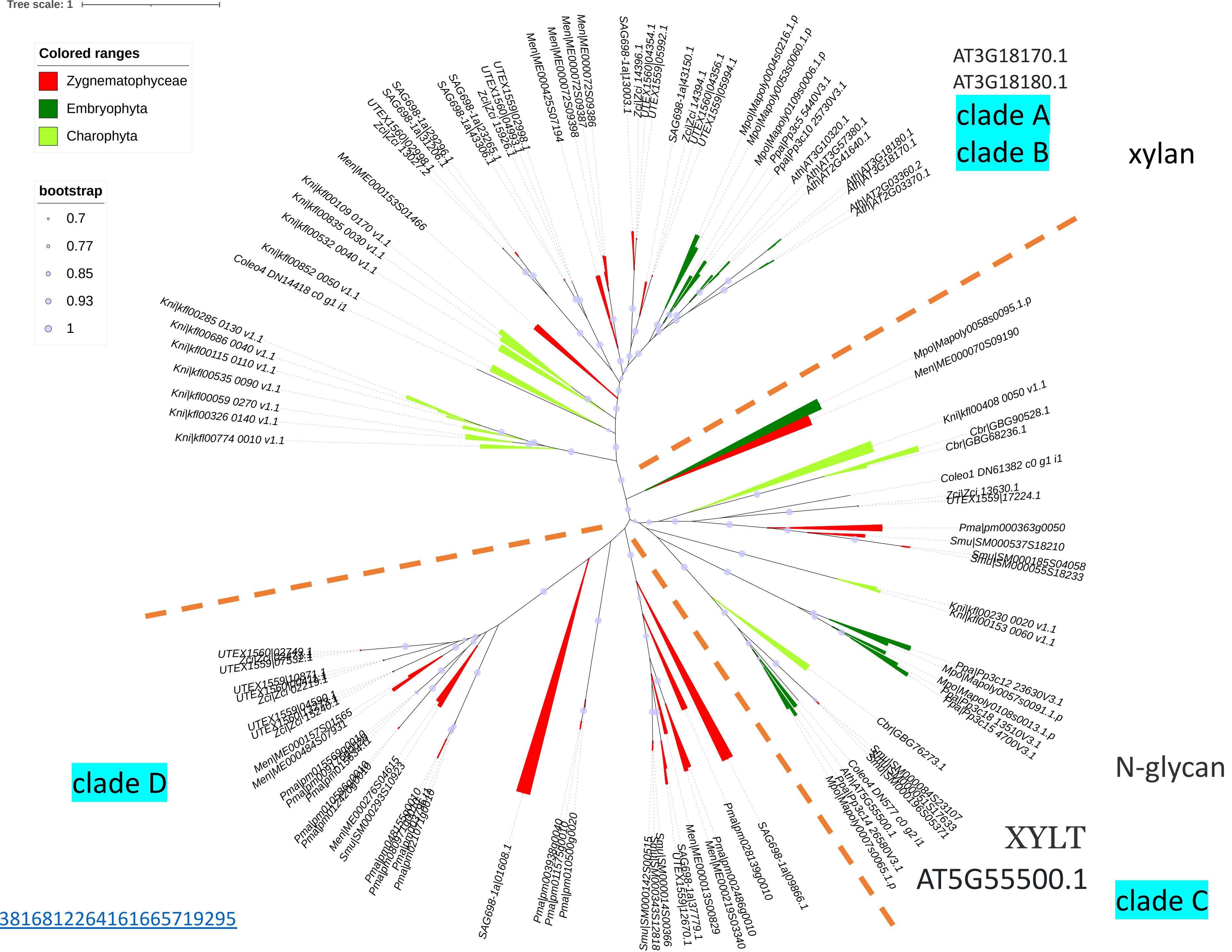


https://itol.embl.de/tree/13423816812264161665719295

Data S-39c

https://itol.embl.de/tree/13423816812471401665758298

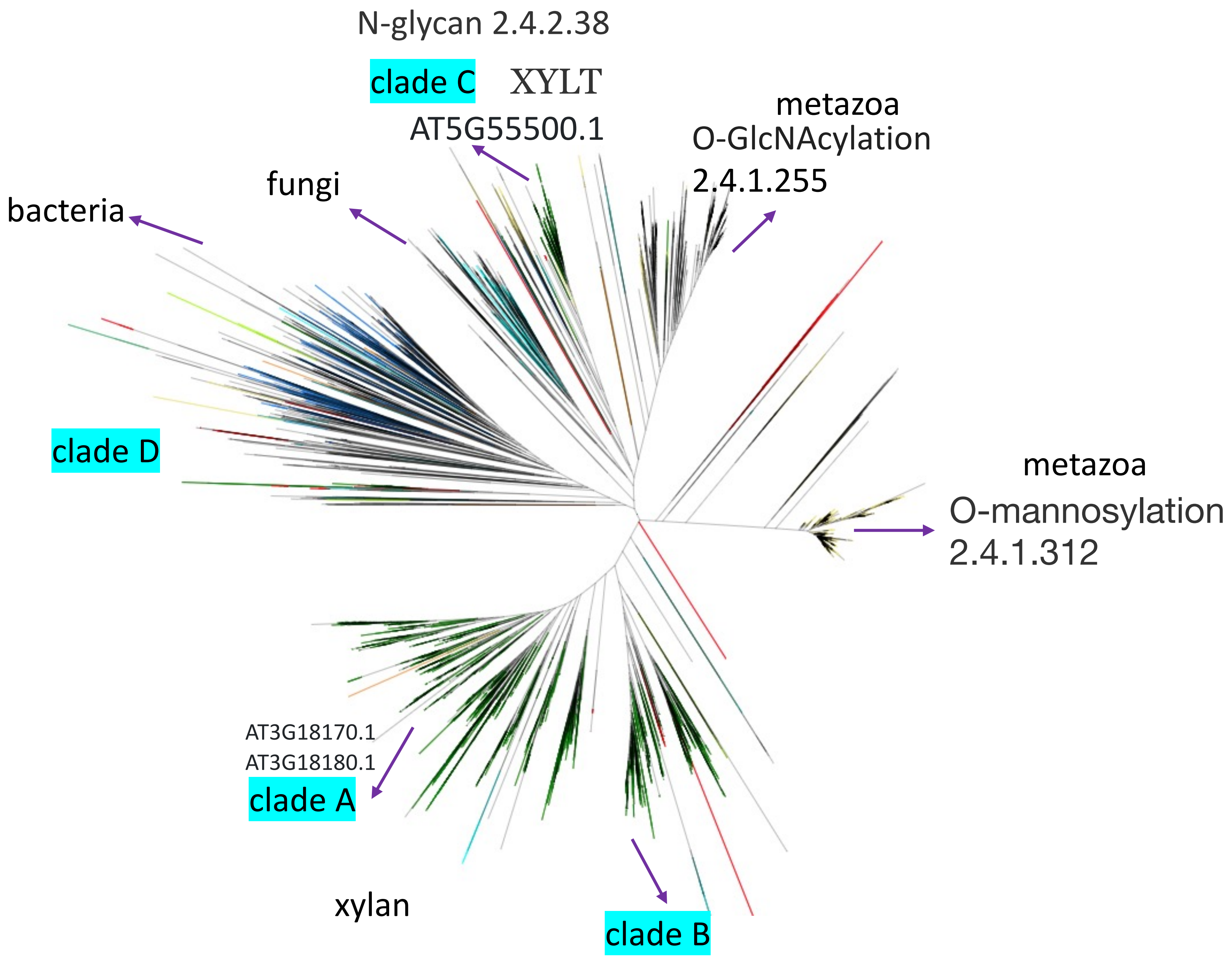

Data S-40

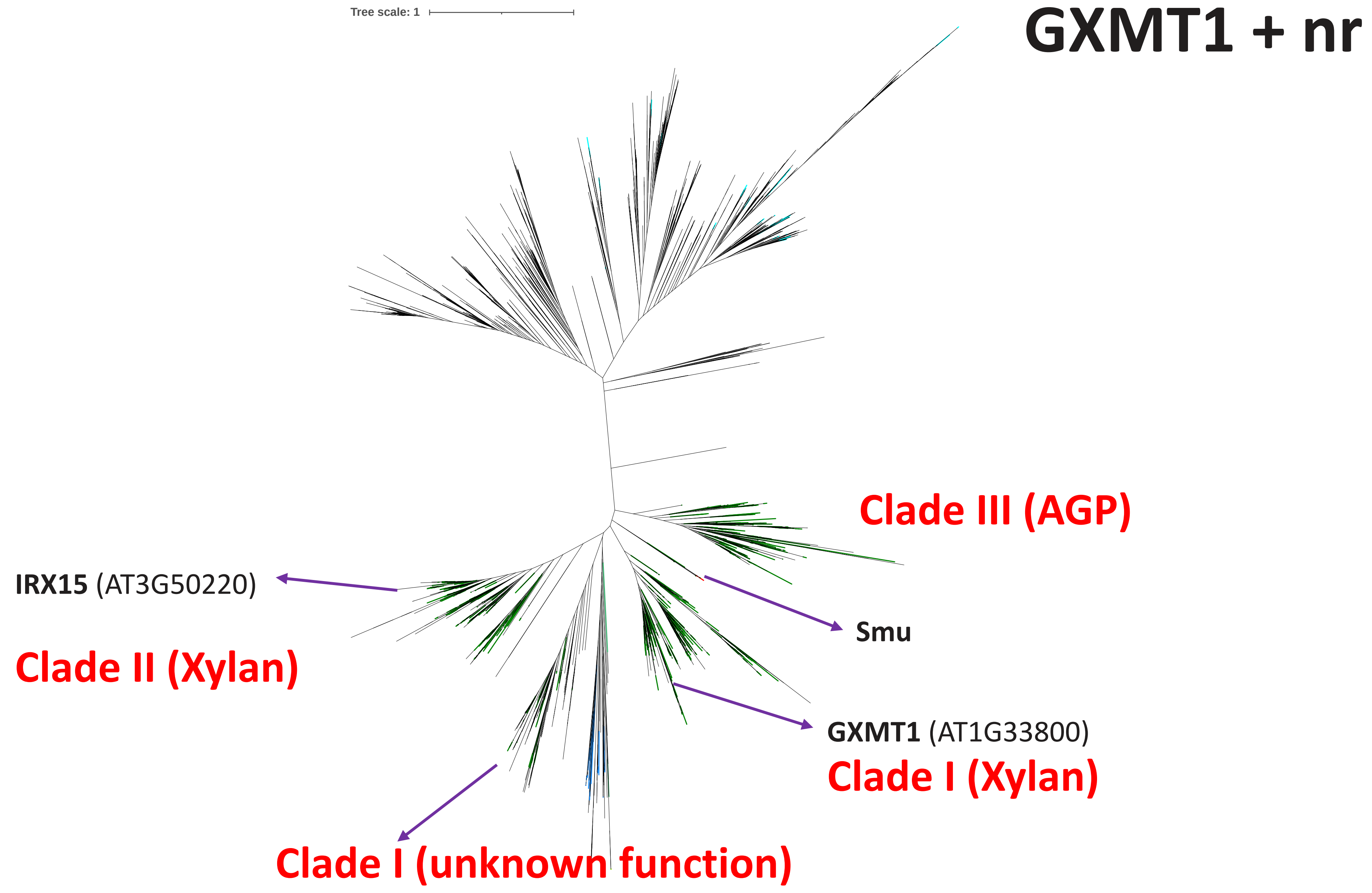
Tree scale: 1
GXMT1 + nr
Clade III (AGP)
IRX15 (AT3G50220)
Smu
Clade II (Xylan)
GXMT1 (AT1G33800)
Clade I (Xylan)
Clade I (unknown function)

Data S-41

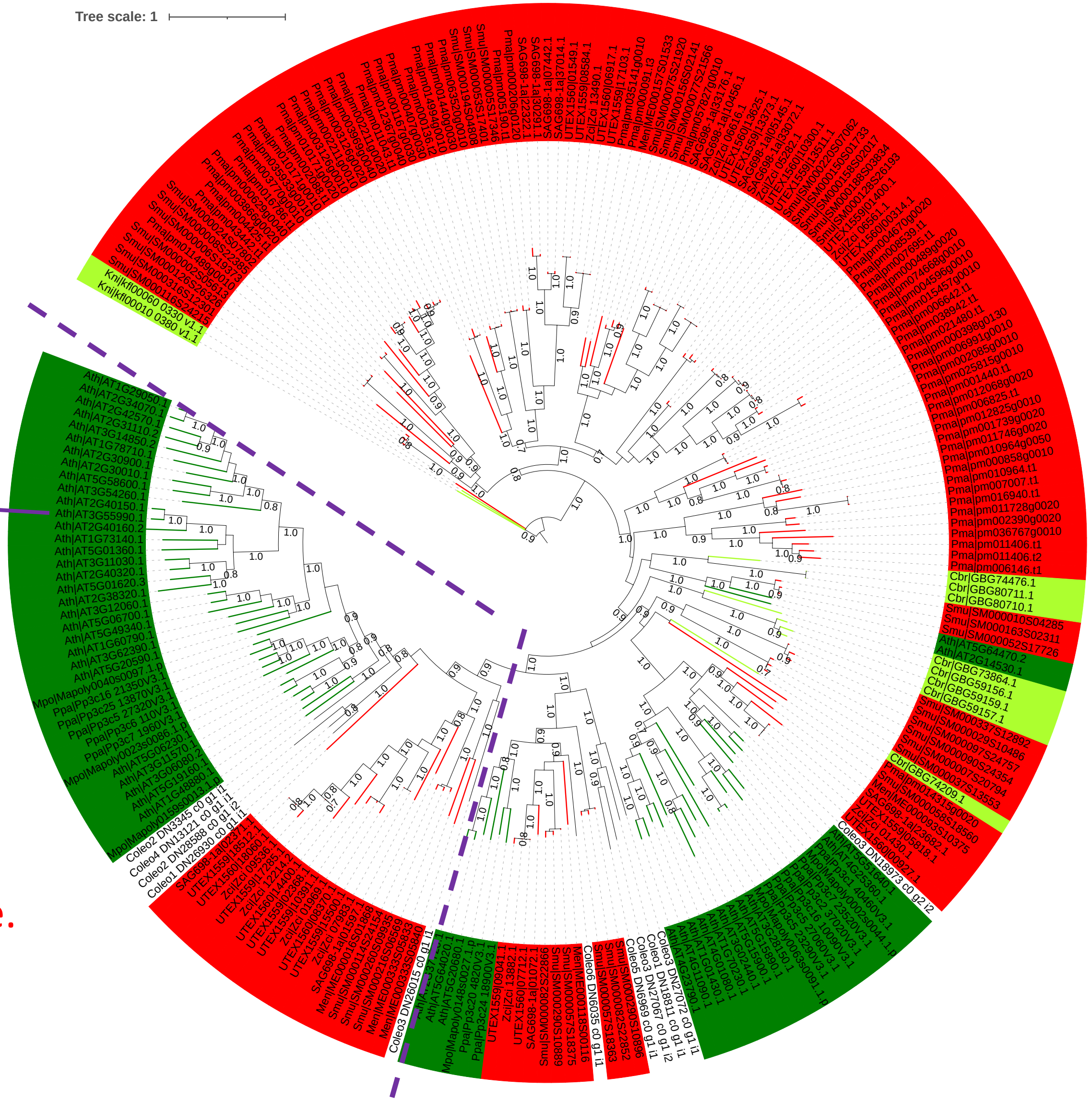


**ESK1 (AT3G55990)**

Zygnema contain the
Xylan O-acetyltransferase.

**References**

https://pubmed.ncbi.nlm.nih.gov/23659919/

https://pubmed.ncbi.nlm.nih.gov/25141999/

Data S-42

### BS1 + DARX1

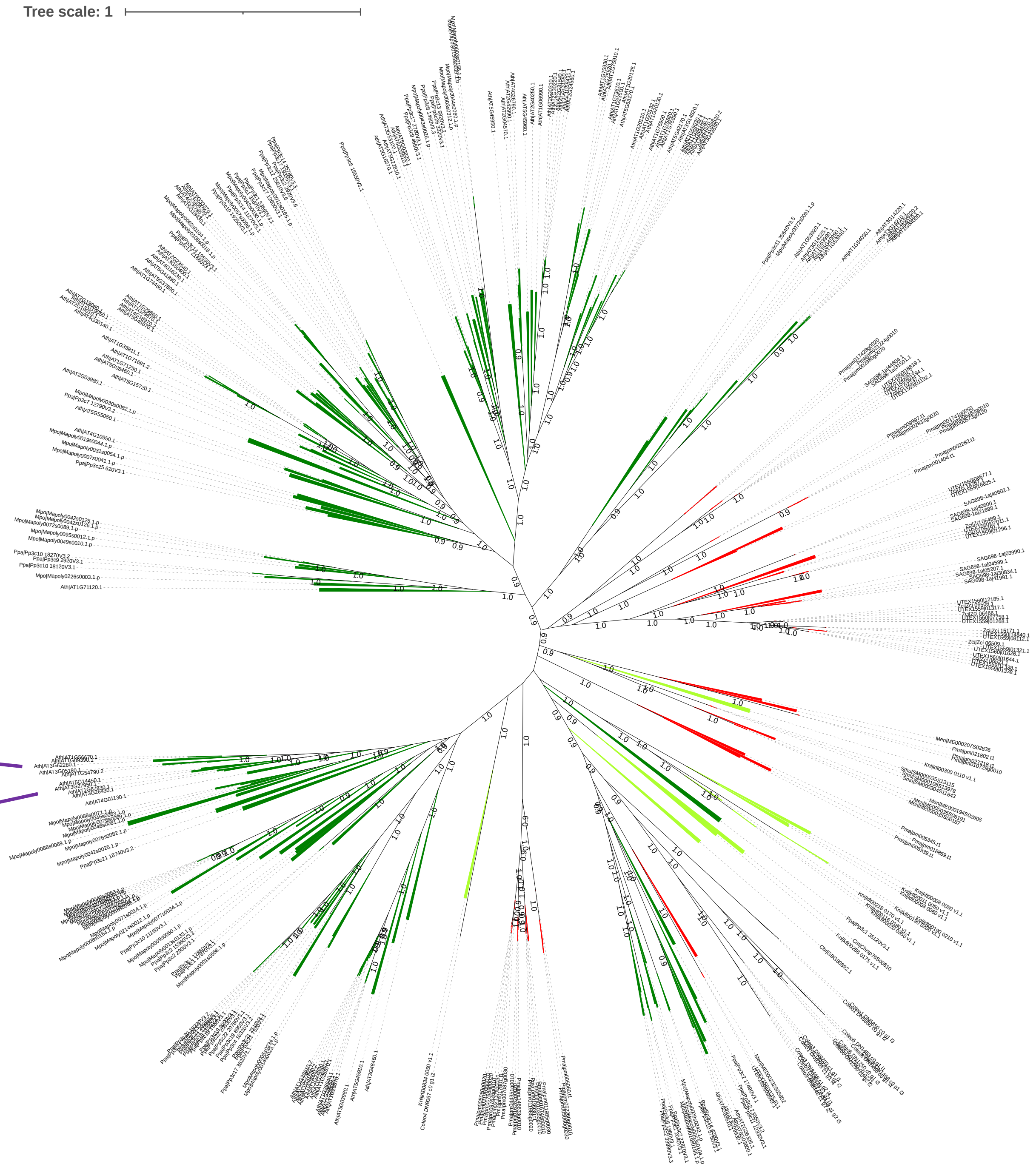

Data S-43

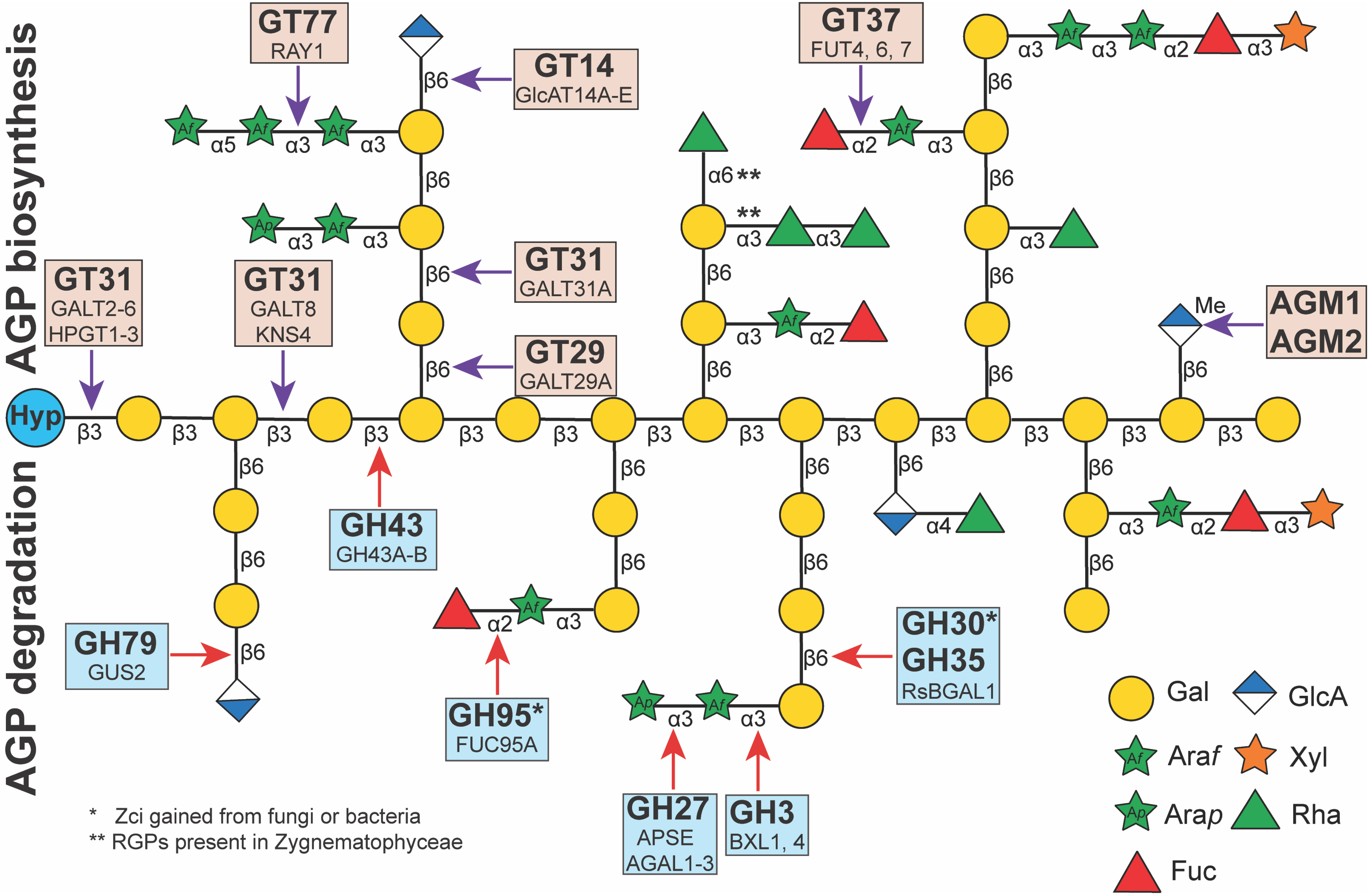
AGP biosynthesis
AGP degradation
GT77
RAY1
GT14
GlcAT14A-E
GT31
GALT31A
GT29
GALT29A
GT31
GALT2-6
HPGT1-3
GT31
GALT8
KNS4
GT37
FUT4, 6, 7
AGM1
AGM2
Me
Hyp
GH43
GH43A-B
GH79
GUS2
GH95*
FUC95A
GH27
APSE
AGAL1-3
GH3
BXL1, 4
GH30*
GH35
RsBGAL1
α6 **
**
Gal
GlcA
Araf
Xyl
Arap
Rha
Fuc
* Zci gained from fungi or bacteria
** RGPs present in Zygnematophyceae

### AGP

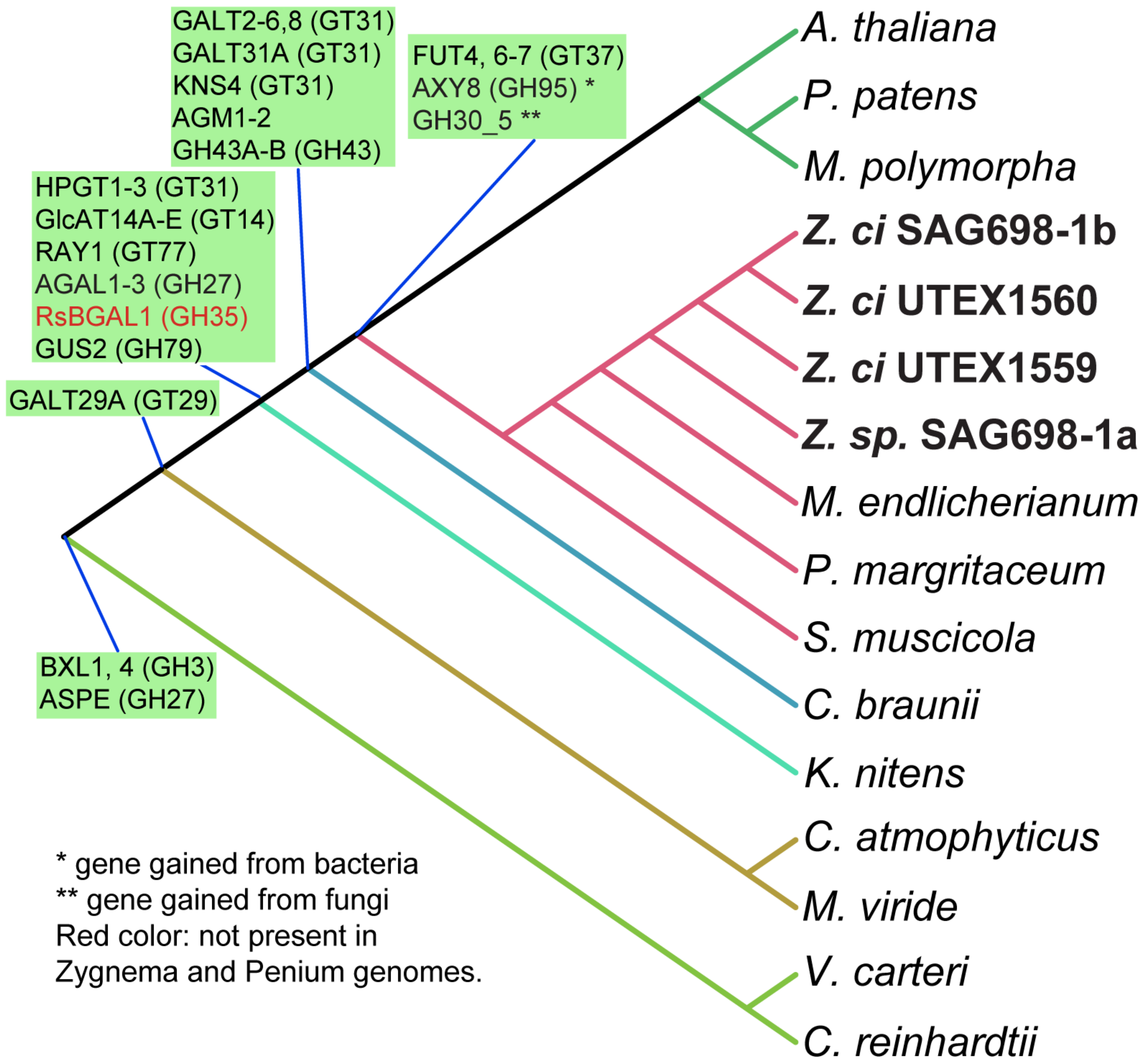
GALT2-6,8 (GT31)
GALT31A (GT31)
KNS4 (GT31)
AGM1-2
GH43A-B (GH43)
FUT4, 6-7 (GT37)
AXY8 (GH95) *
GH30_5 **
HPGT1-3 (GT31)
GlcAT14A-E (GT14)
RAY1 (GT77)
AGAL1-3 (GH27)
RsBGAL1 (GH35)
GUS2 (GH79)
GALT29A (GT29)
BXL1, 4 (GH3)
ASPE (GH27)
A. thaliana
P. patens
M. polymorpha
Z. ci SAG698-1b
Z. ci UTEX1560
Z. ci UTEX1559
Z. sp. SAG698-1a
M. endlicherianum
P. margritaceum
S. muscicola
C. braunii
K. nitens
C. atmophyticus
M. viride
V. carteri
C. reinhardtii
* gene gained from bacteria
** gene gained from fungi
Red color: not present in
Zygnema and Penium genomes.

Data S-44

# GT31

https://itol.embl.de/tree/2081277026466951644607964

Narciso JO, Zeng W, Ford K, Lampugnani ER, Humphries J, Austarheim I, van de Meene A, Bacic A, Doblin MS. Biochemical and Functional Characterization of GALT8, an Arabidopsis GT31 β-(1,3)-Galactosyltransferase That Influences Seedling Development. Front Plant Sci. 2021 May 25;12:678564. doi: 10.3389/fpls.2021.678564.

At1g22015 (*AtGALT8*)

At1g33430 (KNS4/UPEX1)

At1G77810 (β-1,3-GalT)

At1G32930 (*AtGALT31A*)

β-1,6-GalT activity elongating β-1,6-galactan side chains of AGP glycans. Functional AtGALT31A is essential for normal plant embryogenesis.

Geshi, N., Johansen, J. N., Dilokpimol, A., Rolland, A., Belcram, K., Verger, S., et al. (2013). A galactosyltransferase acting on arabinogalactan protein glycans is essential for embryo development in Arabidopsis. Plant J. 76, 128–137. doi: 10.1111/tpj.12281

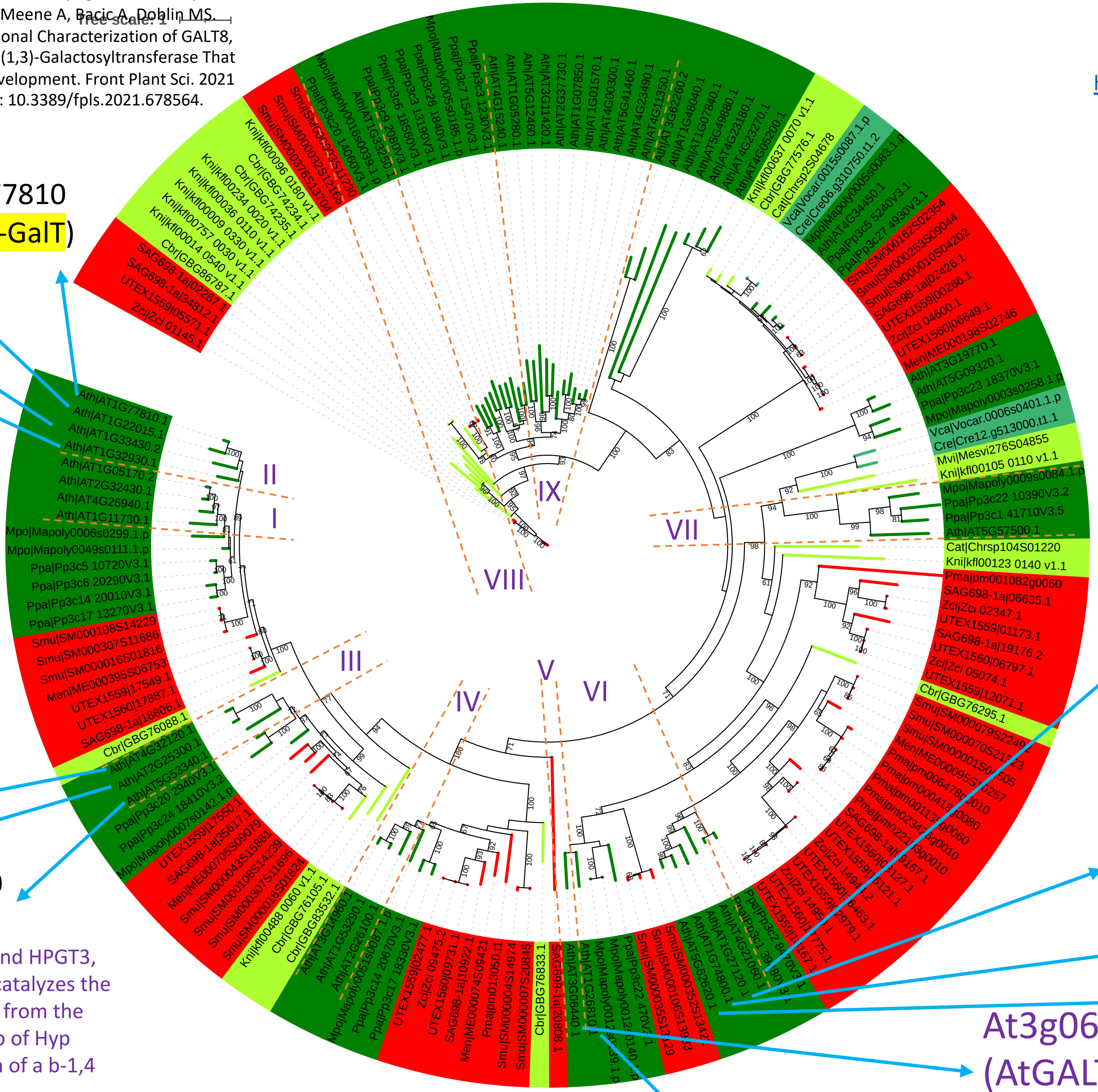


Basu, D., Liang, Y., Liu, X., Himmeldirk, K., Faik, A., Kieliszewski, M., et al. (2013). Functional identification of a hydroxyproline-o-galactosyltransferase specific for arabinogalactan protein biosynthesis in Arabidopsis. J. Biol. Chem. 288, 10132–10143. doi: 10.1074/jbc.M112.432609

Basu D, Wang W, Ma S, DeBrosse T, Poirier E, Emch K, Soukup E, Tian L, Showalter AM. Two Hydroxyproline Galactosyltransferases, GALT5 and GALT2, Function in Arabinogalactan-Protein Glycosylation, Growth and Development in Arabidopsis. PLoS One. 2015 May 14;10(5):e0125624. doi: 10.1371/journal.pone.0125624.

Basu D, Tian L, Wang W, Bobbs S, Herock H, Travers A, Showalter AM. A small multigene hydroxyproline-O-galactosyltransferase family functions in arabinogalactan-protein glycosylation, growth and development in Arabidopsis. BMC Plant Biol. 2015 Dec 21;15:295. doi: 10.1186/s12870-015-0670-7.

At4g21060 (AtGALT2)

transfer of Gal to hydroxyproline residues

At1g27120 (AtGALT4)

At1g74800 (AtGALT5)

At5g62620(AtGALT6)

At3g06440 (AtGALT3)

At1g26810 (AtGALT1)

At4G32120 (HPGT2)

At2G25300 (HPGT3)

At5G53340 (HPGT1)

Hyp O-galactosyltransferases, HPGT1, HPGT2 and HPGT3, from Arabidopsis microsomal fractions. HPGT catalyzes the transfer of a D-galactopyranosyl (Galp) residue from the sugar donor UDP-a-D-Gal to the hydroxyl group of Hyp residues of peptides, leading to the forma- tion of a b-1,4 linkage. They lack a galectin domain.

Ogawa-Ohnishi M, Matsubayashi Y. Identification of three potent hydroxyproline O-galactosyltransferases in Arabidopsis. Plant J. 2015 Mar;81(5):736-46. doi: 10.1111/tpj.12764.

GALT1 is both sufficient and essential for the addition of b1,3-linked galactose residues to N-glycans and thus is required for the biosynthesis of Lewis a structures in Arabidopsis.

Strasser R, Bondili JS, Vavra U, Schoberer J, Svoboda B, Glössl J, Léonard R, Stadlmann J, Altmann F, Steinkellner H, Mach L. A unique beta1,3-galactosyltransferase is indispensable for the biosynthesis of N-glycans containing Lewis a structures in Arabidopsis thaliana. Plant Cell. 2007 Jul;19(7):2278-92. doi: 10.1105/tpc.107.052985.

Data S-45

https://itol.embl.de/tree/208127240222157721668394053

At4G32120
(HPGT2)

At2G25300
(HPGT3)

At5G53340
(HPGT1)

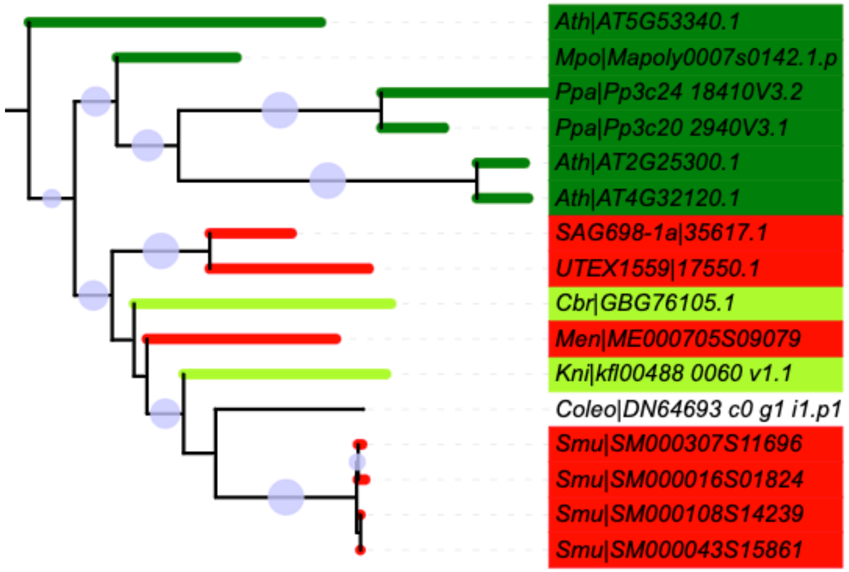


β-1,3-galactan (backbone)
initiate

GALT2-6 At4g21060

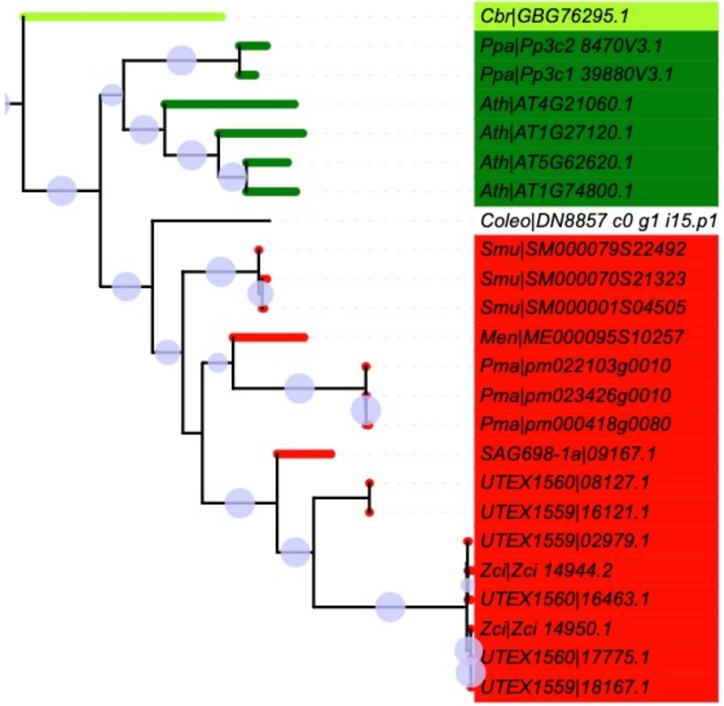


β-1,3-galactan
initiate

At1g33430
(KNS4/UPEX1)

At1g22015
(*AtGALT8*)

At1G32930
(*AtGALT31A*)

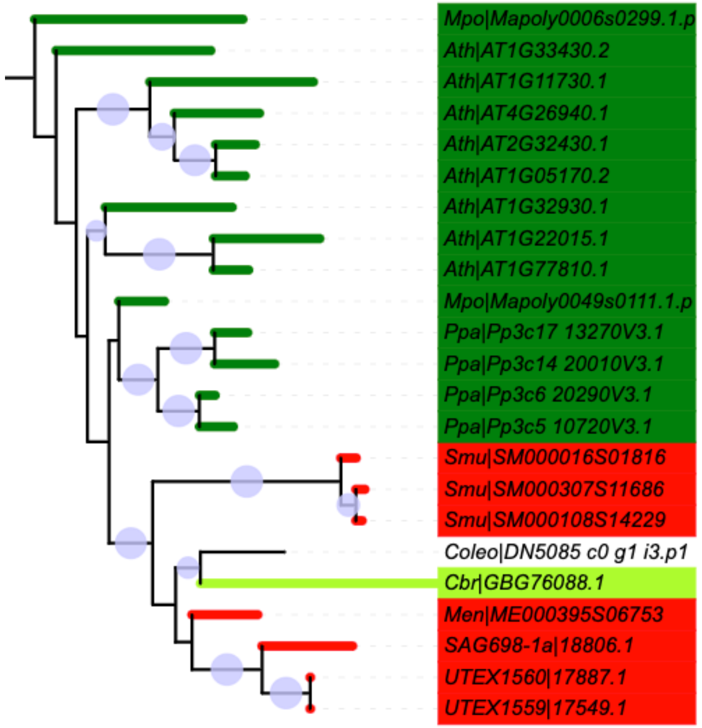


β-1,3-galactan
β-1,6-galactan (sidechain)
elongate

Data S-46

### GT29 + Coleochaete

https://itol.embl.de/tree/2081278323533483166511609

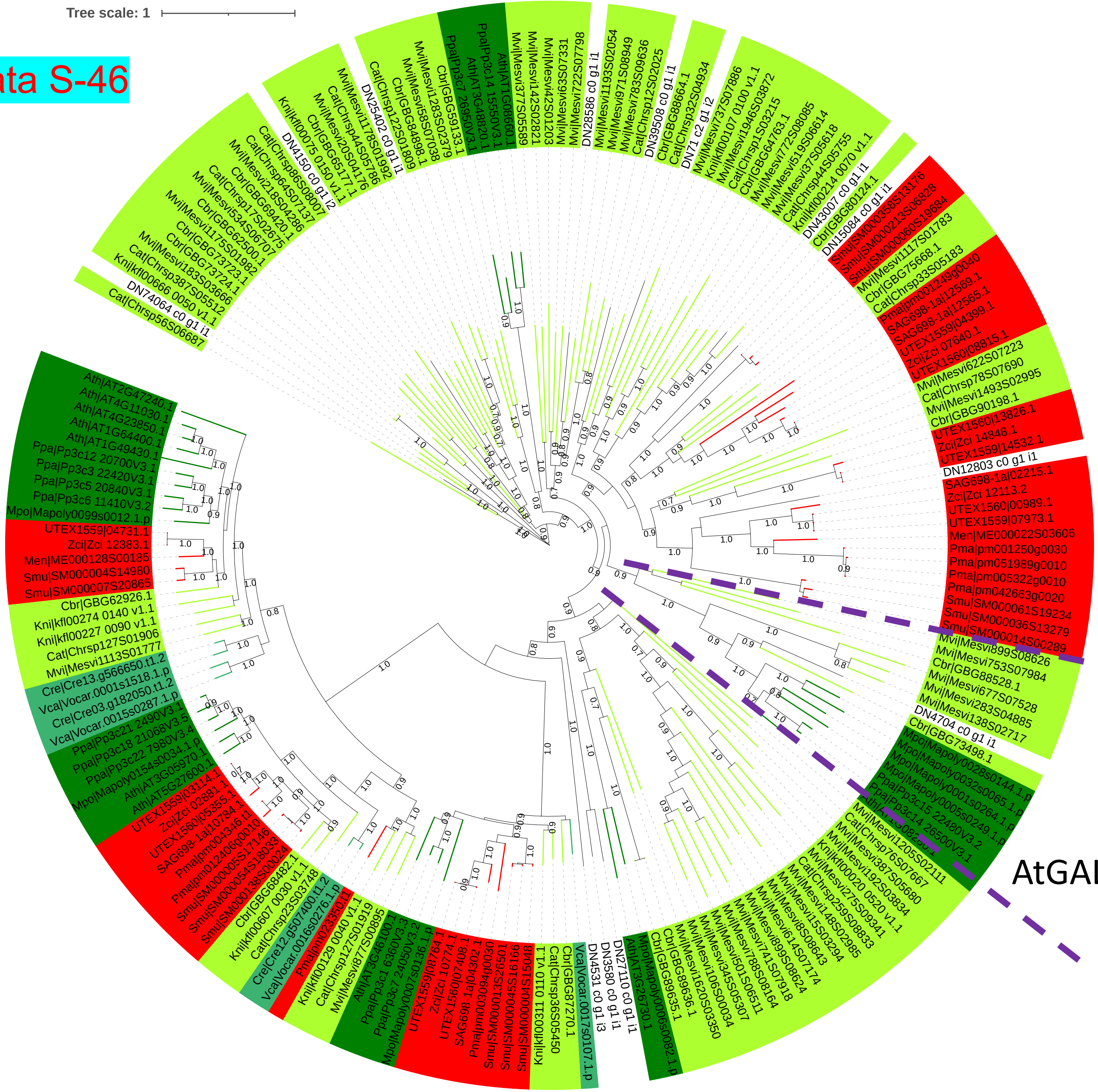


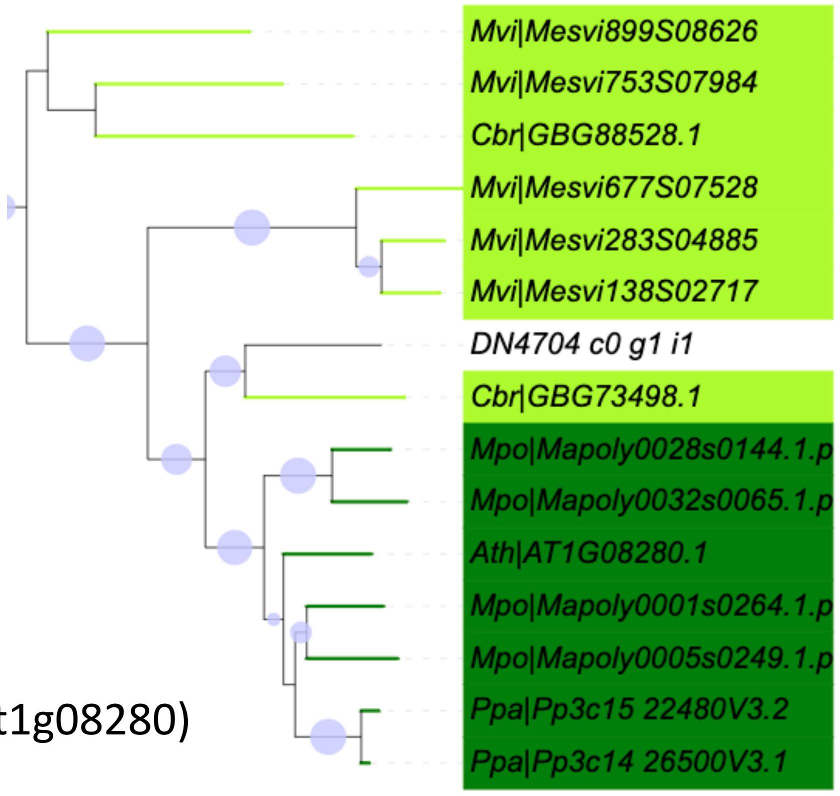

Data S-47

https://itol.embl.de/tree/134238164171236071665177614

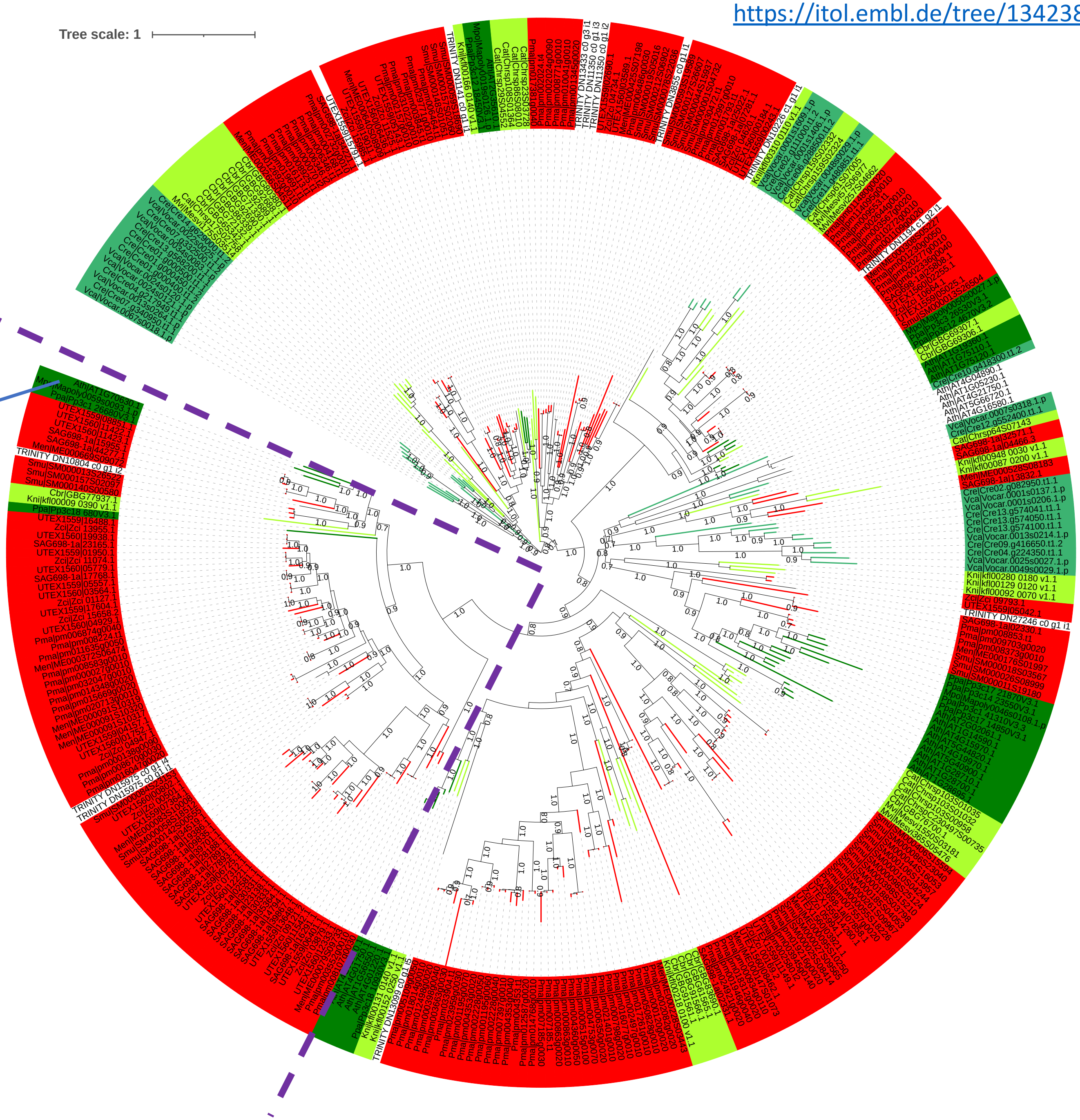


**AGP**

At1g70630 (named REDUCED ARABINOSE YARIV1, **RAY1**)

a putative arabinofuranosyltransferase since the mutation caused a reduced level of arabinofuranose (Araf ) in its AGPs.

Gille, S., Sharma, V., Baidoo, E. E. K., Keasling, J. D., Scheller, H. V., and Pauly, M. (2013). Arabinosylation of a yariv-precipitable cell wall polymer impacts plant growth as exemplified by the arabidopsis glycosyltransferase mutant ray1. Mol. Plant 6, 1369–1372. doi: 10.1093/mp/sst029

### GT77 + Coleochaete

Data S-48

# GT14

AtGlcAT14D (AT3G24040)

AtGlcAT14C (At2g37585)

AtGlcAT14E (AT3G15350)

AtGlcAT14A (At5g39990)

AtGlcAT14B (At5g15050)

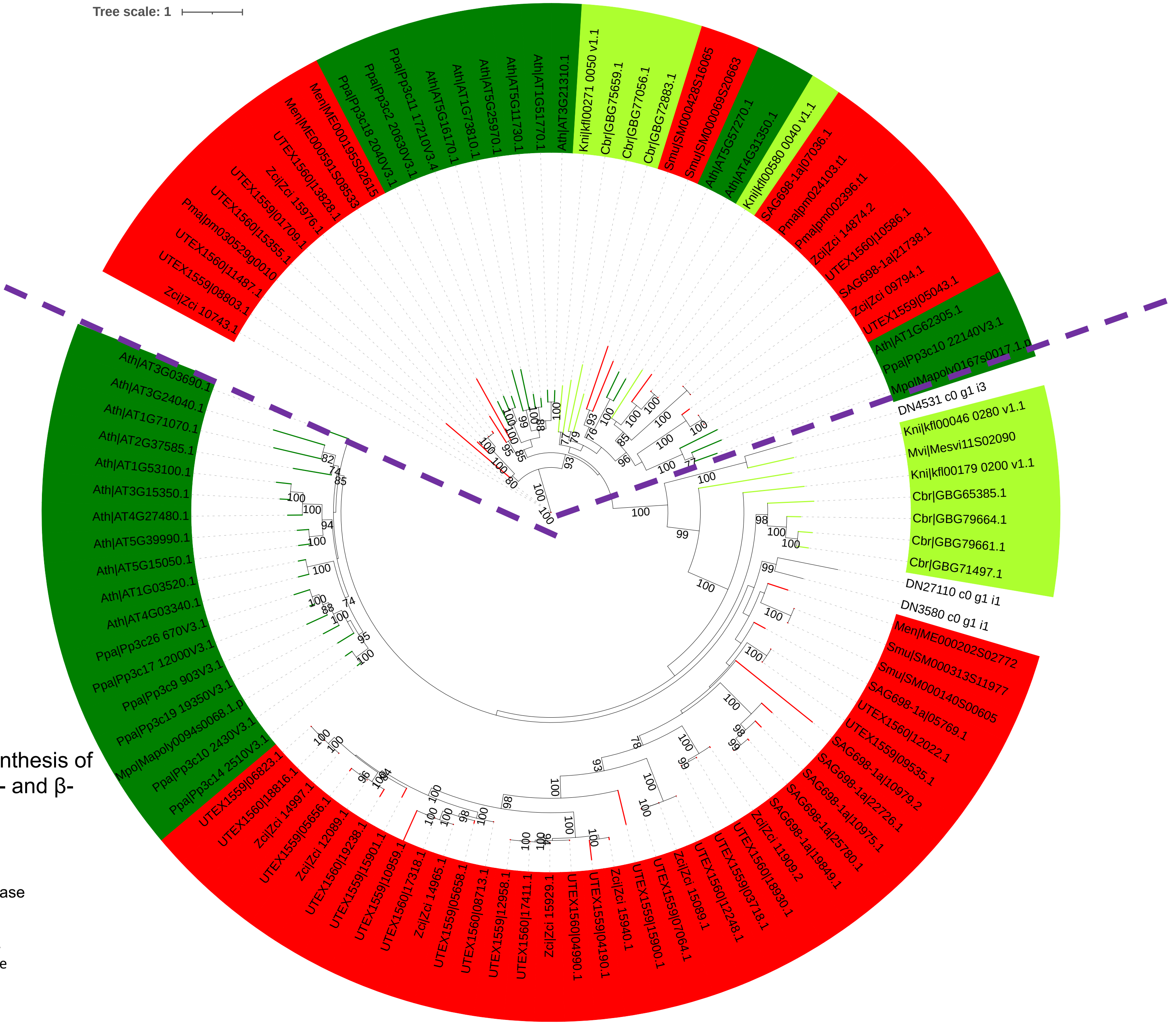


identified as a glucuronosyltransferase involved in the biosynthesis of AGP glycans. β-GlcAT activity by adding GlcA to both β-1,6- and β-1,3-galactan.

Dilokpimol, A., and Geshi, N. (2014). Arabidopsis thaliana glucuronosyltransferase in family GT14. Plant Signal. Behav. 9:e28891. doi: 10.4161/psb.28891

Knoch, E., Dilokpimol, A., Tryfona, T., Poulsen, C. P., Xiong, G., Harholt, J., et al. (2013). A β-glucuronosyltransferase from Arabidopsis thaliana involved in biosynthesis of type II arabinogalactan has a role in cell elongation during seedling growth. Plant J. 76, 1016–1029. doi: 10.1111/tpj.12353

Data S-49

# GT37

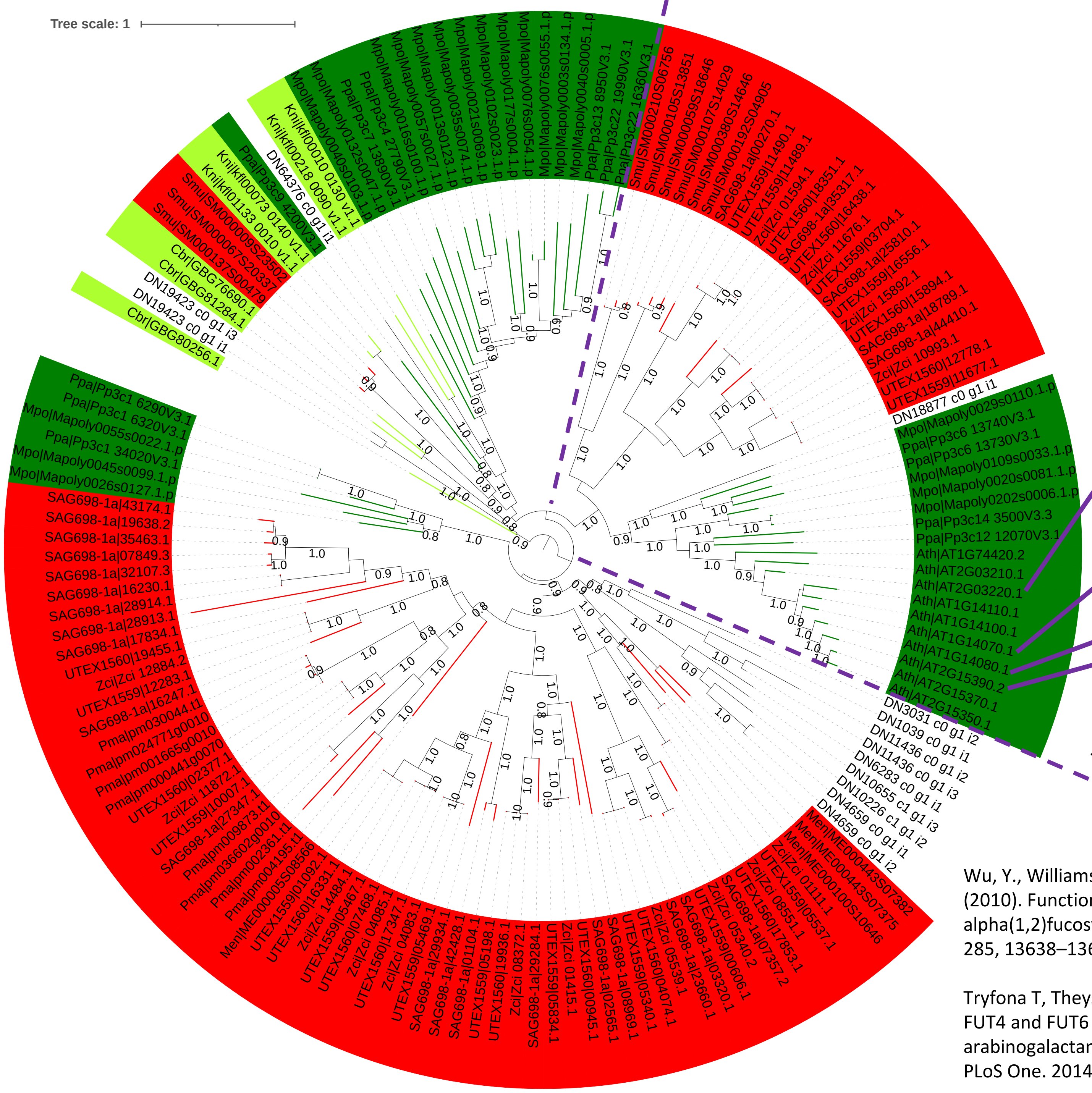


Vanzin GF, Madson M, Carpita NC, Raikhel NV, Keegstra K, Reiter WD. The mur2 mutant of Arabidopsis thaliana lacks fucosylated xyloglucan because of a lesion in fucosyltransferase AtFUT1. Proc Natl Acad Sci U S A. 2002 Mar 5;99(5):3340-5. doi: 10.1073/pnas.052450699.

AtFUT1 (*At2g03220*)

AtFUT7 (At1g14070)

AtFUT6 (*At1g14080*)

AtFUT4 (*At2g15390*)

Two Arabidopsis GTs from family GT37 were identified as α-1,2-fucosyltransferases involved in the biosynthesis of AGP glycans

Wu, Y., Williams, M., Bernard, S., Driouich, A., Showalter, A. M., and Faik, A. (2010). Functional identification of two nonredundant Arabidopsis alpha(1,2)fucosyltransferases specific to arabinogalactan proteins. J. Biol. Chem. 285, 13638–13645. doi: 10.1074/jbc.M110.102715

Tryfona T, Theys TE, Wagner T, Stott K, Keegstra K, Dupree P. Characterisation of FUT4 and FUT6 α-(1 → 2)-fucosyltransferases reveals that absence of root arabinogalactan fucosylation increases Arabidopsis root growth salt sensitivity. PLoS One. 2014 Mar 25;9(3):e93291. doi: 10.1371/journal.pone.0093291.

Data S-50

hmmsearch
PF04669

| | | | | | | | | | |
|---|---|---|---|---|---|---|---|---|---|
| Polysacc_synt_4 | 190 | Ath\|AT1G09610.1 | 282 | 3.8e-70 | 1 | 190 | 79 | 263 | 0.994736842105263 |
| Polysacc_synt_4 | 190 | Ath\|AT1G27930.1 | 289 | 1.8e-74 | 1 | 190 | 83 | 270 | 0.994736842105263 |
| Polysacc_synt_4 | 190 | Ath\|AT1G33800.1 | 297 | 1.5e-71 | 2 | 190 | 93 | 275 | 0.989473684210526 |
| Polysacc_synt_4 | 190 | Ath\|AT1G67330.1 | 291 | 2.5e-75 | 1 | 190 | 86 | 274 | 0.994736842105263 |
| Polysacc_synt_4 | 190 | Ath\|AT1G71690.1 | 295 | 2.4e-77 | 2 | 190 | 92 | 278 | 0.989473684210526 |
| Polysacc_synt_4 | 190 | Ath\|AT2G15440.1 | 329 | 3e-71 | 2 | 190 | 99 | 287 | 0.989473684210526 |
| Polysacc_synt_4 | 190 | Ath\|AT3G50220.1 | 322 | 8.9e-74 | 2 | 189 | 109 | 296 | 0.984210526315789 |
| Polysacc_synt_4 | 190 | Ath\|AT4G09990.1 | 290 | 7.5e-70 | 2 | 190 | 85 | 268 | 0.989473684210526 |
| Polysacc_synt_4 | 190 | Ath\|AT4G24910.1 | 315 | 7.2e-65 | 2 | 190 | 105 | 295 | 0.989473684210526 |
| Polysacc_synt_4 | 190 | Ath\|AT5G67210.1 | 317 | 1.2e-73 | 2 | 189 | 102 | 289 | 0.984210526315789 |
| Polysacc_synt_4 | 190 | Cbr\|GBG71883.1 | 344 | 2.2e-16 | 17 | 176 | 165 | 316 | 0.836842105263158 |
| Polysacc_synt_4 | 190 | Cbr\|GBG71887.1 | 339 | 9.2e-14 | 17 | 175 | 152 | 302 | 0.831578947368421 |
| Polysacc_synt_4 | 190 | Smu\|SM000088S23775 | 194 | 8.5e-21 | 2 | 106 | 84 | 193 | 0.547368421052632 |
| Polysacc_synt_4 | 190 | Smu\|SM000247S08301 | 194 | 5.5e-21 | 2 | 106 | 84 | 193 | 0.547368421052632 |
| Polysacc_synt_4 | 190 | Smu\|SM000302S11676 | 194 | 3.4e-21 | 2 | 106 | 84 | 193 | 0.547368421052632 |

blastp

AGM1 & AGM2

| | | | | | | | | | | | |
|---|---|---|---|---|---|---|---|---|---|---|---|
| Ath\|AT2G15440.1 | AGM2 | 36.604 | 265 | 150 | 8 | 51 | 303 | 33 | 291 | 5.73e-55 | 169 |
| Ath\|AT2G15440.1 | AGM1 | 38.034 | 234 | 133 | 7 | 76 | 303 | 62 | 289 | 1.30e-54 | 168 |
| Ath\|AT1G67330.1 | AGM2 | 100.000 | 291 | 0 | 0 | 1 | 291 | 1 | 291 | 0.0 | 611 |
| Ath\|AT1G67330.1 | AGM1 | 68.905 | 283 | 81 | 5 | 15 | 291 | 8 | 289 | 2.32e-142 | 390 |
| Ath\|AT1G71690.1 | AGM1 | 51.174 | 213 | 102 | 2 | 70 | 281 | 62 | 273 | 2.63e-80 | 233 |
| Ath\|AT1G71690.1 | AGM2 | 45.385 | 260 | 136 | 5 | 38 | 294 | 35 | 291 | 1.42e-78 | 229 |
| Ath\|AT1G33800.1 | AGM2 | 49.784 | 231 | 107 | 4 | 67 | 289 | 62 | 291 | 9.91e-81 | 234 |
| Ath\|AT1G33800.1 | AGM1 | 52.511 | 219 | 99 | 4 | 70 | 284 | 62 | 279 | 5.86e-78 | 227 |
| Ath\|AT1G09610.1 | AGM2 | 51.786 | 224 | 103 | 5 | 62 | 281 | 69 | 291 | 7.58e-81 | 234 |
| Ath\|AT1G09610.1 | AGM1 | 52.133 | 211 | 92 | 3 | 62 | 266 | 66 | 273 | 5.14e-80 | 232 |
| Ath\|AT4G24910.1 | AGM2 | 29.831 | 295 | 187 | 5 | 26 | 307 | 4 | 291 | 1.80e-42 | 136 |
| Ath\|AT4G24910.1 | AGM1 | 35.238 | 210 | 125 | 5 | 105 | 307 | 84 | 289 | 1.04e-40 | 132 |
| Ath\|AT3G50220.1 | AGM2 | 34.768 | 302 | 165 | 12 | 21 | 310 | 10 | 291 | 6.03e-48 | 151 |
| Ath\|AT3G50220.1 | AGM1 | 37.229 | 231 | 130 | 7 | 89 | 310 | 65 | 289 | 7.55e-48 | 150 |
| Ath\|AT1G27930.1 | AGM1 | 100.000 | 289 | 0 | 0 | 1 | 289 | 1 | 289 | 0.0 | 600 |
| Ath\|AT1G27930.1 | AGM2 | 69.258 | 283 | 80 | 5 | 8 | 289 | 15 | 291 | 1.44e-145 | 399 |
| Ath\|AT4G09990.1 | AGM2 | 44.706 | 255 | 129 | 2 | 22 | 272 | 32 | 278 | 3.71e-79 | 230 |
| Ath\|AT4G09990.1 | AGM1 | 51.887 | 212 | 99 | 3 | 68 | 276 | 67 | 278 | 1.19e-75 | 221 |
| Ath\|AT5G67210.1 | AGM2 | 34.932 | 292 | 172 | 9 | 20 | 303 | 10 | 291 | 8.08e-52 | 161 |
| Ath\|AT5G67210.1 | AGM1 | 36.199 | 221 | 134 | 5 | 82 | 300 | 65 | 280 | 3.68e-48 | 151 |
| Cbr\|GBG71883.1 | AGM2 | 31.395 | 172 | 82 | 5 | 166 | 315 | 102 | 259 | 4.28e-23 | 85.1 |
| Cbr\|GBG71883.1 | AGM1 | 30.994 | 171 | 83 | 5 | 166 | 315 | 99 | 255 | 6.38e-21 | 79.0 |
| Cbr\|GBG71887.1 | AGM1 | 29.825 | 171 | 85 | 4 | 153 | 302 | 99 | 255 | 4.98e-21 | 79.0 |
| Cbr\|GBG71887.1 | AGM2 | 30.058 | 173 | 83 | 6 | 153 | 302 | 102 | 259 | 3.13e-19 | 73.9 |
| Smu\|SM000088S23775 | AGM2 | 38.830 | 188 | 98 | 5 | 10 | 193 | 19 | 193 | 1.07e-35 | 115 |
| Smu\|SM000088S23775 | AGM1 | 43.846 | 130 | 65 | 3 | 66 | 193 | 66 | 189 | 2.05e-30 | 101 |
| Smu\|SM000247S08301 | AGM2 | 38.830 | 188 | 98 | 5 | 10 | 193 | 19 | 193 | 1.61e-35 | 114 |
| Smu\|SM000247S08301 | AGM1 | 42.308 | 130 | 67 | 3 | 66 | 193 | 66 | 189 | 3.04e-29 | 98.2 |
| Smu\|SM000302S11676 | AGM2 | 38.298 | 188 | 99 | 5 | 10 | 193 | 19 | 193 | 6.69e-35 | 112 |
| Smu\|SM000302S11676 | AGM1 | 43.077 | 130 | 66 | 3 | 66 | 193 | 66 | 189 | 1.43e-29 | 99.0 |

Data S-51

https://itol.embl.de/tree/208127240222403281665195572

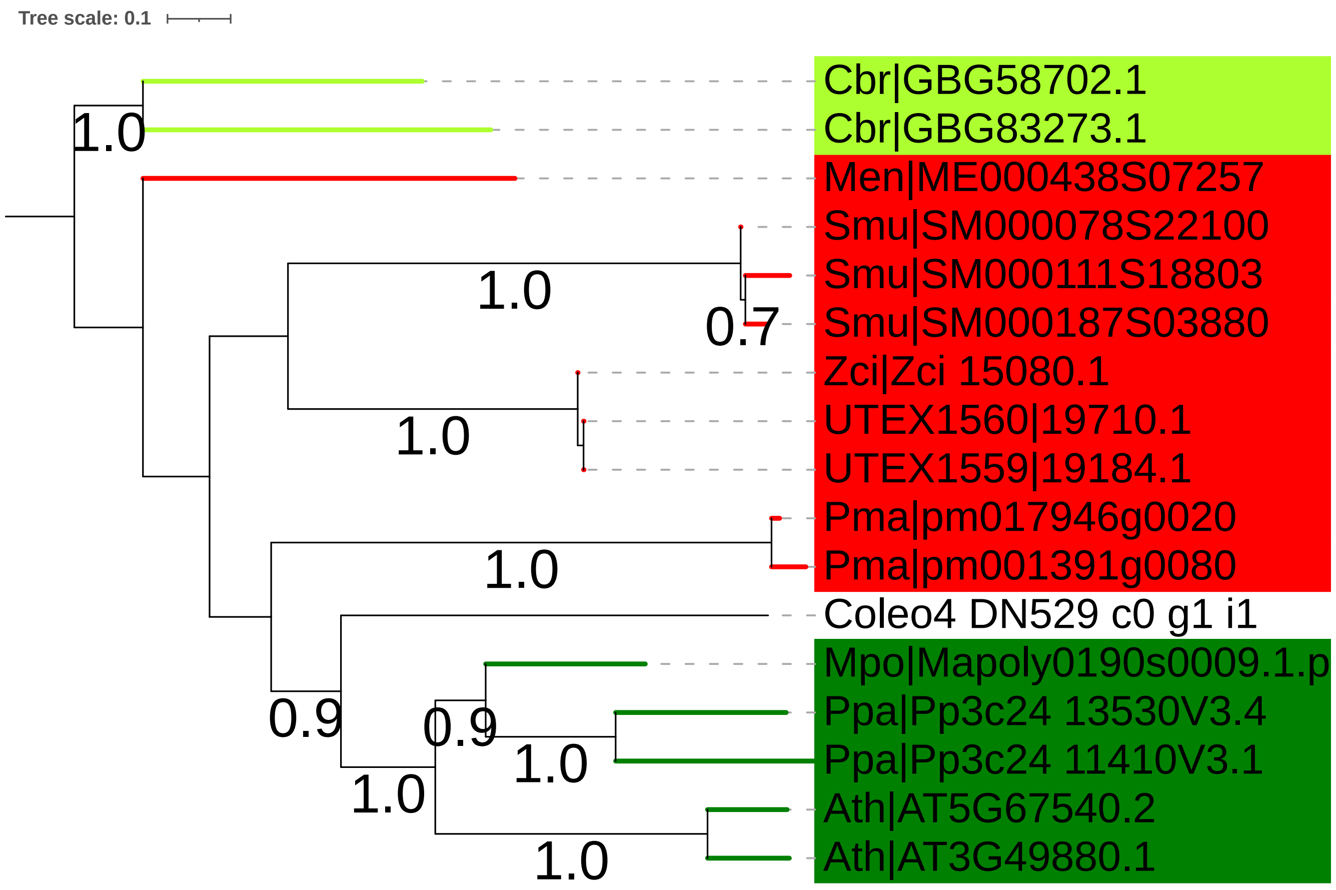


GH43A (AT5G67540)

GH43B (AT3G49880)

### GH43_24 + Coleochaete

Characterization of recombinant GH43 variants revealed that the exo-β-1,3-galactosidase activity of GH43 enzymes is hindered by β-1,6 branches on β-1,3-galactans.

Nibbering P, Petersen BL, Motawia MS, Jørgensen B, Ulvskov P, Niittylä T. Golgi-localized exo-β1,3-galactosidases involved in cell expansion and root growth in Arabidopsis. J Biol Chem. 2020 Jul 31;295(31):10581-10592. doi: 10.1074/jbc.RA120.013878.

Data S-52

# GH43_24 nr

**The Arabidopsis GH43_24 has been characterized to be involved in AGP backbone degradation.**

**GH43_24: exo-b-1,3-galactanase (EC 3.2.1.145), might be HGT from bacteria.**

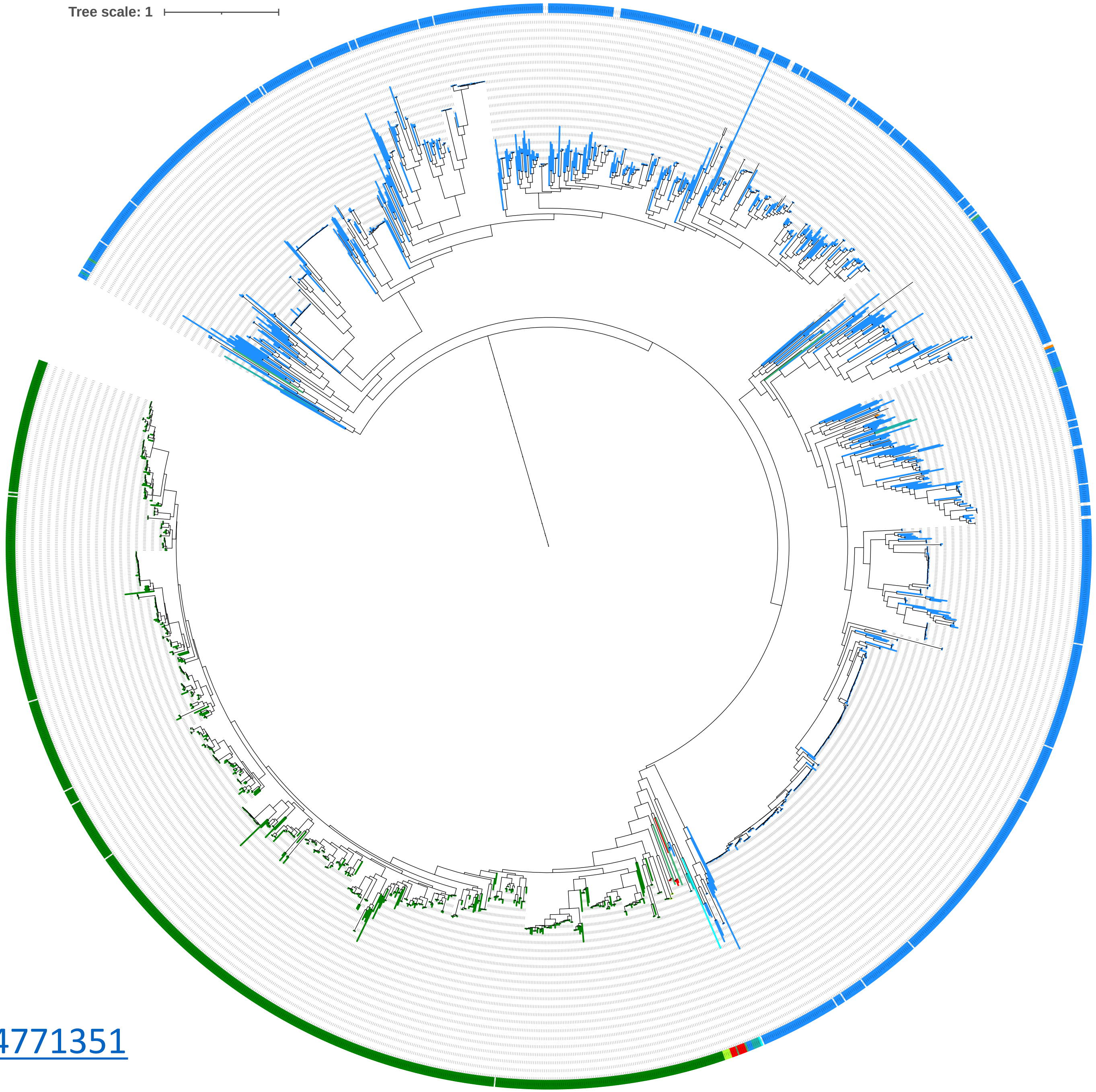


https://itol.embl.de/tree/20812770255164771664771351

Data S-53

# GH30_5

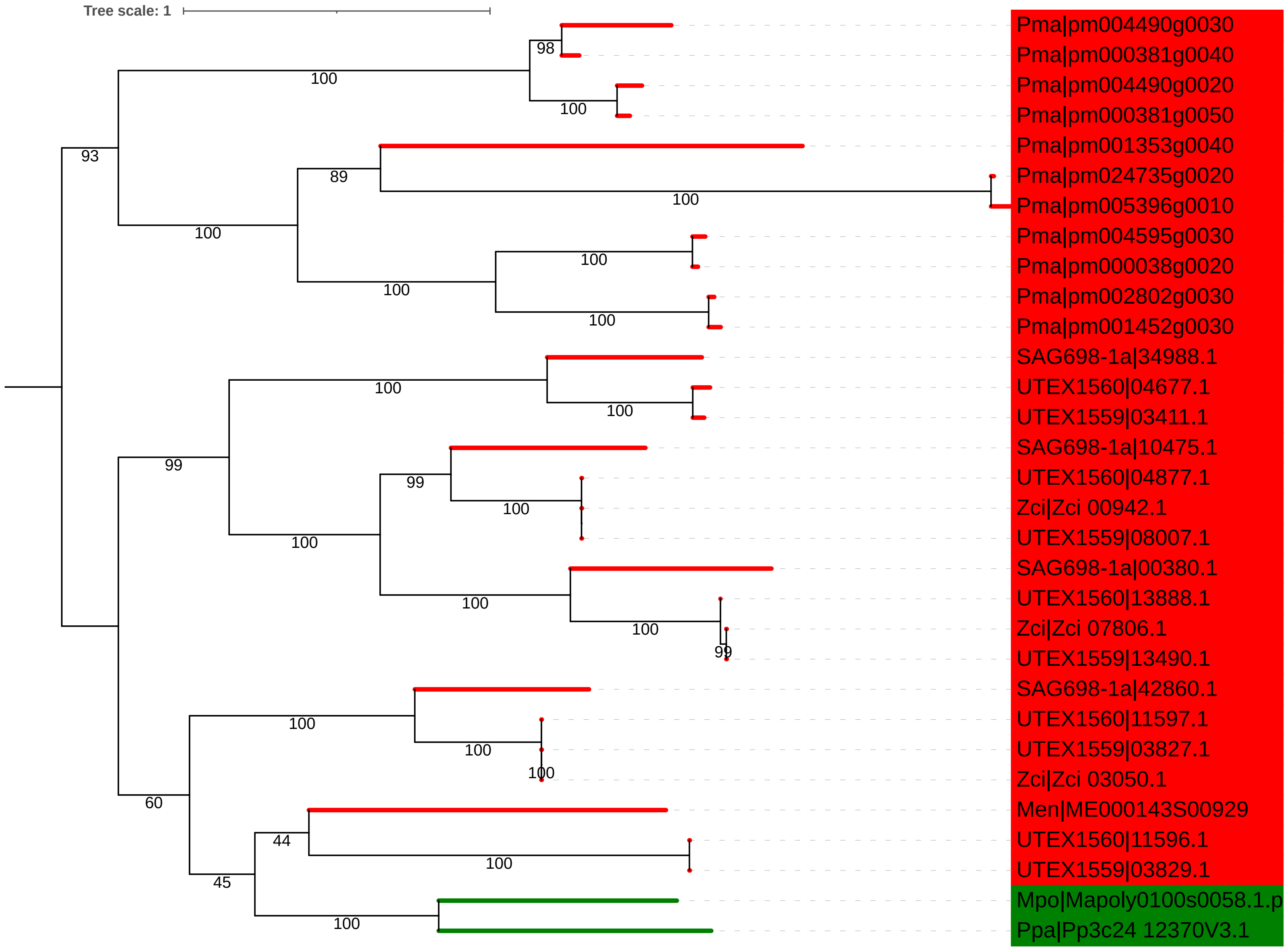


endo-b-1,6-galactanase (EC 3.2.1.164);galactan exo-1,6-b-galactobiohydrolase (non-reducing end) (EC 3.2.1.213)

No Coleochaete

Data S-54

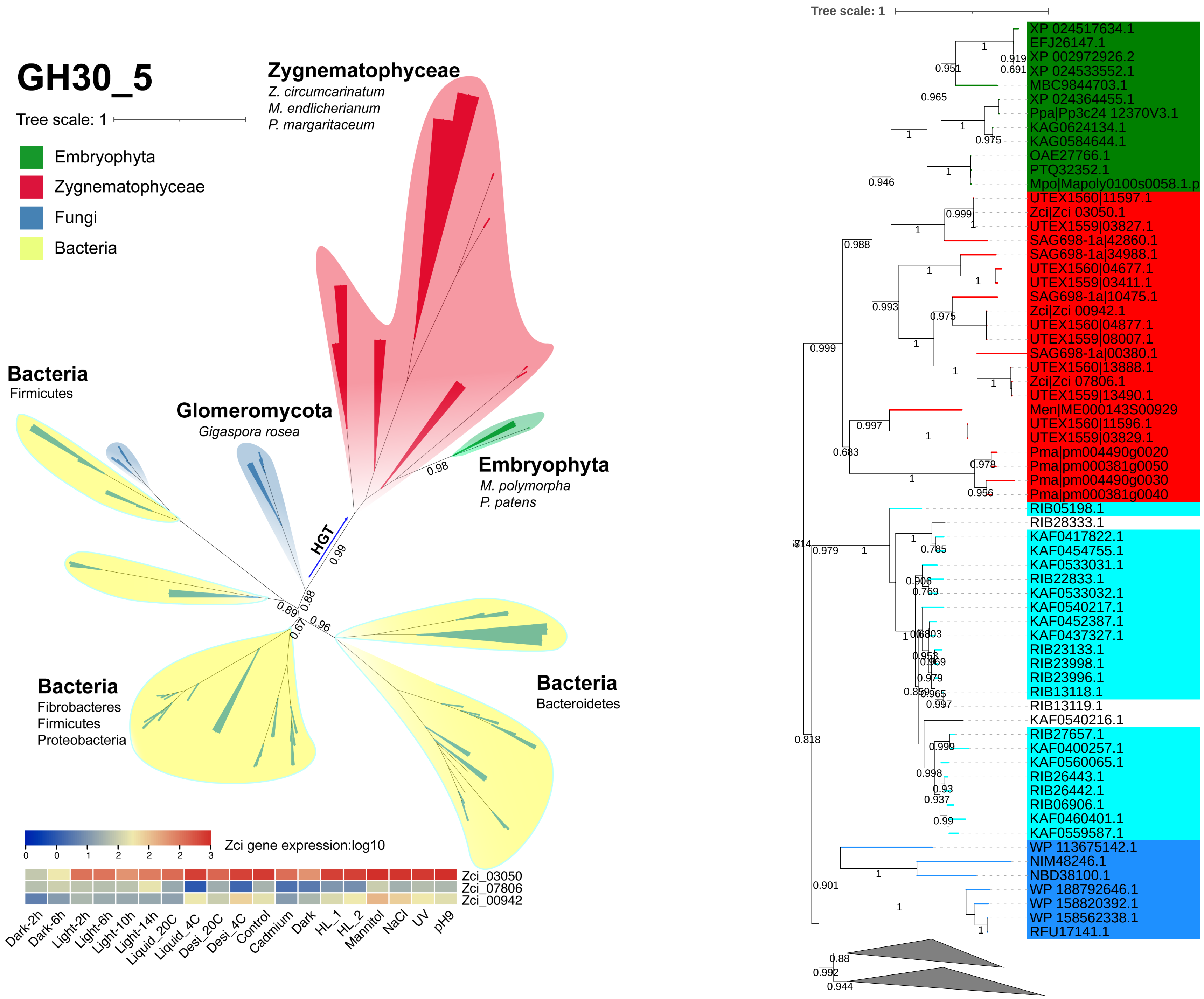
GH30_5
Tree scale: 1
Embryophyta
Zygnematophyceae
Fungi
Bacteria
Zygnematophyceae
Z. circumcarinatum
M. endlicherianum
P. margaritaceum
Bacteria
Firmicutes
Glomeromycota
Gigaspora rosea
Embryophyta
M. polymorpha
P. patens
HGT
Bacteria
Fibrobacteres
Firmicutes
Proteobacteria
Bacteria
Bacteroidetes
Zci gene expression:log10
Zci_03050
Zci_07806
Zci_00942
Dark-2h
Dark-6h
Light-2h
Light-6h
Light-10h
Light-14h
Liquid_20C
Liquid_4C
Desi_20C
Desi_4C
Control
Cadmium
Dark
HL_1
HL_2
Mannitol
NaCl
UV
pH9
Tree scale: 1
XP 024517634.1
EFJ26147.1
XP 002972926.2
XP 024533552.1
MBC9844703.1
XP 024364455.1
Ppa|Pp3c24 12370V3.1
KAG0624134.1
KAG0584644.1
OAE27766.1
PTQ32352.1
Mpo|Mapoly0100s0058.1.p
UTEX1560|11597.1
Zci|Zci 03050.1
UTEX1559|03827.1
SAG698-1a|42860.1
SAG698-1a|34988.1
UTEX1560|04677.1
UTEX1559|03411.1
SAG698-1a|10475.1
Zci|Zci 00942.1
UTEX1560|04877.1
UTEX1559|08007.1
SAG698-1a|00380.1
UTEX1560|13888.1
Zci|Zci 07806.1
UTEX1559|13490.1
Men|ME000143S00929
UTEX1560|11596.1
UTEX1559|03829.1
Pma|pm004490g0020
Pma|pm000381g0050
Pma|pm004490g0030
Pma|pm000381g0040
RIB05198.1
RIB28333.1
KAF0417822.1
KAF0454755.1
KAF0533031.1
RIB22833.1
KAF0533032.1
KAF0540217.1
KAF0452387.1
KAF0437327.1
RIB23133.1
RIB23998.1
RIB23996.1
RIB13118.1
RIB13119.1
KAF0540216.1
RIB27657.1
KAF0400257.1
KAF0560065.1
RIB26443.1
RIB26442.1
RIB06906.1
KAF0460401.1
KAF0559587.1
WP 113675142.1
NIM48246.1
NBD38100.1
WP 188792646.1
WP 158820392.1
WP 158562338.1
RFU17141.1

Data S-55

https://itol.embl.de/tree/208127240222737916651977706

# GH35

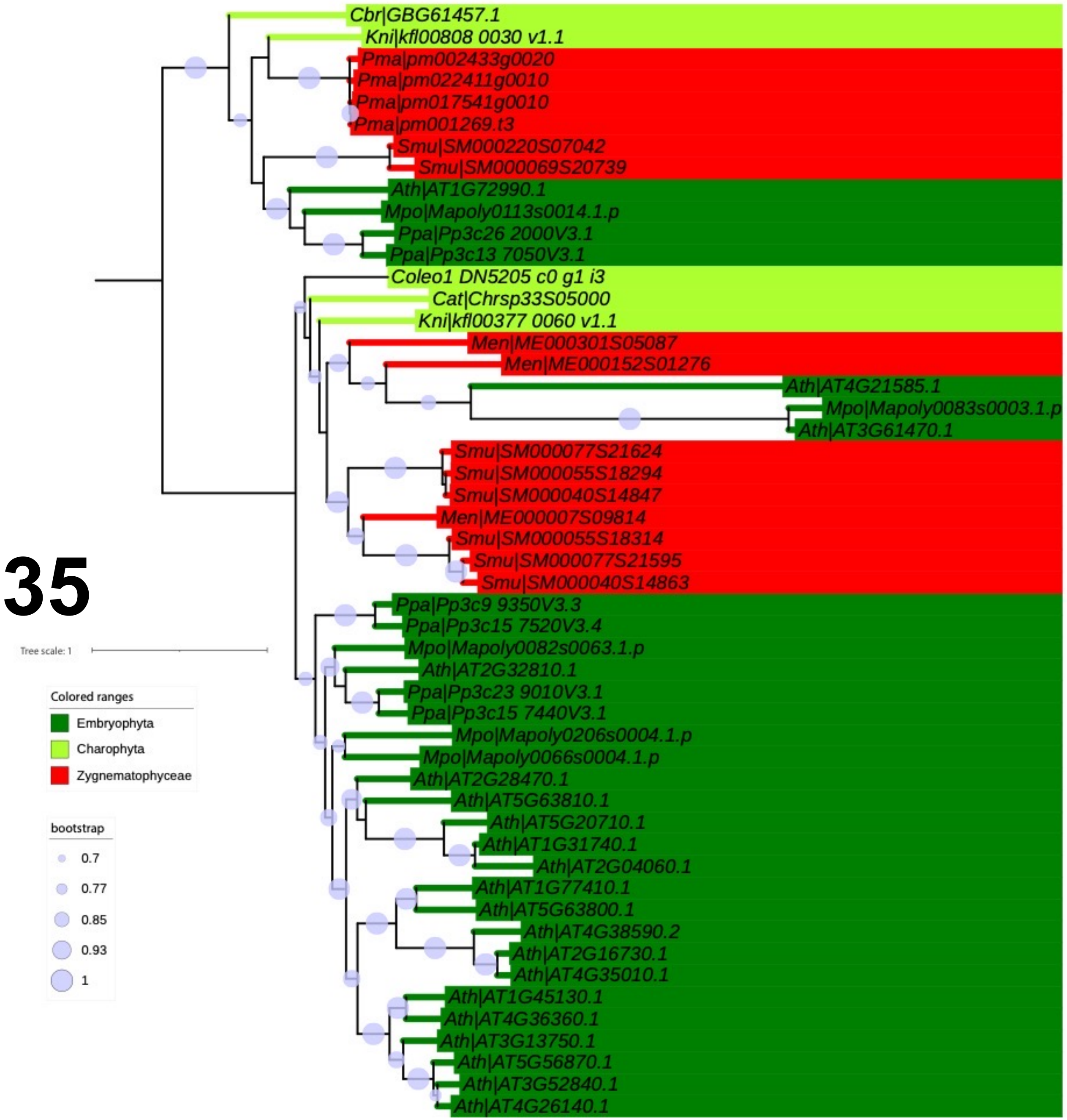


**AT3G61470 (LHCA2):** photosystem I light harvesting complex gene 2

**AT4G21585 (ENDO4): Endonuclease 4**

BGAL8 (AT3G28470)

BGAL10 (AT5g63810)

Kotake T, Dina S, Konishi T, Kaneko S, Igarashi K, Samejima M, Watanabe Y, Kimura K, Tsumuraya Y. Molecular cloning of a {beta}-galactosidase from radish that specifically hydrolyzes {beta}-(1->3)- and {beta}-(1->6)-galactosyl residues of Arabinogalactan protein. Plant Physiol. 2005 Jul;138(3):1563-76. doi: 10.1104/pp.105.062562.

Sampedro et al. AtBGAL10 is the main xyloglucan β-galactosidase in Arabidopsis, and its absence results in unusual xyloglucan subunits and growth defects. Plant Physiol. 2012 Mar;158(3):1146-57. doi: 10.1104/pp.111.192195.

Data S-56

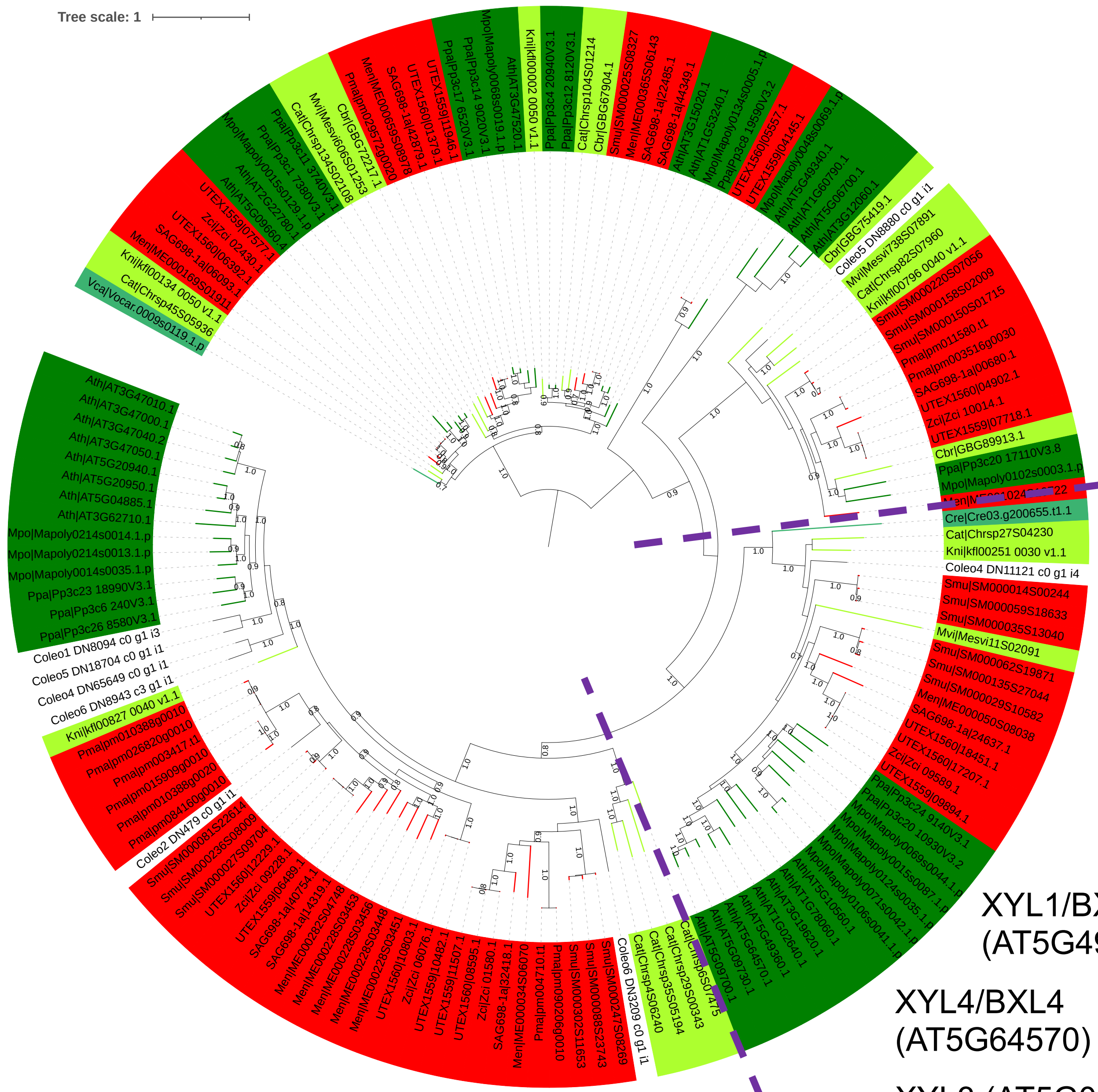


XYL1/BXL1 (AT5G49360)

XYL4/BXL4 (AT5G64570)

XYL3 (AT5G09730)

GH3 + Coleochaete

Minic et al. Purification, functional characterization, cloning, and identification of mutants of a seed-specific arabinan hydrolase in Arabidopsis. J Exp Bot. 2006;57(10):2339-51. doi: 10.1093/jxb/erj205.

Kotake T, Tsuchiya K, Aohara T, Konishi T, Kaneko S, Igarashi K, Samejima M, Tsumuraya Y. An alpha-L-arabinofuranosidase/beta-D-xylosidase from immature seeds of radish (Raphanus sativus L.). J Exp Bot. 2006;57(10):2353-62. doi: 10.1093/jxb/erj206.

Minic Z, Rihouey C, Do CT, Lerouge P, Jouanin L. Purification and characterization of enzymes exhibiting beta-D-xylosidase activities in stem tissues of Arabidopsis. Plant Physiol. 2004 Jun;135(2):867-78. doi: 10.1104/pp.104.041269.

XYL3: released L-arabinose from (1-->5)-alpha-L-arabinofuranobiose, arabinoxylan, sugar beet arabinan, and debranched arabinan.
**RG-I side chain degradation**

**XYL4: release mainly D-Xyl from oat spelt xylan, rye arabinoxylan, wheat arabinoxylan, and oligoarabinoxylans. ARAf and XYL1 can also release D-Xyl from these substrates but less efficiently than XYL4. Moreover, they can also release L-Ara from arabinoxylans and arabinan.**

**AtBXL1: putative bifunctional beta-d-xylosidase/alpha-l-arabinofuranosidase; implicated as beta-d-xylosidase acting during vascular development**
**GH3: hydrolytic activity on radish AGPs, pectic α-1,5-arabinan and arabinoxylan.**

RsAraf1: α-arabinofuranosidase in the GH3 family; heterologously expressed in Arabidopsis, RsAraf1 hydrolyzed α-arabinofuranosyl residues of AGPs (Kotake et al., 2006)

Data S-57

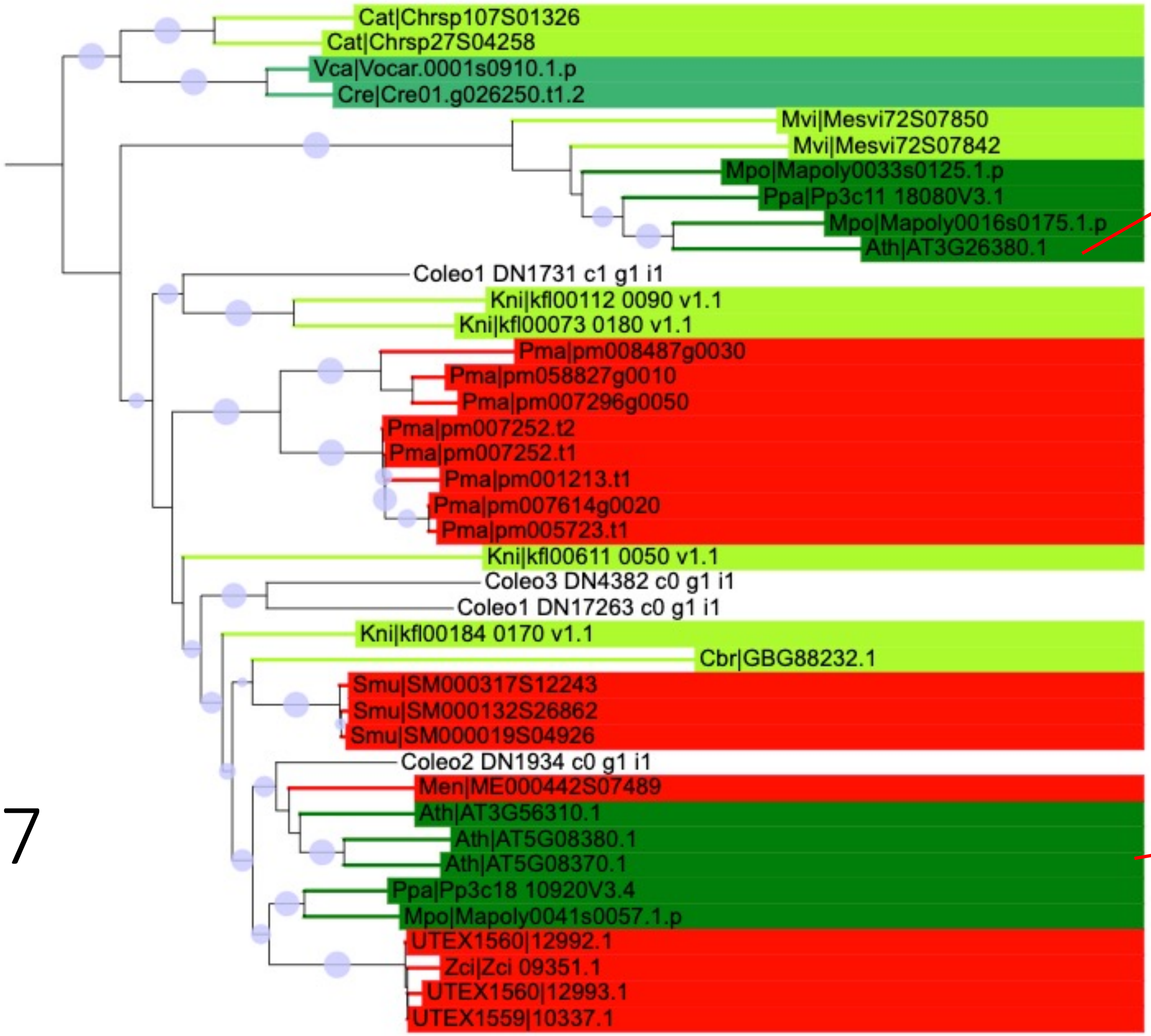


APSE (AT3G26380)

**APSE and AGALs also contribute to the hydrolysis of β-l-Arap residues of pectic α-1,3:1,5-arabinan and type I AG.**

The β-l-Arap residues of AGPs are hydrolysed mainly by APSE and partially by AGALs in Arabidopsis.

Imaizumi et al.: Heterologous expression and characterization of an Arabidopsis β-l-arabinopyranosidase and α-d-galactosidases acting on β-l-arabinopyranosyl residues. J Exp Bot. 2017 Jul 20;68(16):4651-4661. doi: 10.1093/jxb/erx279.

AGAL1 (AT5G08380)
AGAL2 (AT5G08370)
AGAL3 (AT3G56310)

GH27

https://itol.embl.de/tree/2081272402224247916652018640

Data S-58

APSE: might have been gained via HGT from bacteria

# GH27 nr

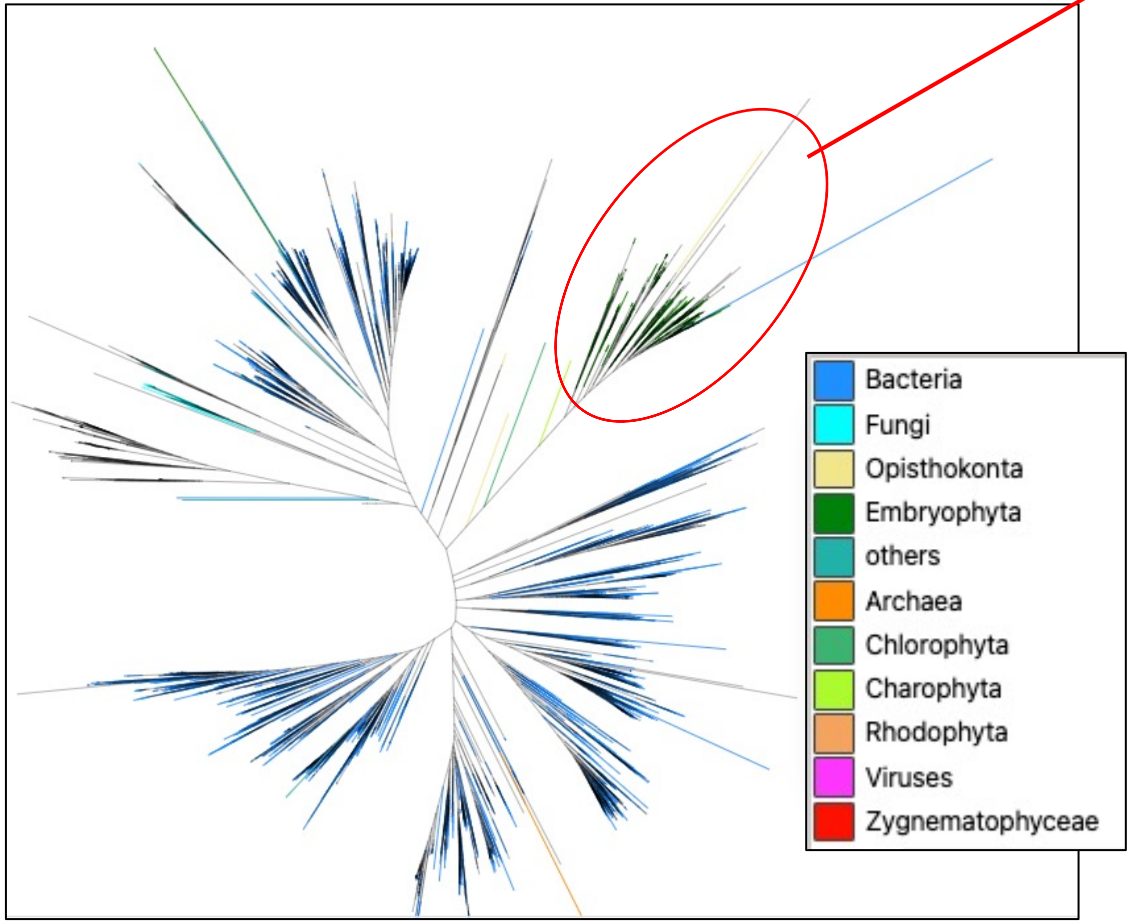


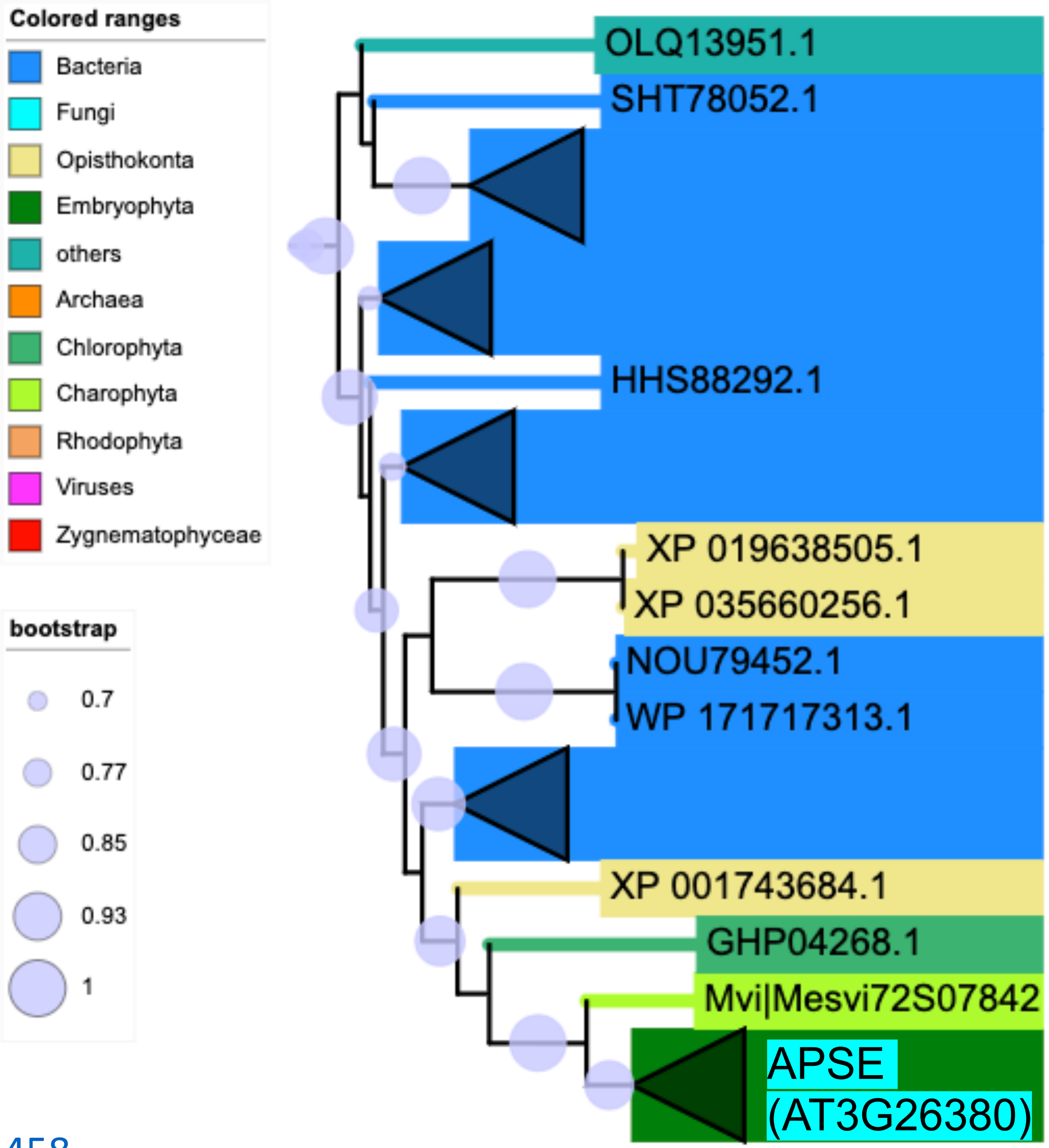


https://itol.embl.de/tree/20812724022256331665203458

Data S-59

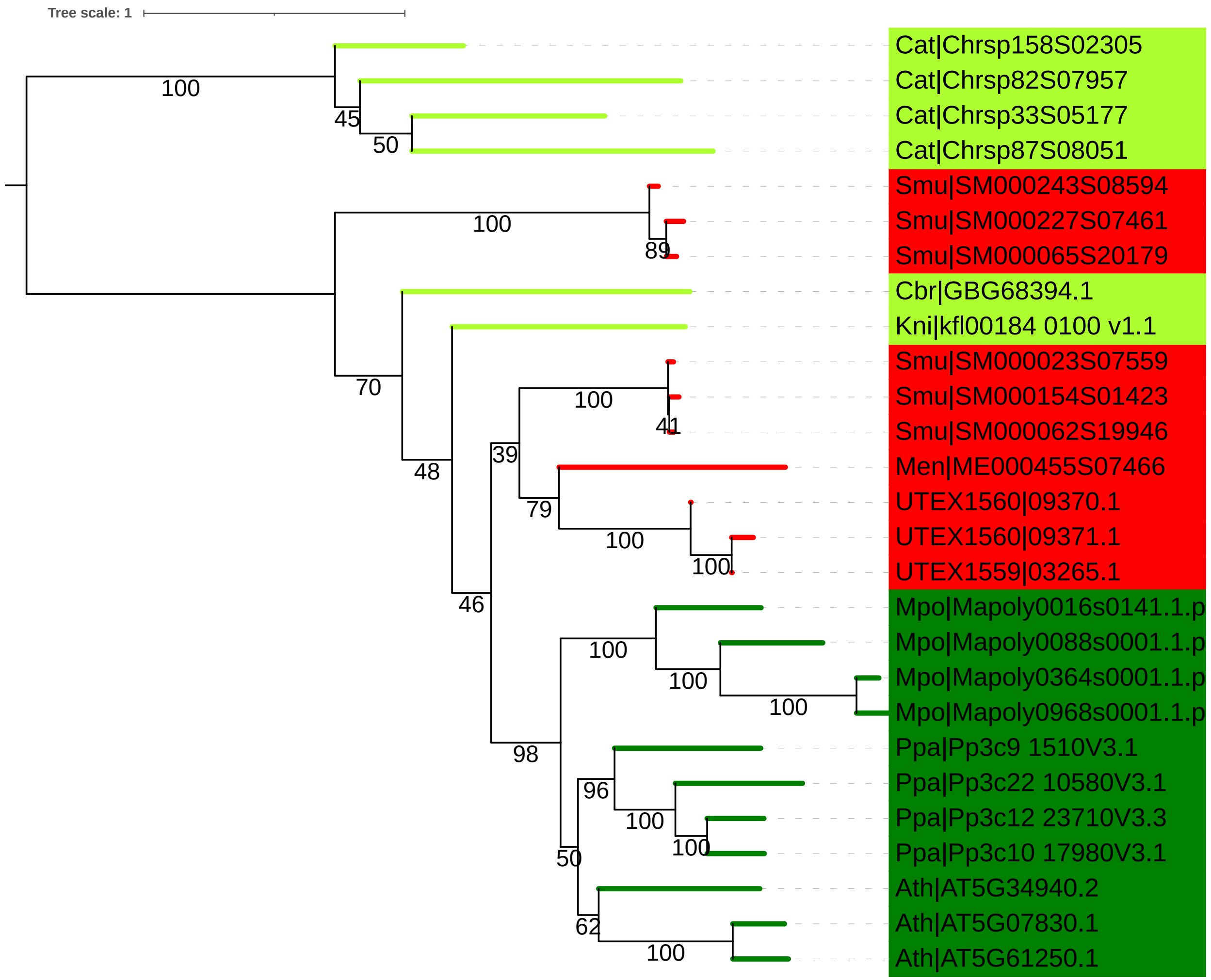


# GH79

Eudes, A., Mouille, G., Thévenin, J., Goyallon, A., Minic, Z., and Jouanin, L. (2008). Purification, cloning and functional characterization of an endogenous beta-glucuronidase in Arabidopsis thaliana. Plant Cell Physiol. 49, 1331–1341. doi: 10.1093/pcp/pcn108

AtGUS1 (AT5G61250)

AtGUS2 (AT5G07830)

AtGUS3 (AT5G34940)

No GH79 was found in the Coleochaete transcriptome sequences.

Data S-60

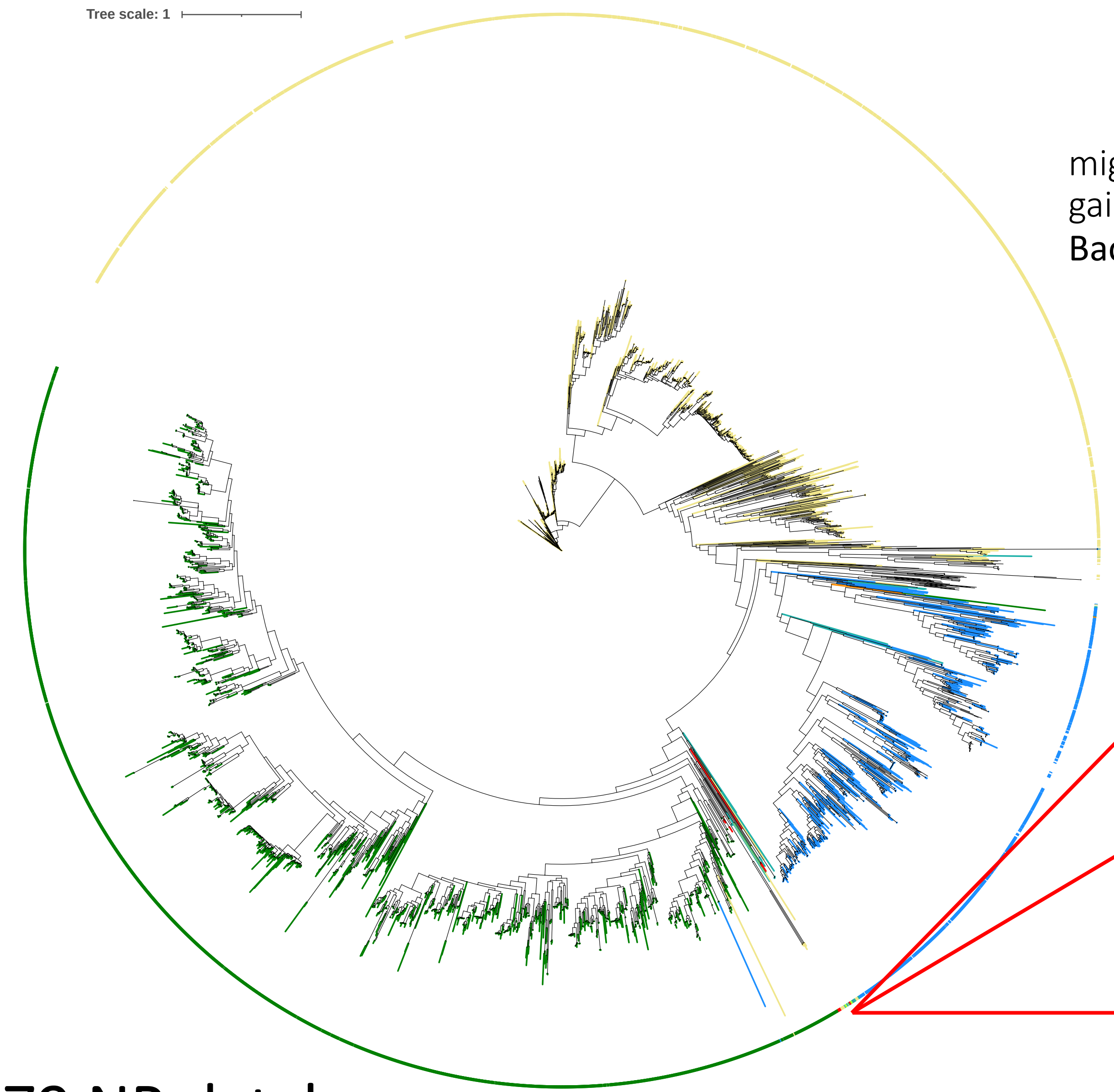


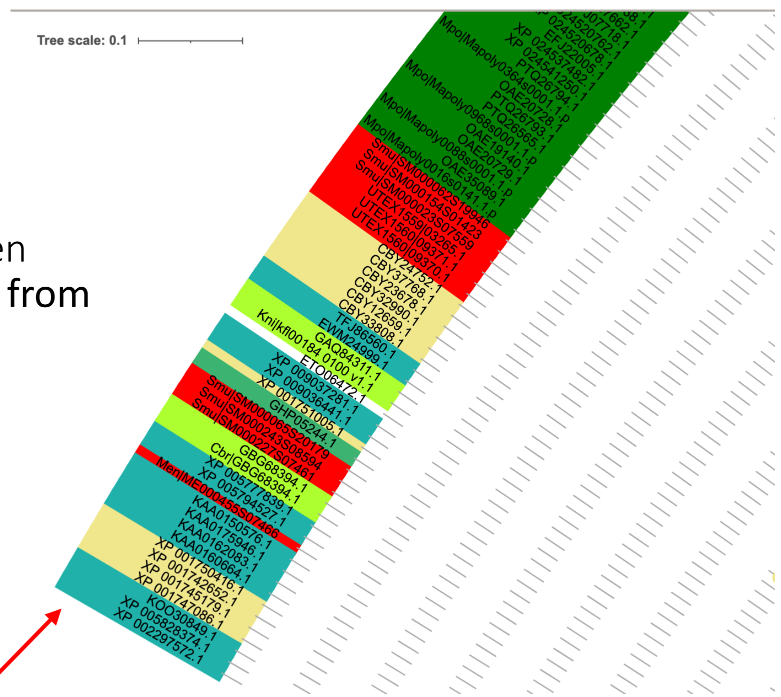


might have been gained via **HGT from Bacteria**

Zynema (UTEX1559, UTEX1560)
Klebsormidium
Chara

**Amorphea** (Monosiga brevicollis)
**Animalia** (Oikopleura dioica)
**SAR** (Nannochloropsis salina, Aureococcus anophagefferens)
**Chlorophyta** (Pycnococcus provasolii)

### GH79 NR database

Data S-61

### Homogalacturonan (HG)

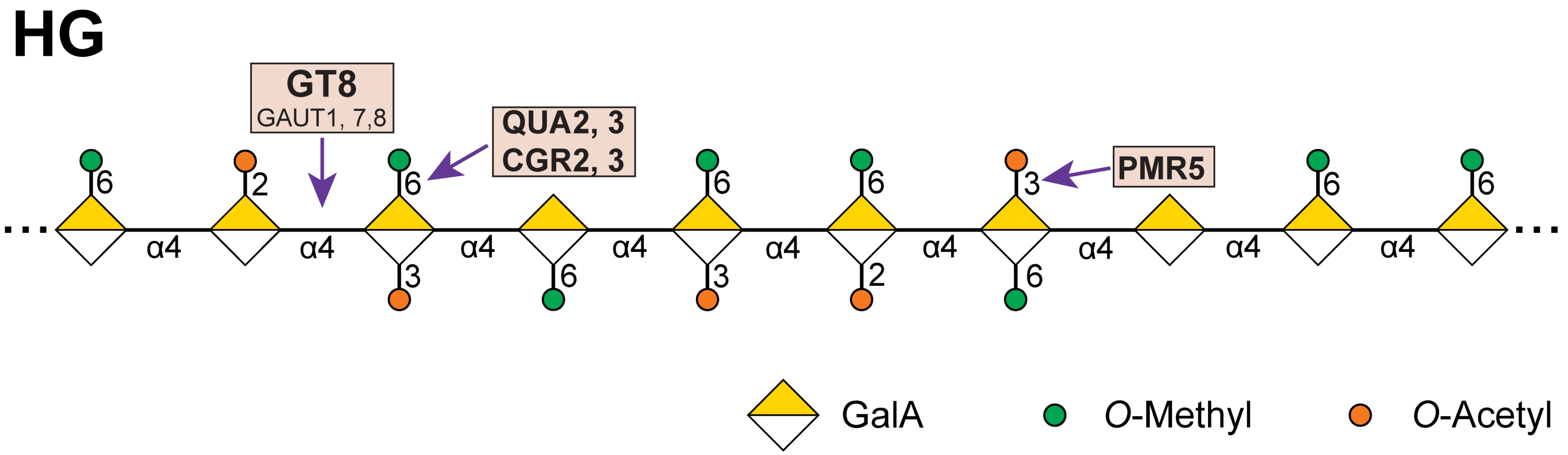


- galacturonic acid (α-D-GalA)
- Approximately **65%** of pectin is HG, which is a homopolymer of D-GalA linked in an α- 1,4 configuration
- GALACTURONOSYL- TRANSFERASE (GAUT)
- **Modifications:**
- Up to 80% of the carboxyl groups may be methylesterified.
- GalA may be *O*-acetylated at *O*-2 or *O*-3.
- Some GalA may be substituted at *O*-3 with β-D-Xyl, forming xylogalacturonan (XGA). The Xyl may be further elongated at the *O*-2 position by another β-D-Xyl; additional extension with Xyl residues has been observed in soybean.
- GalA in some plants, especially aquatic, may be β-substituted with D-Api*f* at *O*-2 and/or *O*-3, forming apiogalacturonan (AGA).
- Possible *trans*-esters have been proposed between the carboxyl group and *O*-2 or *O*-3 hydroxyls.

Data S-62

https://itol.embl.de/tree/208127240222263521665352415

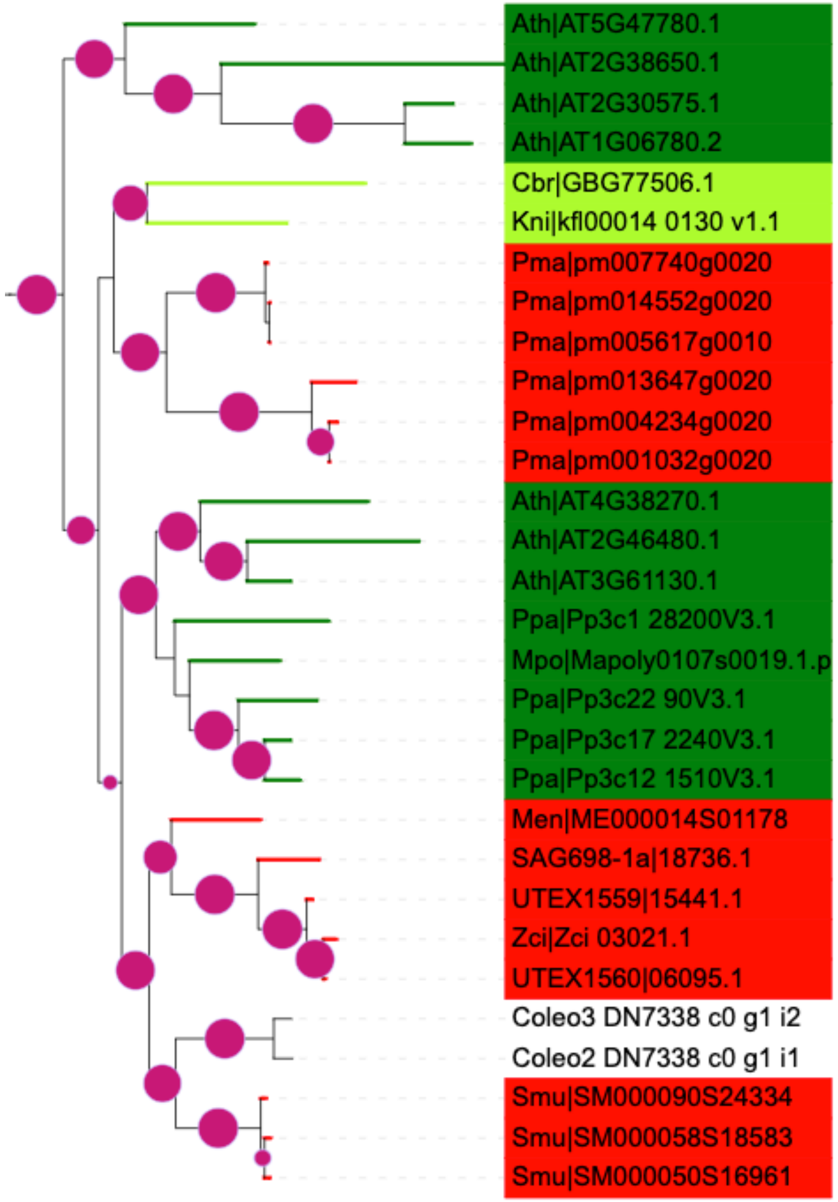


**GAUT6** (AT1G06780)
**GAUT5** (AT2G30575)
**GAUT7** (AT2G38650)
**GAUT4** (AT5G47780)

**GAUT1** (AT3G61130)
**GAUT2** (AT2G46480)
**GAUT3** (AT4G38270)

Atmodjo et al. Galacturonosyltransferase (GAUT)1 and GAUT7 are the core of a plant cell wall pectin biosynthetic homogalacturonan:galacturonosyltransferase complex. Proc Natl Acad Sci U S A. 2011 Dec 13;108(50):20225-30. doi: 10.1073/pnas.1112816108.

Sterling et al. Functional identification of an Arabidopsis pectin biosynthetic homogalacturonan galacturonosyltransferase. Proc Natl Acad Sci U S A. 2006 Mar 28;103(13):5236-41. doi: 10.1073/pnas.0600120103.

Bouton et al. QUASIMODO1 encodes a putative membrane-bound glycosyltransferase required for normal pectin synthesis and cell adhesion in Arabidopsis. Plant Cell. 2002 Oct;14(10):2577-90. doi: 10.1105/tpc.004259.

**GAUT8** (QUA1, AT3G25140)

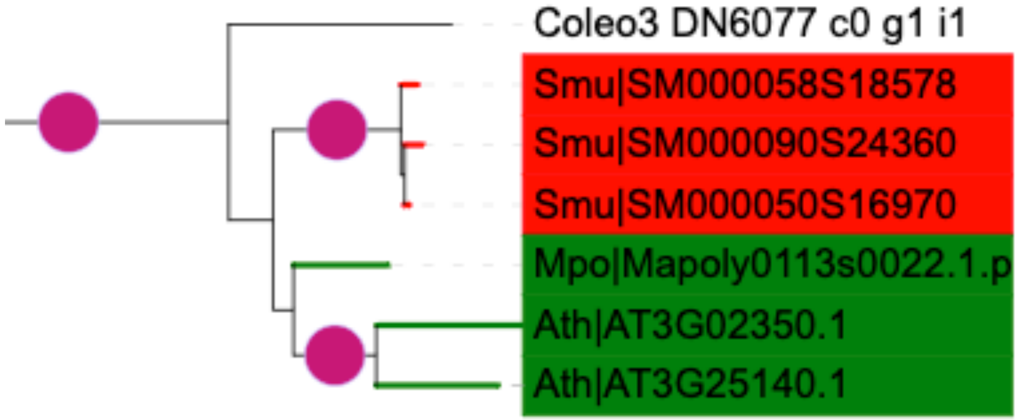

Data S-63

### QUA2 (AT1G78240)

Blast with 16genome + Coleochaete

HG is methylesterified by QUA2, QUA3 and GGR3, of which QUA2 might be first present in Chara that is consistent with the HG in Chara (also experiment verified).

QUA3 (AT4G00740) might be first present in Zygnema

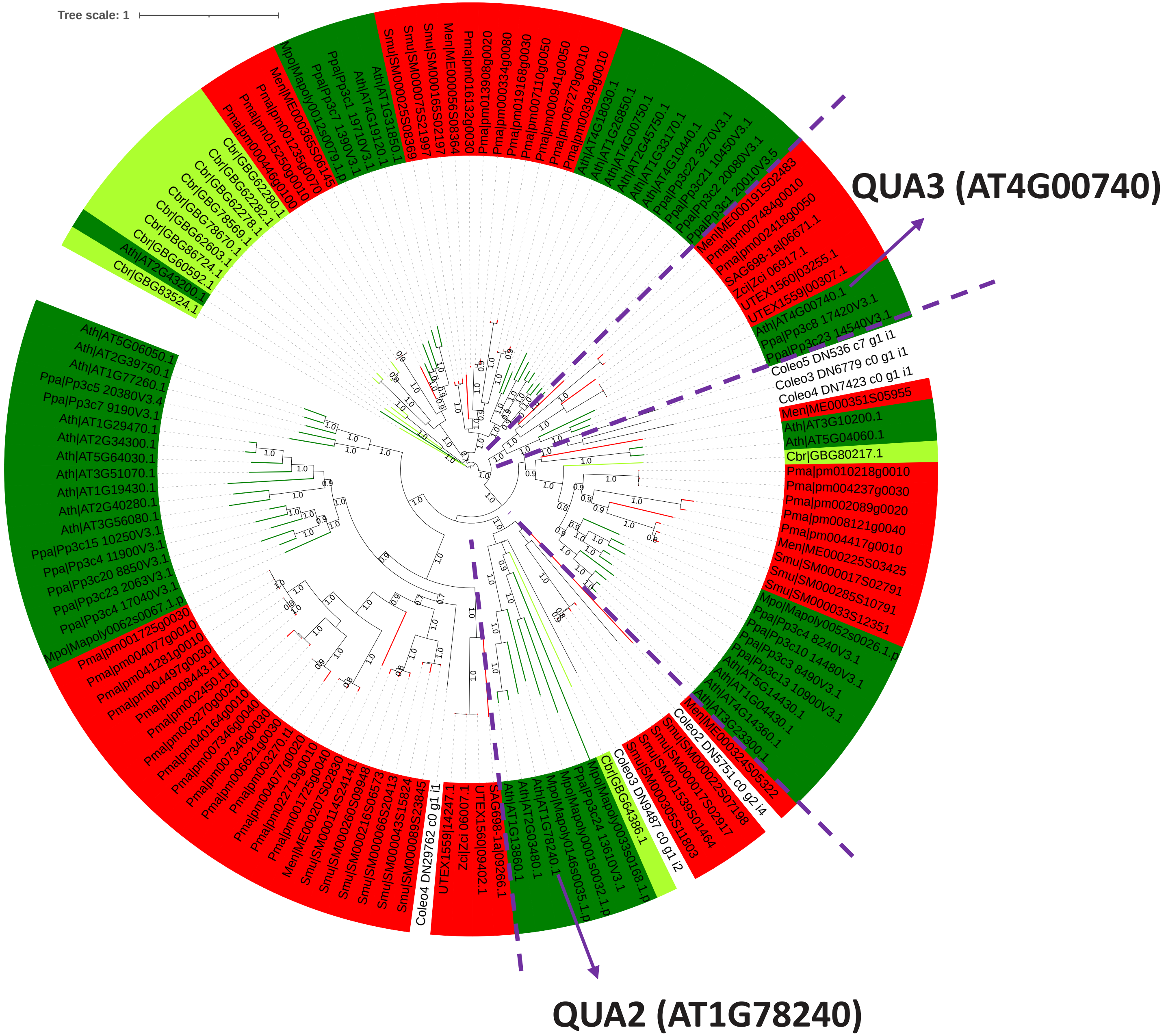

Data S-64

Kim SJ, Held MA, Zemelis S, Wilkerson C, Brandizzi F. CGR2 and CGR3 have critical overlapping roles in pectin methylesterification and plant growth in Arabidopsis thaliana. Plant J. 2015 Apr;82(2):208-20. doi: 10.1111/tpj.12802.

### CGR2/3 homologs (evalue<1e-15)

| | | | | | | | | | | | |
|---|---|---|---|---|---|---|---|---|---|---|---|
| Ath\|AT5G65810.1 | CGR2\|AT3G49720 | 86.590 | 261 | 32 | 1 | 1 | 258 | 1 | 261 | 1.59e-174 | 469 |
| Ath\|AT3G49720.1 | CGR2\|AT3G49720 | 100.000 | 261 | 0 | 0 | 1 | 261 | 1 | 261 | 0.0 | 534 |
| Cbr\|GBG71667.1 | CGR2\|AT3G49720 | 30.075 | 133 | 91 | 2 | 60 | 190 | 94 | 226 | 2.26e-15 | 64.7 |
| Mpo\|Mapoly0082s0061.1.p | CGR2\|AT3G49720 | 34.201 | 269 | 167 | 6 | 40 | 306 | 1 | 261 | 6.71e-49 | 151 |
| Mvi\|Mesvi21S04311 | CGR2\|AT3G49720 | 27.174 | 184 | 119 | 8 | 70 | 243 | 48 | 226 | 1.58e-15 | 61.6 |
| Ppa\|Pp3c12_13990V3.1 | CGR2\|AT3G49720 | 36.397 | 272 | 128 | 6 | 31 | 298 | 27 | 257 | 1.54e-55 | 168 |
| Coleo1_DN62522_c0_g1_i1 | CGR2\|AT3G49720 | 29.487 | 78 | 55 | 0 | 12 | 89 | 149 | 226 | 1.55e-13 | 50.4 |

| | | | | | | | | | | | |
|---|---|---|---|---|---|---|---|---|---|---|---|
| Ath\|AT5G65810.1 | GGR3\|AT5G65810 | 100.000 | 258 | 0 | 0 | 1 | 258 | 1 | 258 | 0.0 | 529 |
| Ath\|AT3G49720.1 | GGR3\|AT5G65810 | 86.590 | 261 | 32 | 1 | 1 | 261 | 1 | 258 | 6.63e-174 | 469 |
| Cbr\|GBG71667.1 | GGR3\|AT5G65810 | 29.927 | 137 | 94 | 2 | 56 | 190 | 87 | 223 | 2.20e-14 | 63.5 |
| Mpo\|Mapoly0082s0061.1.p | GGR3\|AT5G65810 | 35.124 | 242 | 148 | 4 | 67 | 306 | 24 | 258 | 3.60e-49 | 153 |
| Mvi\|Mesvi21S04311 | GGR3\|AT5G65810 | 30.827 | 133 | 89 | 3 | 114 | 243 | 91 | 223 | 1.86e-13 | 57.0 |
| Ppa\|Pp3c12_13990V3.1 | GGR3\|AT5G65810 | 36.765 | 272 | 127 | 6 | 31 | 298 | 24 | 254 | 1.59e-59 | 180 |
| Coleo1_DN62522_c0_g1_i1 | GGR3\|AT5G65810 | 29.487 | 78 | 55 | 0 | 12 | 89 | 146 | 223 | 8.25e-12 | 47.4 |

Data S-65

### **PMR5 (<u>AT5G58600</u>)**

#### Blast with 16genome + Coleochaete

PMR5 (AT5G58600) might be first present in Chara.

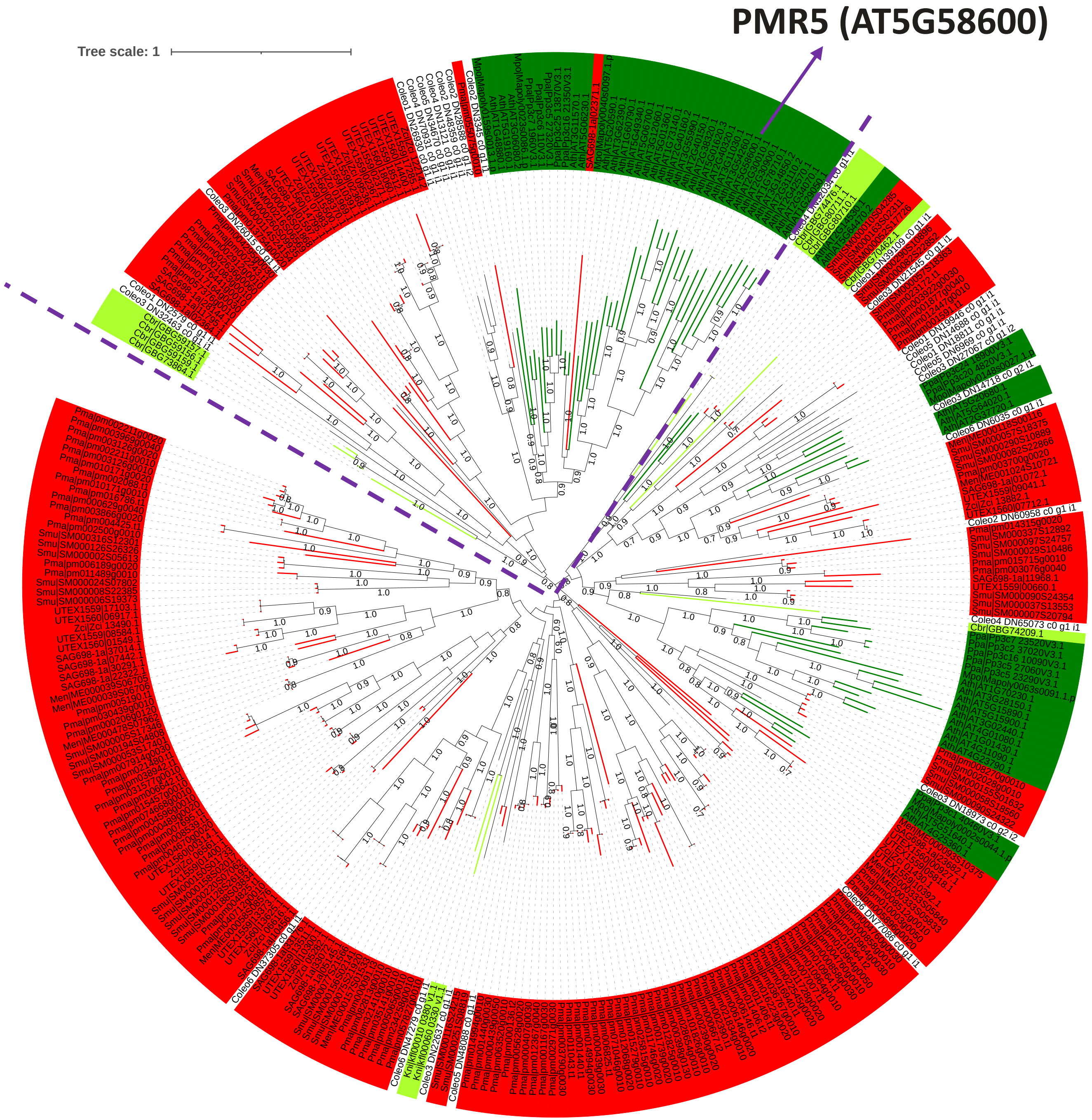


Chiniquy D, Underwood W, Corwin J, Ryan A, Szemenyei H, Lim CC, Stonebloom SH, Birdseye DS, Vogel J, Kliebenstein D, Scheller HV, Somerville S. PMR5, an acetylation protein at the intersection of pectin biosynthesis and defense against fungal pathogens. Plant J. 2019 Dec;100(5):1022-1035. doi: 10.1111/tpj.14497.

Data S-66

https://itol.embl.de/tree/1342381681245654166831464 4

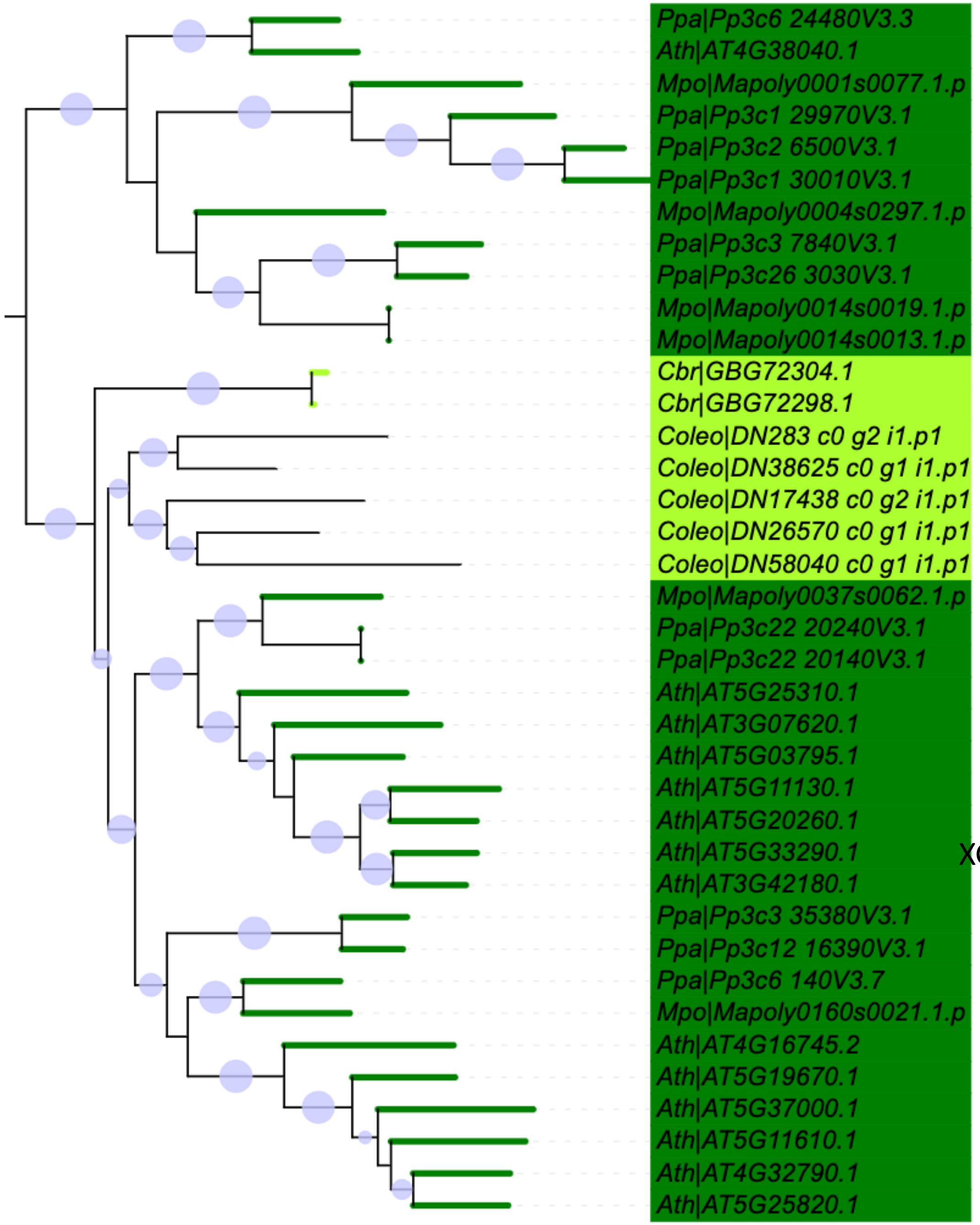


GT47 (Data S-33)

Jensen et al. Identification of a xylogalacturonan xylosyltransferase involved in pectin biosynthesis in Arabidopsis. Plant Cell. 2008 May;20(5):1289-302. doi: 10.1105/tpc.107.050906

XGD1 (AT5G33290)

**XGA**

Data S-67

https://itol.embl.de/tree/1342381641713026616669281512

### CE8: pectin methylesterase

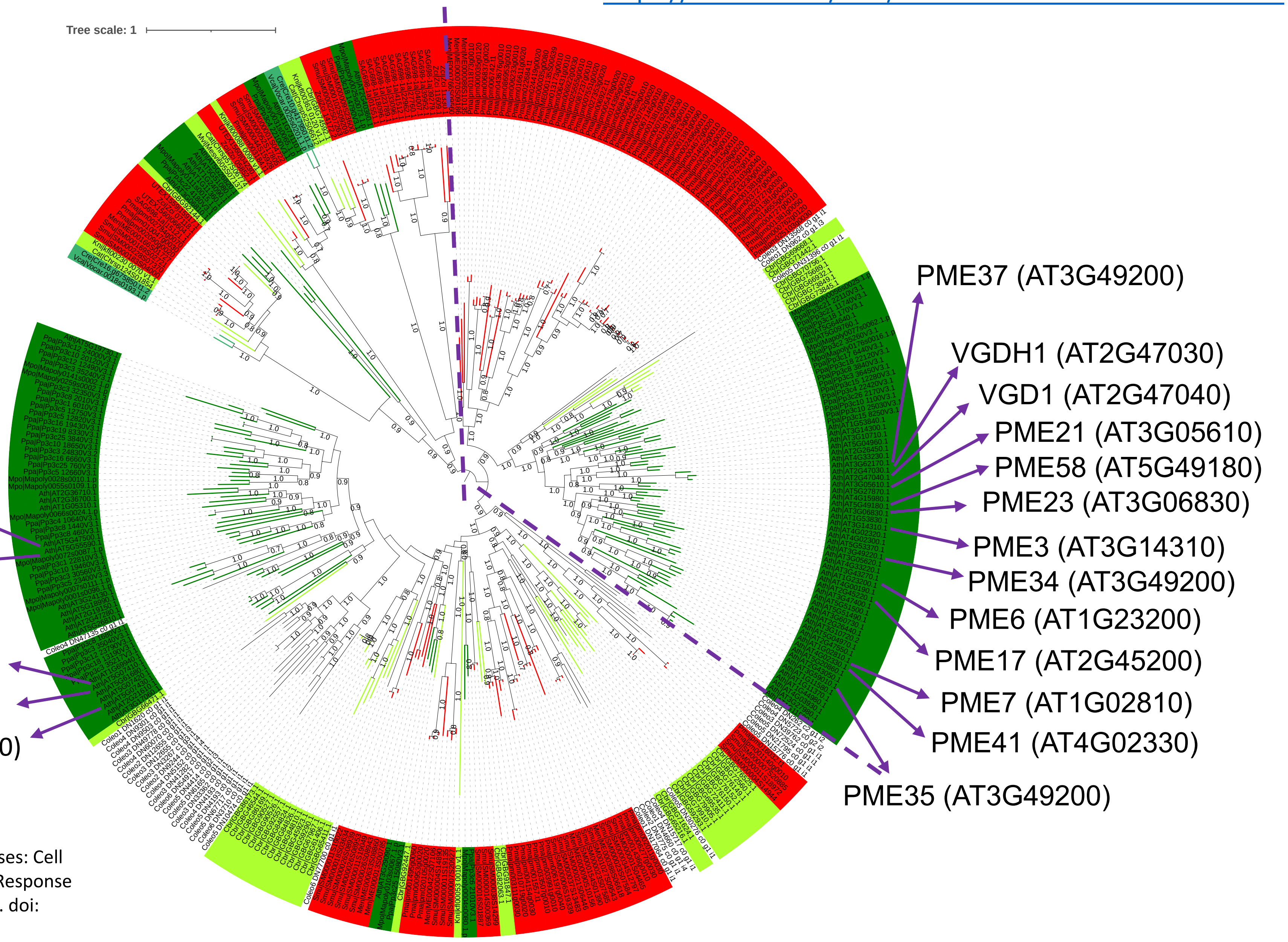


Wu HC, Bulgakov VP, Jinn TL. Pectin Methylesterases: Cell Wall Remodeling Proteins Are Required for Plant Response to Heat Stress. Front Plant Sci. 2018 Nov 6;9:1612. doi: 10.3389/fpls.2018.01612.

Data S-68

### CE13: pectin acetylesterase

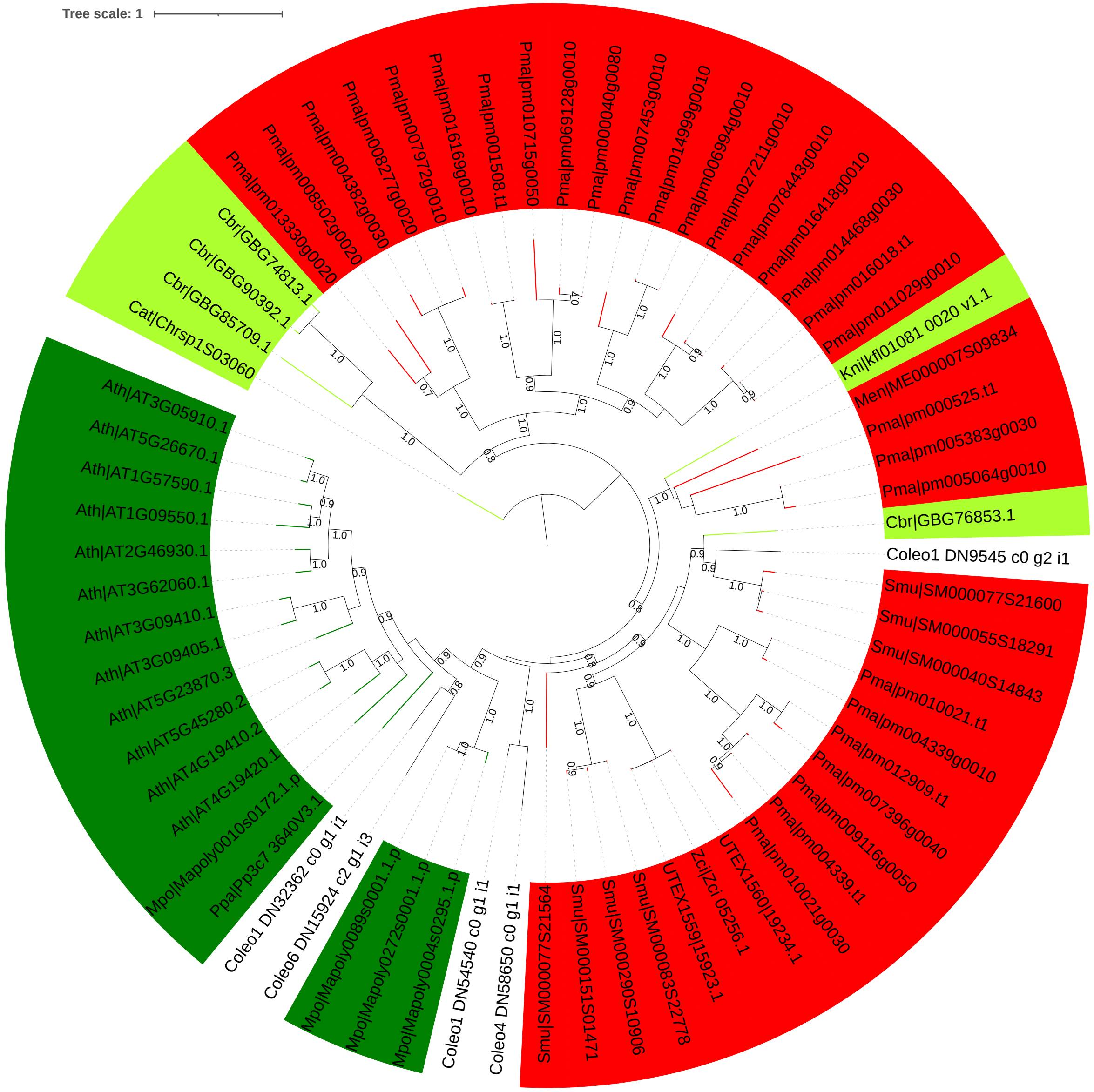


Philippe F, Pelloux J, Rayon C. Plant pectin acetylesterase structure and function: new insights from bioinformatic analysis. BMC Genomics. 2017 Jun 8;18(1):456. doi: 10.1186/s12864-017-3833-0.

Data S-69

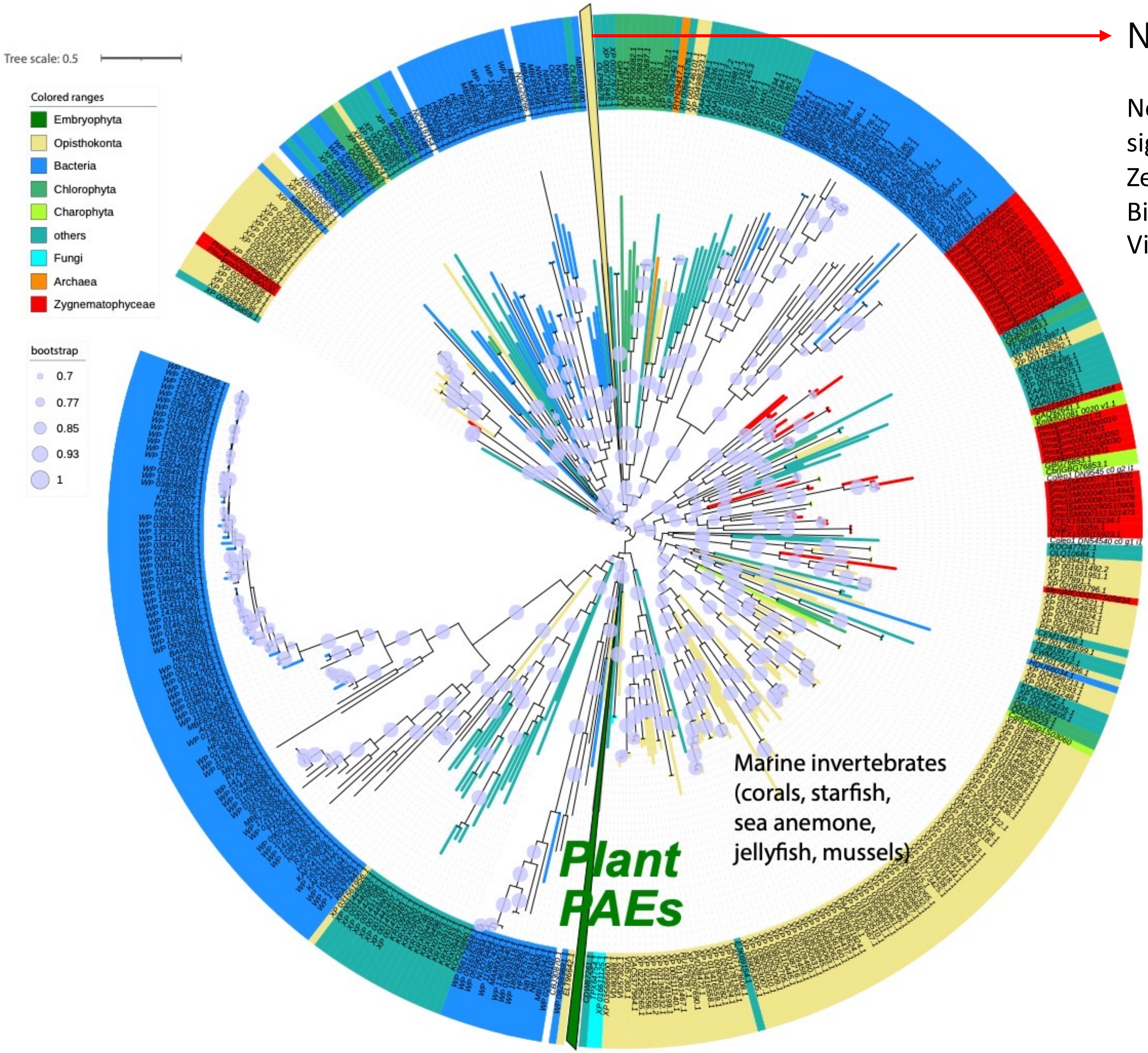


Notum from animals

Notum deacylates Wnt proteins to suppress signalling activity. Kakugawa S, Langton PF, Zebisch M, Howell SA, Chang TH, Liu Y, Feizi T, Bineva G, O'Reilly N, Snijders AP, Jones EY, Vincent JP. Nature 519, 187-92, (2015).

CE13 nr

### Rhamnogalacturonan I (RG-I)

RG-I

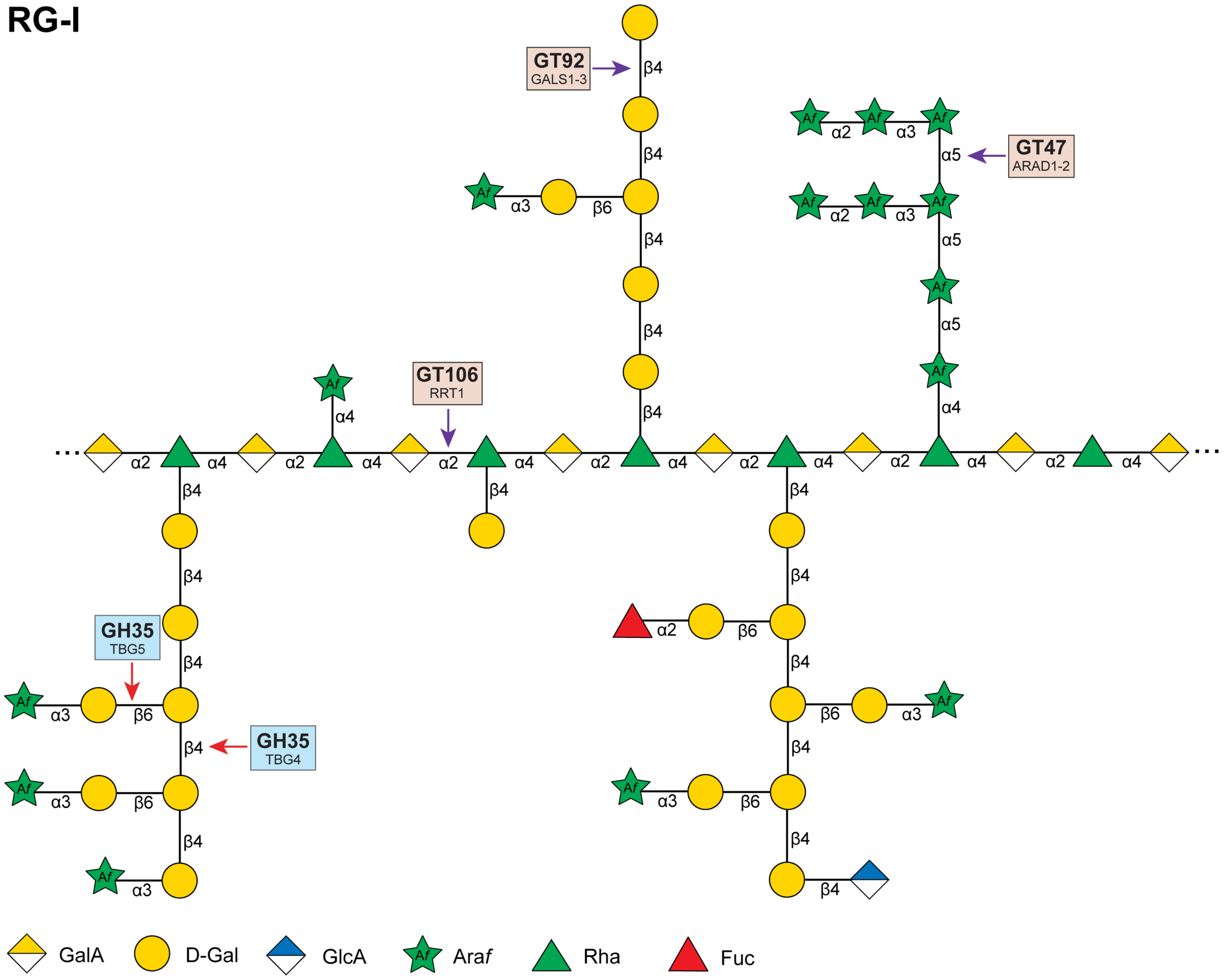


- **RG-I constitutes 20–35% of pectin.**
- GALACTAN SYNTHASE 1 (GALS1) of GT92 has been demonstrated as a β-1,4-galactan:β-1,4-GalT
- ARABINAN DEFICIENT 1 (ARAD1) and its homolog ARAD2 in pectin arabinan biosynthesis

Data S-71

### GT106 + Coleochaete

Takenaka Y, Kato K, Ogawa-Ohnishi M, Tsuruhama K, Kajiura H, Yagyu K, Takeda A, Takeda Y, Kunieda T, Hara-Nishimura I, Kuroha T, Nishitani K, Matsubayashi Y, Ishimizu T. Pectin **RG-I rhamnosyltransferases** represent a novel plant-specific glycosyltransferase family. Nat Plants. 2018 Sep;4(9):669-676. doi: 10.1038/s41477-018-0217-7.

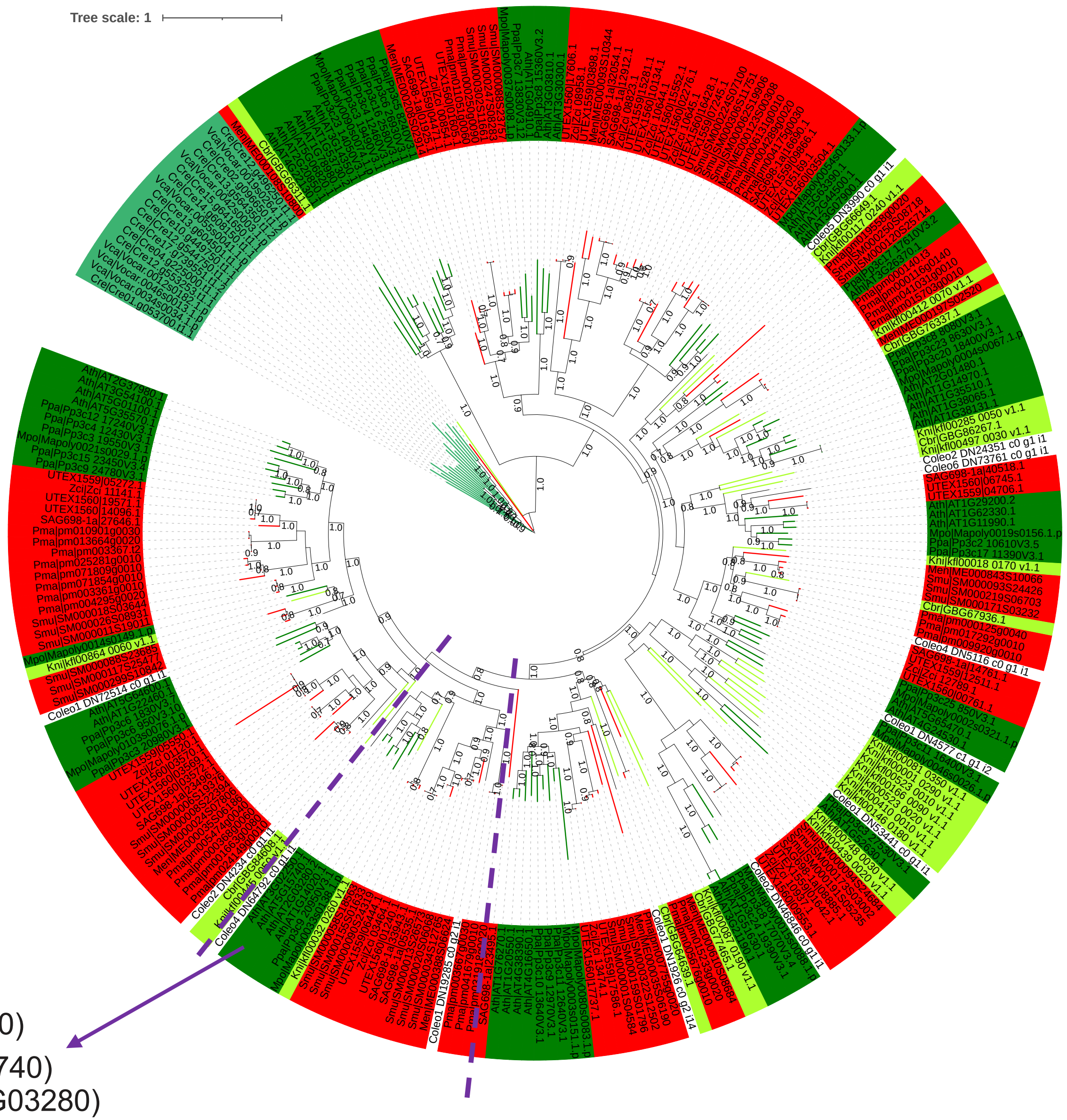

Data S-72

### RG-I: GAT1 (GT116/DUF616)

**Galaturonosyltransferase**

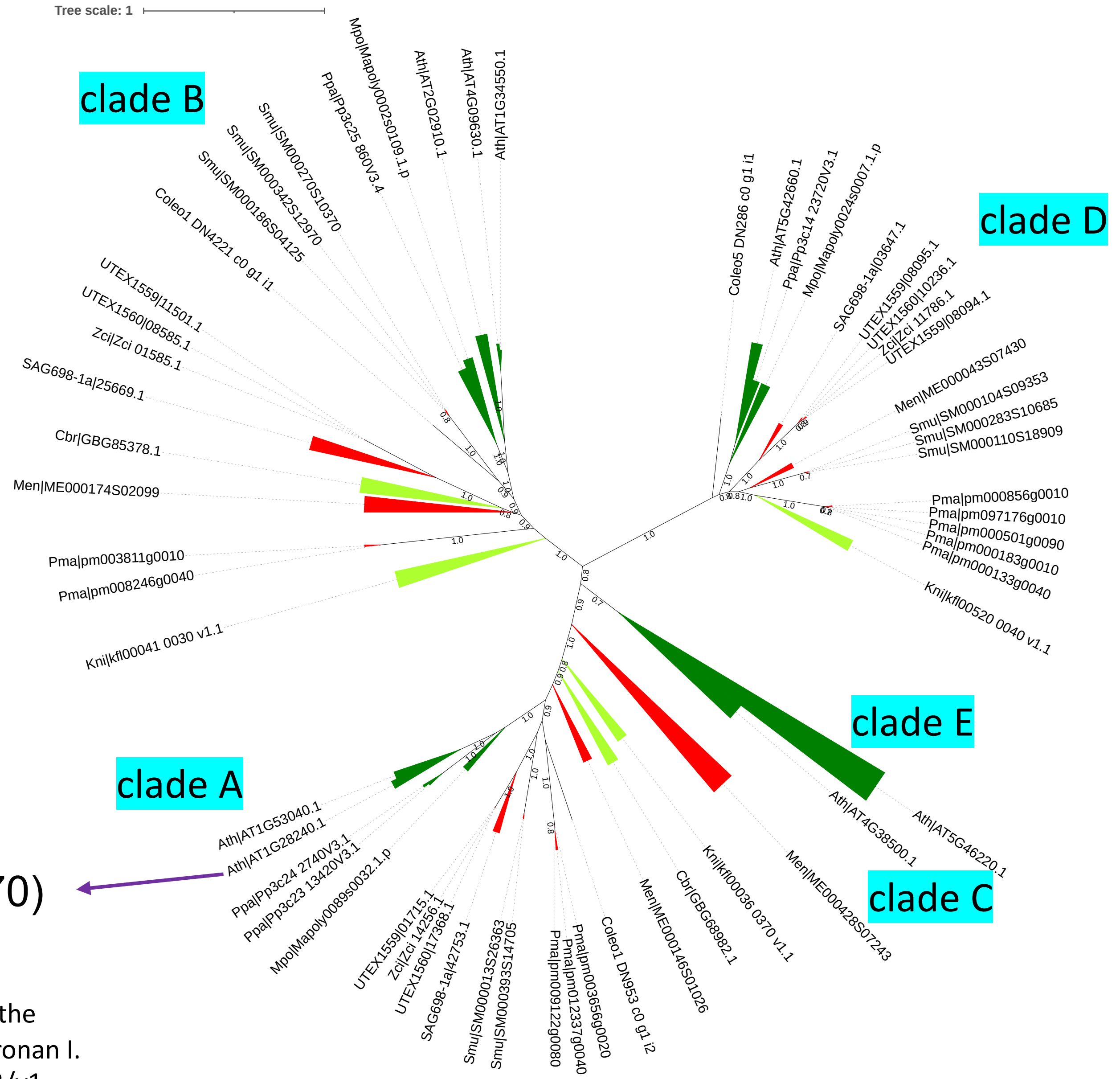


RGGAT1(AT1G28240/MUCI70)

Amos R, Atmodjo M, Huang C, et al. Polymerization of the backbone of the pectic polysaccharide rhamnogalacturonan I. Research Square; 2022. DOI: 10.21203/rs.3.rs-1475173/v1.

Data S-73

### GAT1(AT1G28240/MUCI70)

https://itol.embl.de/tree/13423819014786971667102713

RG-I: GAT1 (DUF616)
**Galaturonosyltransferase**

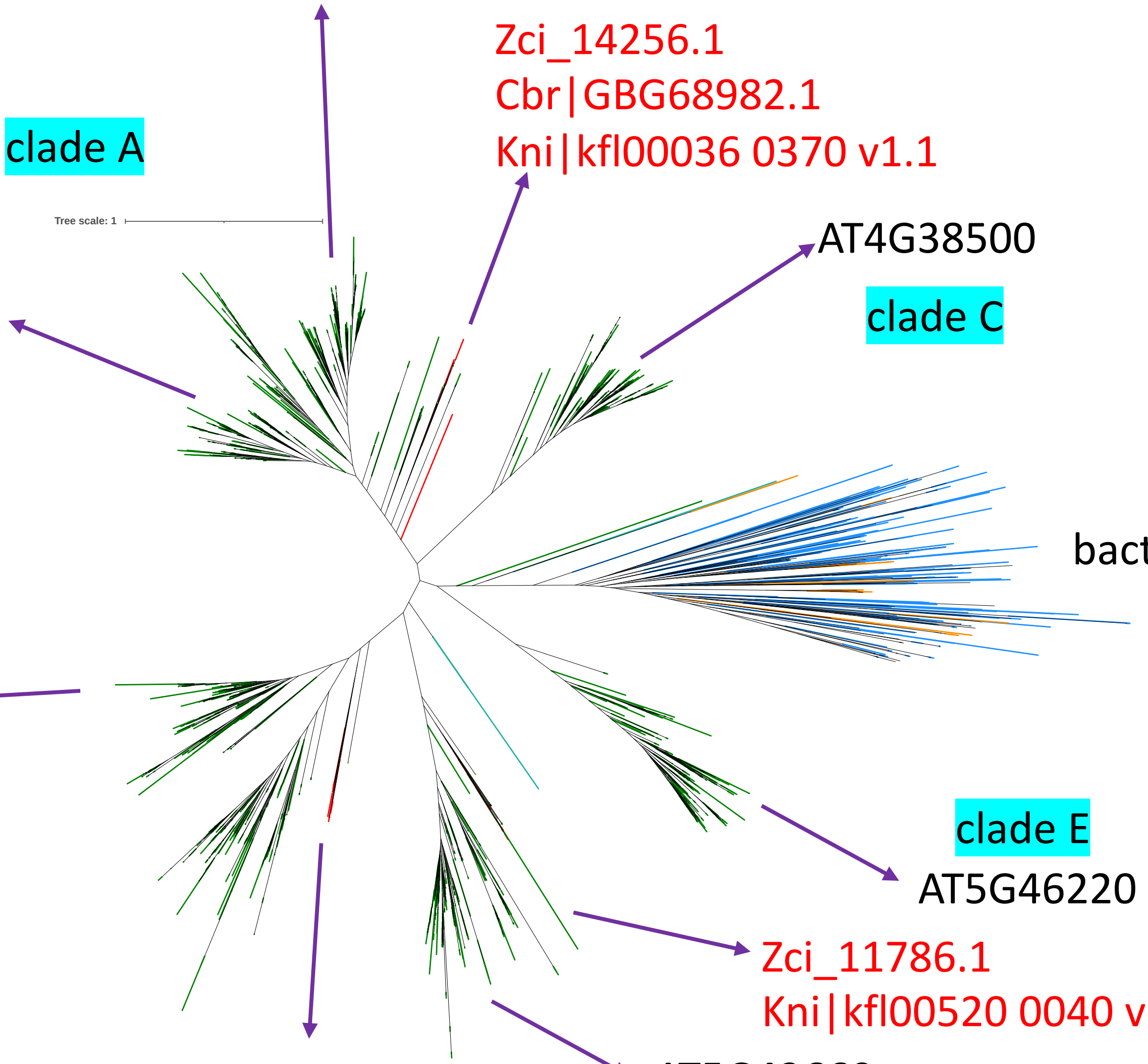


Amos R, Atmodjo M, Huang C, et al. Polymerization of the backbone of the pectic polysaccharide rhamnogalacturonan I. Research Square; 2022. DOI: 10.21203/rs.3.rs-1475173/v1.

Data S-74

https://itol.embl.de/tree/1342381681245654166831464

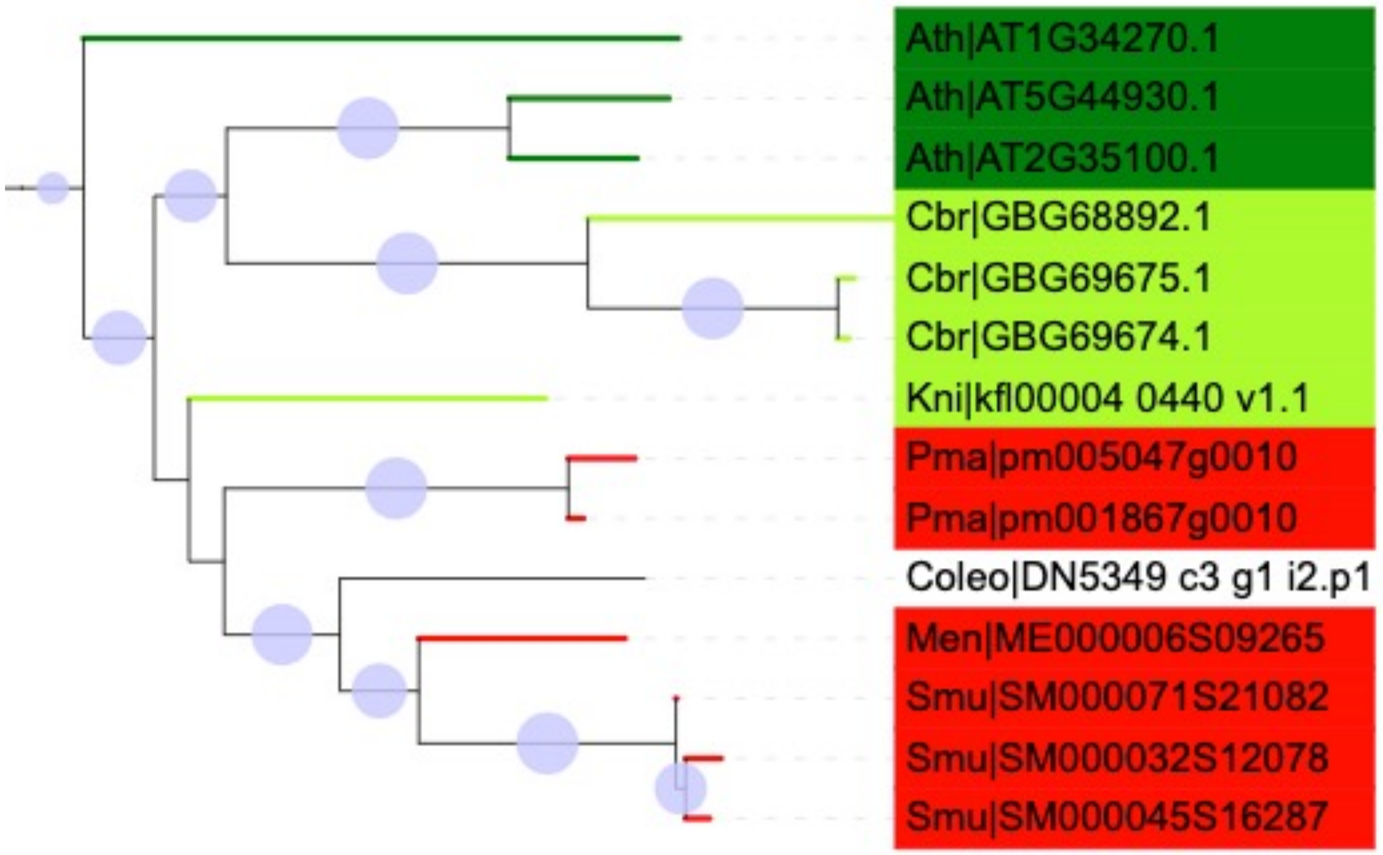


ARAD2 (AT5G44930)
ARAD1 (AT2G35100)

**RG-I**

Harholt et al. ARABINAN DEFICIENT 1 is a putative arabinosyltransferase involved in biosynthesis of pectic arabinan in Arabidopsis. Plant Physiol. 2006 Jan;140(1):49-58. doi: 10.1104/pp.105.072744.

GT47

Data S-75

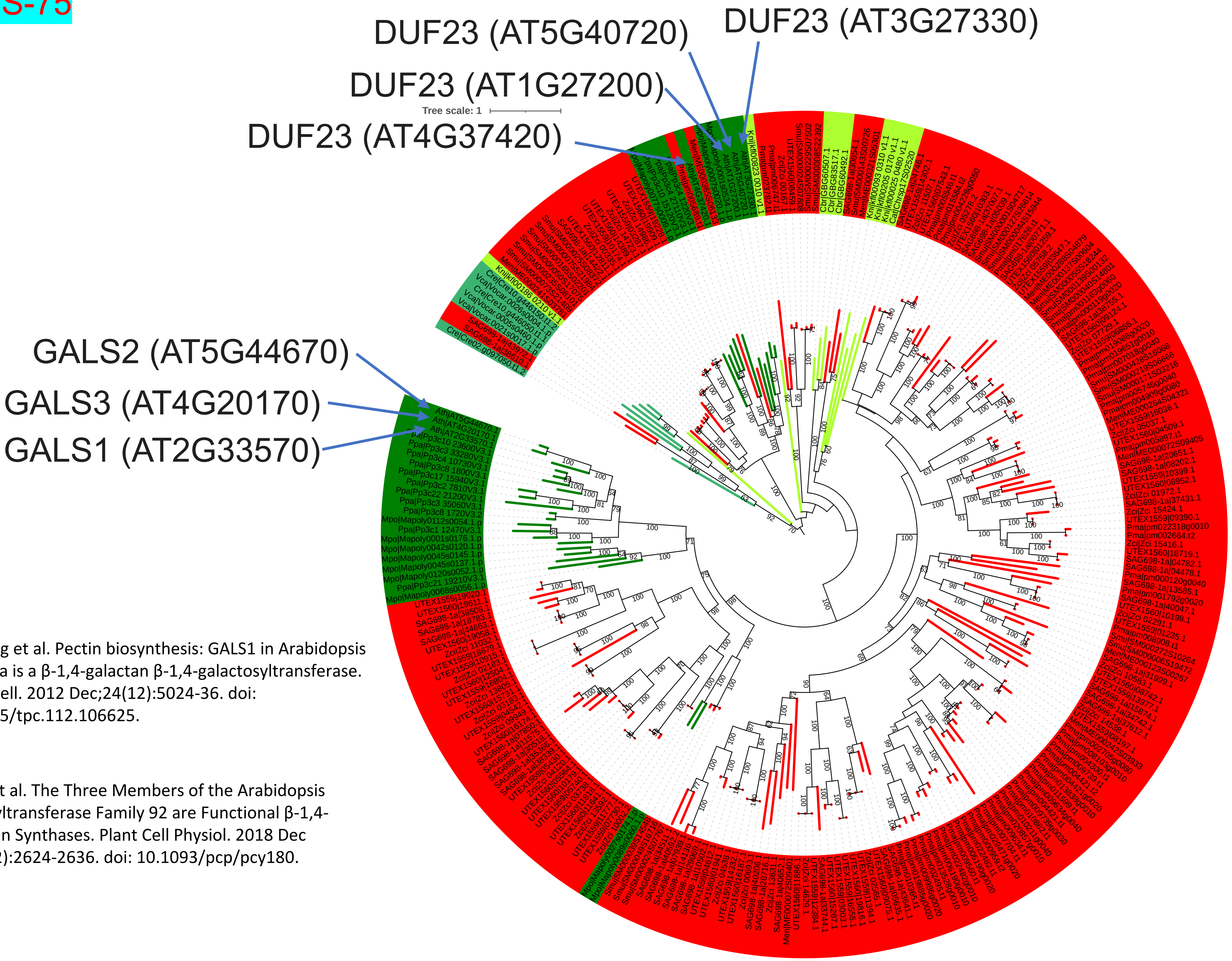


# GT92 (PF01697)

**e-value: e-10**

Liwanag et al. Pectin biosynthesis: GALS1 in Arabidopsis thaliana is a β-1,4-galactan β-1,4-galactosyltransferase. Plant Cell. 2012 Dec;24(12):5024-36. doi: 10.1105/tpc.112.106625.

Ebert et al. The Three Members of the Arabidopsis Glycosyltransferase Family 92 are Functional β-1,4-Galactan Synthases. Plant Cell Physiol. 2018 Dec 1;59(12):2624-2636. doi: 10.1093/pcp/pcy180.

Data S-76

https://itol.embl.de/tree/208127240222183691667018617

#### Polysaccharide Lyase Family 4 / Subf 2
#### rhamnogalacturonan endolyase (EC 4.2.2.23)

Mokshina N, Makshakova O, Nazipova A, Gorshkov O, Gorshkova T. Flax rhamnogalacturonan lyases: phylogeny, differential expression and modeling of protein structure. Physiol Plant. 2019 Oct;167(2):173-187. doi: 10.1111/ppl.12880.

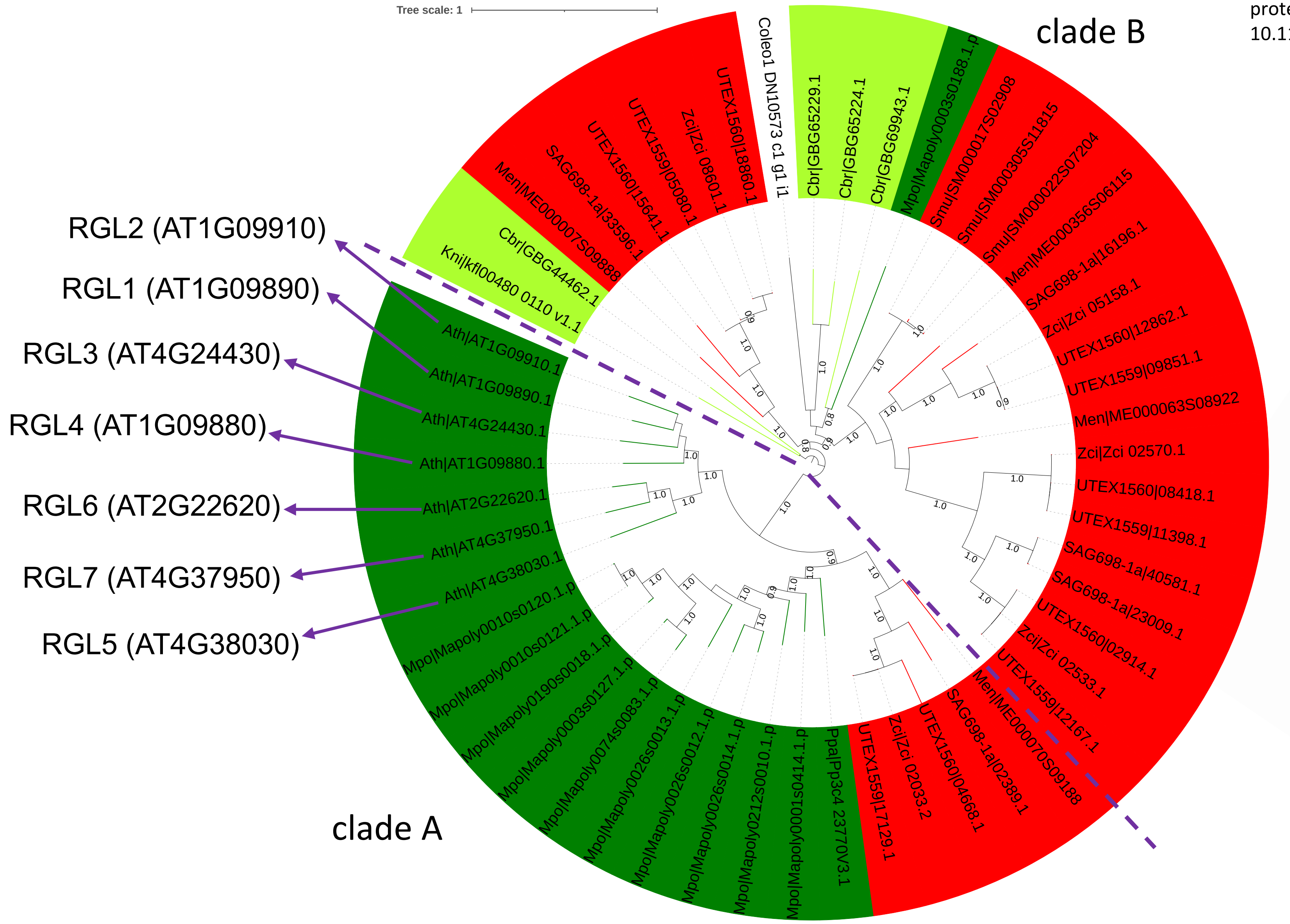


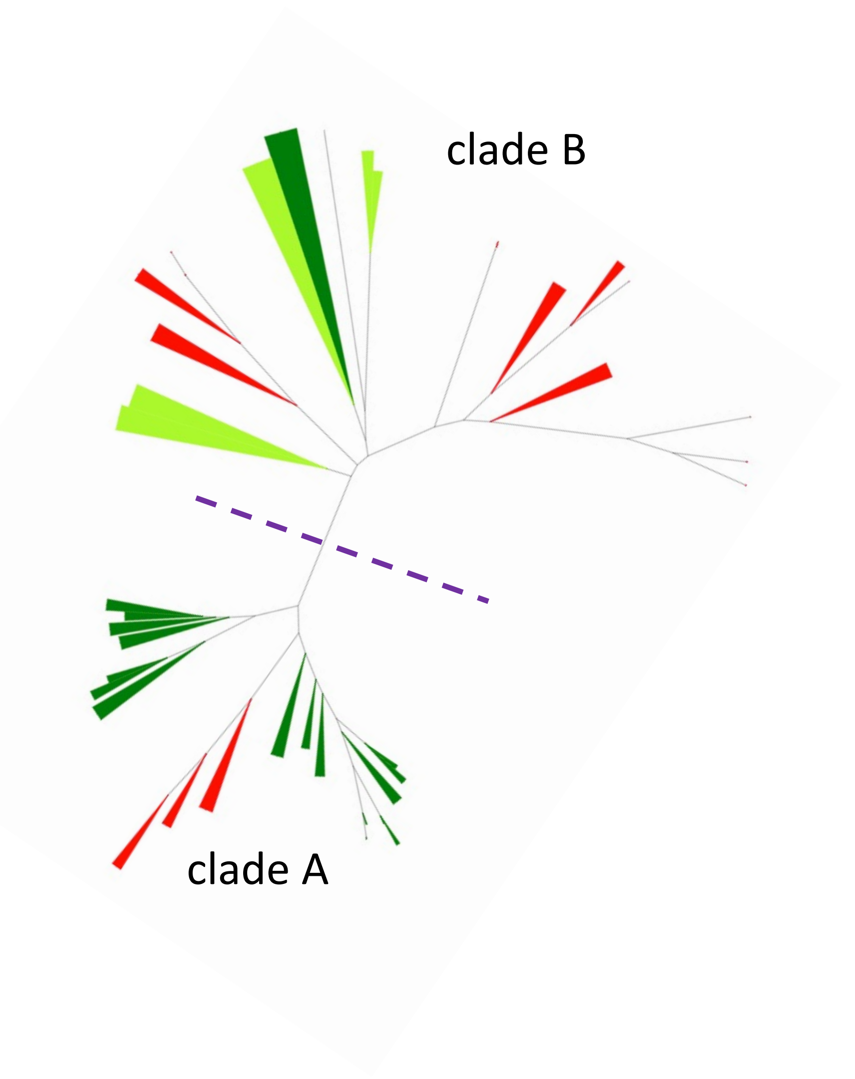

### Data S-77

https://itol.embl.de/tree/2081272402223602516670199130

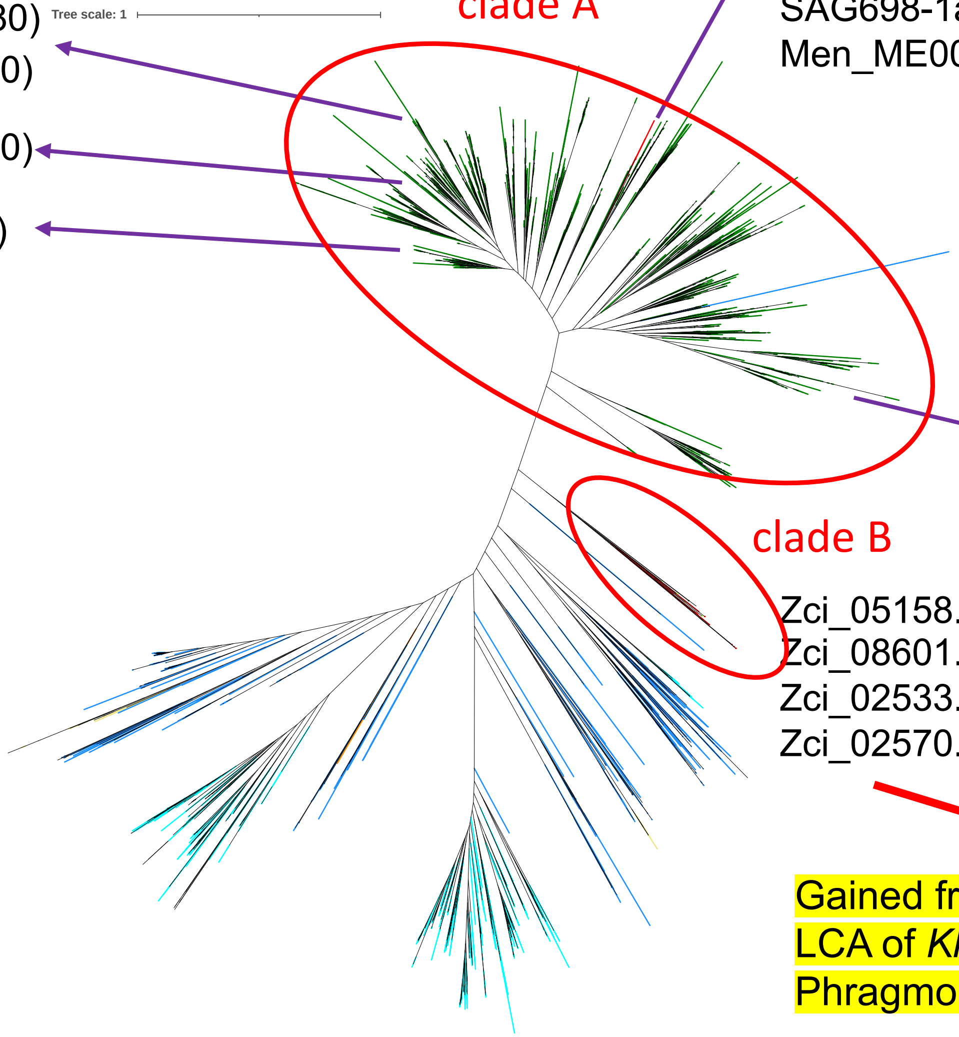


RGLs might have been gained via HGT from bacteria

- clade B is the ancestral clade; emerged in the LCA of Klebsormidiophyceae & Phragmoplastophyta
- clade A first appeared in the LCA of Zygnematophyceae & land plants

Data S-78

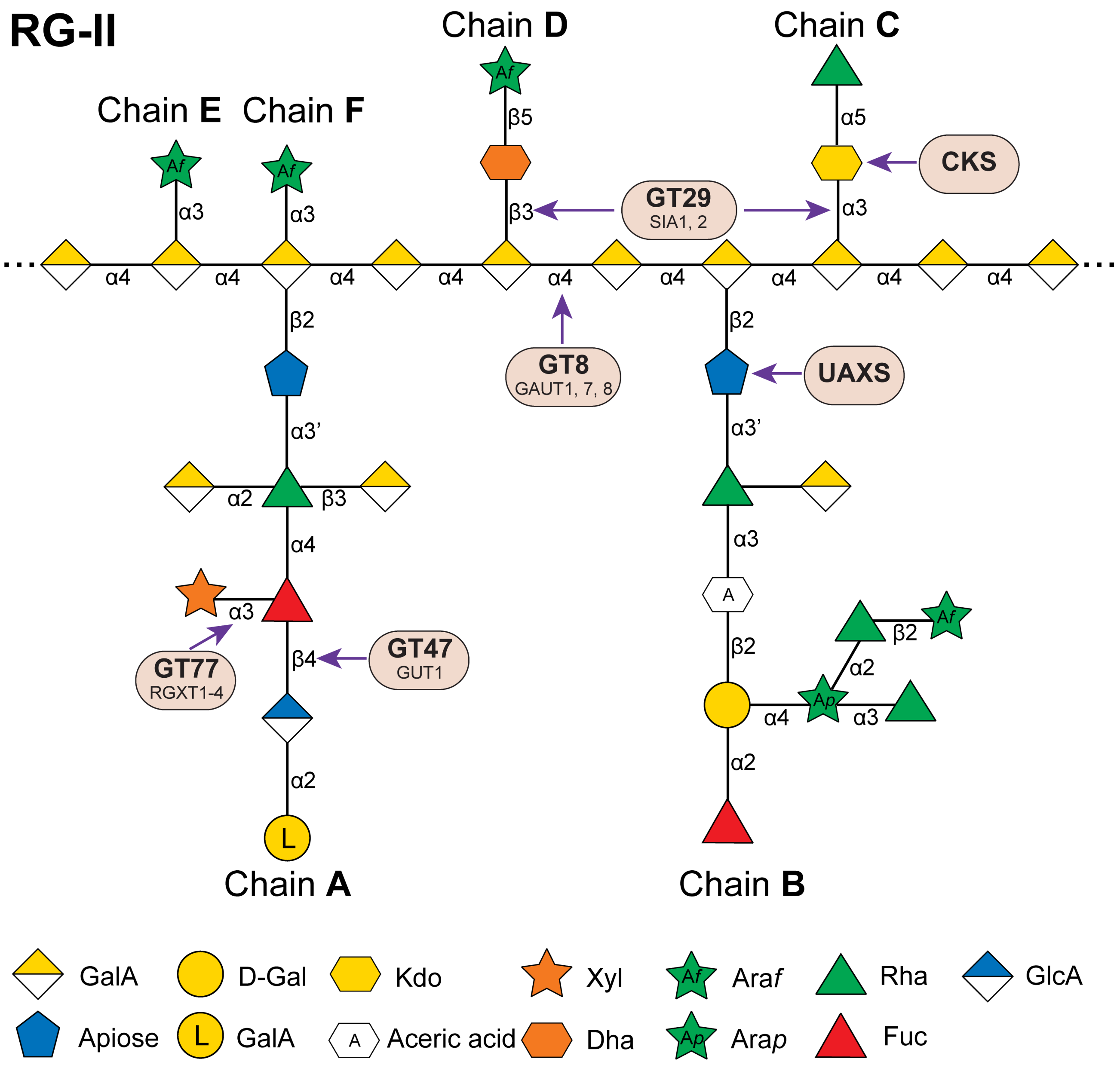


### Rhamnogalacturonan II (RG-II)

- RHAMNOGALACTURONAN XYLOSYL- TRANSFERASES (RGXTs) that have RG- II:α-1,3-xylosyltransferase activity

Gal-Fuc disaccharide structure in sidechain B of RG-II is identical to that found in XyG

terminal non-reducing 2-O-Me-a-L-Fucresidue that is a-(1,2)-linked to D-Gal in sidechain B that is often acetylated

Data S-79

https://itol.embl.de/tree/1342381641712360716651777614

### GT77 + Coleochaete

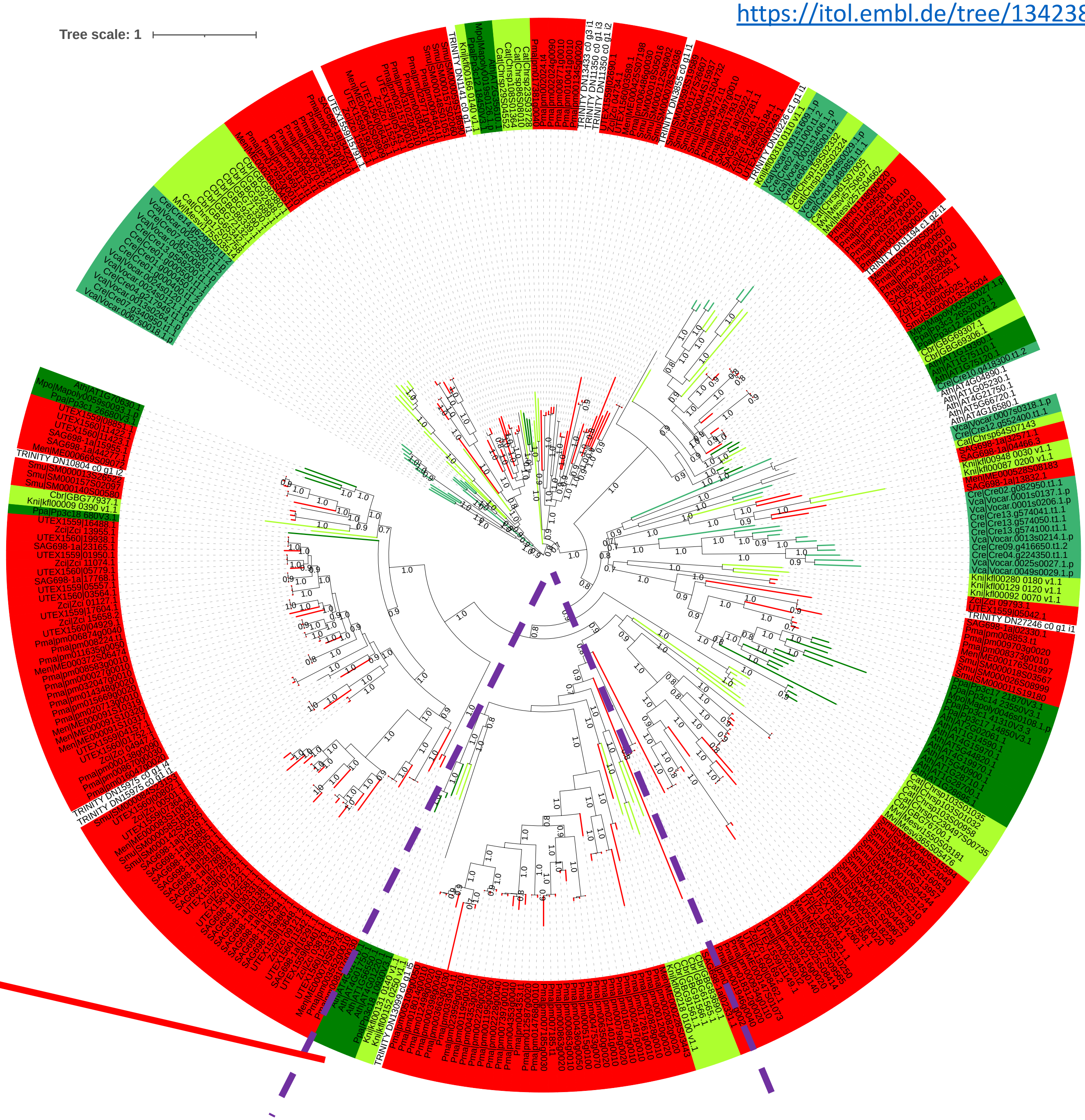


#### α-1,3-xylosyltransferases

**pectic rhamnogalacturonan-II**

Egelund et al. Arabidopsis thaliana RGXT1 and RGXT2 encode Golgi-localized (1,3)-alpha-D-xylosyltransferases involved in the synthesis of pectic rhamnogalacturonan-II. Plant Cell. 2006 Oct;18(10):2593-607. doi: 10.1105/tpc.105.036566.

RGXT3 (At1g56550)

RGXT4/MGP4 (At4g01220)

RGXT2 (At4g01750)

RGXT1 (At4g01770)

Data S-80

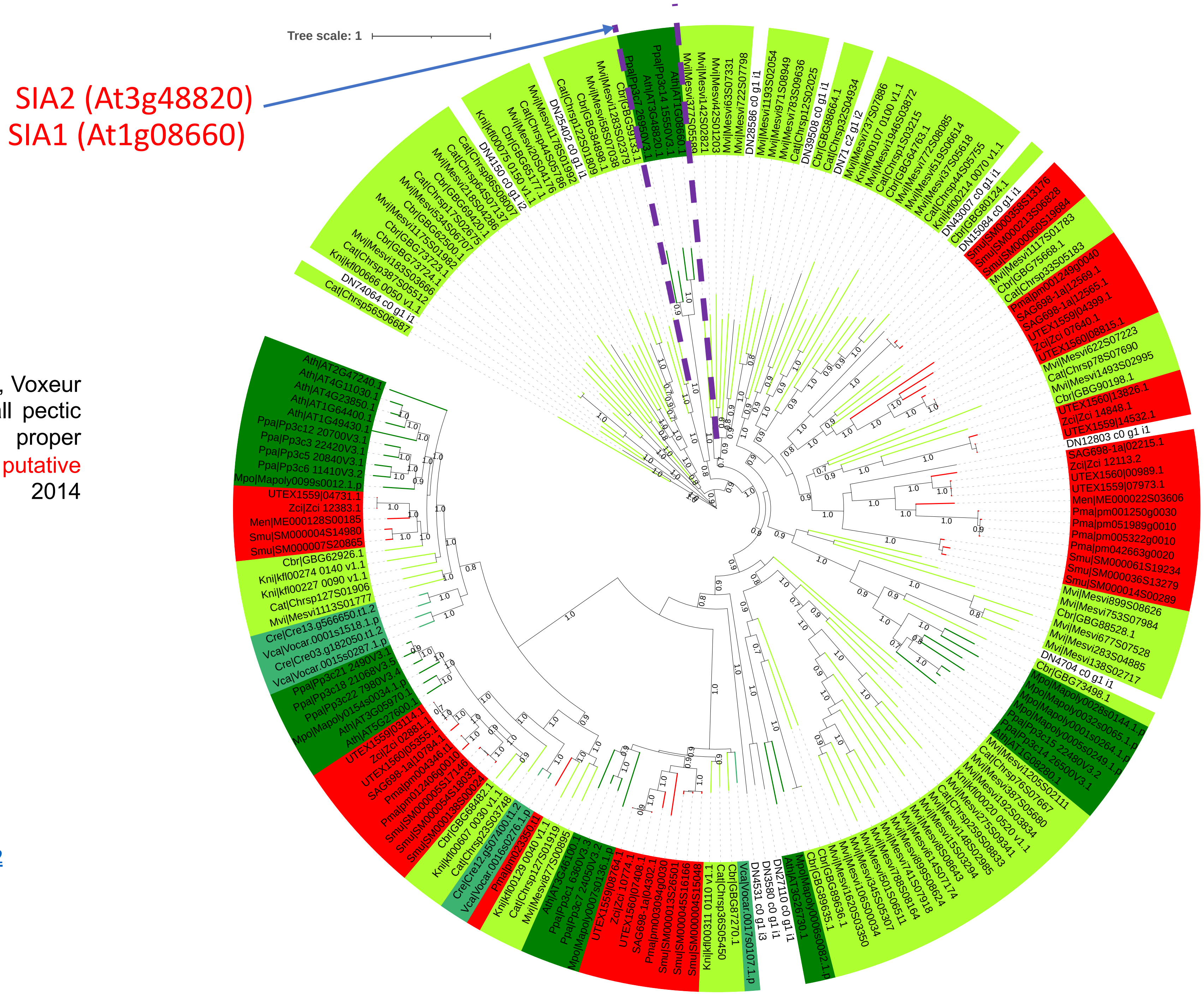


Dumont M, Lehner A, Bouton S, Kiefer-Meyer MC, Voxeur A, Pelloux J, Lerouge P, Mollet JC. The cell wall pectic polymer rhamnogalacturonan-II is required for proper pollen tube elongation: implications of a putative sialyltransferase-like protein. Ann Bot. 2014 Oct;114(6):1177-88. doi: 10.1093/aob/mcu093.

### GT29 + Coleochaete

https://itol.embl.de/tree/20812783235334831665116092

Data S-81

# GH28

GH28.hmm (dbcan) from 16 genomes and Coleochaete with e-value < 1e-10

Rodríguez-Gacio Mdel C, Nicolás C, Matilla AJ. Cloning and analysis of a cDNA encoding an endo-polygalacturonase expressed during the desiccation period of the silique-valves of turnip-tops (Brassica rapa L. cv. Rapa). J Plant Physiol. 2004 Feb;161(2):219-27. doi: 10.1078/0176-1617-01153.

Gallego-Giraldo, L., Liu, C., Pose-Albacete, S., Pattathil, S., Peralta, A.G., Young, J. et al. (2020) ARABIDOPSIS DEHISCENCE ZONE POLYGALACTURONASE1 (ADPG1) releases latent defense signals in stems with reduced lignin content. Proceedings of the National Academy of Sciences of the United States of America, 117, 3281–3290

ADPG1 (At3g57510)

Torki, M., Mandaron, P., Mache, R., and Falconet, D. (2000). Characterization of a ubiquitous expressed gene family encoding polygalacturonase in Arabidopsis thaliana. Gene 242, 427–436. doi: 10.1016/S0378-1119(99)00497-7

PGF8 (Pectin lyase-like superfamily protein, AT3G57790)

E-I

Zci_08289

AT1G19170(PGF1)
AT3G42950(PGF6)

E-V

clade E

Zci_09024

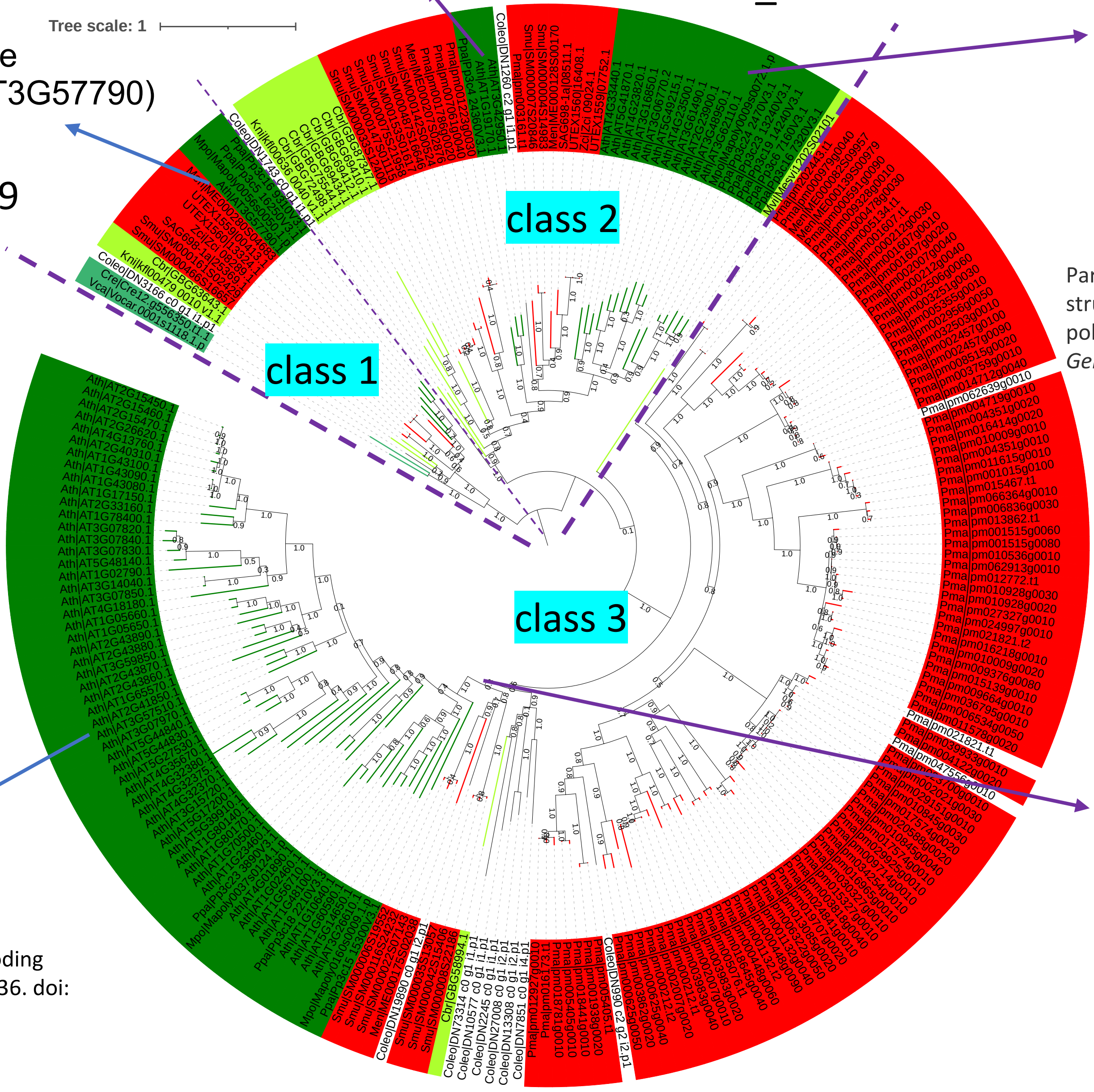


AT4G33440(PGF14) AT5G41870(PGF15)
AT4G23820(PGF13)
AT3G16850(PGF5)
AT3G06770(PGF4)
AT5G49215(PGF16)
AT4G23500(PGF12)
AT3G61490(PGF9)
AT2G23900(PGF3)
AT3G48950(PGF7)
AT3G62110(PGF10)

PMID: 30154820

Park, KC., Kwon, SJ. & Kim, NS. Intron loss mediated structural dynamics and functional differentiation of the polygalacturonase gene family in land plants. *Genes Genom* **32**, 570–577 (2010).

**67 Ath IDs**

AT4G01890
AT1G02460
AT1G56710(PGL1)
AT1G48100(PGX3)
AT1G10640
AT1G60590
AT5G14650(PGLR)
AT3G26610(PGX1)
AT1G23460
AT1G70500
AT1G80170
AT1G80140
AT5G39910
AT5G17200
AT3G15720
AT4G32370

https://itol.embl.de/tree/1342381641712832116730287l5

### Data S-82a

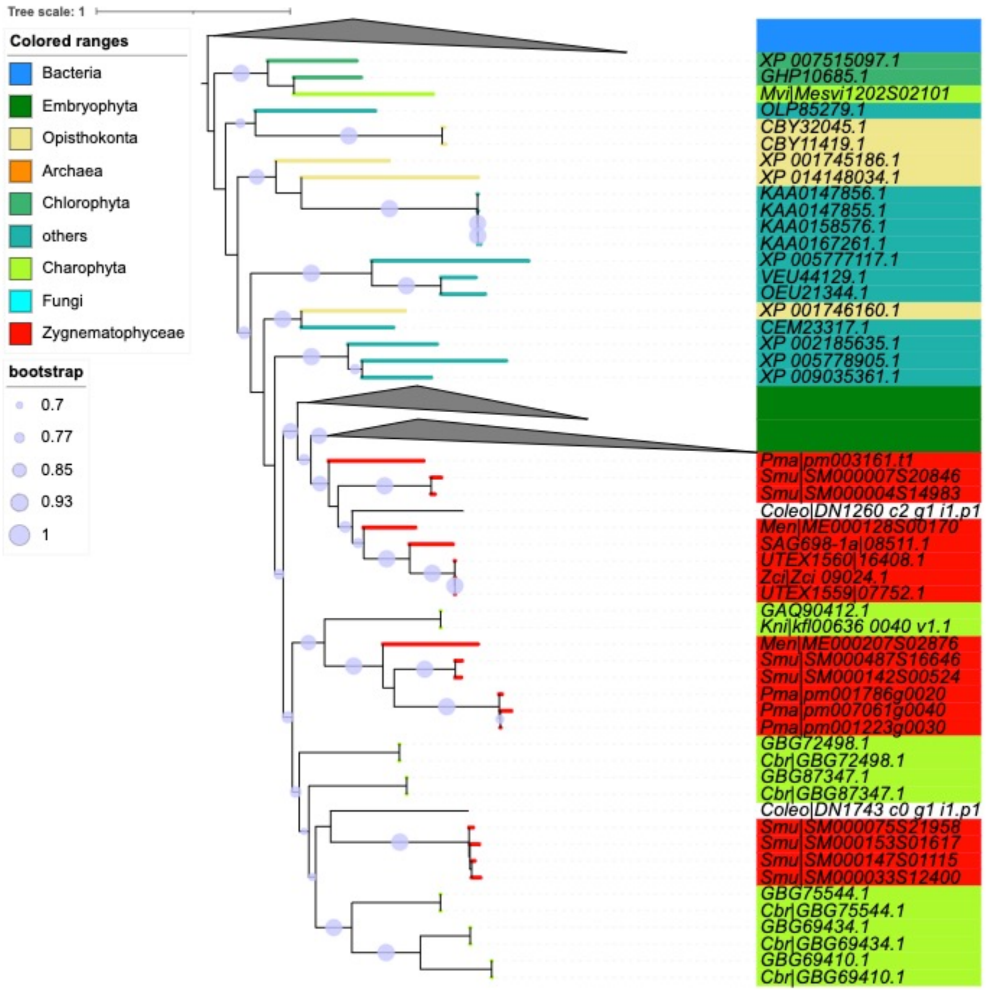


class 2

Zci_09024

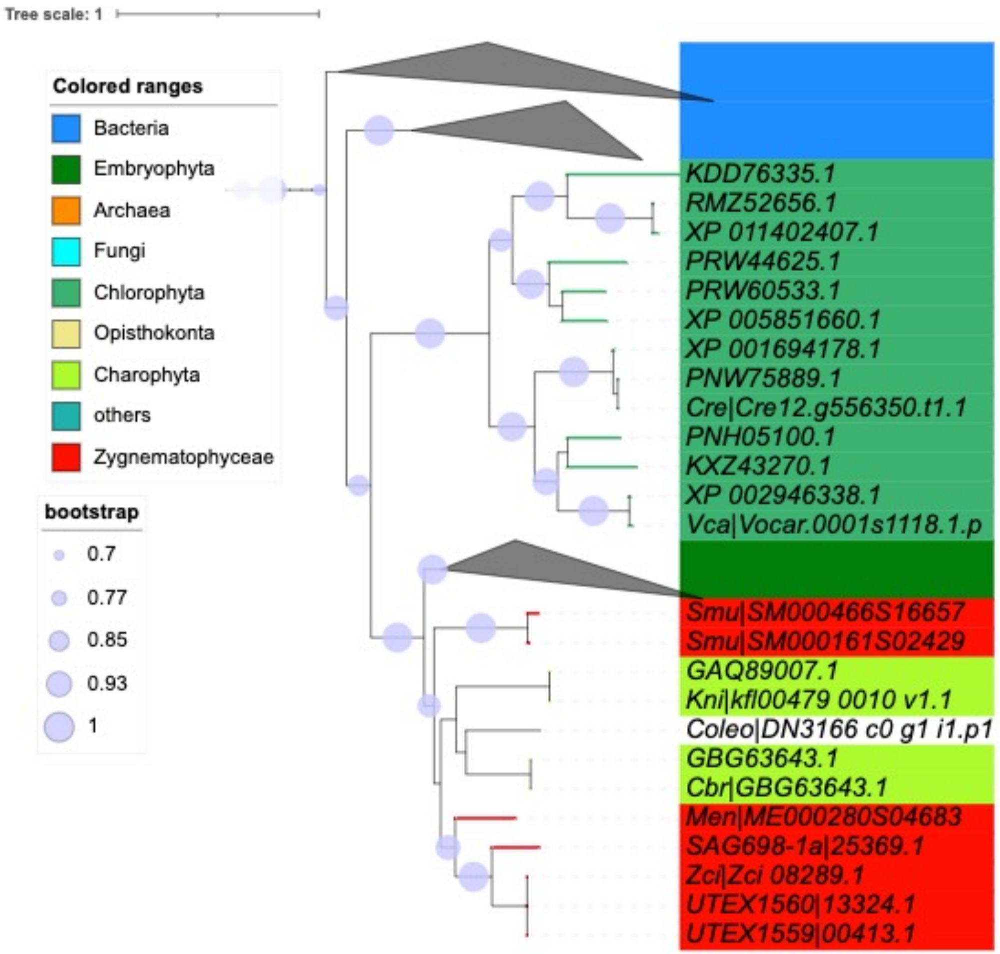


class 1

Zci_08289

Data S-82b

https://itol.embl.de/tree/20812770255475741673199623

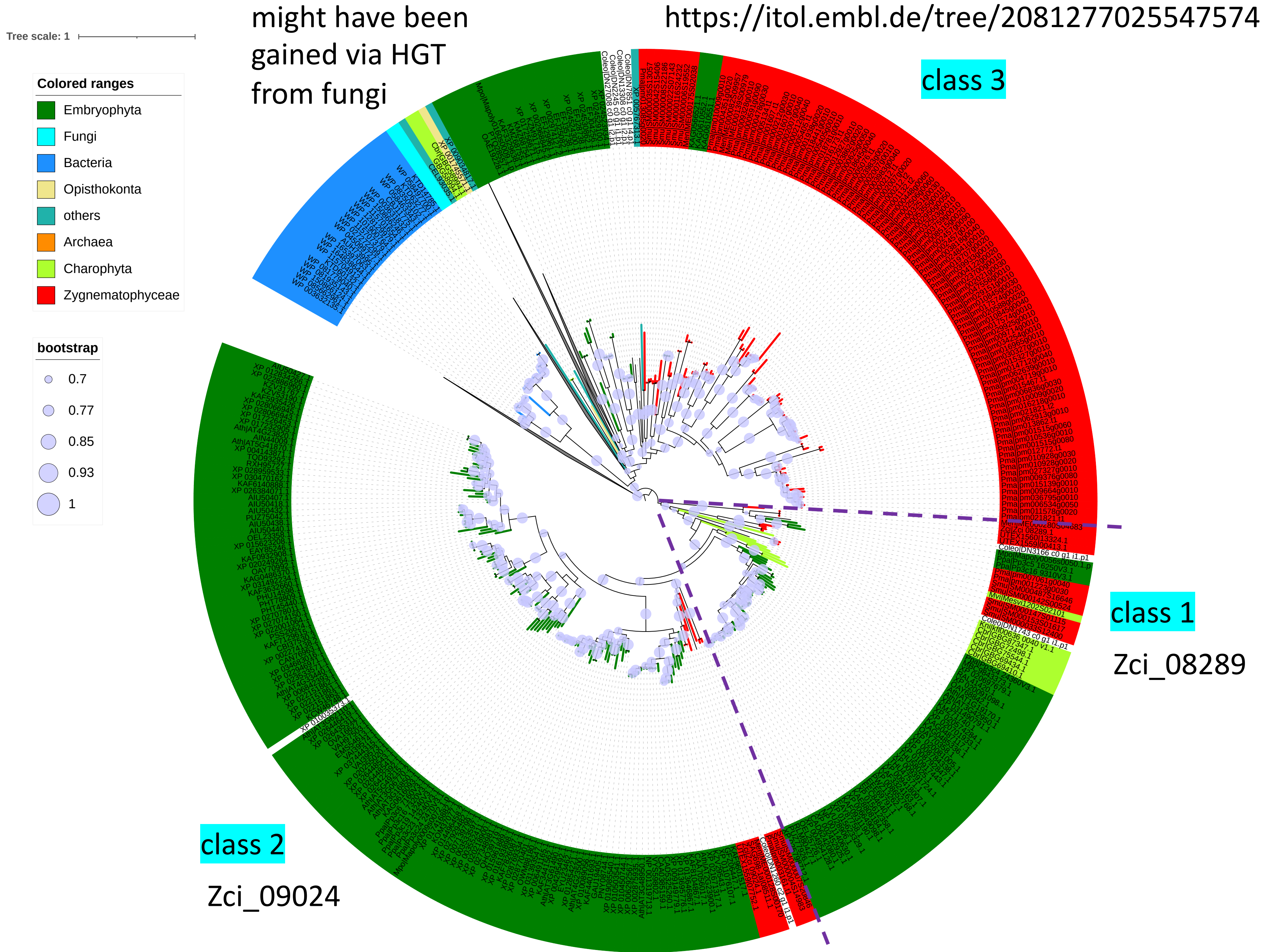

Data S-83a

http://www.cazy.org/e1.html

http://www.cazy.org/PL1_12_characterized.html

| | | |
|---|---|---|
| At1g04680 | PL1 | AAB80622.1 |
| At1g11920 | PL1 | AAC17625.1 |
| At1g14420 | PL1 | AAF43942.1 |
| At1g30350 | PL1 | AAG51103.1 |
| At1g67750 | PL1 | AAL58893.1 |
| At2g02720 | PL1 | AAC05350.1 |
| At3g01270 | PL1 | AAF03499.1 |
| At3g07010 | PL1 | AAF27005.1 |
| At3g09540 | PL1_12 | AAF23285.1 |
| At3g55140 | PL1_12 | CAB75748.1 |
| At3g24230 | PL1 | BAB01365.1 |
| At3g24670 | PL1 | BAB01216.1 |
| At3g27400 | PL1 | NP_189376.1 |
| At3g53190 | PL1 | BAD95042.1 |
| At3g54920 (PMR6) | PL1 | AAM97687.1 |
| At4g13210 | PL1 | CAB41931.1 |
| At4g13710 | PL1 | AAL11586.1 |
| At4g22080 | PL1 | CAA18111.1 |
| At4g22090 | PL1 | CAA18112.1 |
| At4g24780 (A10) | PL1 | AAK25850.1 |
| At5g04310 | PL1 | NP_196051.2 |
| At5g09280 (fragment) | PL1 | CAC05454.1 |
| At5g15110 | PL1 | CAC01830.1 |
| At5g48900 | PL1 | AAK92730.1 |
| At5g55720 | PL1 | BAB09239.1 |
| At5g63180 | PL1 | AAL25610.1 |

Data S-83b

https://itol.embl.de/tree/2081278868212161668015296

# PL1

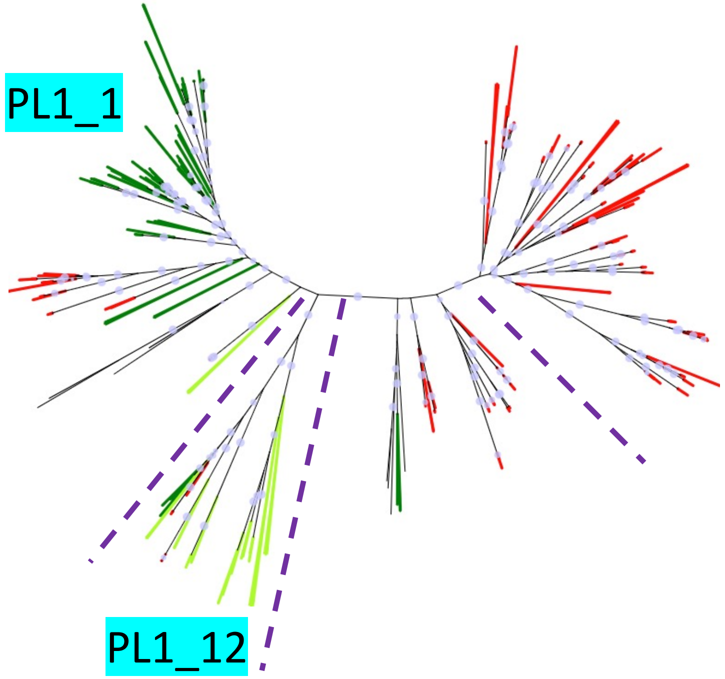


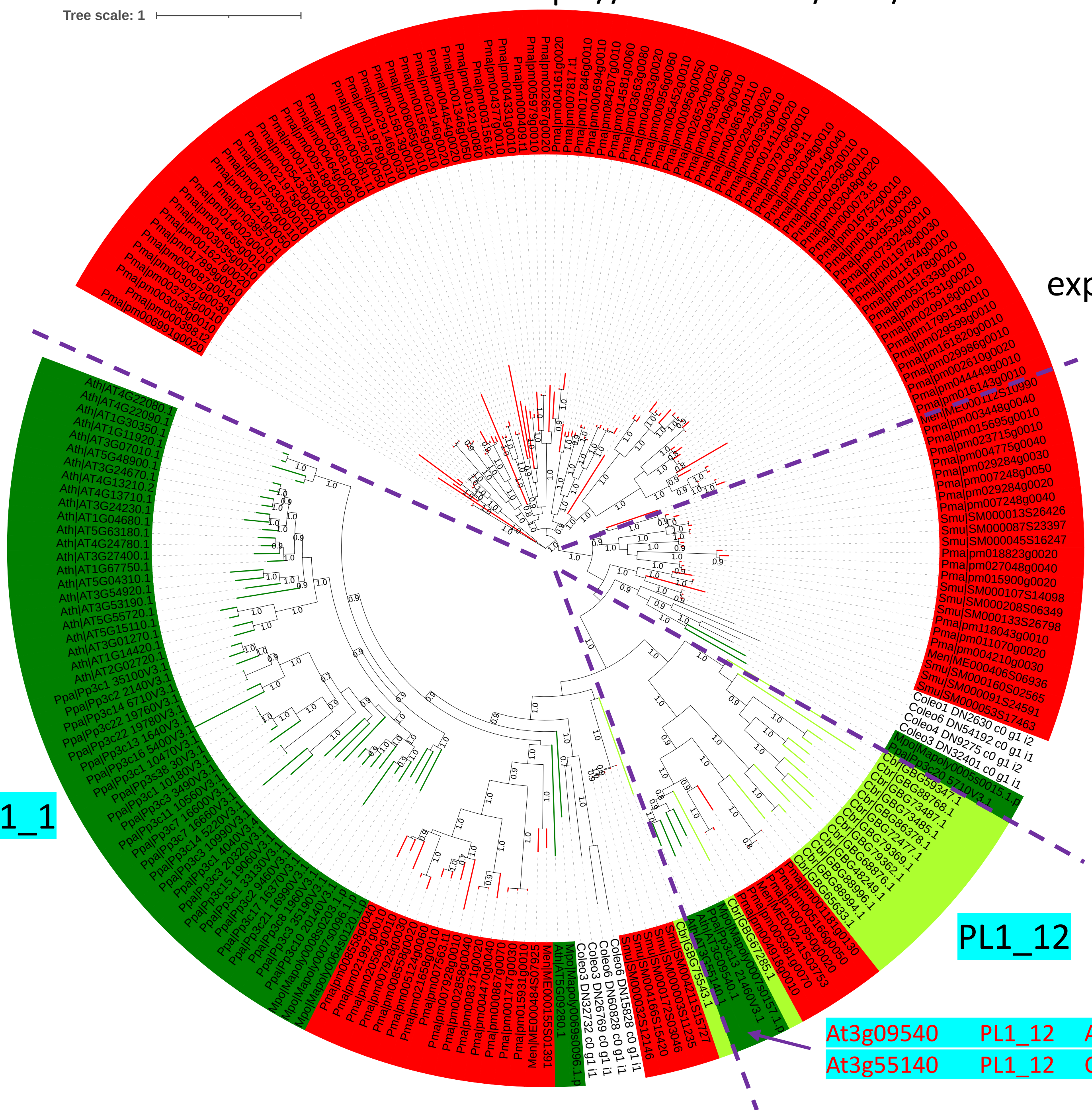


| | | |
|---|---|---|
| At3g09540 | PL1_12 | AAF23285.1 |
| At3g55140 | PL1_12 | CAB75748.1 |

### Data S-84a

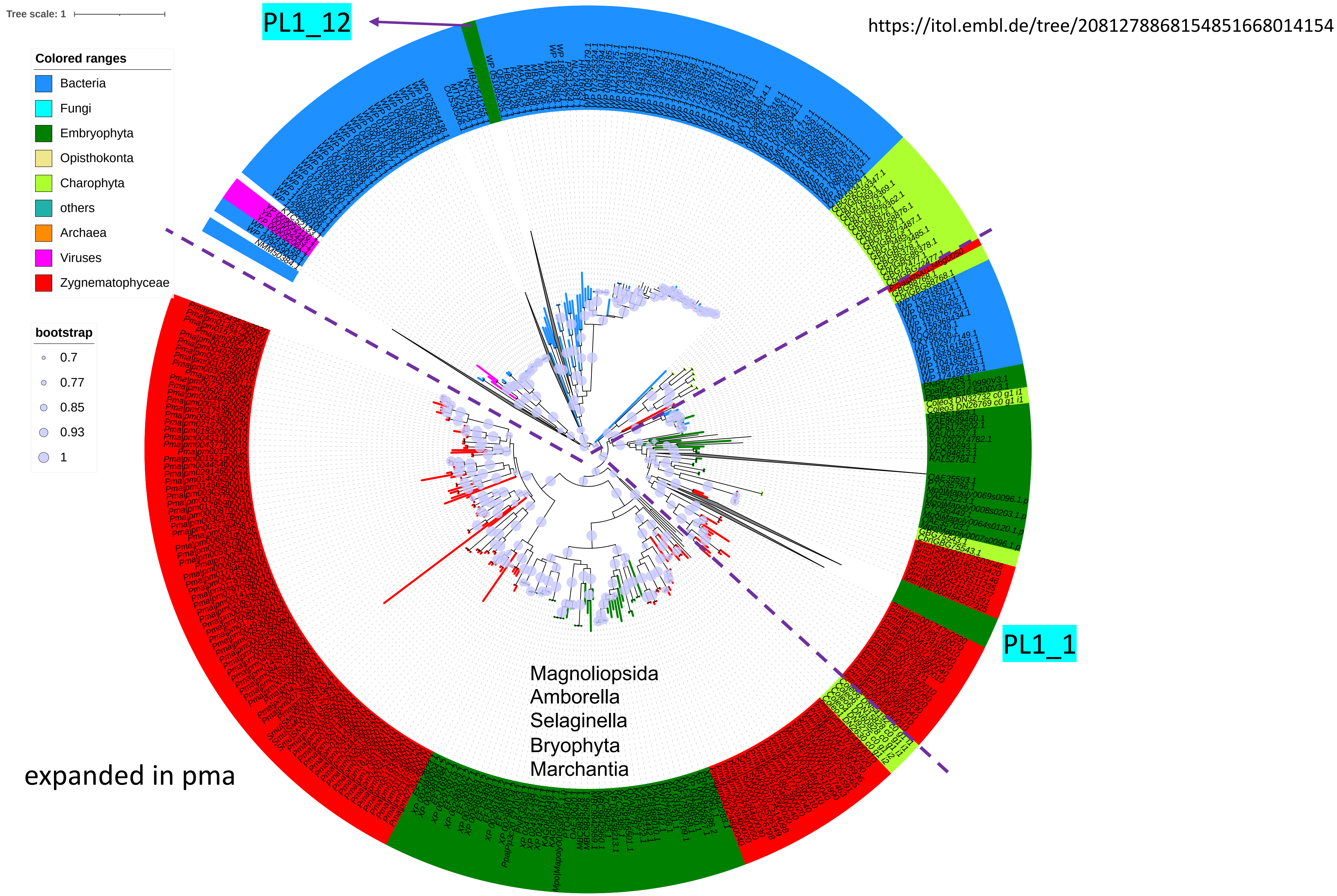

Tree scale: 1
PL1_12
https://itol.embl.de/tree/2081278868154851668014154
Colored ranges
Bacteria
Fungi
Embryophyta
Opisthokonta
Charophyta
others
Archaea
Viruses
Zygnematophyceae
bootstrap
0.7
0.77
0.85
0.93
1
PL1_1
Magnoliopsida
Amborella
Selaginella
Bryophyta
Marchantia
expanded in pma

Data S-84b

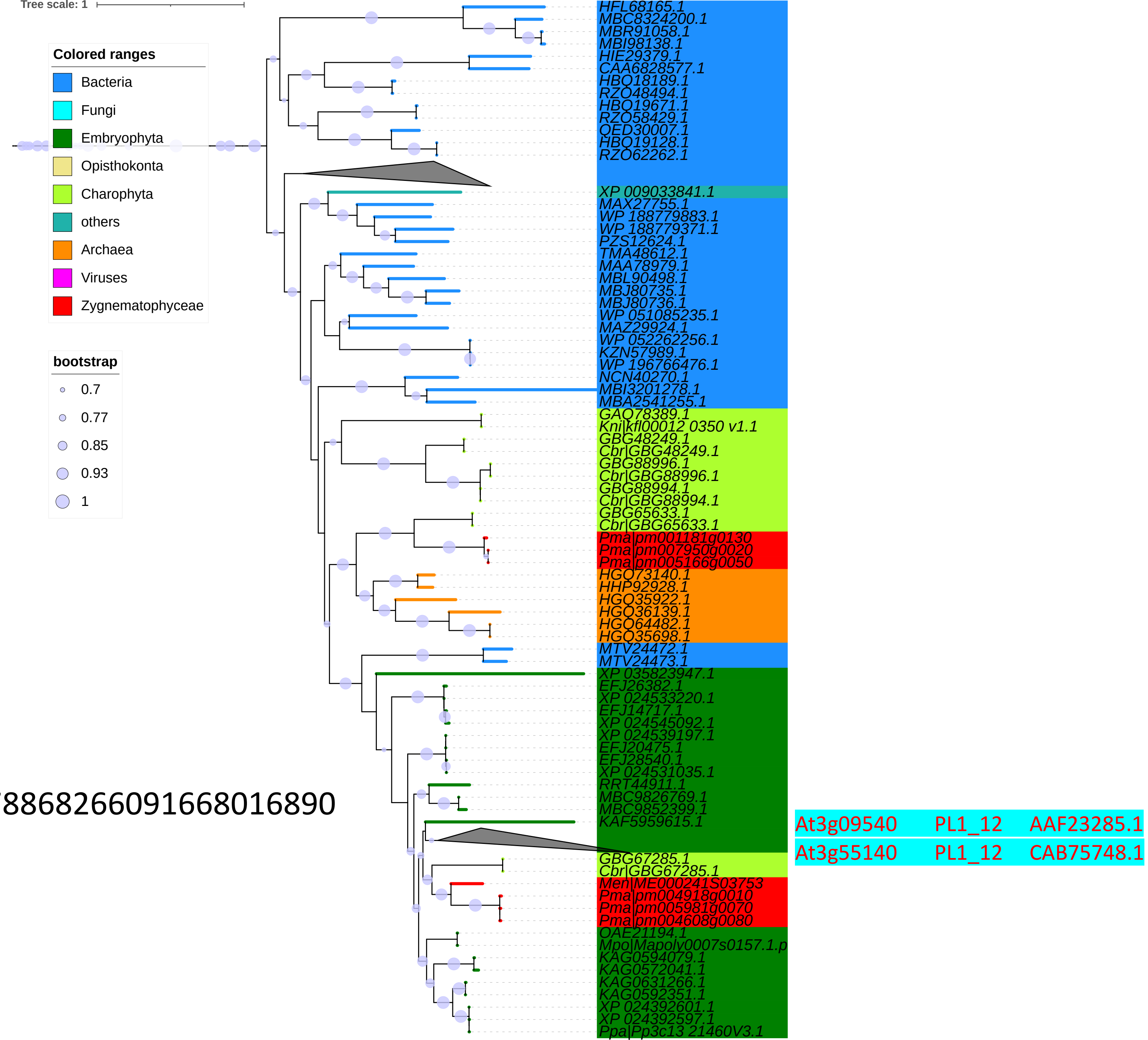


https://itol.embl.de/tree/2081278868266091668016890
