## Supplemental Text and Figures for "Chromosome-level genomes of multicellular algal sisters to land plants illuminate signaling network evolution"

**SUPPLEMENTARY MATERIAL**

**Data S1**. CAZyme gene family analysis.
**Supplementary Text 1**. History of *Zygnema* strains
**Supplementary Text 2**. Supplemental text on cell wall complexity and CAZymes

**Table S1**. Data on genome sequencing, annotation, divergence times, *Closterium* sex hormone homologs and cell wall-related enzymes. (A) Genome and transcriptome sequence Data, (B) Genome size estimation, (C) Summary of genome assembly, (D) Completeness, evaluation of four *Zygnema* genomes, (E) BUSCO evaluation of other genomes, (F) Divergence time estimation of the representative genomes, (G) Gene annotation of SAG 698-1a mitogenome, (H) Comparison of introns and exons between mitogenomes of SAG 698-1a and SAG 698-1b, (I) Summary of gene content in 16 genomes, (J) Repeat elements annotation, (K) *Closterium* sex hormone protein search, (L) Cell wall-related enzyme subfamilies.
**Table S2**. Domain combinations in the Zygnematophyceae and Embryophyta ancestor.
**Table S3**. (A) Orthogroup expansions and contractions (full CAFE results), (B) Significant orthogroup expansions and contractions for the ancestors of Embrophyta + Zygematophyceae, Zygnematophyceae, and *Zygnema*, (C) Results of TAPscan, (D) Identification of cell-division genes in *Zygnema* and across the green lineage.

**Figure S1**. Light micrographs of *Zygnema circumcarinatum* SAG 698-1b fixed and stained with acetocarmine.
**Figure S2**. K-mer plots for genome size estimation.
**Figure S3.** Gene annotation of *Zygnema circumcarinatum* SAG 698-1b mitogenome
**Figure S4**. Mauve alignment of mitogenomes of UTEX 1559 and SAG 698-1b.
**Figure S5**. Gene annotation of *Zygnema* cf. *cylindricum* SAG 698-1a mitogenome.
**Figure S6**. Polyploidy analysis of SAG 698-1b chromosome-level genome and comparison with *Physcomitrium patens*.
**Figure S7**. Maximum likelihood trees inferred from orthogroups obtained from Zygnematophyceae and *Zygnema* genomes.
**Figure S8**. Comparisons of the three *Z. circumcarinatum* chromosome-level genomes.
**Figure S9**. Phylogeny of the O-FucT family.
**Figure S10**. Phylogenetic tree for CCD proteins.
**Figure S11**. Phylogeny of CCD7 homologs.
**Figure S12**. Phylogeny of CCD8 homologs.
**Figure S13.** Identification of ABA in SAG 698-1b.
**Figure S14**. Phylogenies of genes salient to the production of phenylpropanoid-derived specialized metabolites.
**Figure S15**. Maximum Likelihood phylogeny of phytochrome homologs.
**Figure S16**. Maximum Likelihood phylogeny of MADS-domain proteins.

**Supplemental Text 1: History of *Zygnema* strains**

A *Zygnema* sample was collected by V. Czurda in 1929 from a ditch at meadow Poselteich (Poselský rybnik) near Hirschberg (Doksy), Bohemia. From the original samples, a mating type (mt) + and mt – were created, which gave rise to Pringsheim's Prague strains 208 and 209, respectively. Pringsheim's strain 208 (mt +) was then transferred to the Culture Collection of Algae and Protozoa (CCAP), Scotlan under CCAP698/1A and from there to the Culture Collection of Algae at Göttingen University (SAG) under accession SAG 698-1a and to the Culture Collection of Algae at UT-Austin (UTEX) under accession UTEX 42 (=former IUCC 42). A "spontaneous mutant with increased size and numbers of chloroplasts per cell" in UTEX 42 was described by Gauch (1966), deposited under accession UTEX 1559. Pringsheim's strain 209 (mt -) was deposited as CCAP 689/1B and from there into SAG 698-1b and UTEX 43. Again, Gauch (1996) identified a "spontaneous mutant of UTEX 43 with increased size and numbers of chloroplasts per cell", which was deposited under accession UTEX 1560. Therefore, all four *Zygnema* isolated should derive from the same field sample obtained by V. Czurda in 1929. This contrast with our findings that SAG 698-1a is in fact *Zygnema* cf. *cylindricum*, which points to an event of contamination or mislabeling in algal collections at some point in the last ~100 years. The known strain history, however, explains the closer affinity of Z. circumcarinatum UTEX 1559 and SAG 698-1b compared to UTEX 1560.

**Supplemental Text 2: Cell wall complexity and innovation**

The earliest land plants had to overcome stressors such as drastic temperature change, UV, high light, and pathogens not faced in aquatic habitats (Furst-Jansen et al., 2020). To adapt to the land environment, the common ancestors of Streptophyta must have made some major molecular innovations in their cell wall structures (Bowles et al., 2022). Cell walls are the first layer of natural barriers that protect plant cells from various environmental stresses, which differ significantly between aquatic and land environments. Cell walls also can constantly restructure during active cell growth, cell division, and in response to biotic or abiotic stresses. Streptophyta primary cell walls consist of polysaccharides: celluloses are tethered with various hemicelluloses (xylans, xyloglucans, mannans, and beta-glucans) and then embedded within the matrix of highly complex pectic polysaccharides, including homogalacturonan (HG), rhamnogalacturonan I (RG-I), rhamnogalacturonan II (RG-II), and arabinogalactan proteins (AGP) (Sorensen et al., 2011). Celluloses are linear but most hemicelluloses and pectins have very complex sidechains. Besides being the main component of cell walls, some of these polysaccharides can be secreted to the outside of cells to from extracellular matrix and mucilage, critical for the algae to form biofilms, trap water, prevent desiccation, and interact with soil microbes (Domozych and Bagdan, 2022; Herburger et al., 2022).

Carbohydrate active enzymes (CAZymes) are responsible for the syntheses, degradations, and modifications of cell wall polysaccharides (Lombard et al., 2014). CAZymes include glycosyl hydrolase (GH), glycosyl transferase (GT), carbohydrate esterase (CE), and polysaccharide lyase (PL) classes, carbohydrate binding modules (CBMs) and auxiliary activities (AAs). Comparing the CAZyme repertories between 16 genomes through phylogenetic analyses could reveal the evolutionary innovations of cell wall syntheses and modifications from algae to plants (Figure 3A) (e.g., (Mikkelsen et al., 2014; Mikkelsen et al., 2021)). Previous immunocytochemical and biochemical studies suggest that the later branched Streptophyta green algae (especially Zygnemophyceae) had already evolved the terrestrial pre-adapted cell walls and presented similar compositions and properties with land plants (Domozych et al.,

2014; Pfeifer et al., 2022; Sorensen et al., 2011). However, a holistic understanding of cell wall innovations has been missing.

Recent research highlighted gene gains, potentially via horizontal gene transfer (HGT), as a source for genetic innovations underpinning adaptive traits (Cheng et al., 2019; Ma et al., 2022). We hypothesized that HGT may have also contributed to the cell wall innovations. Therefore, we have focused our phylogenetic analyses on CAZymes involved in the syntheses and modifications of cellulose, mannan, beta-glucan, xyloglucan, xylan, AGP, and pectins. Our results revealed that: (i) the cell walls of embryophyte and Zygnemophyceae share all these major polysaccharide components including many of the sidechains and modifications; (ii) much of the cell wall innovations especially for the synthesis of polysaccharide backbones occurred when Klebsormidiophyceae had evolved; (iii) horizontal gene transfer (HGT) has played a major role in the origin of the enzymatic toolbox for the cell wall polysaccharide remodeling; (iv) gene loss is also very common in the cell wall gene families creating scattered distribution in sequenced Streptophyta algal genomes.

**Cellulose**: Cellulose is the major component of plant cell walls, and the most abundant biopolymer on Earth. Cellulose is widely present in various organisms including algae. Plant cellulose synthase (CesA of GT2) subunits form the hexameric rosette CesA complex, responsible for the synthesis of cellulose microfibers (multiple β-1,4-glucose chains tightly bundled together by hydrogen bonds) and support plant growth. Previous studies have suggested that the rosette CesA complex has already evolved in three classes of Streptophyta algae: Zygnemophyceae, Klebsormidiophyceae and Charophyceae (Lampugnani et al., 2019) (Figure 3A). Our recent phylogenetic analysis using Zygnemophyceae transcriptomes (Fitzek et al., 2019) suggested that the ortholog of land plant rosette CesAs first appeared in Zygnemophyceae. Now, with the assembled genomes, the phylogeny clearly (Figure 3B, Data S1-1) showed that SAG 698-1b has two CesAs: ZcCesA1 (Zci_04468) together with CesAs of all other sequenced Zygnemophyceae are the direct orthologs of land plant CesA; all Zygnemophyceae genomes also have a second copy of CesAs, e.g., ZcCesA2 (Zci_03055), which are clustered with CesAs of Klebsormidiophyceae and Charophyceae. The presence of land plant-like CesA only in Zygnemophyceae suggests that the cellulose synthase complex (CSC) in a six-subunit rosette evolved first in Zygnemophyceae. This agrees with earlier experimental observations (electron microscopy and isotope labeling) that Zygnemophyceae but not other algae have the hexameric rosette CSC (Tsekos, 1999). The two CesA clades were further clustered with a newly defined CslQ clade (Zci_13680), which is also present in Streptophyta algae.

The CslD (of GT2) clade, containing the Arabidopsis β-1,4-glucan synthase CSLD3 (Yang et al., 2020), does not have orthologs in any Streptophyta algae except for Coleochaete (Figure 3B). The phylogeny with NCBI-nr hits indicates that the CslQ clade represents the ancestor of the CesA and CslD clades, and was gained by an ancient HGT from bacteria into the common ancestor of Streptophyta algae (Figure 3D). Among the three SAG 698-1b CesA/CslQ homologs, the ZcCesA1 (Zci_04468) has much higher expression than the other two and responds to various stresses (Figure 3C).

The GH9 endo-β-1,4 glucanase KORRIGAN (KOR), a membrane-bound cellulase, has been shown to be important for cellulose synthesis or assembly (Lampugnani et al., 2019). Arabidopsis GH9 proteins have been classified into three structural subclasses according to what functional domains they have: A (cytosolic domain, transmembrane domain, and catalytic domain), B (signal peptide and catalytic domain) and C (signal peptide, catalytic domain, linker region and cellulose binding domain CBM49) (Urbanowicz et al., 2007). A has transmembrane regions so is located on the membrane, such as KOR. B and C has signal peptide so is likely secreted and degrades cellulose extracellularly. Our phylogenetic analysis with plant and algal GH9 proteins shows that there are two major clades (Data S1-2), where clade 1 contains most Arabidopsis subclass B proteins and all subclass C proteins, and clade 2 contains all subclass A proteins and the rest of subclass B proteins. Clade 1 is restricted to land plants and Zygnemophyceae, while clade 2 first appeared in Klebsormidiophyceae (Data S1-2).

SAG 698-1b has eight GH9 genes. Five of them belong to clade 1 and they all have CBM49 domains and signal peptides, and therefore belong to subclass C. Three genes belong to clade 2 (Zci_02466, Zci_12102, Zci_10931) and have much higher expression under various conditions than the five clade 1 genes. Zci_02466 has a CBM49 domain so belongs to subclass C, which might degrade cellulose extracellularly. Zci_12102 and Zci_10931 have TM region but no CBM49 so belong to subclass A (Data S1-2), which may be involved in cellulose synthesis.

**Mannan**: Mannans are widely present in the cell walls of various algae (e.g., Chlorophytes and Charophytes) and plants (Popper and Tuohy, 2010). Mannan has a β-1,4-mannose backbone, while its variant glucomannan has β-1,4-linked glucose mixed in the mannose backbone (Voiniciuc, 2022) (Data S1-3a). Galactomannan and galactoglucomannan are branched mannans with galactose sidechains. The CslA subfamily of GT2 are responsible for the synthesis of land plant mannan backbones (Voiniciuc, 2022). The land plant CslA clade is phylogenetically clustered with a Zygnemophyceae-specific clade CslL (Zci_04551 and Zci_07893) and the CslK clade (Figure 3B, Data S1-1). ClsK was initially thought to be only present in Chlorophytes (Yin et al., 2009; Yin et al., 2014) but recently found to contain sequences from both Chlorophytes and Charophytes (Mikkelsen et al., 2021). This CslK clade may be the ancestor of CslA (mannan backbone), CslL, and CslC (xyloglucan backbone), but this needs further evidence. Including NCBI-nr hits in the phylogeny suggested a very ancient origin of CslK, likely via HGT from bacteria into ancient Archaeplastida (Data S1-3b).

CslK and CslL are likely responsible for the mannan backbone synthesis in green algae (Chlorophytes and Charophytes) given their closer relationship with CslA than CslC (Data S1-1, Figure 2B). AlphaFold predicted protein structure of CslL are more similar to predicted protein structure of CslA (TM-score between Zci_07893 and AT2G35650: 0.85) than to CslC (TM-score between Zci_07893 and AT3G28180: 0.71). The predicted Zci_07893 structure was superimposed with the experimentally solved CesA of bacterial bcsA protein structure (4HG6) which contains its substrate UDP-glucose (Data S1-4). The GDP-mannose (substrate of CslA) was docked into the superimposed structures showing that GDP-mannose fits well in the binding pocket interacting with the key motifs characterized in 4HG6 interacting with UDP-glucose (substrate of CesA). The candidate mannan synthase (Zci_07893, ZcCslL1) has as a high expression comparable to ZcCesA1 (Zci_04468); the other candidate (Zci_04551, ZcCslL2) is also expressed under various treatments at a lower level (Figure 3C). This is supported by the notion that mannan is an abundant hemicellulose in algae (Voiniciuc, 2022).

MSR1 and MSR2 (GT106) are cofactors of ClsAs, and play a role in the synthesis of glucomannan. GT106 contains the pectic RG-I rhamnosyltransferases (RRTs), which share sequence homology with

MSR1 and MSR2. The close homologs of MSR1 and MSR2 are present in Zygnematophyceae and most distant ones in Klebsormidiophyceae (Data S1-5).

GMGT (MUCI10) of GT34 has been characterized as α-1,6-galactosyltransferase for the addition of galactose sidechain to mannans. GMGTs are only found in Zygnematophyceae and land plants (Data S1-6), and are clustered with xyloglucan α-1,6-xylosyltransferases (XXTs) in the GT34 phylogeny. The phylogeny revealed that GT34 homologs are also present in other Charophytes and Chlorophytes, but it was until in Zygnematophyceae that XXTs and GMGTs were separated due to gene duplications in ancient Zygnematophyceae.

For modifications, MOAT1-4 (DUF231) are mannan acetylation enzymes. The DUF231 phylogeny shows that MOAT1-4 first appeared Charophyceae (Data S1-7). The acetylation enzymes for xylans (XOATs) and pectins (PMR5) also belong to the DUF231 family. The phylogeny shows that XOATs and PMR5 are separated in land plants and their orthologs are present in Zygnematophyceae and Coleochaetophyceae.

For degradations, MAN1-3 are characterized as endo-β-1,4-mannanases (GH5_7) (Rodriguez-Gacio Mdel et al., 2012). The homologs of MAN1-3 are missing in the Zygnema genus but are present in Chlorophytes, Charophytes, and land plants (Data S1-8). This is consistent with the wide distribution of mannans in plant and green algae. A phylogeny with NCBI-nr hits suggests an ancient origin of plant and green algal mannanases from bacteria through HGT (Data S1-9). AGAL2 and AGAL3 (GH27) have an α-galactosidase activity and may act on the cleaving off galactose sidechains; their orthologs first appeared in Klebsormidiophyceae and may have gained from fungi through HGT (Data S1-10).

**Mixed linkage glucan and other beta-glucan**: MLG in monocots (grasses and cereals) consists of unbranched and interspersed chains of β-1,3;1,4-glucose polymer (Kim and Brandizzi, 2021). MLG is absent in other angiosperms but also present in ferns, brown and green algae, fungi, and bacteria (Chang et al., 2021). The CslF and CslH of GT2 are responsible for the synthesis of MLGs in monocots but are absent in Zygnematophyceae and other non-monocots. In fungi, *Aspergillus fumigatus* Tft1 (XP_748682.1) has been characterized as the enzyme for the synthesis of MLG, which is a component of fungal cell walls (Samar et al., 2015). In bacteria, *Sinorhizobium meliloti bgsBA* operon encodes a MLG synthase (*bgsA*, SM_b20391) and a putative transporter (*bgsB*) (Perez-Mendoza et al., 2015). The *bgsA* gene was shown to be phylogenetically closer to β-1,3-glucan synthase *crdS* than to bacterial CesA *bcsA/celA* (Perez-Mendoza et al., 2015).

From the phylogeny of GT2 (Figure 3B), there are three newly defined Csl clades consisting of almost exclusively Streptophyta algae. CslO (Zci_07462 and Zci_15850) and CslP (Zci_01910 and Zci_11882) are phylogenetically separated from CslA/C/K/L and CesA/CslD/CslQ. Including NCBI-nr hits found that CslO is also present in non-seed land plants such as moss, spike moss, and fern (Data S1-11). CslO is further clustered with the fungal Tft1, suggesting that CslO is likely transferred from fungi (Figure 3E, Data S1-11). CslO also contains *Physcomitrella patens* Pp3c12_24670 (PNR44324.1) (Roberts et al., 2018), which has been functionally characterized for the synthesis of a novel polysaccharide called arabinoglucan (AGlc) currently only found in *P. patens*. Therefore, it is also possible that CslO functions as AGlc synthase in Streptophyta algae.

Including NCBI-nr hits found that CslP is absent in land plant, but present in various microalgae such as Charophytes and Chlorophytes (Figure 3E, Data S1-11). Interestingly, CslP and CslO are phylogenetically next to each other, and further clustered with known CesAs from oomycetes, tunicates, amoeba, and red algae (Blanton et al., 2000; Blum et al., 2010; Matthews et al., 2010; Matthysse et al., 2004). The microalgal CslP proteins (average 1,474 aa) and the micro-eukaryotic CesAs (average 1,252 aa) are much longer than CslO proteins (average 659 aa) (Data S1-12). The closest bacterial homologs of CslP/O proteins are predominantly from Cyanobacteria and Chloroflexi, which include known Cyanobacterial CesA proteins (Nobles et al., 2001; Zhao et al., 2015) (Data S1-12). An additional clade next to Cyanobacterial CesAs in the phylogeny include CesAs from Proteobacteria, Planctomycetota, Actinobacteria. These CesAs have the hallmark c-di-GMP-binding PilZ domain of the well characterized bacterial cellulose synthase A subunit (bcsA) (Romling and Galperin, 2015) (Data S1-12). Therefore, the phylogenetic clustering with various bacterial CesAs suggests that the eukaryotic CslP/O proteins have originated via HGT from bacterial CesAs (Figure 3E, Data S1-12). After that, new functions of CslO evolved in different eukaryotic organisms (e.g., MLG synthesis in fungi and likely also in various microalgae, AGlc synthesis in mosses) via divergent evolution, while the cellulose synthesis function conserved in oomycetes, tunicates, amoeba, red algae, and possibly also in green algae with CslP. Clearly, the CslP/O-like CesAs had a distinct origin compared to the land plant CesAs, an example of convergent evolution. Among the four SAG 698-1b CslO/P proteins, Zci_01910 (CslP) shows relatively high expression and responses to different stresses (Figure 3C).

CslN (Zci_08939) also shows responses to different stresses (Figure 3C). CslN is closer to CslA/C/K/L, and uniquely found in Zygnematophyceae. Searching against NCBI-nr found no hits in any other algae and plants. The top hits are all bacterial proteins mostly from Actinobacteria, Proteobacteria, and Chloroflexi, suggesting CslN originated from bacteria via HGT (Figure 3F, Data S1-13). However, none of the close bacterial homologs have characterized functions. Interestingly, in addition to the C-terminal GT2 domain, most of these CslN-like bacterial proteins also have an N-terminal PleD region (Data S1-13). PleD contains receiver domains (RECs) that respond to phosphorylation by dimerization, and a diguanylate cyclase domain (DGC or GGDEF) that is activated by dimerization leading to the production of an important secondary messenger c-di-GMP (Jenal et al., 2017). The GT2 domain fused with PleD in these CslN-like proteins is likely a c-di-GMP effector domain. In bcsA proteins, c-di-GMP binds to their C-terminal PilZ domain to activate the N-terminal cellulose synthesis domain (Abidi et al., 2022). Unlike bcsA, these CslN-like bacterial proteins do not have the PilZ domains. However, some CslN-like proteins have genes usually found in the gene neighborhood of bcsA (e.g., bcsC, bcsQ). For example, in the *Polaromonas eurypsychrophila* genome, the CslN-like gene (GGB02139) has an upstream TRP domain-containing bcsC homolog (GGB02133) (Romling and Galperin, 2015). In addition, GT2 domain could also be found in enzymes without the PilZ domains, e.g., pgaC for the synthesis of poly-N-acetylglucosamine (PNAG), a major component of bacterial extracellular polysaccharide matrix for biofilm formation (Abidi et al., 2022). Therefore, the CslN-like bacterial proteins could function as cellulose synthases or PNAG synthases. Given that PNAG is not present in Streptophyta, we infer that CslN in Zygnematophyceae is also a cellulose synthase.

**Xyloglucan (XyG):** XyG is an important hemicellulose that contain complex sidechains (Data S1-14), whose compositions are highly variable among different land plants (Mikkelsen et al., 2021). XyG released from plants can act as soil particle aggregator, which could modify the soils suitable for early plant terrestrialization (Galloway et al., 2018). The four zygnema genomes contain the XyG backbone synthases (CslC of GT2, Data S1-1, Figure 3B), and the sidechain xylosyltransferases (XXT of GT34, Data S1-15), galactosyltransferase (XLT/MUR3 of GT47, Data S1-16, Data S1-17), galacturonosyltransferase (XUT of GT47, Data S1-17), fucosyltransferase (FUT of GT37, Data S1-18). Interestingly, in addition to the FUT clade consisting of only embryophyte and Zygnemophyceae, GT37 family is significantly expanded in Zygnemophyceae with a separate clade that only contains sequences from Zygnemophyceae. GT37 also has distant homologs in Chara and Klebsormidium, as well as in many Mortierellaceae fungi (Telagathoti et al., 2021) (Data S1-19), which are saprotrophs in the soil. This indicates that HGT between fungi and Streptophyta algae may have contributed to the fucosyltransferase origin. Notably, although CslC is also present in Chara and Klebsormidium, GT34 and GT47 are restricted in Zygnemophyceae and embryophytes.

For modifications, the XyG backbone degradation enzyme XTH (GH16_20) family is significantly expanded in Zygnemophyceae (Data S1-20), while it also has homologs in Klebsormidiophyceae and Coleochaetophyceae. XTH might have been involved in HGT between fungi and ancient Streptophyta algae (Data S1-21) (Shinohara and Nishitani, 2021). The fucosidase (AXY8, GH95), involved in the degradation of fucosylated xyloglucans and AGP (Gunl et al., 2011; Wu et al., 2010), might have gained from bacteria into Zygnemophyceae through HGT (Data S1-22). The xyloglucan β-galactosidase (BGAL, GH35, Data S1-23) appears to be present only in land plants. The α-xylosidase (AXY3, GH31, Data S1-24) seems conserved across different Streptophyta and Chlorophyta. All these agree with recent papers (Del-Bem, 2018; Mikkelsen et al., 2021) suggesting that the backbone of XyG might have originated earlier, while the full enzymatic system for XyG has evolved in Zygnemophyceae.

**Callose:** Callose is a polymer of β-1,3-glucan (Piršelová and Matušíková, 2013) (Data S1-25). It is a minor component of cell walls in most plant cells but plays a vital role in cell plate maturation and cytokinesis (Davis et al., 2020). Callose is found in Chlorophyte and Streptophyte algae (Davis et al., 2020; Scherp et al., 2001). Unlike other β-glucans, callose is not synthesized by GT2 proteins. Instead, GSL (glucan synthase-like) enzymes of GT48 are involved in the callose synthesis and deposition. The phylogeny showed that Arabidopsis GSLs form three clades, each with some homologs from Streptophyte algae (Data S1-26). GSL homologs in Chlorophytes are clustered with fungi, brown alga and Omycetes (Data S1-27). GH17 family is responsible for callose degradation. Previous phylogenetic analysis has identified three clades of embryophyte proteins within GH17 (Gaudioso-Pedraza and Benitez-Alfonso, 2014): alpha and gamma are closer and thought to be restricted in embryophytes, while beta also contain Streptophyte algae. Our phylogeny (Data S1-28) confirmed that alpha and gamma clades are separated and expanded in embryophytes, while share ancestors in Streptophyte algae but not earlier than Klebsormidiophyceae. Beta is also present in Streptophyte algae as early as Klebsormidiophyceae. The previous study (Gaudioso-Pedraza and Benitez-Alfonso, 2014) also found GH17 present in fungi and close to beta clade, suggesting possible HGT between fungi and ancient Streptophyte algae.

GH5_14 is a subfamily of GH5 (Aspeborg et al., 2012), which contain two experimentally characterized activities: exo-β-1,3-glucosidase (EC 3.2.1.58) and β-glucosidase (EC 3.2.1.21) according to http://www.cazy.org/GH5_14_characterized.html. Therefore, GH5_14could be involved in the degradation of beta-glucans such as callose, MLG, and cellulose. The rice GH5_14 protein GH5BG was involved in cell wall recycling and stress response. GH5_14 was thought to be restricted to embryophytes (Aspeborg et al., 2012), but was recently found to be prevalent in oomycetes (Liang et al., 2020). In our search against the sequenced streptophyte genomes using the GH5_14 HMM from dbCAN as query (Yin et al., 2012) (E-value < 1e-10), only Zygnemophyceae and land plants have GH5_14 hits, and Zygnemophyceae has significantly more GH5_14 hits than land plants (9 in SAG 698-1b vs. 2 in moss, Data S1-29). When extend the research to NCBI-nr (E-value < 1e-15), still no other algal hits were found. The close non-streptophyte hits are from oomycetes, fungi, amoeba, and Actinobacteria (Data S1-30). The phylogeny suggests that GH5_14 in these eukaryotes were gained from bacteria through HGT. Interestingly, most streptophyte GH5_14 proteins have a functionally unknow N-terminal fascin-like domain in addition to the GH5_14 domain (Aspeborg et al., 2012), while non-streptophyte GH5_14 homologs. The combination of Fascin and GH5_14 domains first occurred in Zygnematophyceae (Data S1-31).

**Xylan**: Xylans are the most abundant hemicellulose in vascular plants with β-1,4-xylose backbone (Data S1-32). The xylose can be decorated with glucuronic acids (GlcA) at O-2 sites to form glucuronoxylan (GX), with L-arabinose at the O-2 or O-3 sites to form arabinoxylan (AX), and with both modifications to form glucuronoarabinoxylan (GAX). Much has been known for the GTs involved in xylan synthesis (Zhong et al., 2019). GT43 and GT47 contain enzymes responsible for the xylan backbone synthesis. Specifically, IRX10 and XYS1 (IRX10L) of GT47, and IRX9, IRX9L, IRX14 and IRX14L of GT43, form a protein complex. GT47 is a large protein family in Streptophyta (Data S1-33). The GT47 phylogeny shows that IRX10 and XYS1 have orthologs in Zygnemophyceae and Klebsormidiophyceae but not in other Streptophyta algae (Data S1-34). The presence of IRX10 and XYS1 orthologs in Klebsormidiophyceae agrees with the verified β-1,4-xylan synthase activity of *Kf*XYS1 from K. flaccidum (Jensen et al., 2018). The clade of IRX10/XYS1 is phylogenetically next to a clade containing IRX7 (FRA8) and F8H (for the synthesis of xylan backbone at the reducing end), which is only present in land plants and Klebsormidiophyceae (Data S1-34).

The GT43 (IRX9, IRX9L, IRX14 and IRX14L) family (Data S1-35) is present in Zygnematophyceae, Coleochaetophyceae, Klebsormidiophyceae and Charophyceae (Taujale and Yin, 2015), but significantly expanded with a clade unique to Zygnematophyceae (Data S1-35). The phylogeny with NCBI-nr hits indicates that GT43 is also widely present in animals encoding β-1,3-glucuronyltransferase for heparan synthesis (Data S1-36). A small number of Rhodotorula fungal hits are found phylogenetically closer to Streptophyta than to animals, suggesting a possibility of HGT between fungi and Streptophyta algae.

GT8 proteins are also involved in xylan backbone (IRX8 and PARVUS) and sidechain (GUX1-5) synthesis (Data S1-37). IRX8 and PARVUS (GATL1) have orthologs in Zygnematophyceae (PARVUS absent in Zygnema, Data S1-38), Coleochaetophyceae, Klebsormidiophyceae and Charophyceae. GUX1-5, which add the GlcA sidechain, has orthologs only in Zygnematophyceae (but not in Zygnema, Data S1-38). Two GT61 proteins (XAX1 and XAT) are involved in α-1,2- and α-1,3-arabinosyl transfer onto xylan backbone to form arabinoxylan (Data S1-39a). The GT61 phylogeny reveals three major clades (Data S1-39b). The previously defined clades A and B (Anders et al., 2012) can be merged into one clade, which contain land plant XAX1 and XAT orthologs, as well as proteins from Zygnematophyceae, Coleochaetophyceae, Klebsormidiophyceae. This is confirmed by a larger phylogeny with NCBI-nr hits

(Data S1-39c). The clade C contains AT5G55500 experimentally characterized for the synthesis of N-glycans (2.4.2.38), which is clustered with characterized animal O-GlcNAcylation enzymes (2.4.2.255). There is a new clade D defined here that only contains proteins from Zygnematophyceae (Data S1-39b). However, the larger phylogeny with NCBI-nr hits shows that clade D contains numerous bacteria proteins, which have not yet been experimentally characterized (Data S1-39c).

The GlcA and xylose in xylans can be methylated or acetylated. The GlcA methyltransferase GXMT1-3 (DUF579) are present in Zygnematophyceae (only in *Spirogloea muscicola*) and Charophyceae (Data S1-40). The xylose acetyltransferase ESK1 (DUF231) is present in Coleochaetophyceae and Zygnematophyceae (Data S1-41). Additionally, the arabinose in the sidechain can be linked to coumaric and ferulic acids (phenolic phytochemicals) catalyzed by BADH (coumaric and ferulic acid transferase). The BADH (PF02458) was found to be restricted to monocot plants (Anders et al., 2012). Lastly, the deacetylation enzymes BS1 (backbone xylosyl deacetylase) and DARX1 (sidechain arabinosyl deacetylase) seem to be present in Klebsormidiophyceae, Coleochaetophyceae and Zygnematophyceae (Data S1-42).

Altogether, the phylogenetic analyses suggest that enzymes for the synthesis of xylan backbones have already evolved in Klebsormidiophyceae, while the synthesis of sidechains and modifications have evolved later. For example, GUX1-5 for the GlcA sidechain appeared in Zygnematophyceae, and XAX1 and XAT for the addition of arabinose sidechains are restricted in land plants.

**Arabinogalactan protein (AGP):** In plant cell walls, arabinogalactan-proteins (AGPs) are highly glycosylated hydroxyproline-rich glycoproteins (Data S1-43). AGPs have been shown in seed plants to play important roles in plant growth, morphogenesis, cell division, apoptosis, pattern formation, abiotic and biotic stresses, sexual reproduction, and plant-microbe interaction (Seifert and Roberts, 2007). AGPs in lower land plants (e.g., liverworts, mosses, and ferns) contain highly branched galactan and unusual monosaccharides such as acofriose (3-O-Me-Rha) (Happ and Classen, 2019). Recently, significant modifications of AGPs were found in *Spirogyra pratensis* of Zygnemophyceae to have less arabinoses but much more rhamnoses in the sidechains (Pfeifer et al., 2022), leading to the proposed presence of "rhamnogalactan-protein" or RGP.

At least five GT families are known for the synthesis of AGP backbone and sidechains (Data S1-43) (Knoch et al., 2014; Silva et al., 2020). Our phylogenetic analyses (Data S1-44) suggest that the enzymes for the synthesis of the β-1,3-galactan backbone and β-1,6-galactan sidechain of AGP are present in Zygnemophyceae, Klebsormidiophyceae, Coleochaetophyceae, and Charophyceae. These enzymes include: (1) GALT2-6 and HPGT1-3 (GT31), which add β-1,3-galactan to Hyp residues of protein backbone, (2) GALT8 and KNS4 (GT31) that elongate the β-1,3-galactan backbone, and (3) GALT31A (GT31) that elongates the β-1,6-galactan sidechain (Data S1-45). GALT29A (GT29) is also involved in elongation of β-1,6-galactan but is absent in Zygnemophyceae (Data S1-46). The β-1,6-galactan sidechains can be highly decorated (Data S1-43) with glucuronic acid (GlcA), fucose (Fuc), rhamnose (Rha), arabinose (Ara) and xylose (Xyl). The decorations vary significantly among different plants and algae. The arabinosyltransferase (RAY1, GT77, Data S1-47) and β-glucuronosyltransferases (GlcAT14A-E, GT14, Data S1-48) first appeared in Klebsormidiophyceae, but were significantly expanded and diversified in Zygnemophyceae. The α-1,2-fucosyltransferases (FUT4,6,7, GT37) first appeared and significantly expanded in Zygnemophyceae (Data S1-49). Arabinogalactan methylesterases (AGM1 and AGM2), which methylate the GlcA residues in the sidechains, do not have homologs in Zygnemophyceae except for Smu (Data S1-50). The lack of methylation of the acidic GlcA may explain the excessive mucilage in Zygnema. All these suggest that the biosynthesis machinery for AGP has already evolved in Charophyceae but undergone significant expansion in Zygnemophyceae, which explains the much higher structural complexity present in Zygnemophyceae and land plants.

Plants also encode GH enzymes to break glycosidic linkages for modifying cell wall structures. At least seven GH families are found in Zygnemophyceae for AGP modifications. The AGP β-1,3-galactan backbone can be broken by GH43A and GH43B (GH43_24, Data S1-51), which are present in Zygnemophyceae, Coleochaetophyceae, and Charophyceae. Phylogenetic analysis shows that GH43_24 evolved from bacteria through HGT (Data S1-52). For the β-1,6-galactan sidechain degradation, GH30_5 and GH35 are responsible. Interestingly, GH30_5 is only found in Zygnemophyceae, mosses, and liverworts, and absent in any other Streptophyta and algae (Data S1-53). More interestingly, including NCBI-nr hits in phylogenetic analysis shows that streptophyte and arbuscular mycorrhizas (AM) fungal (Glomeromycota) GH30_5 homologs are next to each other, and may have evolved through HGT from bacteria (Data S1-54). The beta-galactosidases (BGALs) of GH35, on the other hand, seem to be only present in land plants, while other GH35 homologs are found in streptophyte algae (but not in Zygnema) (Data S1-55).

As for sidechain decorations, α-arabinofuranosidases XYL1 and XYL4 (GH3) are found in all plants and algae including Chlorophyta (Data S1-56). The β-l-arabinopyranosidases AGALs (GH27) first appeared in Klebsormidiophyceae (Data S1-57), while the other β-l-arabinopyranosidases APSE (GH27) are found in lands plants and Mesostigmatophyceae. Including NCBI-nr hits in the phylogeny shows that HGT from bacteria is plausible for the origin of APSE in Streptophyta (Data S1-58). GH95 can specifically cleave α-1,2- linage of fucose and first appeared in Zygnemophyceae; our phylogenetic analysis shows that it was gained from bacteria through HGT (Data S1-22). GUS2 (GH79) has been demonstrated to cleave GlcA from AGP, and first appeared in Klebsormidiophyceae (Data S1-59), and seems to have an origin from bacteria via HGT (Data S1-60).

**Pectin:** Pectins include a class of the most complex cell wall polysaccharides (Atmodjo et al., 2013). Pectins play a variety of roles in plant growth, development, cell wall plasticity, response to abiotic stress, cell-cell adhesion and communication, and innate immunity. Pectins are loosely classified into three groups: homogalacturonan (HG), rhamnogalacturonan I (RG-I), and rhamnogalacturonan II (RG-II). HGs exist mostly without sidechains, but in some plants or algae, there are HGs with one single xylose or apiose sidechains, called xylogalacturonan (XGA) and apiogalacturonan (AGA).

HG is a polymer of α-1,4 galacturonic acid (GalA), and makes up 65% of pectins in plant cell walls (Data S1-61). AtGAUT1 and AtGAUT7 of GT8 form a protein complex responsible for the HG biosynthesis (Data S1-37). AtGAUT1-7 orthologs first appeared in Klebsormidiophyceae (Data S1-62), although HG was not chemically detected in Klebsormidium (Sorensen et al., 2011). AtGAUT8 (QUA1) is also involved in HG biosynthesis and has orthologs in Coleochaetophyceae and Zygnematophyceae (not in Zygnema though). GalA in HGs is often methylated or acetylated. The HG methyltransferase QUA2 is absent in Zygnema but present in other Zygnematophyceae, as well as Coleochaetophyceae and

Charophyceae (Data S1-63), while QUA3 is only found in Zygnematophyceae and land plants. CGR2 and CGR3 are also methyltransferases but their homologs are absent in Zygnematophyceae (homologs in Coleochaetophyceae, Charophyceae, and Mesostigmatophyceae, Data S1-64). The HG acetyltransferase PMR5 first appeared in Charophyceae (Data S1-65). These findings indicated that the full machinery for HG synthesis has already evolved in Charophyceae, which is consistent with the immunocytochmemical and chemical evidence by (Sorensen et al., 2011). The β-1,3-xylosyltransferase XGD1 (GT47, Data S1-33) for XGA synthesis appears to be present in Coleochaetophyceae and Charophyceae but absent in Zygnematophyceae (Data S1-66). Pectin methylesterases (PMEs) of CE8 and acetylesterases (PAEs) of CE13 can release the methyl and acetyl groups from the pectins, respectively. Homologs of PMEs have been found in Klebsormidium, and were reported to have been gained from bacteria through HGT (Ma et al., 2022). The land plant PME genes form two groups in our CE8 phylogeny (Data S1-67), and have many orthologs in streptophytes as early as Klebsormidiophyceae. However, PME is entirely absent in the genus Zygnema (Data S1-67). PAEs are also found in streptophyte algae (including Zygnema) as early as Klebsormidiophyceae (Data S1-68). The phylogeny with NCBI-nr hits shows that PAEs in plants are closer to bacteria and Chlorophytes than to streptophyte algae, suggesting ancient HGT from bacteria (Data S1-69). The PAEs from streptophyte algae are instead clustered with a group of marine invertebrates (e.g., corals, starfish, sea anemone, jellyfish, mussels) and early branching eukaryotes (e.g., Emiliania huxleyi) and haptophyte microalgae (Data S1-69).

RG-I constitutes 20–35% of pectin and has a backbone composed of repeating units of GalA and rhamnose (Rha). Its sidechains contain arabinans (α-1,5-linked) and galactans (β-1,4-linked) and/or arabinogalactans (Data S1-70). RG-I rhamnosyltransferases AtRRT1-4 and galaturonosyltransferase AtRGGAT1/MUCI70 are responsible for the synthesis of RG-I backbone (Amos et al., 2022; Takenaka et al., 2018). AtRRT belongs to a large GT106 family (Data S1-71), which also contain other characterized proteins for the mannan and arabinogalatan syntheses (Data S1-5). AtRRT orthologs are found in Zygnematophyceae, Coleochaetophyceae, and Klebsormidiophyceae (Data S1-71). AtRGGAT1 and their homologs define a new GT family GT116 (DUF616), and were phylogenetically classified into five clades (Amos et al., 2022). AtRGGAT1 belongs to clade A (Data S1-72), which contains orthologs in Zygnematophyceae, Coleochaetophyceae, Charophyceae, and Klebsormidiophyceae. Although other clades may have different functions, the GT116 family has NCBI-nr hits restricted within streptophytes and bacteria, suggesting plant GT116's origin from bacteria via HGT (Data S1-73).

The ARAD1 and ARAD2 of GT47 are involved in the synthesis of the sidechain α-1,5-arabinan of RG-I. They appeared first in Klebsormidiophyceae but absent in Zygnema (Data S1-74, S33). GALS1-3 (GT92) are responsible for the sidechain β-1,4-galactan synthesis (Ebert et al., 2018). Their orthologs first appeared in Zygnematophyceae and significantly expanded in the four Zygnema genomes (9 genes in each genome) (Data S1-75). GT92 phylogeny also revealed a separate cluster with proteins almost exclusively from Zygnematophyceae. The significantly increased GT92 family size in Zygnema may be related to the high mucilage content in Zygnematophyceae to facilitate water fixation.

TBG4/5 (GH35) specifically hydrolyzes RG-I β-galactan sidechains (Ishimaru et al., 2009) and are present only in land plants (Data S1-55). Polysaccharide lyase family 4 subfamily 2 (PL4_2) can cleave α-1,4 backbone of RG-I. PL4_2 contains seven Arabidopsis genes (AtRGL1-7). Our phylogeny shows that streptophyte PL4_2 homologs form two clades: AtRGL1-7 are in clade A, which only contain land plants and Zygnematophyceae (Zci_02033 in SAG 698-1b), and clade B contains liverworts and streptophyte algae as early as Klebsormidiophyceae (Data S1-76). Including NCBI-nr hits (Data S1-77) confirms that clade B represents the ancestral RGL clade, which originated from bacteria via HGT, and clade A remains to be present only in land plants and Zygnematophyceae.

RG-II is the most structurally complex class of pectins and has the same backbone as HG (Data S1-78). RG-II contains up to 13 different mono-sugars and more than 20 different glycosidic linkages, forming highly diverse sidechains. Enzymes for most of the glycosidic linkages in RG-II are currently unknown except for three GT families. GT8 (GAUT1 and GAUT7) has been discussed above for HG backbone synthesis (Data S1-62). GT77 (RGXT1-4) has been shown to possess α-1,3-xylosyltransferase activity, which adds xylose to fucose in the sidechain. AtRGXT1-4 have orthologs in Klebsormidiophyceae and Zygnematophyceae except for Zygnema (Data S1-79). GT37 fucosyltransferase (FUT) can synthesize α-1,2 linked Gal-Fuc disaccharide structure in RG-II sidechain, which is also present in XyG and AGP. FUT first appeared in and significantly expanded in Zygnematophyceae (Data S1-49). SIA1 and SIA2 (GT29) can transfer rare sugars Dha or Kdo onto the RG-II sidechains (Dumont et al., 2014) but their orthologs are only found in land plants (Data S1-80).

GH28 family contains 67 polygalacturonases (PGs) for pectin backbone degradation in Arabidopsis, which were classified into endo-PGs, exo-PGs, and rhamono-PGs (Park et al., 2010; Yang et al., 2018). Phylogenetic analyses in these previous papers have revealed six clades, where clade E is the only one that contain algal sequences and corresponds to rhamono-PGs. Our phylogeny (Data S1-81) suggests that GH28 in Viridiplantae should be reclassified into three classes: (i) class 1 contains not only proteins from land plants (including AT3G57790 and Zci_08289), streptophyte algae, but also chlorophyte algae, and represents the earliest GH28 class in green plants; (ii) class 2 contains most clade E proteins from land plants and streptophyte algae (Zci_09024) as early as Klebsormidiophyceae; (iii) class 3 contains proteins of other previous defined clades (A-D,F) (Park et al., 2010; Yang et al., 2018) from streptophytes as early as Charophyceae, but is absent in Zygnema. Phylogenies were also built to include NCBI-nr hits (Data S1-82). The three classes have different origins. Class 1 (Zci_08289) is also present in chlorophytes and likely evolved from bacteria via HGT. Class 2 (Zci_09024) is found in more diverse algae (e.g., diatom, chlorophytes) and bacterivorous marine flagellate, and also likely evolved from bacteria via HGT. Class 3 (ADPG1/At3g57510) is clustered with fungi, so likely evolved from fungi via HGT.

According to the CAZy database (www.cazy.org), PL1_1 and PL_12 are pectate lyases and present in plants. Three plant PL1_1 enzymes have been characterized to play important roles in fruit ripening, rhizobial infection, cell elongation and differentiation (Domingo et al., 1998; Marin-Rodriguez et al., 2003; Xie et al., 2012). Arabidopsis has 24 PL1_1 genes and 2 PL_12 genes (Data S1-83a). These 26 genes form two clades in the phylogeny (Data S1-83b), which shows that PL1_1 is first present in Chara and might be gained from bacteria via HGT (Data S1-84a). PL1_12 (At3G09540 and At3G55140) has a different origin than PL1_1 but might be also gained bacteria via HGT (Data S1-84b). There is also a different clade significantly expanded in Penium, which is also found in Coleochaetophyceae and early diverging land plants (e.g., *Magnoliopsida, Amborella, Selaginella, Bryophyta, Marchantia*). Interestingly, PLs are absent in the four Zygnema genomes.

In conclusion, pectic polysaccharides probably firstly appeared in Charophyceae and evolved into the complex structures that resemble to land plants. The gradual evolved cell wall structures paved the way from water to terrestrial environments.

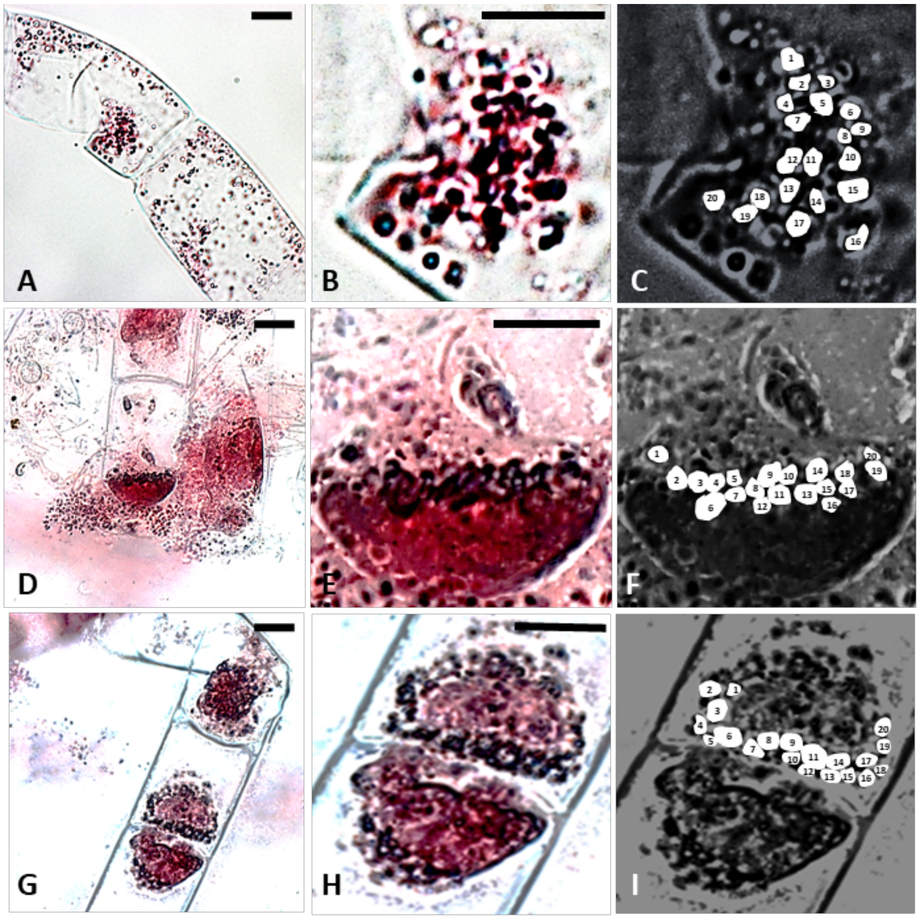


**Figure S1**. Light micrographs of *Zygnema circumcarinatum* SAG 698-1b fixed and stained with acetocarmine. (A-C) are slides without crush preparation, (D-I) are crush prepared slides. (A) overview of prophase, (B) detail from A with chromosomes visible, (C) counted chromosomes from B represented as sketches, (D) overview of metaphase, (E) detail from D with chromosomes visible, (F) counted chromosomes from E represented as sketches, (G) overview of telophase, (H) detail from G with chromosomes visible, (I) counted chromosomes from H represented as sketches. Scale bars: 10 µm.

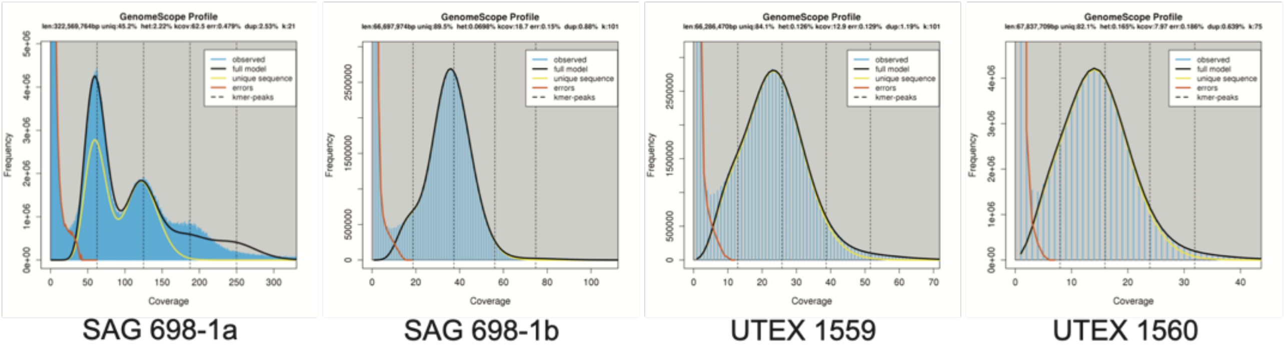


**Figure S2**. Genome size estimation of genomes of *Zygnema* cf. *cylindricum* SAG 698-1a, *Z. circumcarinatum* SAG 698-1b, UTEX 1559 and UTEX 1560 with k-mer analysis. Analysis details are described in the Methods. The input data is given in Table S1B.

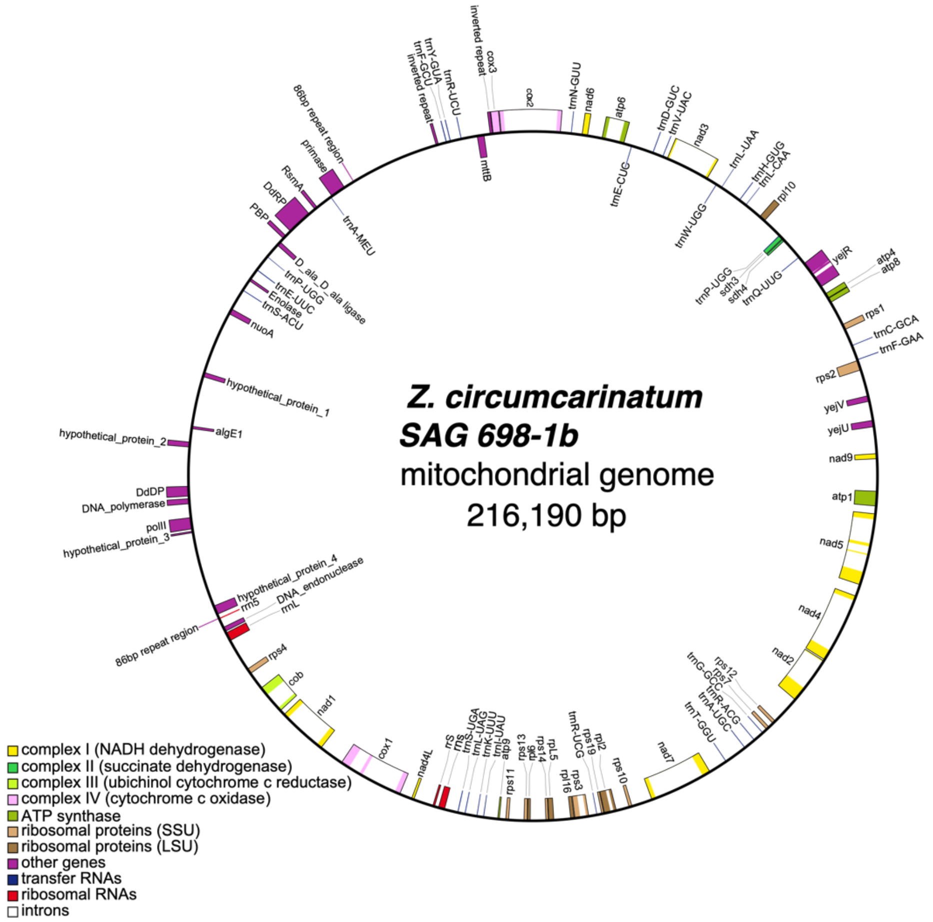


**Figure S3.** Gene annotation of *Zygnema circumcarinatum* SAG 698-1b mitogenome. Analysis details are described in the Methods. The UTEX 1560 mitogenome is identical to that of SAG 698-1b, and slightly different from that of UTEX 1559 (MT040698, 215,954 bp).

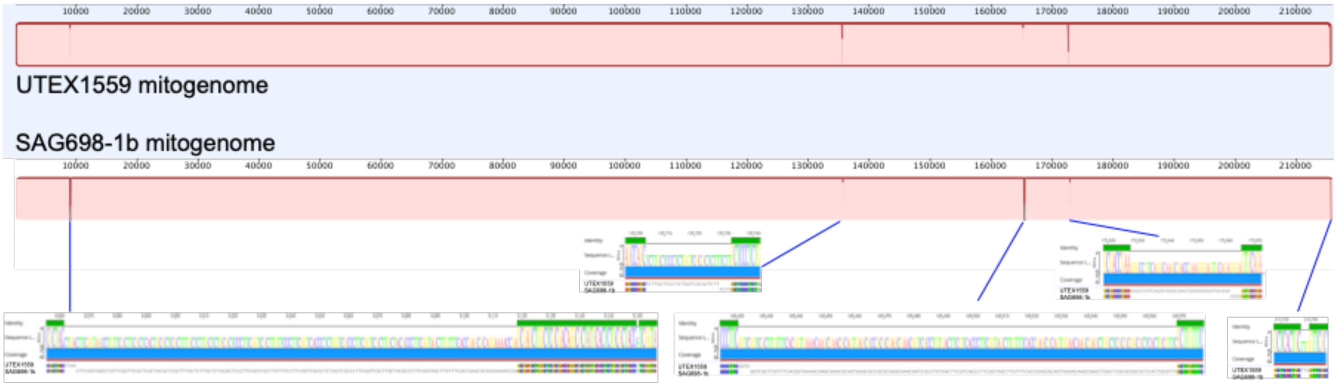


**Figure S4**. Mauve alignment of mitogenomes of UTEX 1559 (top) and SAG 698-1b (bottom). The five different regions are shown with enlarged views.

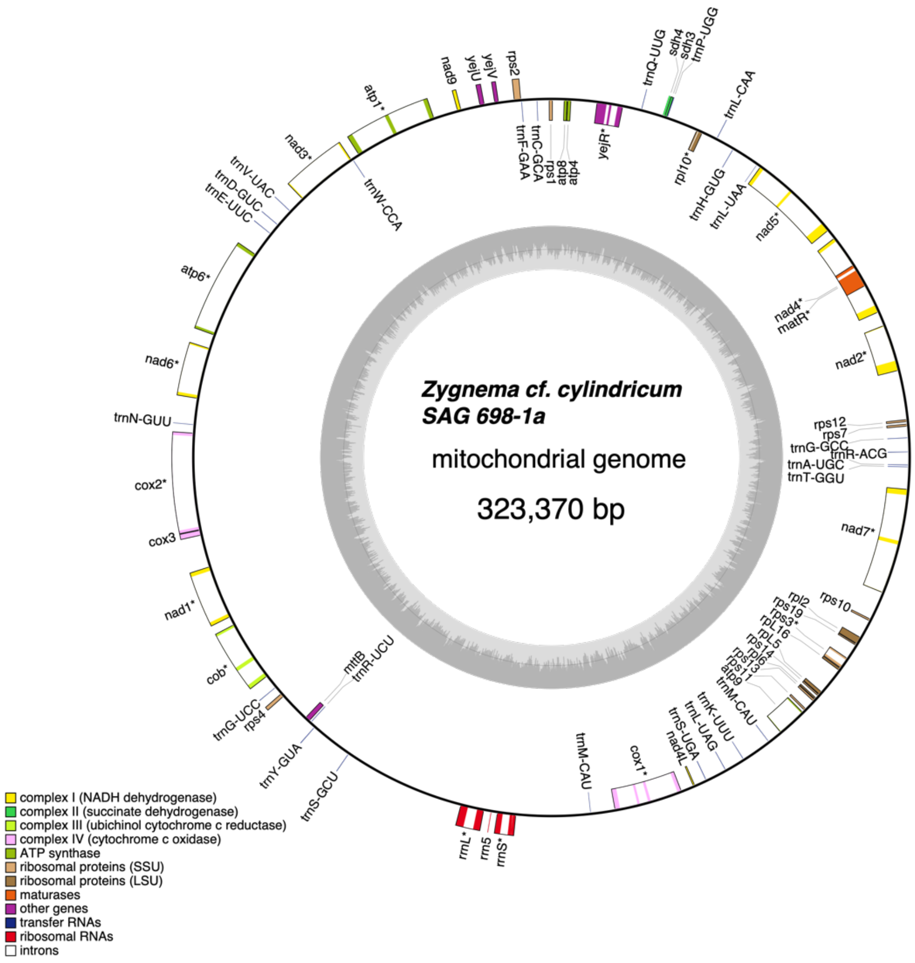


**Figure S5**. Gene annotation of *Zygnema* cf. *cylindricum* SAG 698-1a mitogenome. Analysis details are described in the Methods. The mitogenome comparison between SAG 698-1a and SAG 698-1b is given in Table S1G,H.

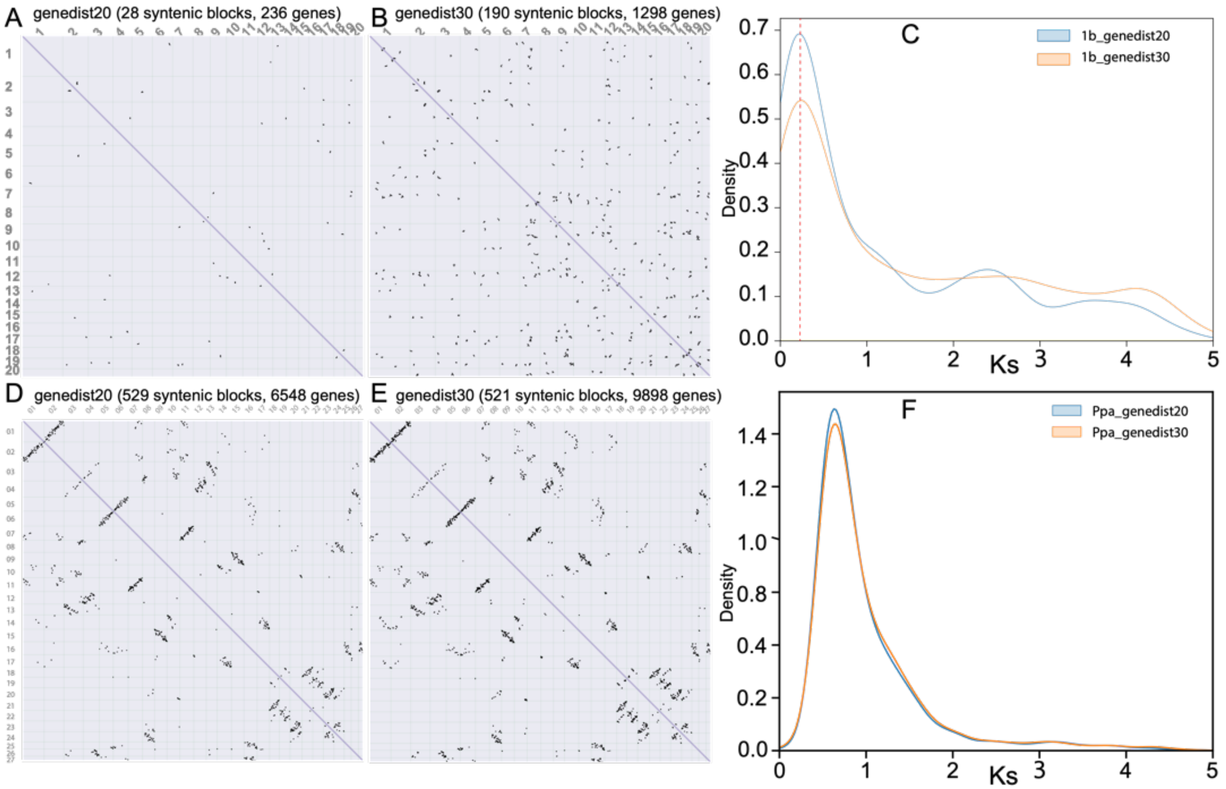


**Figure S6**. Polyploidy analysis of SAG 698-1b chromosome-level genome and comparison with *Physcomitrium patens*. (A) Dot plot of syntenic blocks in SAG 698-1b genome. The syntenic block regions were identified by MCscan with the parameter that the distance between two colinear genes within a syntenic block < 20 genes. All the paralogous genes were identified using protein RBBH (reciprocal best BLASTP hits) with E-value < 10-6. (B) Dot plot of syntenic blocks in SAG 698-1b genome. The syntenic block regions were identified by MCscan with the parameter that the distance between two colinear genes within a syntenic block < 30 genes. (C) The Ks distribution of all paralog RBBH pairs in SAG 698-1b genome. (D) Dot plot of syntenic blocks in *P. patens* genome with distance threshold < 20 genes. (E) Dot plot of syntenic blocks in P. patens genome with distance threshold < 30 genes. (F) The Ks distribution of all paralog RBBH (Reciprocal Best Blast Hit) pairs in the *P. patens* genome.

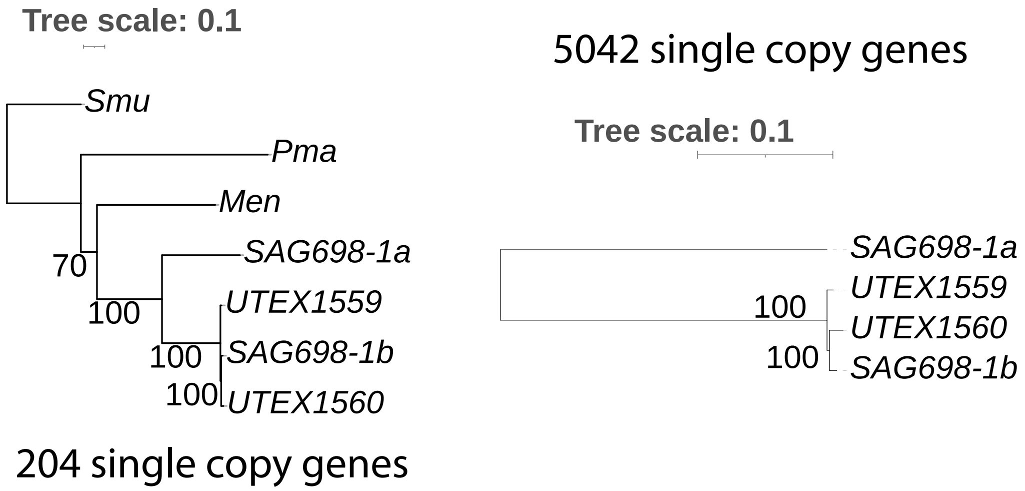


**Figure S7**. Maximum likelihood trees inferred from orthogroups obtained from Zygnematophyceae (left) and *Zygnema* (right) genomes. The numbers of single copy orthogroups used in the phylogenetic reconstructions are indicated. Analysis details are described in the Methods.

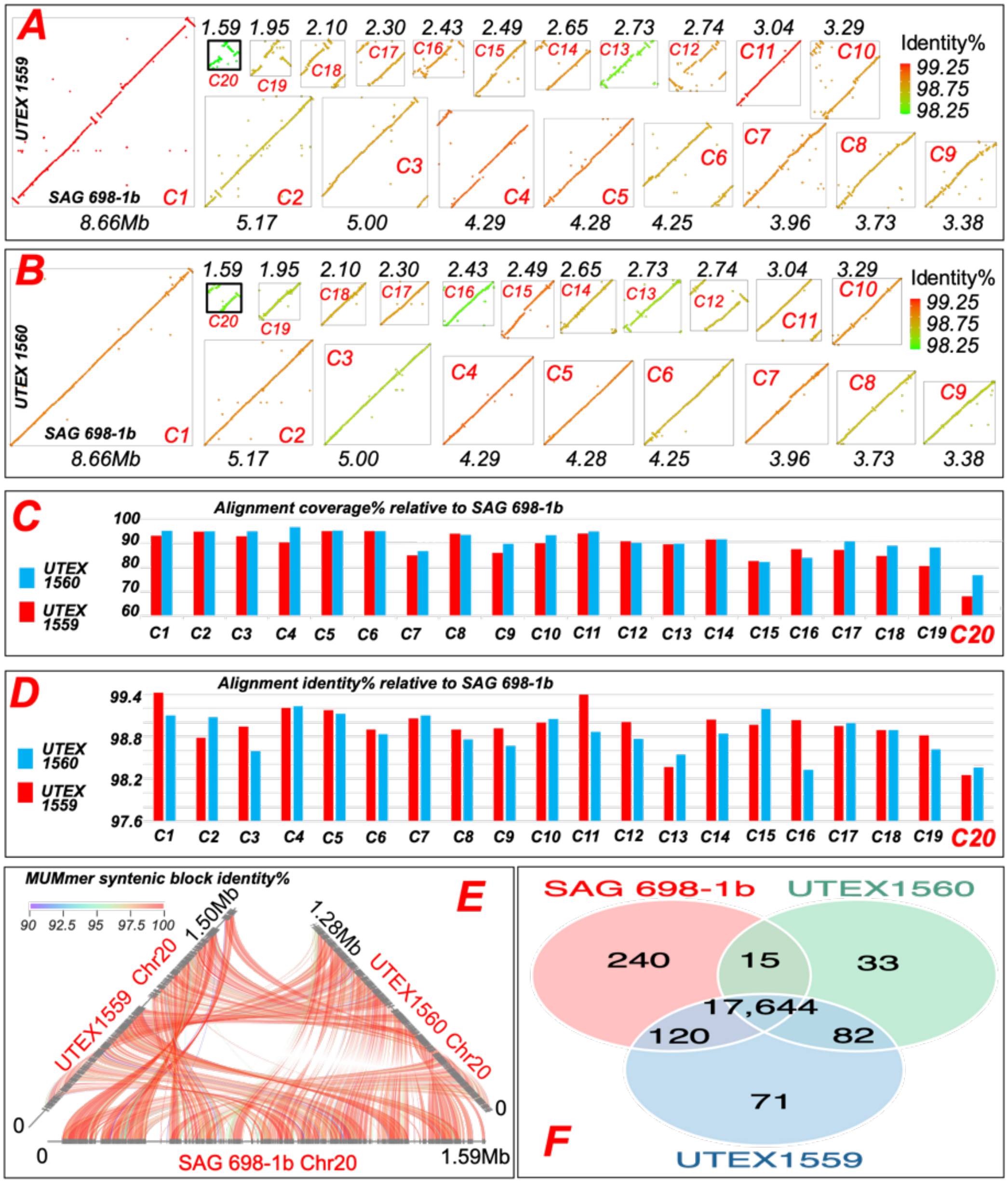


**Figure S8**. Comparisons of the three *Z. circumcarinatum* chromosome-level genomes. (A) Dot plots of NUCmer (MUMMER) identified colinear DNA blocks (size > 1000bp) between SAG 698-1b (x-axis) and UTEX 1559 (y-axis) for all the 20 chromosomes. The dots are colored based on the average NUCmer calculated nucleotide identity of all blocks in the chromosomes (Chr). Chromosome lengths are shown for SAG 698-1b (Mb). Chromosomes are drawn in proportion to their lengths. (B) Same as A but for between SAG 698-1b (x-axis) and UTEX 1560 (y-axis) comparisons. (C) Alignment coverage (the summed length of colinear DNA blocks divided by the total length of the chromosome). (D) The average alignment identity of all NUCmer aligned colinear DNA blocks of the chromosome. (E) The 3-way colinear plot of all NUCmer colinear DNA blocks of the chromosome 20 among the three genomes. (F) The venn diagram to shows the counts of core genes, shell genes, and unique genes from the gene content comparisons among the three genomes. Details are provided in the Methods.

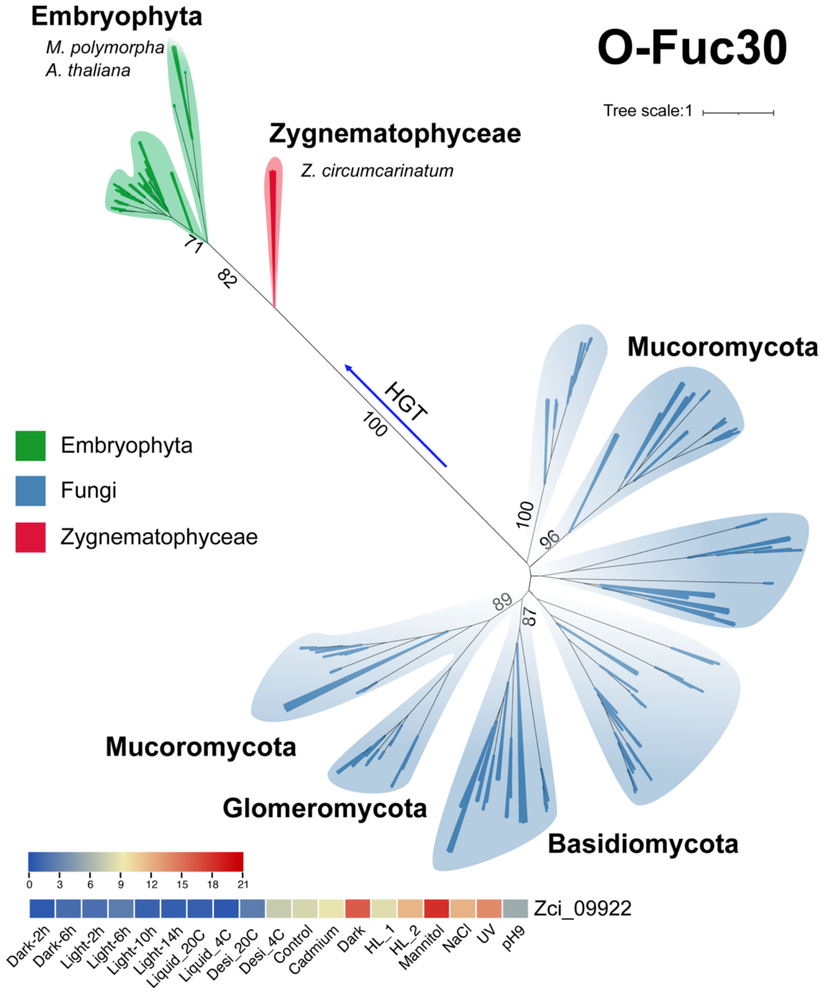


**Figure S9**. Phylogeny of the O-FucT family. The Pfam O-FucT domain (PF10250) was used to search against NCBI-nr database (E-value < 1e-10) to gather homologs, which were combined with homologs in the 16 genomes to build the tree. Zci_09922 contains a conserved O-FucT (GDP-fucose protein O-fucosyltransferase) domain (Pfam: PF10250), and was identified as a HGT candidate from fungi (Fig. 4A). The Pfam search of O-FucT found in total  Interestingly, under drought and cold conditions, the expression levels of this gene (Zci_09922) were upregulated, indicating that it was involved in response to stresses. Furthermore, our previous results showed that the fucose metabolism genes were enriched in Zygnema (Fitzek et al., 2019). In *Arabidopsis*, the O-FucT involved in cell wall integrity and cell adhesion (Verger et al., 2016). This acquired gene from fungi might facilitate the algae in adaption in extreme environments, such as drought, UV radiation.

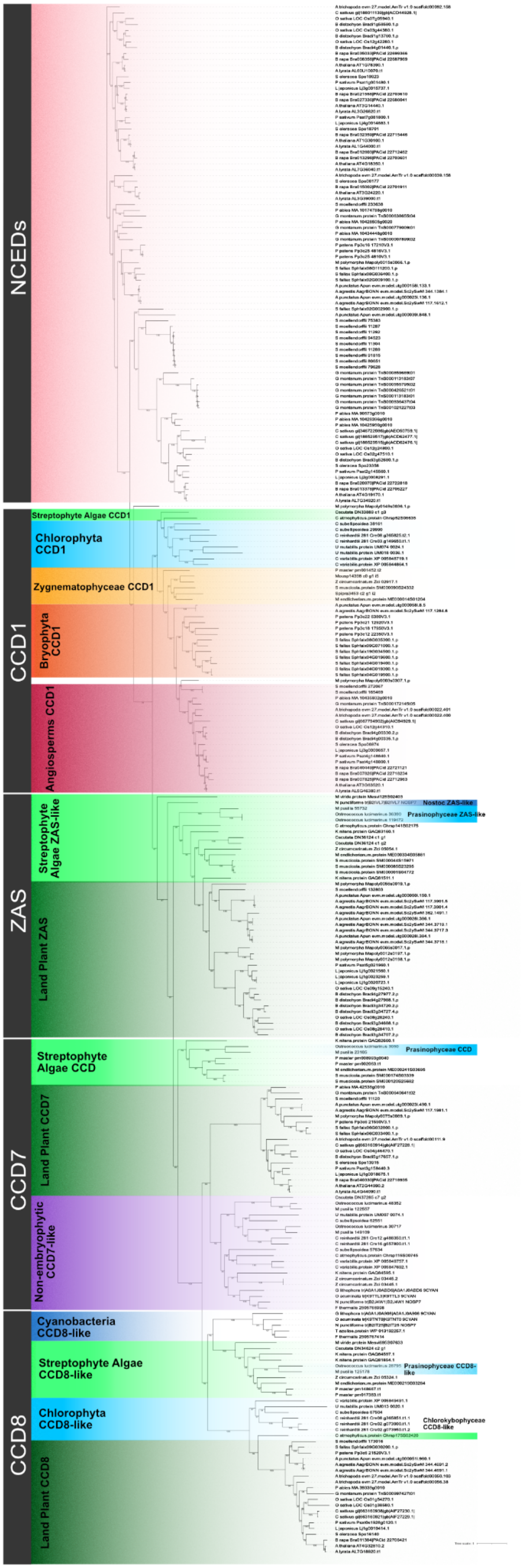


**Figure S10**. Phylogenetic tree for CCD proteins. The tree was constructed by aligning 267 CCD proteins from 41 species including *Crocus* using the MAFFT program and subjected to phylogenetic analysis by the maximum likelihood method using the IQ-TREE software. The numbers on the nodes indicate the bootstrap values after 100 replicates.

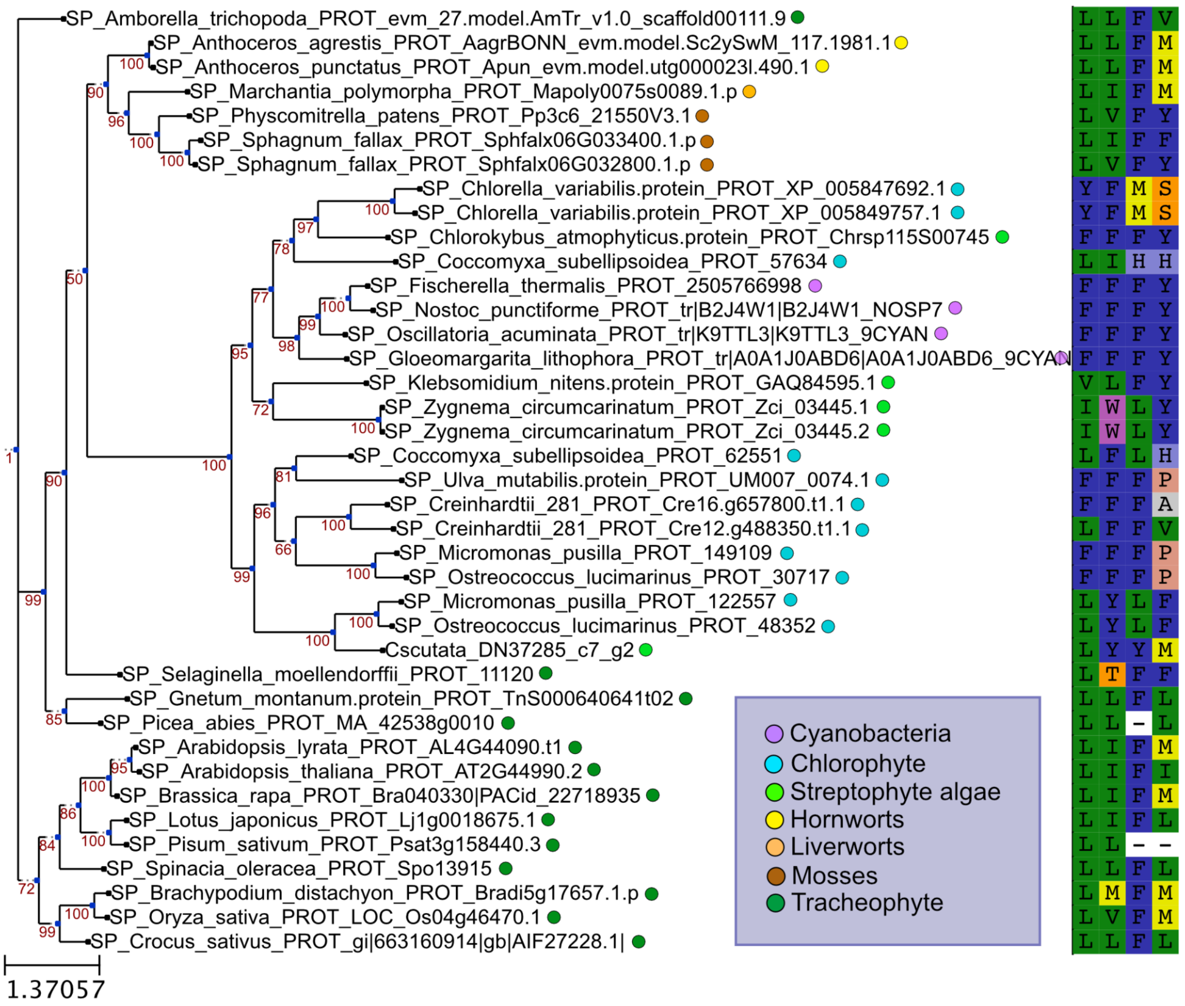


**Figure S11**. Phylogeny of CCD7 homologs. On the left, amino acids correspond to the positions of Phe-171, Phe-411, Val-478 and Phe-589 of ZmVP14. These amino acids were proposed to be crucial for substrate specificity of all CCDs (Messing et al., 2010). Dashes are gaps.

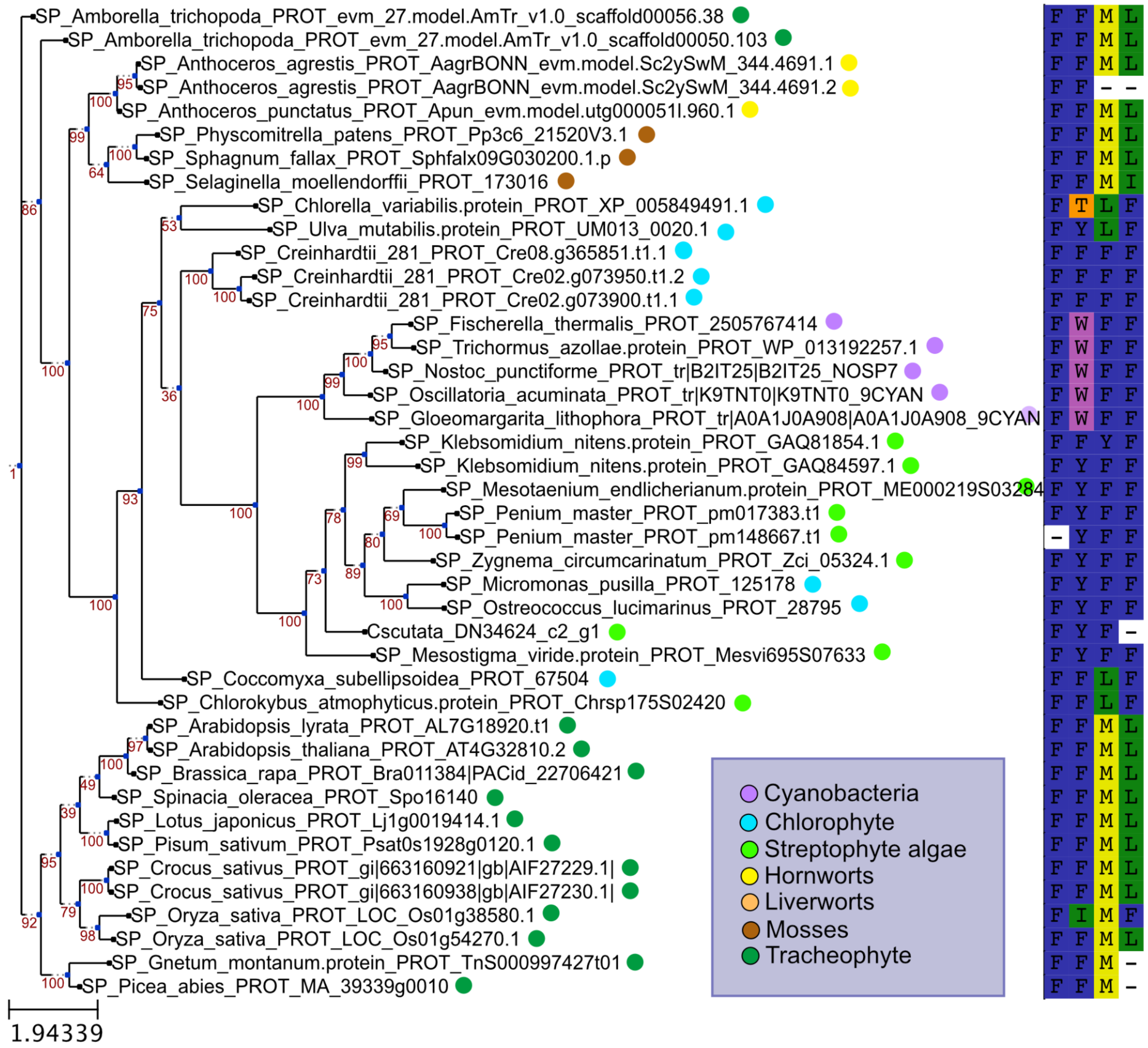


**Figure S12**. Phylogeny of CCD8 homologs. On the left, amino acids correspond to the positions of Phe-171, Phe-411, Val-478 and Phe-589 of ZmVP14. These amino acids were proposed to be crucial for substrate specificity of all CCDs (Messing et al., 2010). Dashes are gaps.

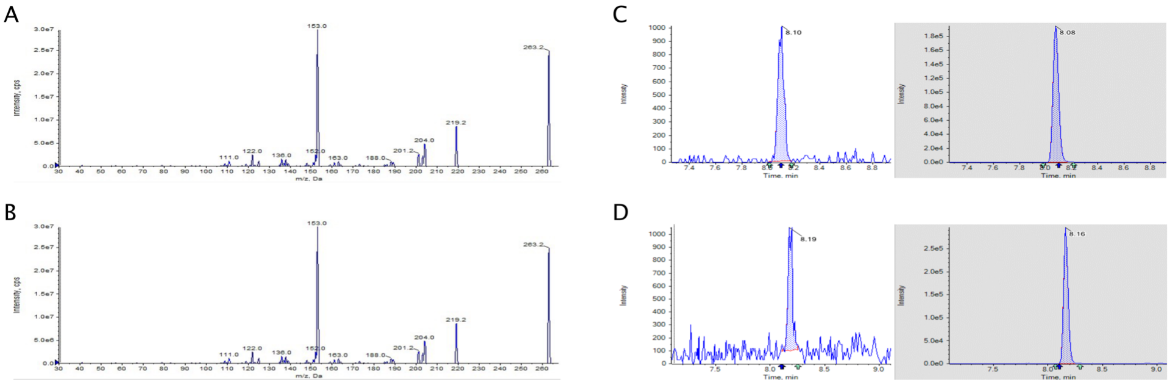


**Figure S13.** Identification of ABA in SAG 698-1b using an internal standard. Negative ESI MS/MS spectrum of abscisic acid (A) or abscisic acid-D6 (B) fragment with m/z 263.2 (C,D left) and 269.2 (C,D) respectively were used for quantification ABA and the deuterated standards. LC-MS/MS (MRM) chromatograms of 0.1 ng/ml abscisic acid (C, left) calibration standard and the internal standard (C,. right). Detection of ABA using LC-MS/MS (MRM) chromatograms of the abscisic acid (D, left) internal standard (D, right) detected in the sample.

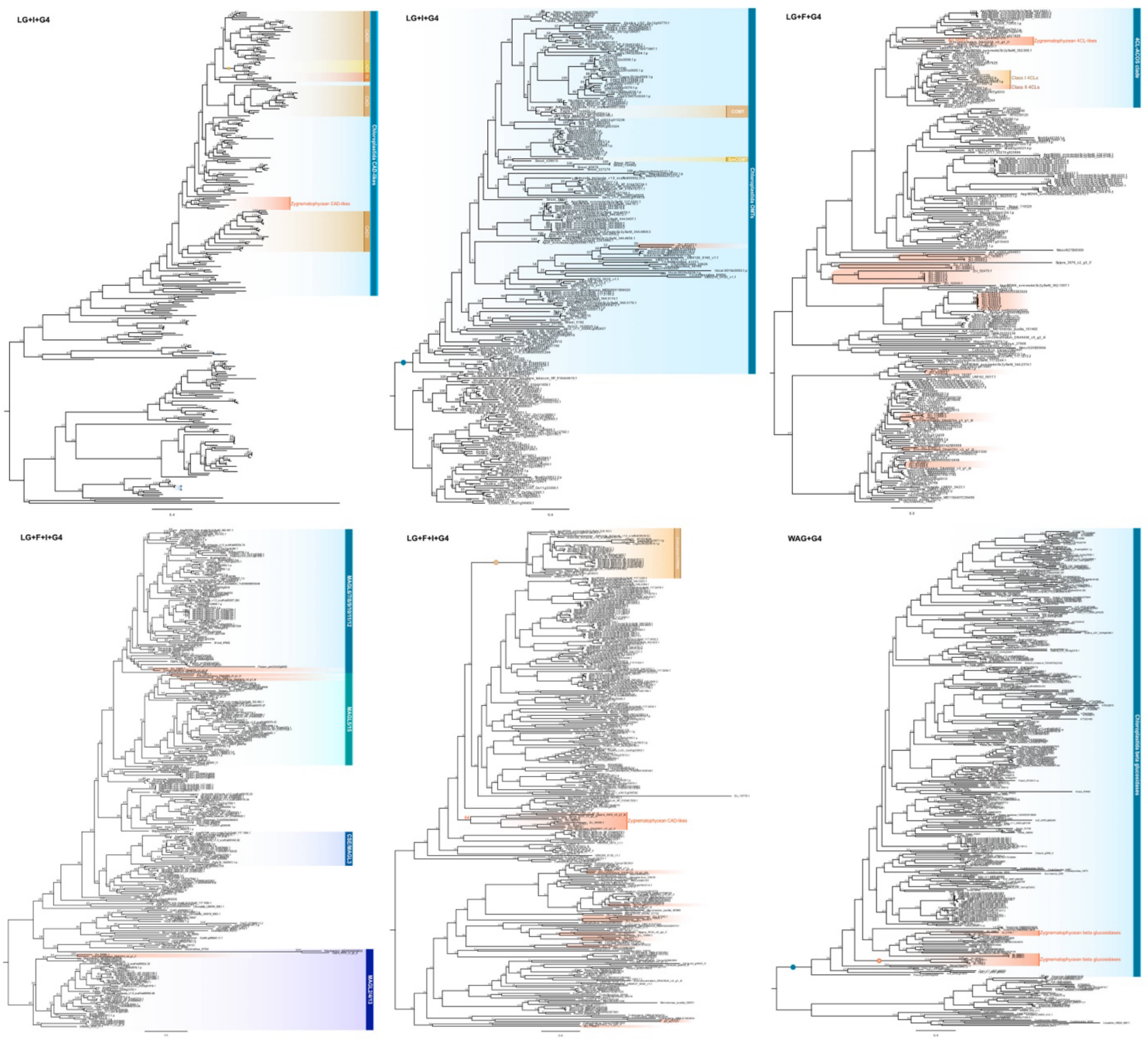


**Figure S14**. Phylogenies of genes salient to the production of phenylpropanoid-derived specialized metabolites. Best-fit models of protein evolution are noted in the top left corner.

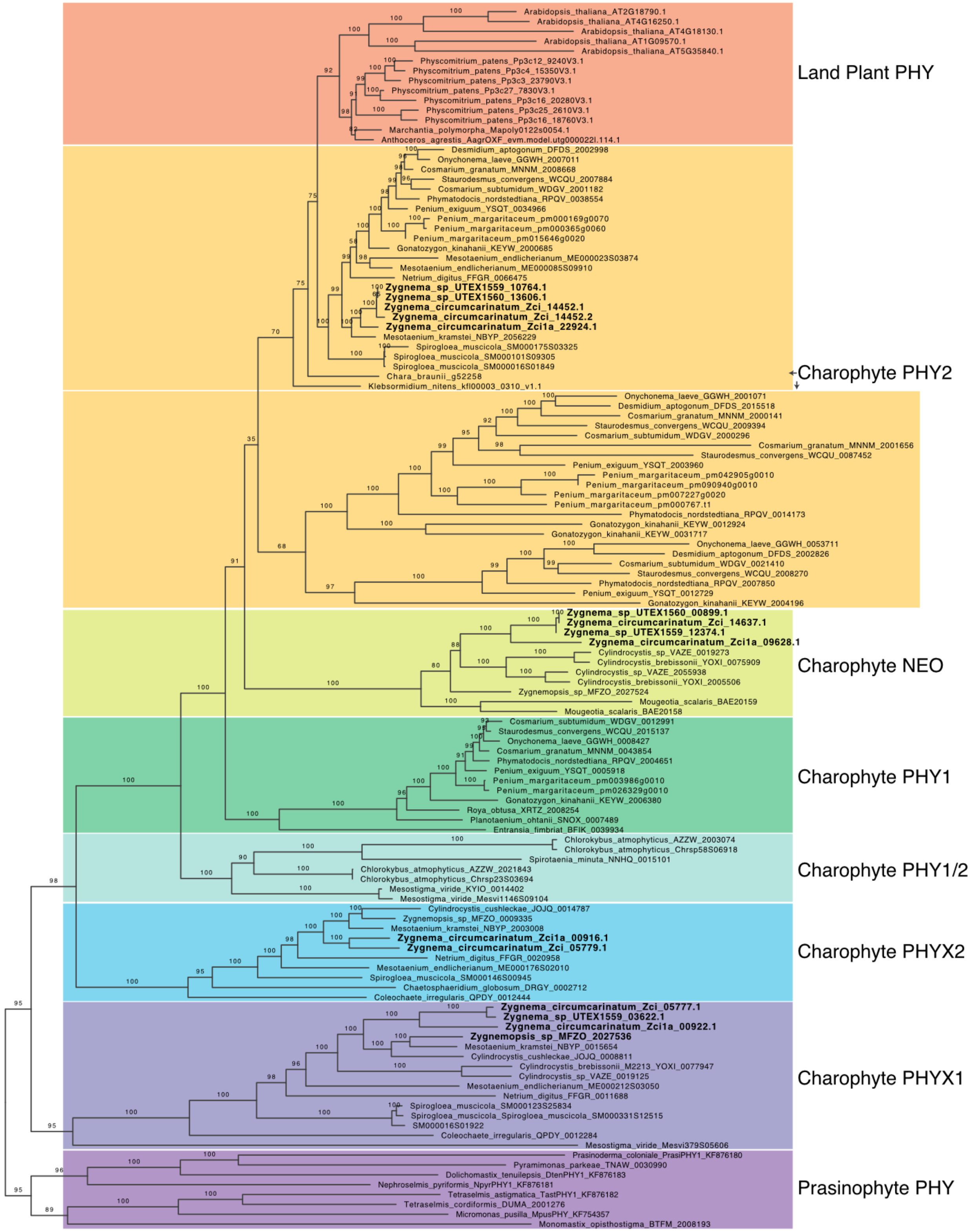


**Figure S15**. Phylogeny of phytochromes. The Zygnema sequences from this study were bolded. The dataset and clade nomenclature were derived from Li et al., 2015

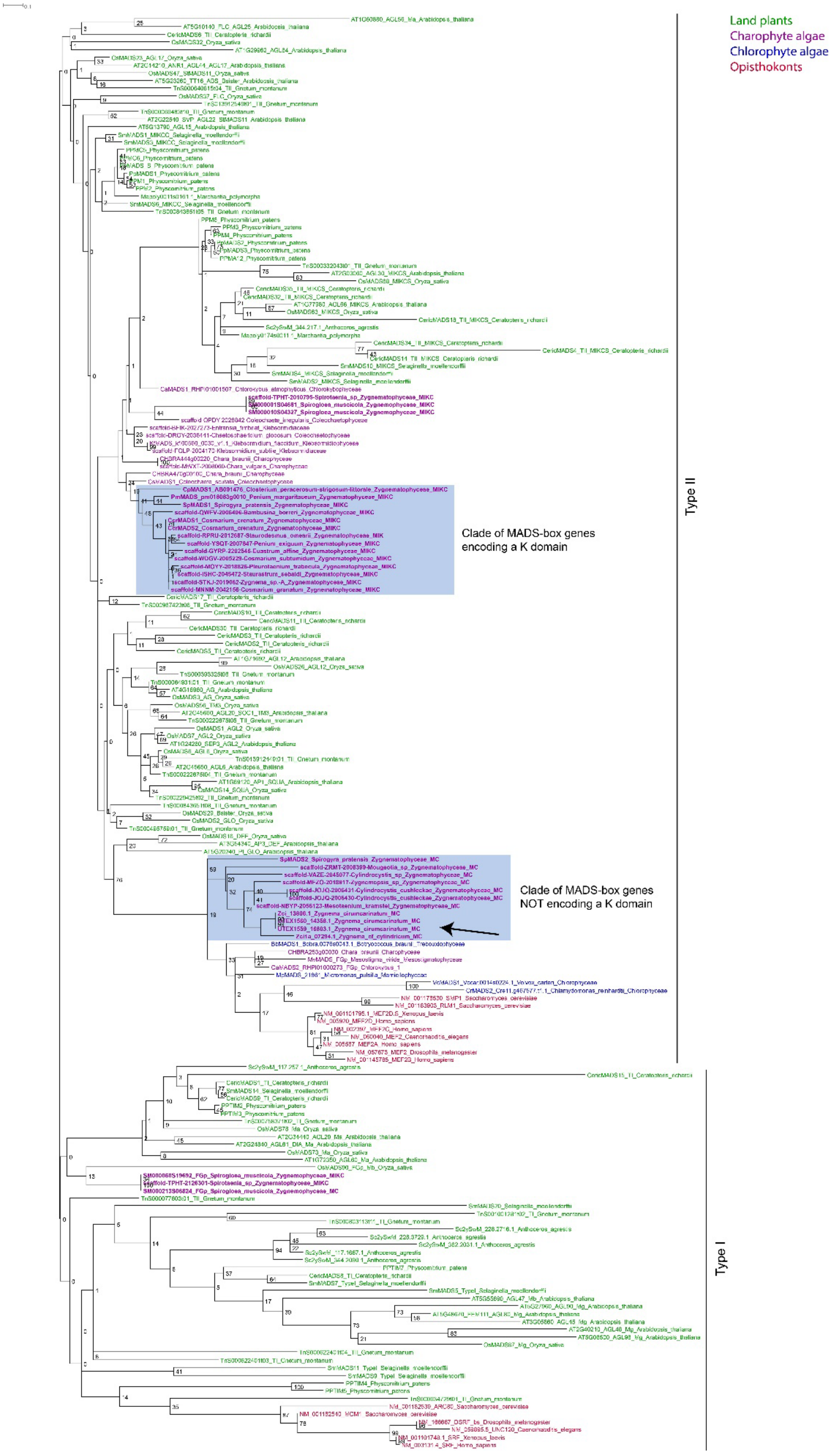

Land plants
Charophyte algae
Chlorophyte algae
Opisthokonts
Type II
Clade of MADS-box genes encoding a K domain
Clade of MADS-box genes NOT encoding a K domain
Type I

**Figure S16**. Maximum Likelihood phylogeny of MADS-domain proteins. The phylogeny was reconstructed using RAxML. Names of proteins are colored as follows: green, proteins of land plants, purple, proteins of charophytes, blue, proteins of chlorophytes, red, proteins of opisthokonts. The two major clades of Type II MADS-domain proteins in Zygnematophyceae, one comprised of proteins containing the K domain and the other including proteins without a K domain, are highlighted by shading. The positions of the MADS-domain proteins of the Zygnema genomes sequenced here is indicated by an arrow. The clades of Type I and Type II MADS-domain proteins is indicated on the right. Bootstrap values are given on the nodes.
